# Proximity-induced protein deglycosylation by endogenous O-GlcNAcase

**DOI:** 10.64898/2026.08.25.746915

**Authors:** Haixing Xu, Bowen Ma, Yunpeng Huang, Billy Wai-Lung Ng

**Affiliations:** Guangdong-Hong Kong-Macao Joint Laboratory for New Drug Screening, School of Pharmacy, The Chinese University of Hong Kong, Hong Kong, China; Li Ka Shing Institute of Health Sciences, Faculty of Medicine, The Chinese University of Hong Kong, Hong Kong, China; Gerald Choa Neuroscience Institute, The Chinese University of Hong, Kong, Hong Kong, China; Peter Hung Pain Research Institute, Faculty of Medicine, The Chinese University of Hong Kong, Hong Kong, China

**Keywords:** Bifunctional molecules, O-GlcNAcylation, Enzymes, Protein modification, inhibitors

## Abstract

O-GlcNAcylation is an important post-translational modification that regulates numerous cellular processes, yet tools enabling selective removal of O-GlcNAc from individual proteins via endogenous O-GlcNAcase (OGA) in living cells remain limited. Here, we report De-O-GlcNAcylation-targeting chimeras (DOGTACs), a chemically induced proximity strategy that selectively reduces O-GlcNAc from target proteins by recruiting endogenous OGA. Initial designs incorporating potent competitive OGA inhibitors efficiently engaged OGA but failed to induce de-O-GlcNAcylation, revealing that catalytic competence is essential for productive proximity-driven editing. By attenuating inhibitor potency while retaining sufficient OGA engagement, we developed optimized DOGTACs that promote concentration- and time-dependent, target-specific de-O-GlcNAcylation in living cells without perturbing global O-GlcNAc levels. Furthermore, we successfully applied DOGTAC to additional target proteins across multiple cell lines. Collectively, this work established attenuated competitive inhibitors as effective recruitment modules for catalytic enzyme engagement and a novel framework, DOGTAC, for targeted de-O-GlcNAcylation via endogenous OGA recruitment in living cells.

**Table of Contents:** Graphical Abstract
Bifunctional molecules recruit endogenous O-GlcNAcase to target proteins, enabling selective de-O-GlcNAcylation in living cells without perturbing global O-GlcNAc homeostasis. This work developed a programmable chemical strategy for targeted O-GlcNAc editing and framework for functional enzymatic recruitment via competitive enzyme inhibitors.

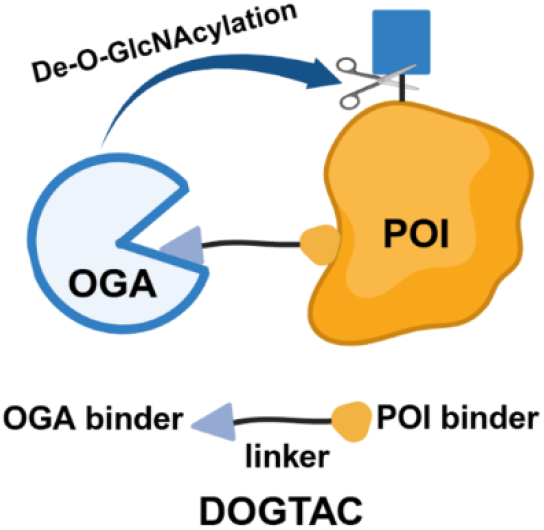

## Introduction

O-Linked β-N-acetylglucosamine (O-GlcNAc) is a ubiquitous monosaccharide post-translational modification (PTM) installed on serine and threonine residues of thousands of nucleocytoplasmic proteins in human cells.^1^ This dynamic modification, termed O-GlcNAcylation, is regulated by a single pair of enzymes: O-GlcNAc transferase (OGT), which installs the modification, and O-GlcNAcase (OGA), which removes it.^2^ Through reversible cycling, O-GlcNAcylation modulates diverse cellular processes by influencing protein localization, enzymatic activity, stability, and protein-protein interactions.^3,4^ Dysregulation of O-GlcNAc homeostasis has been implicated in numerous diseases, including neurodegeneration, diabetes, and cancers, underscoring the therapeutic relevance of precise control over O-GlcNAc modulation.^3,5–7^

Current strategies to modulate O-GlcNAcylation largely rely on genetic manipulation or small-molecule inhibitors targeting OGT or OGA. Although effective at altering cellular O-GlcNAc levels, these approaches inevitably induce global perturbations and trigger compensatory changes in OGT and OGA expression, complicating the functional interpretation of O-GlcNAcylation on individual proteins and limiting translational potential.^8–10^ To improve specificity, semisynthetic strategies developed by Pratt, Cole, and co-workers enabled site-defined O-GlcNAc installation on full-length proteins in vitro.^11–13^ In parallel, genetically encoded tools such as the light-activated Opto-OGT reported by Yang and colleagues provided spatiotemporal control of OGT activity in cells.^14^

More recently, induced-proximity approaches have emerged as powerful means to selectively manipulate O-GlcNAcylation on target proteins. Woo and co-workers employed nanobody-fused OGT or split-OGA systems to modulate O-GlcNAcylation of tagged proteins in cells.^15–17^ Zhu and Hart developed dual-specificity RNA aptamers to recruit endogenous OGT to β-catenin, resulting in selective O-GlcNAcylation.^18^ We previously also reported a chemically induced proximity strategy, termed O-GlcNAcylation-targeting chimeras (OGTACs), that selectively enhances O-GlcNAcylation of target proteins.^19–21^

Despite these advances, existing induced-proximity strategies have primarily focused on the installation of O-GlcNAc. In contrast, tools capable of selectively removing O-GlcNAcylation from specific proteins utilizing endogenous OGA remain scarce. This gap is particularly striking given the strong association of elevated OGT expression and increased O-GlcNAc levels with enhanced proliferation, survival, chemoresistance, and metastatic potential in many cancers.^22^ Targeted de-O-GlcNAcylation therefore represents an attractive yet not well-explored strategy for functional interrogation and future therapeutic exploration.

Chemically induced proximity is a powerful strategy to modulate protein functions and holds considerable promise for therapeutic development. One central challenge in this area, however, is the conversion of competitive enzyme inhibitors into effective enzyme recruiters without abolishing catalytic activity, particularly for enzymes that perform essential and nonredundant cellular functions. OGA, the only known enzyme responsible for removing O-GlcNAc modifications in human cells, exemplifies this challenge. Identifying strategies to recruit endogenous OGA while preserving its enzymatic function is therefore critical for both mechanistic studies and therapeutic modulation of O-GlcNAc signaling.

Here we report a chemically induced proximity strategy termed De-O-GlcNAcylation-Targeting Chimeras (DOGTACs) that enables selective removal of O-GlcNAc from target proteins in living cells by directly recruiting endogenous OGA (Figure 1A-B). We initially converted reported OGA inhibitors into bifunctional molecules capable of inducing proximity between OGA and a protein of interest (NUP98-HOXA9). Although these molecules successfully promoted OGA recruitment, they failed to reduce target O-GlcNAcylation, suggesting that excessive OGA inhibition compromises catalytic turnover (Figure 1C). Guided by this insight, we rationally attenuated the inhibitory potency of OGA-binding motifs to generate optimized DOGTACs that recruit OGA while preserving its enzymatic activity. These DOGTACs selectively decreased O-GlcNAcylation of NUP98-HOXA9 in a dose- and time-dependent manner, without significantly perturbing global O-GlcNAc levels in living cells or OGA activity *in vitro*. Mechanistic studies further demonstrated that DOGTAC activity depends on ternary complex formation, as co-treatment with an OGA ligand or the FKBP12^F36V^ ligand AP1867 effectively rescued the DOGTAC effect. Furthermore, we used DOGTAC to successfully reduce the O-GlcNAcylation of NUP98-KDM5A and CK2α in K562 and HEK293T cells, suggesting the potential of DOGTAC for other systems. Collectively, this work establishes DOGTAC as a chemical strategy for protein-specific de-O-GlcNAcylation in living cells and reveals an optimization strategy for catalytic proximity systems based on competitive enzymatic inhibitors.

**Figure 1.**
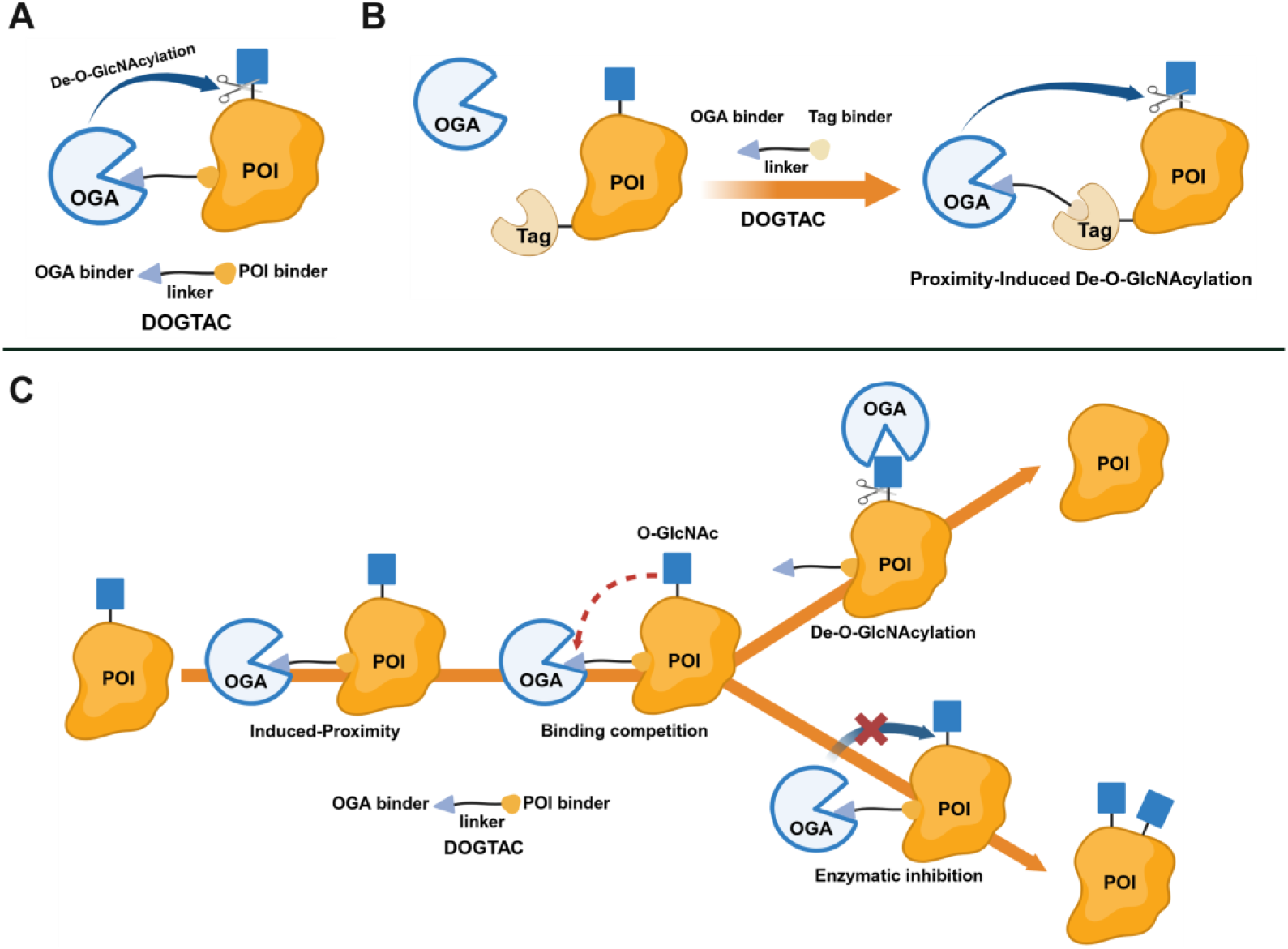
The schematic illustration of DOGTAC. (A) The design concept of the bifunctional DOGTAC to reduce O-GlcNAcylation from protein of interest (POI) using endogenous OGA. (B) Model of using Tag-fused POI for proof-of-concept of DOGTAC using OGA binder and Tag binder. (**C**) Schematic illustration of potential mechanism of DOGTAC-mediated de-O-GlcNAcylation and enzymatic inhibition.

## Results and Discussion

### Recruitment of OGA using potent OGA inhibitors is insufficient for functional de-O-GlcNAcylation

To evaluate the feasibility of DOGTAC-mediated de-O-GlcNAcylation, a highly O-GlcNAcylated protein is ideal as an initial target. Nucleoporins are among the most extensively O-GlcNAcylated proteins in cells, and nucleoporin 98 (NUP98) is a well-characterized example.^23^ NUP98 frequently forms oncogenic fusion proteins, including NUP98-HOXA9 (NHA9) and NUP98-KDM5A (NK5A), which play key roles in leukemogenesis.^24^ We therefore chose NHA9 as a model substrate for DOGTAC validation. Because no small-molecule ligand for NHA9 is available, we engineered an N-terminal FKBP12^F36V^ tag to enable chemical recruitment and appended a 2×HA tag for detection (Figure 2A). Lentiviral transduction and single-clone selection yielded a K562 cell line stably expressing FKBP12^F36V^-2HA-NHA9 (hereafter FKBP-NHA9).

**Figure 2.**
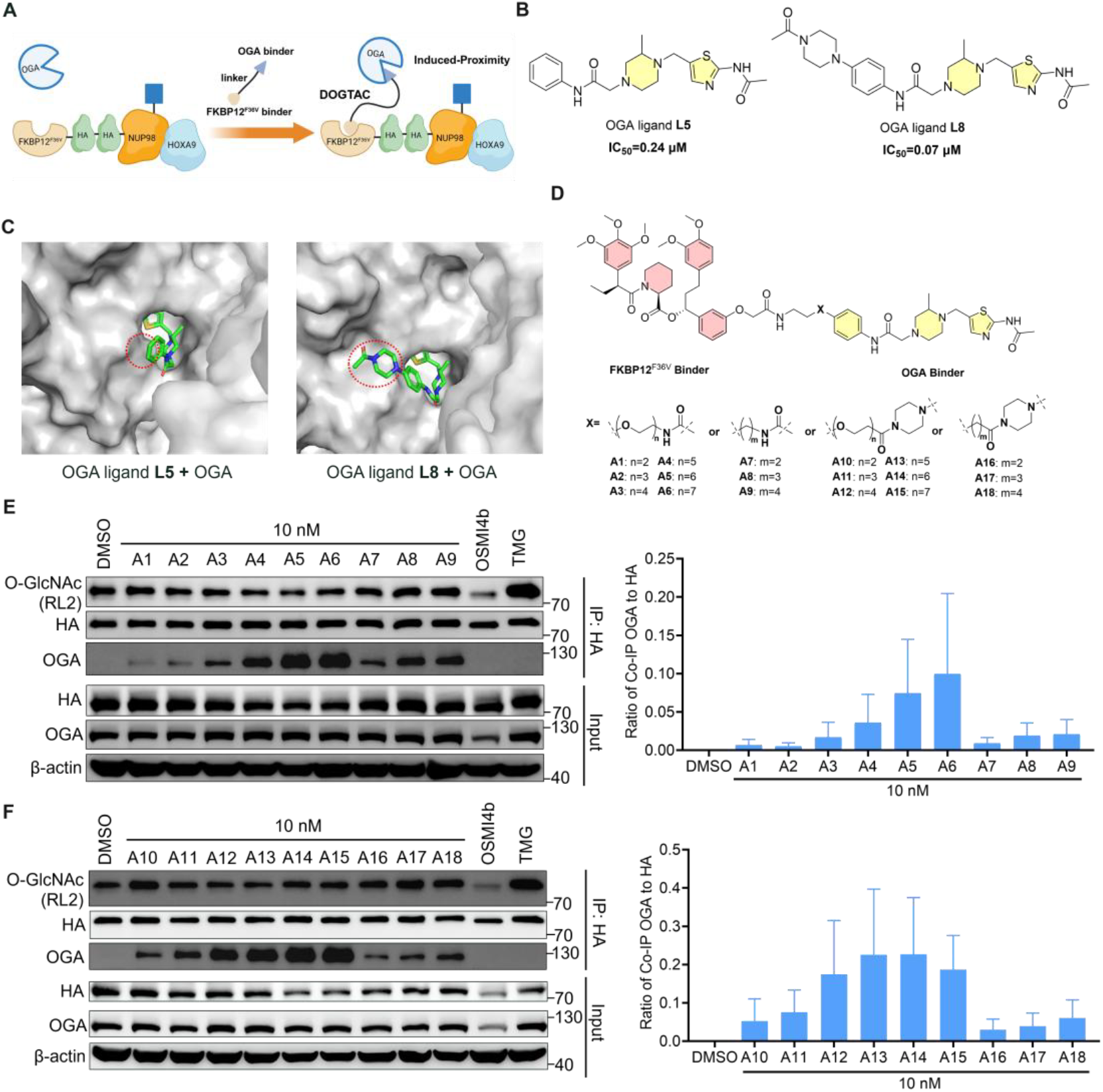
Bifunctional molecules induced proximity between OGA and FKBP-NHA9. (A) Schematic illustration of bifunctional molecules incorporating h-OGA inhibitors and FKBP12^F36V^ binder to induce the proximity between endogenous h-OGA and FKBP-NHA9. (B) Chemical structures of OGA small molecule inhibitors L5 and L8. (C) Molecular docking of OGA ligand L5 and L8 in h-OGA (gray, PDB ID: 5UHL). (D) Chemical structures of synthesized bifunctional molecules A1-A18. (E, F) Immunoprecipitation analysis to assess the effect of bifunctional molecules A1-A18 on reducing O-GlcNAcylation of FKBP-NHA9 and inducing proximity between OGA and FKBP-NHA9. FKBP-NHA9 K562 cells were treated with 10 nM A1-A18 for 24 h. 10 μM OGT inhibitor OSMI-4b and OGA inhibitor Thiamet-G (TMG) were also included as control. Then FKBP-NHA9 was enriched by HA antibody and probed for O-GlcNAc by RL2 antibody. Co-IP of OGA was detected by OGA antibody. The results in (E and F) were representative of two biologically independent replicates. Data are presented as mean ± s.e.m.

As for OGA-recruiting elements, we initially selected two reported OGA inhibitors (OGA ligands L5 and L8; Figure 2B).^25^ Structure-activity relationship analysis and molecular docking indicated that the benzyl moieties of both inhibitors are solvent-exposed (Figure 2C, Table S1), providing suitable exit vectors for linker attachment. Accordingly, we introduced a para-formyl substituent on the benzyl ring of L5 and removed the acetyl group from the piperazine of L8. These modified OGA binders were conjugated to AP1867, a high-affinity ligand for FKBP12^F36V^,^26,27^ using either polyethylene glycol (PEG) or alkyl linkers to generate a panel of bifunctional molecules (A1-A18; Figure 2D).

All compounds were screened in FKBP-NHA9 K562 cells at 10 nM to minimize OGA inhibition. FKBP-NHA9 was immunoprecipitated, and O-GlcNAc levels were assessed using the RL2 antibody, with OSMI-4b and Thiamet-G (TMG) included as OGT and OGA inhibitor controls.^9,10^ All bifunctional molecules successfully induced proximity between endogenous OGA and FKBP-NHA9, as evidenced by co-immunoprecipitation (Figure 2E-F). Compounds bearing longer linkers (e.g., A5-A6 and A14-A15) exhibited enhanced OGA recruitment (Figure 2E-F). Despite effective proximity induction, none of these compounds significantly reduced FKBP-NHA9 O-GlcNAcylation. Given the potent inhibitory activity of the parent OGA ligands (IC_50_ = 70-240 nM), we hypothesized that excessive OGA inhibition counteracted enzymatic de-O-GlcNAcylation, necessitating attenuation of inhibitor potency. Thus, enforced proximity alone was insufficient in this case: these first-generation recruiters behaved primarily as OGA inhibitors tethered to the target, which is incompatible with turnover-dependent de-O-GlcNAcylation. We therefore redesigned the OGA-binding element to reduce inhibitory potency while maintaining moderate OGA engagement.

### Optimization of OGA inhibitors to attenuate inhibitory activity

Human OGA functions as a homodimer, with substrate binding occurring in a V-shaped cleft formed between the catalytic domain of one monomer and the stalk domain of the other.^28–31^ Docking studies revealed that the N-(thiazol-2-yl)acetamide motif of OGA ligand L5 serves as the primary binding element, while the N-phenylformamide substituent linked via a 2-methylpiperazine is solvent-exposed (Figure 3A, Table S1). Guided by these insights, we pursued three complementary optimization strategies to reduce inhibitory potency while maintaining OGA binding. Strategy I exploited the stereochemistry of the piperazine methyl group. Because the R and S isomers may differentially affect OGA inhibition while preserving binding, we resolved compound A5 into its two stereoisomers, B1 (S) and B2 (R) (Figure 3B). Strategy II involved removal of the piperazine methyl group altogether. This modification yields OGA ligand L1, which displays ~10-fold weaker inhibition (IC_50_ = 2.6 μM).^25^ Using this scaffold, we synthesized a series of bifunctional molecules with extended linkers (C1–C4) (Figure 3C). Strategy III leveraged the minimal binding motif, as the N-(thiazol-2-yl)acetamide core alone retains OGA affinity even with substantial modification of the solvent-exposed region (e.g., OGA ligand L7). We therefore directly conjugated this core to AP1867 to generate bifunctional molecules D1–D6 (Figure 3D).

**Figure 3.**
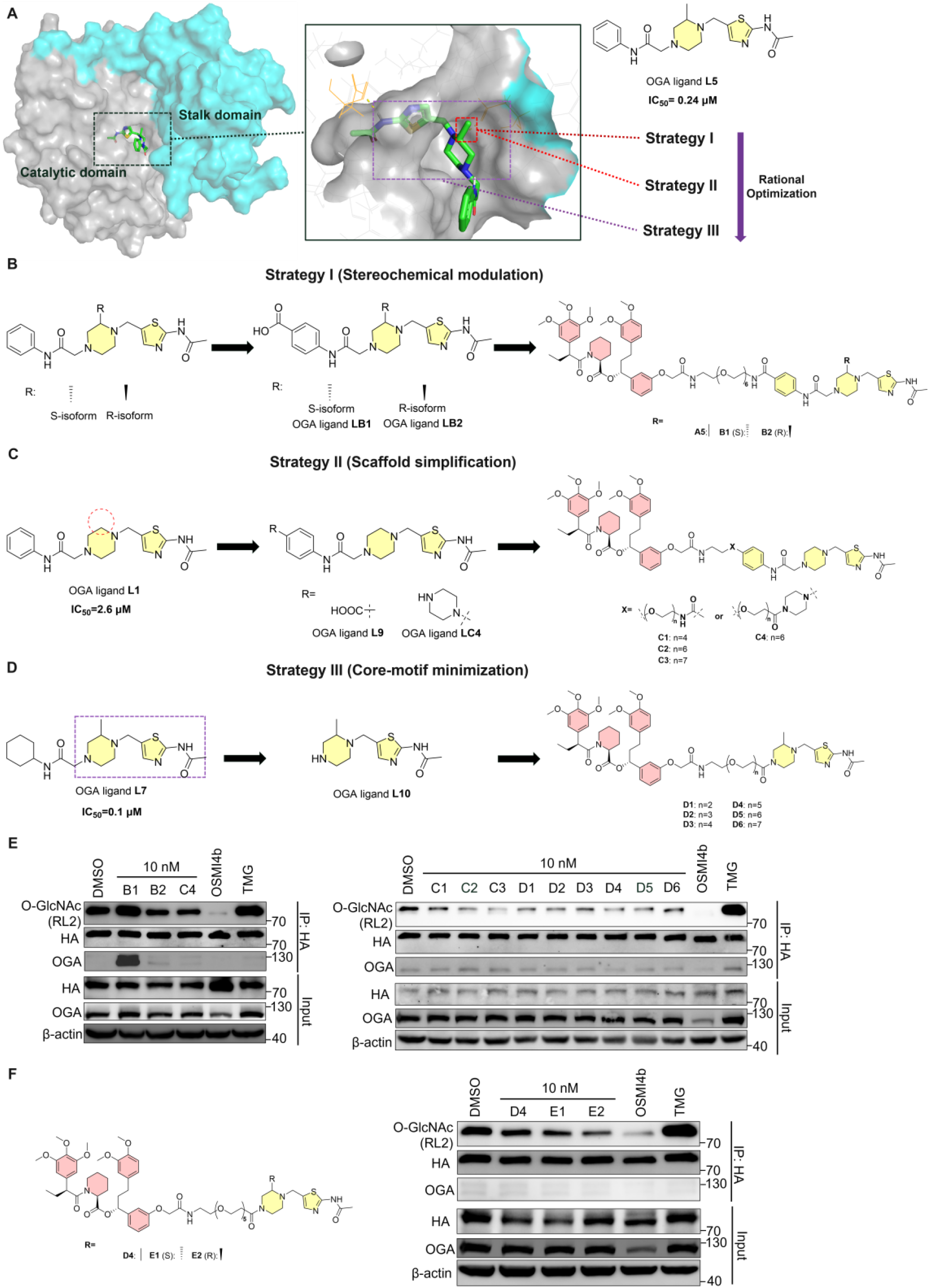
Rational optimization to generate DOGTACs reducing the O-GlcNAcylation of FKBP-NHA9. (A-D) Structure-based rational optimization and design for DOGTACs with attenuated OGA inhibitory activity. Molecular docking revealed the overview binding mode of OGA ligand L5 in h-OGA (A). Three strategies were generated step by step via rational optimization. Optimized DOGTACs were designed and synthesized via Strategy I (B), Strategy II (C) and Strategy III (D). (E) Immunoprecipitation analysis to assess the effect of DOGTACs B1-B2 and C1-C4 in reducing the O-GlcNAcylation of FKBP-NHA9. FKBP-NHA9 K562 cells were treated with 10 nM DOGTACs for 24 h. (F) The isoforms of D4 were further evaluated for DOGTACs effect. The results in (E and F) were representative of two biologically independent replicates.

### DOGTAC reduces FKBP-NHA9 O-GlcNAcylation in a concentration-and time-dependent balance between recruitment and catalysis

Screening of the optimized bifunctional molecules at 10 nM in FKBP-NHA9 K562 cells revealed that all new compounds effectively reduced target O-GlcNAcylation (Figure 3E). Within Strategy I, the R-isomer B2 exhibited stronger DOGTAC activity than the S-isomer B1, despite B1 showing superior OGA recruitment. For Strategy II, all C-series compounds were active, and C4 was selected for further study. For Strategy III, D4 emerged as the most effective compound and its stereoisomers E1 and E2 were subsequently synthesized, with the R-isomer E2 displaying superior activity (Figure 3F). Compounds B2, C4, and E2 were therefore advanced for detailed characterization.

Dose-response studies demonstrated that B2 and C4 were active at subnanomolar concentrations, with maximal effects observed between 2 and 10 nM, followed by reduced efficacy at higher concentrations (Figure 4A-B), consistent with a hook effect. Notably, OGA coimmunoprecipitation was detectable at higher concentrations (10-50 nM), indicating that proximity induction alone does not directly correlate with functional de-O-GlcNAcylation. We propose that at low concentrations, DOGTAC recruit OGA while allowing O-GlcNAcylated substrates to outcompete the ligand for the catalytic pocket, enabling turnover. At higher concentrations, persistent occupancy of the active site by the DOGTAC inhibits catalysis despite enhanced proximity (Figure 1C).

**Figure 4.**
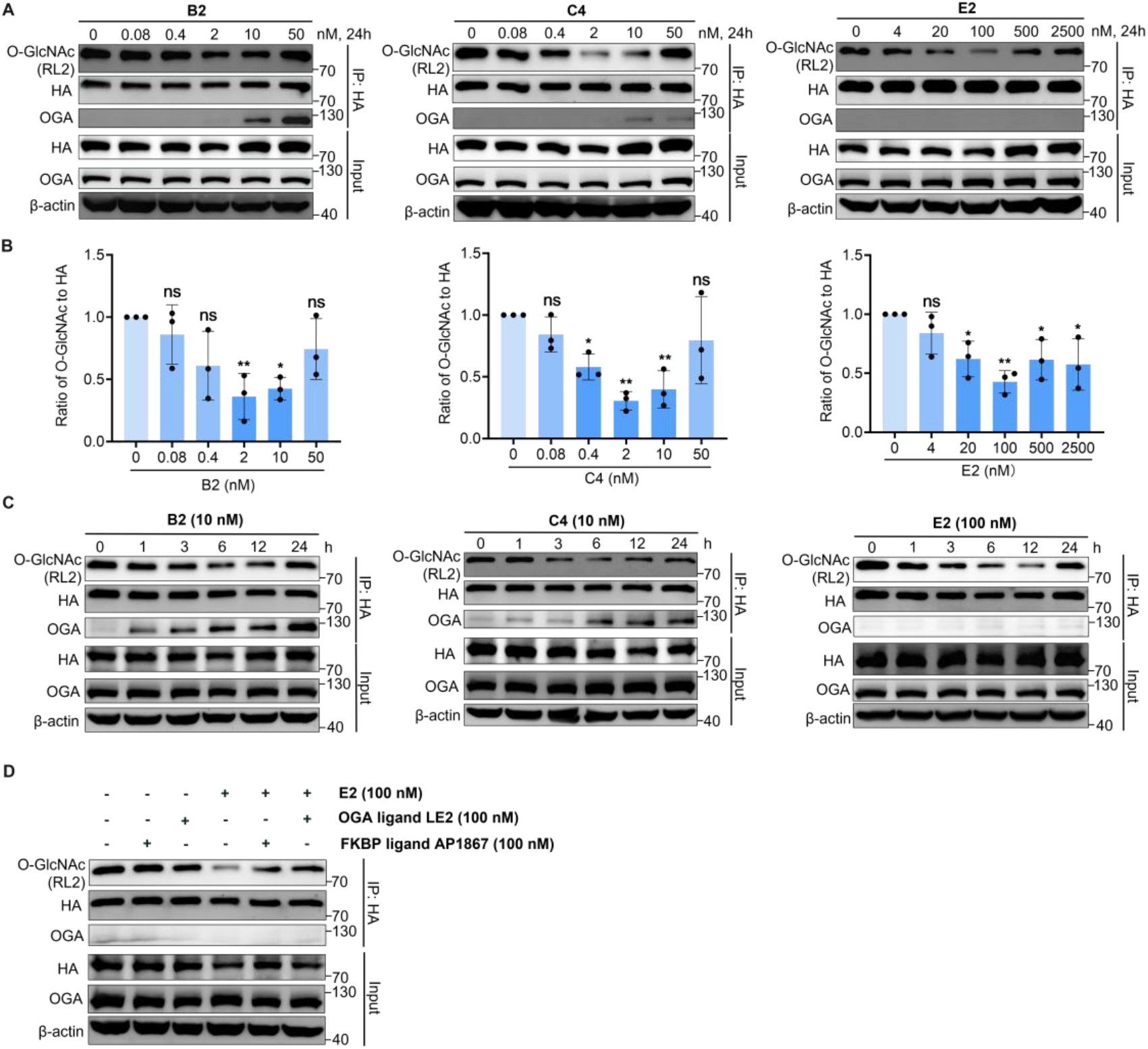
DOGTAC reduced O-GlcNAcylation of FKBP-NHA9 in a dose- and time-dependent manner. (A-B) Concentration-dependent de-O-GlcNAcylation of FKBP-NHA9 by DOGTACs in K562 cells. K562 cells were treated with increasing concentrations of B2, C4, and E2 for 24 h. Then FKBP-NHA9 was enriched by HA antibody and probed for O-GlcNAc by RL2 antibody. (B) showed the quantifications of the immunoblots signal of RL2 relative to FKBP-2HA-NHA9 and normalized to DMSO treatment as the mean ± s.e.m. of n = 3 biologically independent experiments. Statistical significance was calculated with ordinary one-way ANOVA using GraphPad Prism. Differences were considered statistically significant at p < 0.05. ns, *, **, ***: no significance, p < 0.05, p < 0.01, p < 0.001. (C) Time-dependent de-O-GlcNAcylation of FKBP-NHA9 by DOGTAC in K562 cells. K562 cells were treated with B2 (10 nM), C4 (10 nM), and E2 (100 nM) for 0, 3, 6, 12, 24 h. (D) Immunoprecipitation analysis of O-GlcNAcylation level of FKBP-NHA9. K562 cells were treated with the parent OGA ligand (LE2, 100 nM), FKBP12^F36V^ ligand (AP1867, 100 nM) as control and co-treated with E2 (100 nM) and DMSO, LE2 (100 nM), or AP1867 (100 nM) for 12 h which block E2-mediated de-O-GlcNAcylation.

Consistent with this model, E2, derived from a weaker OGA binder LE2, exhibited a shifted activity window, with optimal effects observed at 100-500 nM and no detectable OGA coimmunoprecipitation even at 2.5 μM, resulting in a broader effective concentration range.

Time-course experiments revealed rapid DOGTAC activity, with a reduction in FKBP-NHA9 O-GlcNAcylation observed within 1-3 h for all three compounds (Figure 4C). DOGTACs B2, C4, and E2 exhibited sustained activity, with maximal effects observed between 6 and 12 h. Notably, prolonged treatment with B2 and C4 resulted in increased OGA recruitment over time, which did not correlate with the extent of de-O-GlcNAcylation. To further examine this behavior, we performed additional time-course experiments with B2 and C4 at 2 nM. Similar trends were observed: both compounds were active as early as 1-3 h, with optimal de-O-GlcNAcylation occurring between 3-6 h for B2 and 6-12 h for C4, while OGA recruitment continued to increase over time (Figure S1A-B). This divergence suggests that DOGTAC-mediated de-O-GlcNAcylation is a dynamic process in which enzyme recruitment alone is insufficient to sustain catalytic turnover. These kinetic features are consistent with the highly dynamic cycling of O-GlcNAcylation governed by the opposing activities of OGT and OGA.

### DOGTAC activity requires ternary complex formation

For B2 and C4, they could recruit OGA to targeted protein observed via Co-IP experiments at a dose- and time-dependent manner (Figure 4A and 4C). However, these results were not observed for E2. We hypothesized that the weak inhibition effect of E2 also resulted in weak OGA recruitment ability. Although endogenous OGA recruitment by E2 was below the detection limit of Co-IP under these conditions, co-expression of myc-OGA enabled detection of E2-dependent OGA recruitment to the target protein, supporting target engagement by the attenuated recruiter (Figure S2).

To probe the mechanism of action, we also performed competition experiments using E2. Cotreatment with either an OGA ligand LE2 or the FKBP12^F36V^ ligand AP1867 significantly attenuated the DOGTACs effect, whereas neither competitor alone affected FKBP-NHA9 O-GlcNAcylation (Figure 4D). These results support a ternary-complex-dependent mechanism requiring simultaneous engagement of OGA and the target protein.

### DOGTAC does not perturb Global O-GlcNAcylation

To assess potential global OGA inhibition, we determined the IC_50_ values of DOGTACs and their corresponding OGA ligands using an *in vitro* enzymatic assay (Figure 5A). E2 and its parent ligand LE2 exhibited IC_50_ values greater than 10 μM, indicating minimal inhibition of OGA. B2 and its ligand LB2 displayed weak OGA inhibition (IC_50_ = 12.8 ± 5.2 μM and over 20 μM, respectively). In contrast, C4 and its ligand LC4 showed moderate OGA inhibition, with IC_50_ of 0.83 ± 0.09 μM and 1.66 ± 0.18 μM, respectively; these values remain substantially higher than their effective cellular concentrations.

**Figure 5.**
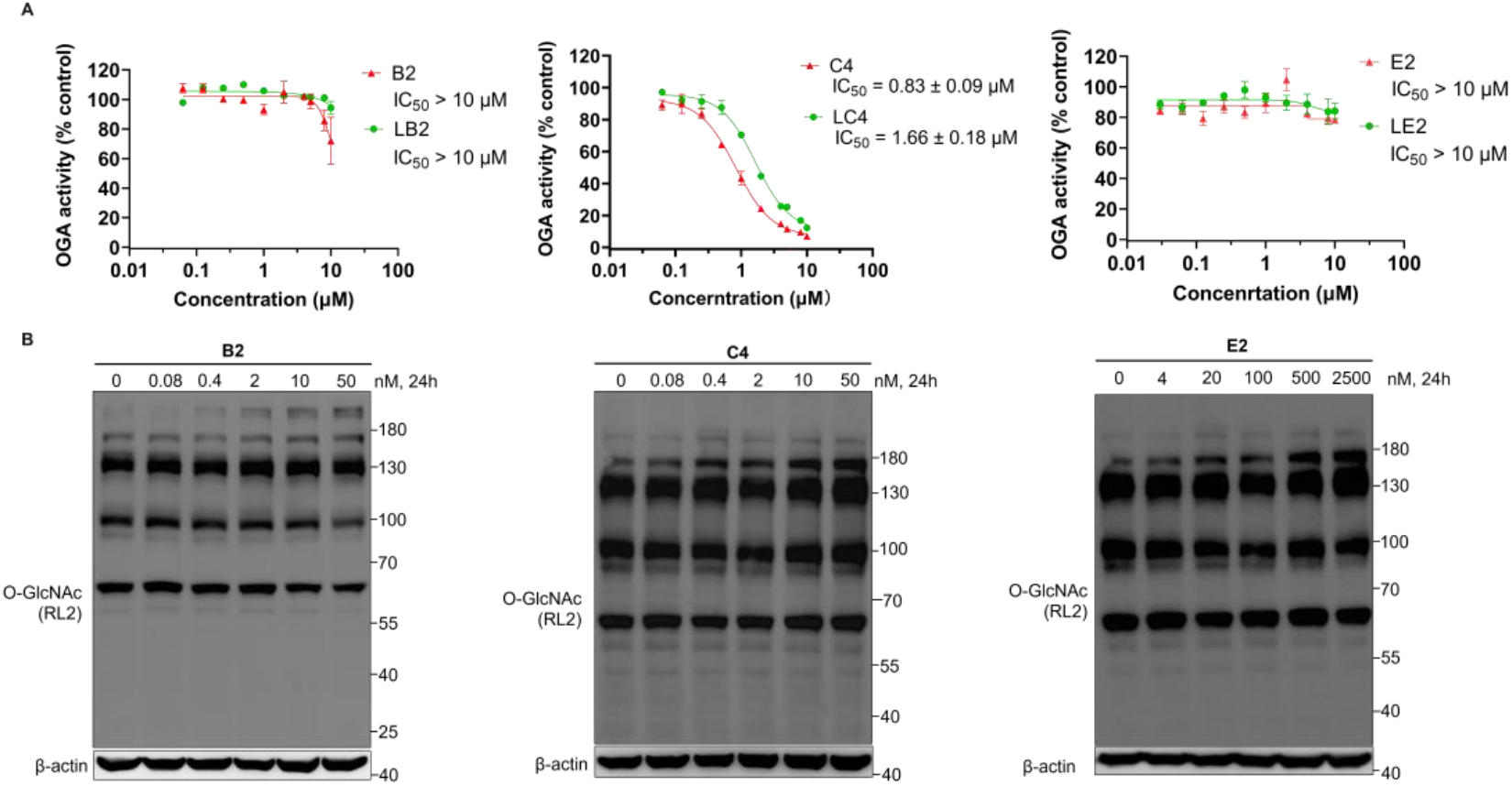
The OGA inhibitory activity of DOGTACs. (A) The OGA inhibitory activity of DOGTACs (B2, C4, and E2) and their parent OGA ligands (LB2, LC4, and LE2) *in vitro*. (B) Effects of DOGTACs B2, C4, and E2 on OGA activity in K562 cells. Immunoblot analysis of the global O-GlcNAcylation level in lysates from K562 cells treated with DOGTACs at the indicated concentrations. The results in (B) were representative of three biologically independent replicates.

We further evaluated structural isoforms of B2 and E2 along with their parent ligands (Figure S3). The S-isoform B1 and its parent ligand LB1 exhibited substantially stronger inhibition (IC_50_ of 128.7 ± 8.9 nM and 344.5 ± 45.6 nM, respectively). Notably, both B1 and B2 displayed enhanced inhibitory potency relative to their corresponding parent ligands, suggesting that the bifunctional scaffold engages additional interactions within or adjacent to the OGA binding pocket, consistent with available binding space near the catalytic site. Similarly, the S-isoform E1 and its parent ligand LE1 also showed stronger inhibitory effect (IC_50_ of 39.50 ± 1.60 μM and 39.67 ± 1.60 μM, respectively), compared to E2 and LE2.

Consistent with these findings, treatment of K562 cells with B2, C4, or E2 at their working concentrations did not significantly alter global O-GlcNAc levels after 24 h, as assessed by RL2 immunoblotting (Figure 5B).

### DOGTAC is applicable to multiple FKBP-tagged proteins across cell lines

Encouraged by these results, we next evaluated whether the de-O-GlcNAcylation activity of DOGTAC is generalizable beyond NHA9 and K562 cells. We focused on E2 and first asked whether its activity depends on the identity of the target protein. To this end, we constructed FKBP12^F36V^-2HA-NUP98-KDM5A (FKBP-NK5A), another oncogenic NUP98 fusion protein, and generated K562 cell lines stably expressing this construct. Treatment with E2 resulted in a dose-dependent reduction of O-GlcNAcylation on FKBP-NK5A after 8 h. Notably, compared with FKBP-NHA9, E2 exhibited a broader effective concentration window for FKBP-NK5A, with an optimal concentration of approximately 500 nM, slightly higher than that required for FKBP-NHA9 (Figure 6A).

**Figure 6.**
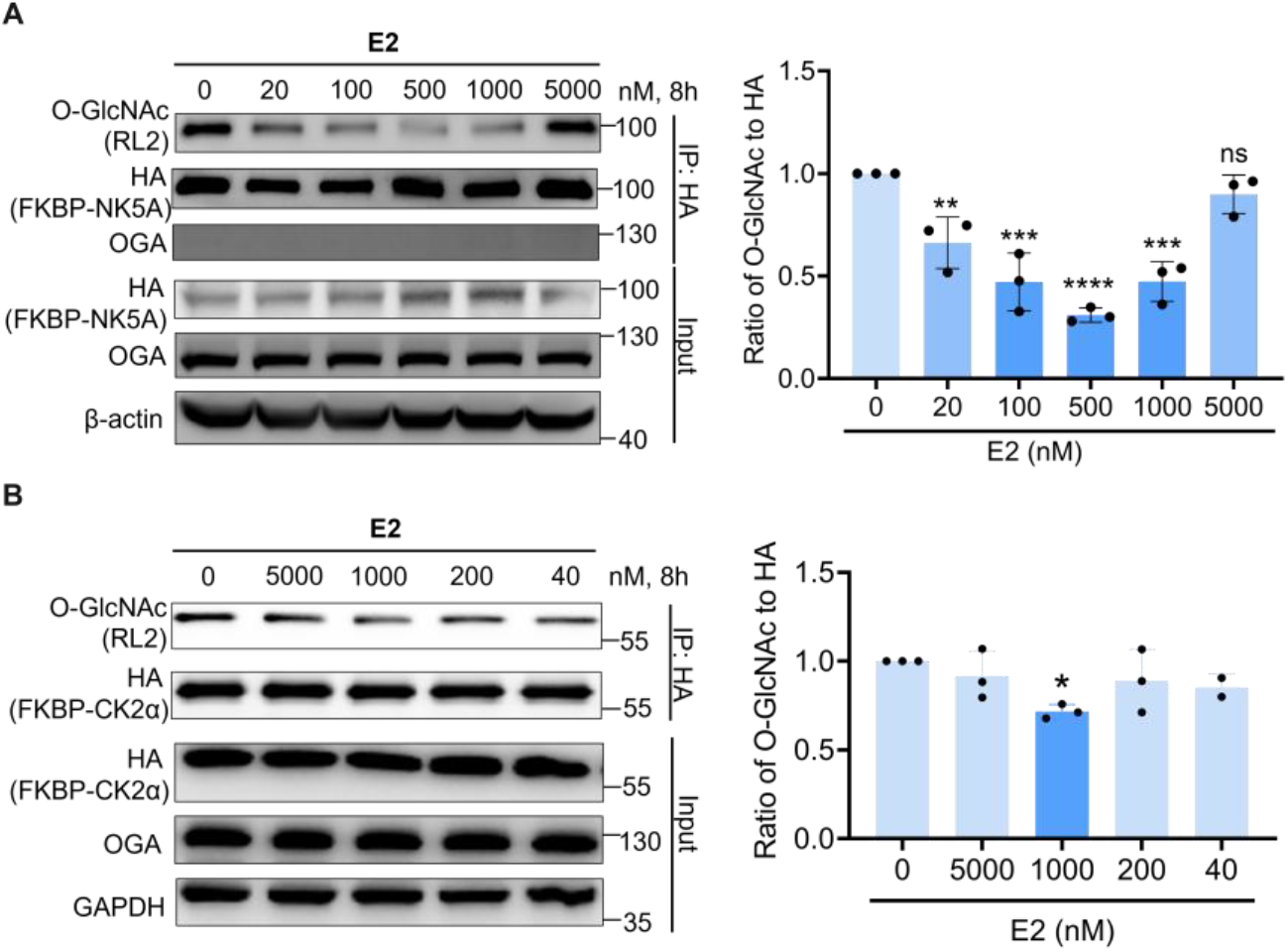
DOGTAC can be applied to other proteins. (A) E2 reduces O-GlcNAcylation of FKBP-NK5A in K562 cells. K562 cells were treated with increasing concentration of E2 for 8 h. FKBP-NK5A was enriched by HA antibody and probed for O-GlcNAc by RL2 antibody. (B) E2 reduces O-GlcNAcylation of FKBP-CK2α in HEK293T cells. FKBP-CK2α was transfected into HEK293T cells with or without treatment with increasing concentration of E2 for 8 h. FKBP-CK2α was enriched by HA antibody and probed for O-GlcNAc by RL2 antibody. Differences were considered statistically significant at p < 0.05. ns, *, **, ***, ****: no significance, p < 0.05, p < 0.01, p < 0.001, p < 0.0001. The results in (A and B) were representative of three biologically independent replicates.

We next examined whether E2 could function in a different cellular context and on an unrelated O-GlcNAcylated protein. Casein kinase 2α (CK2α) is a highly O-GlcNAcylated protein whose modification status regulates global phosphorylation signaling.^32^ We constructed FKBP-2HA-CK2α (FKBP-CK2α) and transiently transfected it into HEK293T cells. Treatment with E2 led to a dose-dependent decrease in O-GlcNAcylation of FKBP-CK2α after 8 h, with an optimal working concentration of approximately 1 μM—higher than that observed for FKBP-NHA9 in K562 cells—highlighting protein- and cell-type-dependent differences in DOGTAC efficacy (Figure 6B).

Collectively, these results demonstrate that DOGTAC can selectively reduce O-GlcNAcylation of multiple FKBP-tagged target proteins across distinct cell lines, supporting the broader feasibility of this chemically induced proximity strategy in tagged-protein models.

## Conclusions

In this work, we developed a chemical strategy for the selective removal of O-GlcNAcylation from target proteins in living cells through heterobifunctional small molecules that recruit endogenous OGA. Starting from reported OGA inhibitors (OGA ligands L5 and L8), we generated bifunctional molecules capable of inducing proximity between OGA and FKBP12^F36V^-tagged NUP98-HOXA9 (NHA9). Although these initial constructs efficiently recruited OGA, they failed to reduce target O-GlcNAcylation, revealing a key mechanistic distinction between proximity induction and functional de-O-GlcNAcylation. These results underscore that simple recruitment of an enzyme is insufficient when the recruiting ligand also potently inhibits the enzyme’s catalytic activity.

We propose that DOGTAC activity is governed by a dynamic competition between the bifunctional recruiter and substrate O-GlcNAc for access to the OGA substrate-binding pocket. At early time points, DOGTAC rapidly recruits OGA to the protein of interest (POI), positioning the enzyme in proximity to its glycosylated substrate. Productive de-O-GlcNAcylation then depends on whether the substrate O-GlcNAc moiety can effectively compete with the OGA-binding element of the DOGTAC for catalytic engagement. If the recruiter binds too tightly, OGA remains occupied in a nonproductive complex, suppressing catalytic turnover despite enforced proximity. Under such conditions, net de-O-GlcNAcylation is diminished and may even shift toward increased O-GlcNAcylation due to perturbation of the OGT–OGA dynamic equilibrium. Conversely, when OGA inhibition is sufficiently attenuated, transient binding permits substrate exchange and enables productive catalysis. This mechanistic balance is further modulated by DOGTAC’s concentration and treatment duration, giving rise to the observed concentration- and time-dependent behaviors (Figure 1C).

Guided by this mechanistic framework, we implemented three complementary strategies to weaken OGA inhibition while preserving recruitment. Stereochemical modulation (Strategy I) revealed that subtle changes in OGA-binding geometry can decouple recruitment from inhibition, as exemplified by the superior DOGTAC activity of the R-isomer B2. Scaffold simplification (Strategy II) and core-motif minimization (Strategy III) further reduced inhibitory potency, yielding multiple DOGTACs that effectively decreased FKBP-NHA9 O-GlcNAcylation. Across all strategies, weaker OGA inhibitors consistently enabled productive de-O-GlcNAcylation, in agreement with prior proximity-based platforms such as DUBTAC and TCIP, which similarly rely on attenuated competitive inhibitors to preserve enzymatic function.^33–36^

Mechanistically, DOGTAC activity depends on ternary complex formation, as demonstrated by rescue experiments with free OGA and FKBP12^F36V^ ligands. Also, Co-IP experiments supported that B2 and C4 could recruit OGA to the targeted protein at a dose- and time-dependent manner. Importantly, the optimized DOGTACs operate at concentrations well below their OGA inhibitory IC_50_ values and do not perturb global O-GlcNAcylation in cells. This separation of local, target-specific de-O-GlcNAcylation from global OGA inhibition represents a key advance over traditional enzyme inhibitors. Moreover, we explored the generality of DOGTAC. Using E2, we successfully reduced the O-GlcNAcylation of FKBP-NK5A in K562 stable cells and FKBP-CK2α in HEK293T cells. The magnitude of de-O-GlcNAcylation varied among targets, with a weaker response observed for FKBP-CK2α than FKBP-NK5A. This difference may reflect target- and cell-context-dependent parameters, including target abundance, basal O-GlcNAcylation stoichiometry, accessibility of modified sites, and the relative abundance of endogenous OGA.

Furthermore, we also used GalT-pulldown assay to validate the DOGTAC effects beyond RL2. And the results suggested that 500 nM E2 treatment could also reduce the O-GlcNAcylation level of FKBP-NK5A, supporting our observations using RL2 antibody (Figure S4).

Several considerations remain for the broader implementation of DOGTAC. First, our current studies are limited to FKBP12^F36V^-fused model proteins, and generalizability to endogenous targets remains to be established. Future efforts should therefore focus on disease-relevant proteins that are highly O-GlcNAcylated, such as PGK1 and YTHDF1/3,^37,38^ to evaluate the applicability of DOGTAC in more native biological contexts.

Second, pronounced hook effects were observed in certain cases, particularly for recruiters with stronger intrinsic OGA inhibition (e.g., B2 and C4). Excessive occupancy of the OGA active site likely limits productive turnover, consistent with our mechanistic model. Further attenuation of OGA inhibition, as exemplified by E2, may mitigate this effect. Accordingly, systematic concentration optimization will be necessary when designing DOGTAC for new targets, considering target abundance across different cell lines. Treatment duration also requires careful control. In general, DOGTAC activity was detectable within 1-3 h and reached maximal effects between 6 and 12 h, suggesting that ~8 h may serve as a practical starting point for initial optimization.

Third, the downstream functional consequences of DOGTAC-mediated de-O-GlcNAcylation remain incompletely understood. RT-qPCR analysis of reported NHA9 downstream genes (*KBTBD10, PLN*, and *PBX3*) in K562 cells revealed no significant transcriptional changes following DOGTAC E2 treatment or global O-GlcNAc suppression by the OGT inhibitor OSMI-4b over several time points, in contrast to NHA9 degradation induced by dTAG-13 (Figure S5A-C).^39^ These results suggest that acute, target-specific or global de-O-GlcNAcylation of NHA9 may be insufficient to drive measurable transcriptional responses under these conditions. Longer treatment durations, additional downstream targets, different cell lines or alternative functional readouts may be required.

Finally, selectivity represents an important consideration. Our biochemical and cellular data indicate that DOGTAC treatment does not induce global alterations in O-GlcNAcylation. Although OGA is the only known enzyme to remove O-GlcNAcylation in human cells, off-target interactions of OGA-binding probes cannot be excluded. Comprehensive selectivity profiling, such as biotin pull-down assays or glycoproteomics, will be important in future studies. As the present work establishes a proof-of-concept framework and identifies initial OGA recruitment probes, such investigations will be critical for advancing DOGTAC toward endogenous targets and broader biological applications.

In summary, we establish DOGTAC as a chemically induced proximity strategy for target-selective protein de-O-GlcNAcylation using endogenous OGA in living cells without global disruption of O-GlcNAc homeostasis. More broadly, this work reveals a key design principle for catalytic proximity systems: competitive enzyme inhibitors can be converted into productive recruiters only when their inhibitory potency is sufficiently attenuated to permit substrate turnover. This balance between recruitment and catalysis should inform the development of future proximity-based strategies for editing dynamic post-translational modifications in living cells.

## Supporting Information

The Supporting information is available.

Supplementary figures and tables; Additional experimental details and methods; and compound synthesis.

## Author Information

### Author Contributions

H.X, B.M, and Y.H contributed equally to this work.

### Notes

The authors declare the following competing financial interest(s): H.X., B.M., and B.W.-L.N. are inventors on a U.S. provisional patent application submitted by The Chinese University of Hong Kong that covers the related work in this study. All other authors declare no competing interests.

## Acknowledgments

The authors sincerely acknowledge Prof. Zhong Zuo (School of Pharmacy, CUHK), Prof. Kam Tong Leung (Department of Paediatrics, CUHK), Prof. Sharon Shui Yee Leung (School of Pharmacy, CUHK), Prof. Xiaoyu Yan (School of Pharmacy, CUHK), Dr. Stephan Scheeff (School of Pharmacy, CUHK), Ms. Josefina Xeque Amada (School of Pharmacy, CUHK), and Ms. Khadija Shahed Khan (School of Pharmacy, CUHK) for scientific discussions. B.W.-L.N. acknowledges funding support from CUHK (CU Medicine Faculty Innovation Award; FIA2020/A/03), Guangdong-Hong Kong-Macao Joint Laboratory for New Drug Screening, Hong Kong RGC (General Research Fund; 14308724), the Peter Hung Pain Research Institute (PHPRI) Research Fund, and the Gerald Choa Neuroscience Institute (GCNI) Research Fund.

