## Supporting information for "Proximity-induced protein deglycosylation by endogenous O-GlcNAcase"

### Table of Contents


1. Table S1. The Structure-Activity-Relationship of reported OGA inhibitors.
2. Figure S1. Time dependent experiments of B2 and C4 at 2 nM.
3. Figure S2. E2 leads to co-IP of OGA in 293T cells with co-transient overexpression of myc-OGA and FKBP-NK5A.
4. Figure S3. The OGA inhibitory activity of TMG, B1, E1, E2 and their ligand LB1, LE1, LE2 in vitro.
5. Figure S4. Target-specific deglycosyation effect of E2 was validated by GalT followed by biotin pulldown assay
6. Figure S5. E2 does not affect the mRNA level of downstream genes of NHA9 in cells.
7. Figure S6. Amino acid sequences of FKBP12F36V-2HA-NUP98-HOXA9 and FKBP12F36V-2HA-NUP98-KDM5A.
8. Experimental Methods.
9. General Chemistry Methods and characterizations.

**Table S1. The Structure-Activity-Relationship of reported OGA inhibitors.[1]**

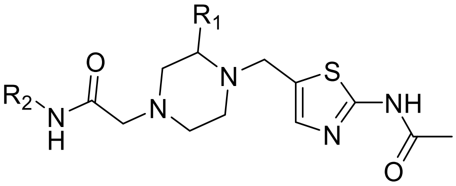

| Compound | $R_1$ | $R_2$ | $IC_{50}$ (μM) |
|---|---|---|---|
| **L1** | H | 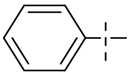 | 2.60 |
| L2 | H | 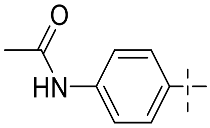 | 1.25 |
| L3 | H | 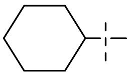 | 1.40 |
| L4 | H | 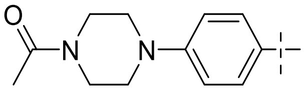 | 1.04 |
| **L5** | Me | 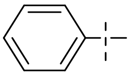 | 0.24 |
| L6 | Me | 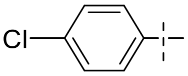 | 0.04 |
| **L7** | Me | 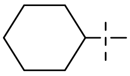 | 0.1 |
| **L8** | Me | 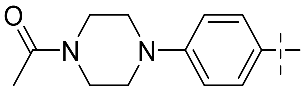 | 0.07 |

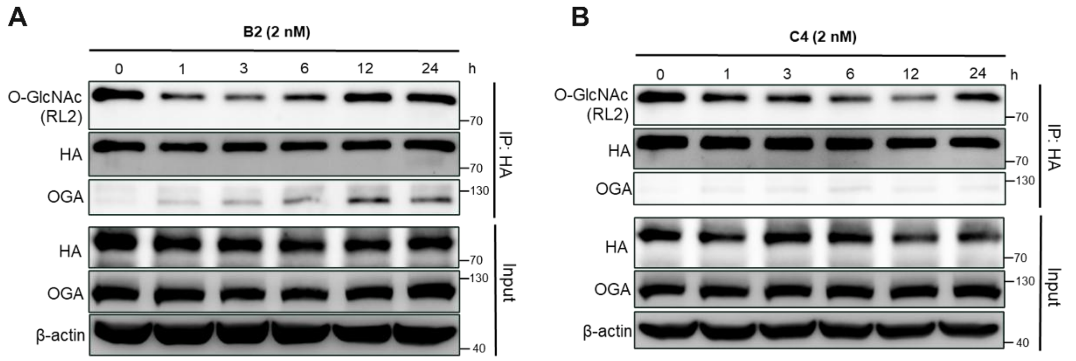


**Figure S1**. **Time dependent experiments of B2 and C4 at 2 nM.** Time-dependent De-O-GlcNAcylation of FKBP-NHA9 by DOGTACs in K562 cells. FKBP-NHA9 K562 cells were treated with B2 (2 nM) and C4 (2 nM) for 0, 3, 6, 12, 24 h.

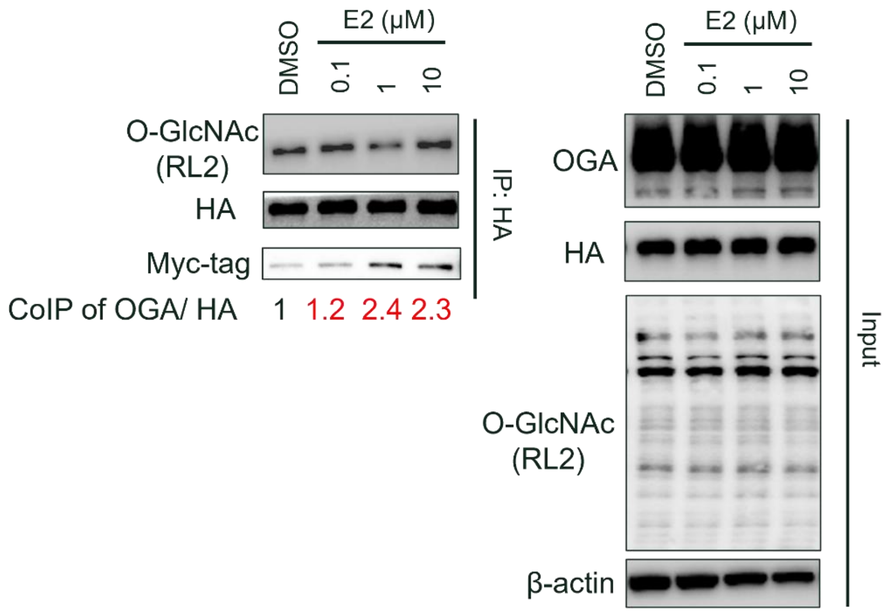


**Figure S2**. **E2 leads to Co-IP of OGA in 293T cells with co-transient overexpression of myc-OGA and FKBP-NK5A.** The myc-OGA and FKBP-NK5A were co-transfected into 293T for 24h. Then E2 were treated for 8h.

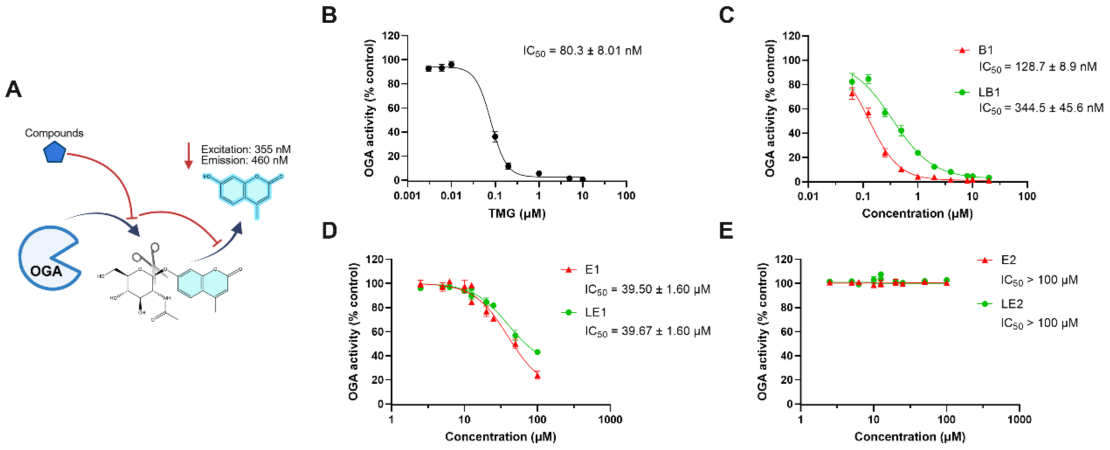


**Figure S3**. **The OGA inhibitory activity of TMG, B1, E1, E2 and their ligand LB1, LE1, LE2 *in vitro*.**

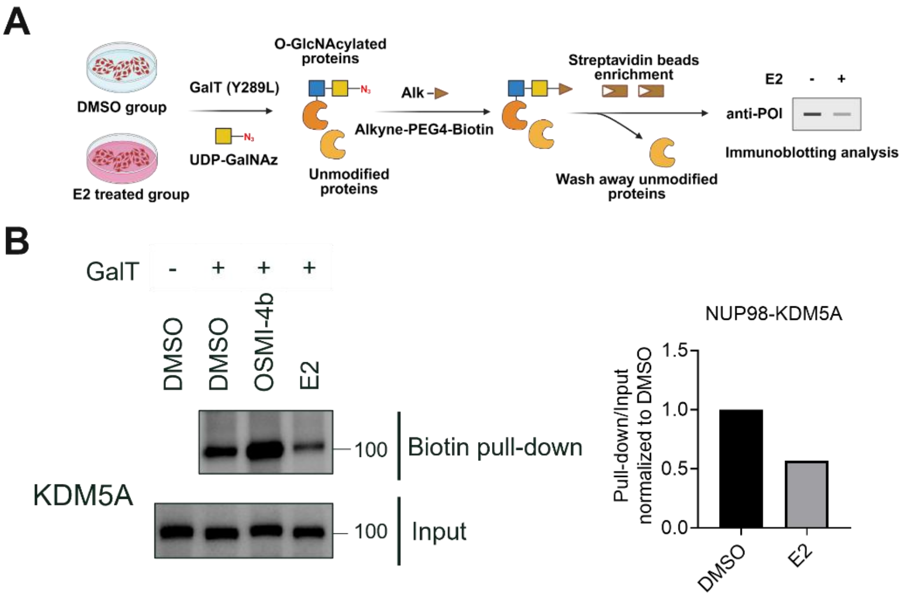


**Figure S4**. **Target-specific deglycosyation effect of E2 was validated by GalT followed by biotin pulldown assay.** 500 nM E2 was used to treat K562-NK5A for 8h.

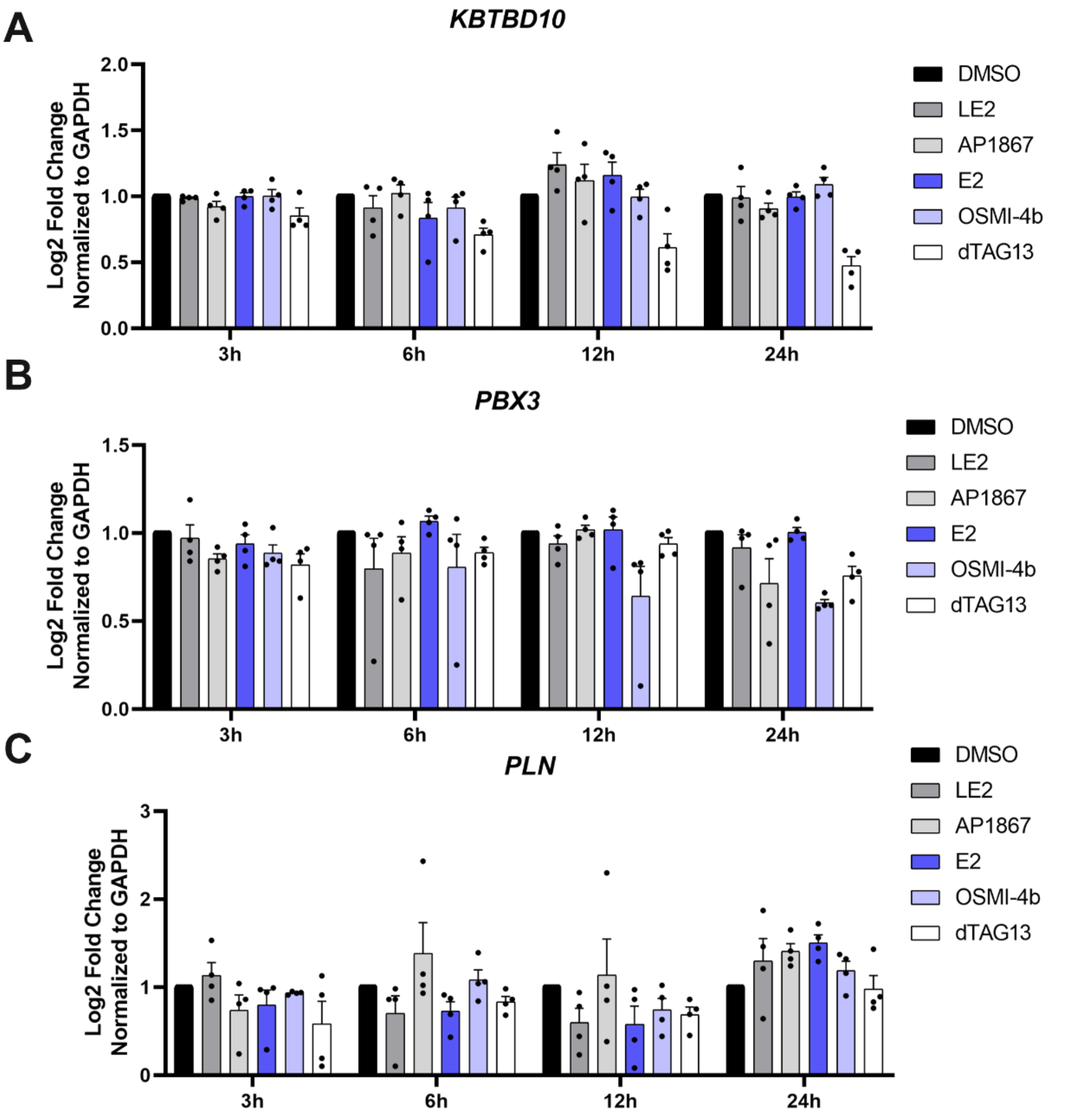


**Figure S5. E2 does not affect the mRNA level of downstream genes of NHA9 in cells.** FKBP-NHA9 K562 cells were treated with 100 nM E2 for 3, 6, 12, 24 hours, followed by RT-qPCR analysis for the mRNA level of KBTBD10 (**A**), PBX3 (**B**), and PLN (**C**).

MGVQVETISPGDGRTFPKRGQTCVVHYTGMLEDGKKVDSSRDRNKPFKFMLGKQ
EVIRGWEEGVAQMSVGQRAKLTISPDYAYGATGHPGIIPPHATLVFDVELLKLEGGYP
YDVPDYAGYPYDVPDYAMFNKSFGTPFGGGTGGFGTTSTFGQNTGFGTTSGGAF
GTSAFGSSNNTGGLFGNSQTKPGGLFGTSSFSQPATSTSTGFGFGTSTGTANTLFG
TASTGTSLFSSQNNAFAQNKPTGFGNFGTSTSSGGLFGTTNTTSNPFGSTSGSLFG
PSSFTAAPTGTTIKFNPPTGTDTMVKAGVSTNISTKHQCITAMKEYESKSLEELRLED
YQANRKGPQNQVGAGTTTGLFGSSPATSSATGLFSSSTTNSGFAYGQNKTAFGTST
TGFGTNPGGLFGQQNQQTTSLFSKPFGQATTTQNTGFSFGNTSTIGQPSTNTMGL
FGVTQASQPGGLFGTATNTSTGTAFGTGTGLFGQTNTGFGAVGSTLFGNNKLTTFG
SSTTSAPSFGTTSGGLFGFGTNTSGNSIFGSKPAPGTLGTGLGAGFGTALGAGQAS
LFGNNQPKIGGPLGTGAFGAPGFNTTTATLGFGAPQAPVVDREKQPSEGAFSENNA
ENESGGDKPPIDPNNPAANWLHARSTRKKRCPYTKHQTLELEKEFLFNMYLTRDRR
YEVARLLNLTERQVKIWFQNRRMKMKKINKDRAKDE*

**FKBP12$^{F36V}$-2HA-NUP98-HOXA9**

MGVQVETISPGDGRTFPKRGQTCVVHYTGMLEDGKKVDSSRDRNKPFKFMLGKQ
EVIRGWEEGVAQMSVGQRAKLTISPDYAYGATGHPGIIPPHATLVFDVELLKLEGGYP
YDVPDYAGYPYDVPDYAGMFNKSFGTPFGGGTGGFGTTSTFGQNTGFGTTSGGA
FGTSAFGSSNNTGGLFGNSQTKPGGLFGTSSFSQPATSTSTGFGFGTSTGTANTLF
GTASTGTSLFSSQNNAFAQNKPTGFGNFGTSTSSGGLFGTTNTTSNPFGSTSGSLF
GPSSFTAAPTGTTIKFNPPTGTDTMVKAGVSTNISTKHQCITAMKEYESKSLEELRLE
DYQANRKGPQNQVGAGTTTGLFGSSPATSSATGLFSSSTTNSGFAYGQNKTAFGTS
TTGFGTNPGGLFGQQNQQTTSLFSKPFGQATTTQNTGFSFGNTSTIGQPSTNTMG
LFGVTQASQPGGLFGTATNTSTGTAFGTGTGLFGQTNTGFGAVGSTLFGNNKLTTF
GSSTTSAPSFGTTSGGLFGFGTNTSGNSIFGSKPAPGTLGTGLGAGFGTALGAGQA
SLFGNNQPKIGGPLGTGAFGAPGFNTTTATLGFGAPQAPVALTDPNASAAQQAVLQ
QHINSLTYSPFGDSPLFRNPMSDPKKKEEDDSMEEKPLKVKGKDSSEKKRKRKLEK
VEQLFGEGKQKSKELKKMDKPRKKKLKLGADKSKELNKLAKKLAKEEERKKKKEKA
AAAKVELVKESTEKKREKKVLDIPSKYDWSGAEESDDENAVCAAQNCQRPCKDKV
DWVQCDGGCDEWFHQVCVGVSPEMAENEDYICINCAKKQGPVSPGPAPPPSFIM
SYKLPMEDLKETS*

**FKBP12$^{F36V}$-2HA-NUP98-KDM5A**

**Figure S6. Amino acid sequences of FKBP12$^{F36V}$-2HA-NUP98-HOXA9 and FKBP12$^{F36V}$-2HA-NUP98-KDM5A**

#### EXPERIMENTAL METHODS

##### Cell culture, DNA constructs, and Reagents

HEK293T cells (kind gift from Prof. Sze Lok Cheng, CUHK) were maintained and cultured at 37 °C with 5% $CO_2$ in Dulbecco's modified Eagle's medium (DMEM) supplemented with 10% fetal bovine serum and 1% penicillin and streptomycin. K562 cells were maintained and cultured at 37 °C with 5% $CO_2$ in Roswell Park Memorial Institute 1640 Medium (RPMI1640) supplemented with 10% fetal bovine serum and 1% penicillin and streptomycin. $FKBP^{F36V}$-CK2α plasmid were constructed by subclone from HTN-CK2α,[2] then homologous recombination into pLVX-puro-$FKBP^{F36V}$ backbone by Beijing Tsingke Biotech Co., Ltd. Transient transfection was conducted according to the manufacture's protocol using Lip2000 TR (AboRo, RL0401).

##### Lentivirus Production and Transduction

To generate lentivirus, we co-transfected HEK293T cells with 5 μg of pLVX-$FKBP12^{F36V}$-2HA-NUP98-HOXA9 or pLVX-$FKBP12^{F36V}$-2HA-NUP98-KDM5A, 5 μg of pΔR8.9, and 1 μg of pCMV-VSVG using JetPRIME. After 40 hours, the viral supernatant was collected, filtered through a 0.45 μm filter, and concentrated with PEG8000-NaCl solution. For transduction, K562 in 6-well plates were incubated with the lentiviral supernatant plus 8 μg/mL polybrene for 16 hours, after which the medium was replaced with fresh medium. 48 hours later, cells were selected with 2 μg/mL puromycin for 48 hours, and expression was confirmed by Western blot. Single-cell clones were isolated by limiting dilution in 96-well plates. After 10 days, clones were expanded, and $FKBP12^{F36V}$-2HA-NUP98-HOXA9 or $FKBP12^{F36V}$-2HA-NUP98-KDM5A expression was assessed by Western blot. The highest-expressing clone was chosen for all subsequent experiments.

##### Western blot assay

Cells were lysed using 100 μL RIPA lysis buffer (Thermo Scientic, 89901) or 200 μL IP lysis buffer (Thermo Scientific, 87788) supplemented with 20 μM OGA inhibitor (Thiamet G, Bidepharm, BD571819) and 100× protease inhibitor (MedChemExpress, HY-K0010). Protein concentrations were determined by bicinchoninic acid (BCA) assay. Samples with equal protein content were boiled in 8× SDS gel loading buffer at 95 °C for 10 minutes. Proteins were separated by 8% SDS-polyacrylamide gel electrophoresis (SDS-PAGE) at 60V for 30 mins and further 140V for 90 mins, transferred to a PVDF membrane at 250mA for 120 mins. Membranes were incubated for 1 h at room temperature with 3% BSA in Tris-buffered saline with 0.1% Tween-20 (TBST) and then immunoblotted with the indicated specific primary antibodies at 4°C overnight. After washing in TBST, the secondary antibodies were incubated with the membranes at room temperature for 1 h. The membranes were washed with TBST for three times (10 mins/each) and visualized on a Bio-Rad ChemiDoc MP Imaging System with Ultra High Sensitivity ECL Kit (MedChemExpress, HY-K1005). Rabbit anti HA (Proteintech, 51064-2-AP, 1:2000 dilution), Mouse anti HA (Abclonal, AE008, 1:1000 dilution), Rabbit anti RL2 (abcam, AB2739, 1:1000 dilution), Rabbit anti β-actin (ABclonal, AC026, 1:10000 dilution), Rabbit anti OGA (Proteintech, 14711-1-AP, 1:2000 dilution), anti-mouse-HRP secondary antibody (cell signaling, 7076S, 1:10000 dilution), anti-rabbit-HRP secondary antibody (cell signaling, 7074S, 1:10000 dilution), anti-rabbit-DyLight 488 (Invitrogen, 35552, 1:6000 dilution). The western blot was quantified using Image Lab software.

##### Immunoprecipitation

For transient overexpression experiments using HEK293T, 5 X $10^5$ HEK293T cells were seed in each well of 6-well plate. After cells attached, transfection reagents and plasmids (0.3 μg for FKBP12$^{F36V}$-CK2α and 1.2 μL TR for each well in 6 well plate) were prepared and added in DMEM. After 24 h transfection, media were replaced by fresh warm complete media with DMSO or probes. After treatment for indicated duration, media were aspired and 200 μL IP lysis buffer was added to each well. The lysate was incubated on ice for 20 min, spun down at 14,000 RPM at 4 °C for 15 mins, and the supernatants were collected for bicinchoninic acid (BCA) assay and normalized to a final 2 mg/mL. 40 μL was saved as input. The remaining samples were subjected to anti-HA nanobody beads (10 μL each sample (Shenzhen KangTi Biopharm Technology Co., Ltd, KTSM1335)) and incubated at 4°C for overnight. For the co-transfection experiment, cells in 6-well plate were transfected with FKBPF36V-NK5A (0.5 μg) and Myc-OGA (0.4 μg) for 24h. The beads were washed with PBS containing 0.1% Tween-20 (PBST) for three times and further resuspended in 25 μL 2× SDS loading buffer, boiling at 95 °C for 10 mins, and followed by western blotting analysis.

For K562 stable cell lines, 2.5 × $10^6$ cells were split into 6 cm diches in indicated media. The probes were treated for indicated time. The cells in media were transferred to 15 mL tubes and spun down at 1200 RPM at room temperature for 3 mins 30 s. Then media in the supernatant was removed and cells were transferred to 1.5 mL tubes using 1 mL PBS and spun down again at 1200 RPM, room temperature for 3 mins 30 s to remove PBS. The cells were lysed using 200 μL IP lysis buffer at ice for 30 mins, spun down at 14,000 RPM at 4 °C for 15 mins, and the supernatants were collected for bicinchoninic acid (BCA) assay and normalized to equal protein contents for each, 45 μL was saved as input. The remaining samples were subjected to anti-HA nanobody beads (10 μL each sample) and incubated at 4 °C for overnight. The beads were washed with PBS containing 0.1% Tween-20 (PBST) for three times and further resuspended in 25 μL 2× SDS loading buffer, boiling at 95 °C for 10 mins, and followed by separation with western blot.

For western blot, we used anti-mouse-HRP (Cell signaling, #7076) as secondary antibody for RL2 and anti-rabbit-DyLight 488 (Invitrogen, #35552) as secondary antibody for total HA being IPed.

**OGA *in-vitro* enzymatic assay**

Human OGA was purified from HEK293T cells that had been transfected with the Halotag-fused full length OGA construct. The enzyme was purified according to the Halotag® Protein Purification System protocol. For each 15 cm dish of HEK293T cells, 8 X $10^6$ cells were seeded, and then 15 μg of HTN-OGA plasmid with 60 μL of TR were well mixed and added into cells. After transfection for 30 h, cells were collected and HTN-OGA was enriched by pre-washed HaloLink resin (200 μL suspension). The lysate beads mixture was incubated overnight and washed robustly by wash buffer (1 X PBS, 1 mM DTT, 0.02% NP40) and the beads were resuspended in 300 μL wash buffer.

Enzymatic reactions were carried out in reaction containing 50 mM $NaH_2PO_4$, 100 mM NaCl, 0.01% Tween 20, and 1 mM dithiothreitol (DTT, pH 7.4) using 2 mM 4-methylumbelliferyl N-acetyl-β-D-glucosaminide dihydrate (TargetMol, T37571) as a substrate. The amount of human OGA used in the reaction was 5 μL (on the HaloLink resin). Test compound of varying concentrations was added to the enzyme prior to initiation of the reaction. TMG was tested as positive control. The reaction was performed at room temperature (RT) in a black 96-well assay plate (COSTAR, 3916) and was initiated by the addition of substrate. After 2 h of reaction, the fluorescence intensity of the product was monitored every 5 min for 60 min using a multimode plate reader (CLARIOstar), with excitation at 355 ± 15 nm and emission at

460 ± 20 nm. The detector gain was optimized to ensure that the signal remained within the linear detection range and did not reach the upper reading limit. Raw fluorescent values were background (control wells without OGA)-subtracted prior to calculating. Data analysis was done with GraphPad Prism 9.5.1 (GraphPad Software, San Diego, CA) statistical analysis software using a nonlinear regression for a sigmoidal dose-response with a variable slope. The $IC_{50}$ value was defined as the concentration inhibiting the activity by 50%.

**GalT followed by biotin pull-down assay**

Cells were harvested and lysed in RIPA buffer supplemented with protease inhibitors and TMG. Following sonication and clearing by centrifugation, protein concentrations were normalized to 2 mg/mL. Lysates were reduced with DTT (56 °C, 30 min), alkylated with iodoacetamide (room temperature, 30 min, dark), and subjected to methanol/chloroform precipitation. For enzymatic labeling, 1 mg of the resuspended protein pellet was incubated with UDP-GalNAz, $MnCl_2$, and mutant Y289L GalT enzyme in labeling buffer (50 mM Hepes pH 7.9, 125 mM NaCl, 5% NP-40) for 24 h at 4 °C. Following a second methanol/chloroform precipitation, the GalNAz-labeled proteins were conjugated to Alkyne-PEG4-Biotin via copper-catalyzed azide-alkyne cycloaddition (CuAAC) using a $CuSO_4$-BTTAA complex and sodium ascorbate for 1 h at 32 °C. Proteins were precipitated a final time, redissolved in 0.2% SDS in PBS, and incubated with pre-equilibrated streptavidin magnetic beads for 2 h at room temperature. Finally, the beads were stringently washed with SDS in PBS (0.2%) three times and Tween-20 in PBS (0.5%) buffer three times, and the enriched glycoproteins were eluted using 2X loading buffer heated at 95 degree for 10 mins for downstream analysis.

**RT-qPCR**

After 24 hours of DOGTAC treatment, cells were collected for RNA isolation. Total RNA was extracted using the RNAfast200 Kit (Fastagen 220010) and reverse transcribed into cDNA with the PrimeScript™ RT Reagent Kit (Perfect Real Time) (TaKaRa RR037A). Real-time quantitative PCR was performed in triplicate using TB Green (TaKaRa, RR830) on the LightCycler® PRO system (Roche). Relative fold changes were calculated using the comparative ΔΔCt method, with GAPDH as the housekeeping gene for normalization.

The primers used are listed below:

| Gene | (5'-3') Forward primer | (5'-3') Reverse primer |
|---|---|---|
| *GAPDH* | GAGTCAACGGATTTGGTCGT | TTGATTTTGGAGGGATCTCG |
| *KBTBD10* | GGAGGACTATATGTGGATGAAGAA | ACCGAAGAGACACCTGGCTGAA |
| *PLN* | AGCACGTCAAAAGCTACAGAATCT | CTGATGTGGCAAGCTGCAGATC |
| *PBX3* | CATCACAGTGTCACAGGTATCC | CAGCATAGAGGTTGGCTTCTT |

**Molecular Docking**

Molecular docking was performed using the Glide module (XP) within the Schrödinger Suite (Version 2021-2). Compounds were docked with the human OGA protein (PDB ID: 5UHL). The protein was prepared using the Protein Preparation Wizard suite. Ligand structures were prepared using the LigPrep suite. The reported grid defining the docking site was subsequently generated according to the binding pocket of original TMG. Default parameters using XP modes were used for docking. The top-ranked results were used for analysis. Visualization was performed using PyMOL (version 3.0.3).

**General Chemistry methods**

Chemicals and reagents were purchased from commercial vendors (Bide pharm, Hao Yuan Chemexpress, Dieckmann, Meryer) and used without further purification. Common solvents and reagents used for reactions (DCM, PhMe, AcOH, MeCN, DMF, DIPEA, TEA) were purchased from Meryer in anhydrous conditions. Temperature above room temperature referred to the oil bath temperature. For cooling to 0 °C, a water/ice bath was used.

All reactions were monitored using TLC and visualized by UV light (254 nM and 365 nM) or sained with iodine, $KMnO_4$ (1.5 g of $KMnO_4$. 10 g $K_2CO_3$, and 1.25 mL 10% NaOH in 200 mL water). Reaction purifications were carried out using Flash Chromatography (230-400 mesh silica gel 60), Biotage® or thin layer chromatography (pTLC, PLC Silica Gel 60 GF254, Leyan, 0.4-0.5 mm thickness) and solvents were used without prior purification. For reverse-phase chromatography purification, Agilent 1260 Infinity II Preparative LC System was used with HP C18 RediSep Rf columns (5 μM), and the gradient was set to 40% of methanol in $H_2O$ containing 0.1% TFA progressing to 90% of methanol in $H_2O$ containing 0.1% TFA, maintaining a flow rate of 20 mL/min and utilizing a UV detector set at 220/254/365 nm.

NMR spectra were recorded on Burker AVANCE III 700 MHz spectrometer in the deuterated solvents ($CDCl_3$, DMSO-$d_6$). Multiplicities are reported with the following abbreviations: s singlet; d doublet; t triplet; q quartet; p pentet; m multiplet; br broad; dd doublet of doublets; dt doublet of triplets; td triplet of doublets; ddd doublet of doublet of doublets; dq doublet of quartet; ddt doublet of doublet of triplets; qd quartet of doublets; dtd doublet of triplet of doublets; tdt triplet of doublet of triplets; Chemical shifts are reported in ppm relative to the residual solvent peak and J values are reported in Hz. Mass spectrometry data were collected on an Agilent 6479 Triple Quadrupole LC/MS instrument (ESI, low resolution) or Thermo Q Exactive Focus Orbitrap Mass Spectrometer (ESI, positive, HRMS). The purity of all final products was confirmed to be >95% through HPLC analysis using a Waters ACQUITY Premier HPLC with a 2998 PDA detector, and the gradient was set to 10% acetonitrile in $H_2O$ containing 0.1% TFA progressing to 100% of acetonitrile at a flow rate of 0.5 mL/min.

**General Method A: Boc deprotection of amines**

The Boc protected amine was dissolved in DCM/TFA (4:1) resulting in a 0.1 M solution. After completion of the reaction (usually 1-2 h), the solvent was removed directly under reduced pressure to give the amines without further purification.

**General Method B: coupling for amines and acids**

The amines (1.0 eq), acids (1.0 eq), and HATU (1.2 eq) were dissolved in DCM resulting in a 0.1 M solution, DIPEA (5.0 eq) was added, and the mixture was stirred at rt. After completion of the reaction (usually overnight, monitored by TLC), the mixture was diluted with DCM (usually 3 mL), washed with saturated sodium bicarbonate solution (usually 5 mL), saturated ammonium chloride aqueous solution (usually 5 mL), brine (usually 5 mL) and concentrated under reduced pressure. The resulting residue was purified with flash column chromatography (silica gel) to give the products.

**Scheme S1.** Synthesis of OGA ligand **5 (L10)**

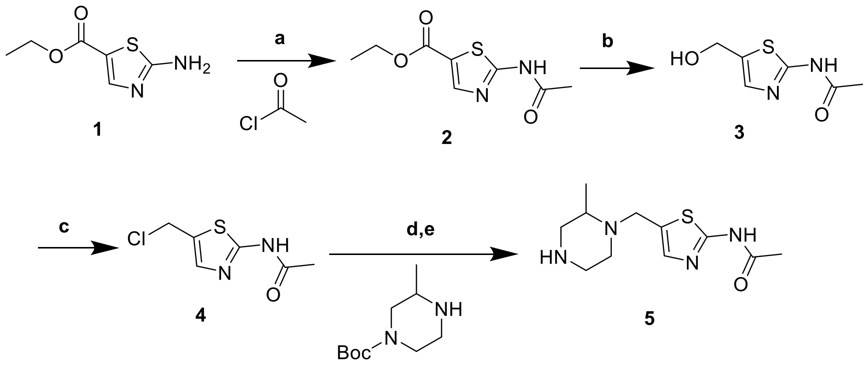

Reaction and conditions: a) TEA, dry DCM, 0°C-rt, 5 h, 53%; b) lithium triethylhydroborate, dry PhMe,-15°C, 4 h, 96%; c) $SOCl_2$, dry DCM, 0°C-rt, 2.5 h, 98%; d) TEA, MeCN, rt, 12 h, 71%; e) TFA, DCM, rt, 1.5 h.

**Ethyl 2-acetamidothiazole-5-carboxylate (2)**

The Ethyl 2-aminothiazole-5-carboxylate (**1**) (1.0 g, 5.81 mmol, 1.0 eq) and TEA (1.61 mL, 11.61 mmol, 2.0 eq) were dissolved in dry DCM (18 mL) and the mixture was stirred at rt for 10 minutes. The acetyl chloride (500 μL, 6.97 mmol, 1.2 eq) was added slowly at 0 °C. And then the mixture was stirred at rt for 5 h until the reaction was complete (monitored by TLC). The mixture was filtered and the solid was washed by water (10 mL) and dried to give the crude without further purification. (white solid, 655 mg, 53%). $^1H$ NMR (700 MHz, DMSO-$d_6$) δ 12.59 (s, 1H), 8.13 (s, 1H), 4.27 (q, $J$ = 7.1 Hz, 2H), 2.19 (s, 3H), 1.29 (t, $J$ = 7.1 Hz, 3H).

**N-(5-(hydroxymethyl)thiazol-2-yl)acetamide (3)**

The intermediate **2** (5.0 g, 23.34 mmol, 1.0 eq) was dissolved in dry PhMe (75 mL), lithium triethylhydroborate (1M in THF, 94 mL, 93.35 mmol, 4.0 eq) was added dropwise at -15 °C. The mixture was stirred at -15 °C for 4 h until the reaction was complete (monitored by TLC). The mixture was quenched with ice water (50 mL) and extracted with EA (50 mL). The aqueous layer was acidified AcOH (100 mL) and then was extracted with EA (120 mL) for 12 times. The organic layer was combined and concentrated to give the crude as intermediate compound **3** without further purification (Pale yellow solid, 3.854 g, 96%). $^1H$ NMR (700 MHz, DMSO-$d_6$) δ 12.11 (s, 1H), 7.26 (s, 1H), 4.57 (s, 2H), 2.13 (s, 3H).

**N-(5-(chloromethyl)thiazol-2-yl)acetamide (4)**

The intermediate **3** (3.85 g, 22.38 mmol, 1.0 eq) was dissolved in dry DCM (20 mL), $SOCl_2$ (3.25 mL, 44.76 mmol, 2.0 eq) was added slowly at 0 °C. The mixture was stirred at rt for 2.5 h until the reaction was complete (monitored by TLC). The mixture was concentrated under reduced pressure and the $SOCl_2$ was removed completely using DCM (10 mL) for multiple times. The resulting residue was further purified by slurring with anhydrous ether, followed by filtration and drying to give intermediate compound **4** (brown solid, 4.18 g, 98%). $^1H$ NMR (700 MHz, DMSO-$d_6$) δ 12.41 (s, 1H), 7.54 (s, 1H), 5.04 (d, $J$ = 17.2 Hz, 1H), 4.60 (d, $J$ = 11.3 Hz, 1H), 2.18 (t, $J$ = 14.7 Hz, 3H).

**N-(5-((2-methylpiperazin-1-yl)methyl)thiazol-2-yl)acetamide (5) (L10)**

The intermediate **4** (2.0 g, 10.49 mmol, 1.0 eq) was dissolved in MeCN (20 mL), TEA (7.29 mL, 52.45 mmol, 5.0 eq) and tert-butyl 3-methylpiperazine-1-carboxylate (2.53 mL, 12.59 mmol, 1.2 eq) were added. The mixture was stirred at rt for overnight (12 h) until the reaction was complete (monitored by TLC). The mixture was concentrated under reduced pressure and purified with flash column chromatography (silica gel, MeOH: DCM=1:100 to 1:50) to give **Boc-5** (white solid, 2.6277 g, 71%). LC-MS (ESI) m/z= 355.2. $^1$H NMR (700 MHz, DMSO-$d_6$) δ 11.99 (s, 1H), 7.27 (s, 1H), 3.91 (d, $J$ = 14.5 Hz, 1H), 3.64 (d, $J$ = 14.5 Hz, 1H), 3.53 (s, 2H), 3.03 (q, $J$ = 9.2, 8.6 Hz, 1H), 2.84 (s, 1H), 2.71 (s, 1H), 2.62 (dt, $J$ = 11.8, 3.3 Hz, 1H), 2.34 (s, 1H), 2.11 (s, 3H), 1.38 (s, 9H), 1.03 (d, $J$ = 6.2 Hz, 3H).

**5** was synthesized from **Boc-5** (2.63 g, 7.41 mmol) using **General Mothed A**. gray solid. $^1$H NMR (700 MHz, DMSO-$d_6$) δ 12.22 (s, 1H), 7.49 (s, 1H), 3.54 – 3.47 (m, 1H), 3.34 (d, $J$ = 12.8 Hz, 2H), 3.22 – 3.16 (m, 1H), 3.16 – 3.11 (m, 1H), 3.09 (td, $J$ = 7.3, 4.7 Hz, 1H), 2.97 (t, $J$ = 13.0 Hz, 1H), 2.89 (s, 1H), 2.75 (s, 1H), 2.15 (s, 3H), 1.29 (d, $J$ = 5.9 Hz, 3H), 1.25 (d, $J$ = 6.4 Hz, 1H).

**Scheme S2.** Synthesis of Intermediate **6a**

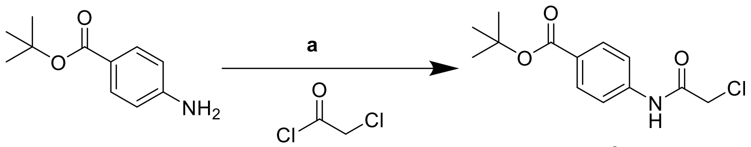


**6a**

Reaction and conditions: a) TEA, DCM, 0 °C-rt, 3 h, 95%.

**Tert-butyl 4-(2-chloroacetamido)benzoate (6a)**

The tert-butyl 4-aminobenzoate (600 mg, 3.10 mmol, 1.0 eq) was dissolved in DCM (12 mL), TEA (475 μL, 3.42 mmol, 1.1 eq) was added. Then 2-chloroacetyl chloride (275 μL, 3.42 mmol, 1.1 eq) was added at 0 °C. The mixture was stirred at rt for 3 h until the reaction was complete (monitored by TLC). The mixture was quenched with Water (15 mL) and the organic layer was concentrated. The resulting residue was purified with flash column chromatography (silica gel, EA: Hexane=1:50 to 1:5) to give **6a** (white solid, 796.1 mg, 95%). $^1$H NMR (700 MHz, $CDCl_3$) δ 8.37 (s, 1H), 7.99 – 7.97 (m, 2H), 7.63 – 7.61 (m, 2H), 4.21 (s, 2H), 1.59 (s, 9H).

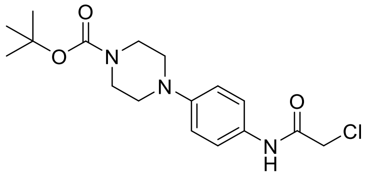


**6b**

**Tert-butyl 4-(4-(2-chloroacetamido)phenyl)piperazine-1-carboxylate (6b)**

Intermediate **6b** was synthesized following the same procedure for preparing intermediate **6a** from tert-butyl 4-(4-aminophenyl)piperazine-1-carboxylate (600 mg, 2.16 mmol). Column chromatography (silica gel, EA: Hexane=1:30 to 1:2) to give **6b** (white solid, 616.6 mg, 81%). $^1$H NMR (700 MHz, $CDCl_3$) δ 8.15 (s, 1H), 7.44 – 7.42

(m, 2H), 6.93 – 6.90 (m, 2H), 4.18 (s, 2H), 3.57 (t, *J* = 5.1 Hz, 4H), 3.12 – 3.07 (m, 4H), 1.48 (s, 9H).

**Scheme S3.** Synthesis of OGA ligand **7a**

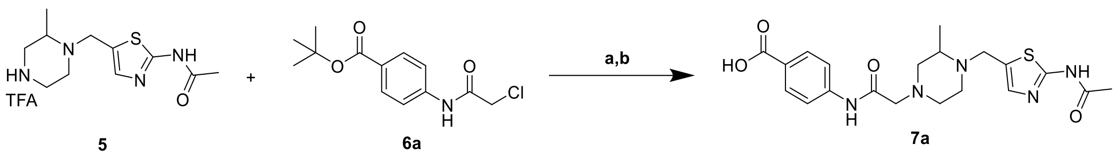


Reaction and conditions: a) $K_2CO_3$, MeCN, rt, 17 h, 29%; b) TFA, DCM, rt, 2 h.

##### 4-(2-(4-((2-acetamidothiazol-5-yl)methyl)-3-methylpiperazin-1-yl)acetamido)benzoic acid (7a)

The intermediate **5** (200 mg, 0.74 mmol, 1.0 eq), **6a** (287 mg, 0.82 mmol, 1.1 eq), and Potassium carbonate (515 mg, 3.71 mmol, 5.0 eq) were dissolved in MeCN (2 mL). The mixture was stirred at rt for overnight (17 h) until the reaction was complete (monitored by TLC). The mixture was diluted with water (3 mL) and extracted with EA (5 mL). The organic layer was concentrated. The resulting residue was purified with flash column chromatography (silica gel, Methanol: DCM =1:100 to 1:30) and further purified via reverse-ISCO to give the **t-Bu-7a** (white solid, 106.1 mg, 29%). $^1$H NMR (700 MHz, $CDCl_3$) δ 12.16 (s, 1H), 9.25 (s, 1H), 7.96 – 7.93 (m, 2H), 7.61 – 7.57 (m, 2H), 7.22 (s, 1H), 4.04 (d, *J* = 14.5 Hz, 1H), 3.71 (d, *J* = 14.5 Hz, 1H), 3.16 – 3.07 (m, 2H), 2.85 – 2.81 (m, 1H), 2.73 (t, *J* = 10.8 Hz, 2H), 2.63 (s, 1H), 2.55 – 2.47 (m, 1H), 2.45 – 2.40 (m, 1H), 2.33 (s, 3H), 2.30 – 2.26 (m, 1H), 1.58 (s, 9H), 1.19 (t, *J* = 6.4 Hz, 3H).

The **t-Bu-7a** (100 mg. 0.21 mmol) was dissolved in TFA/DCM (1:4, 8 mL). The mixture was stirred at rt for 2h until the reaction was complete (monitored by TLC). The mixture was concentrated under reduced pressure to give the intermediate compound **7a** without further purification (white solid).

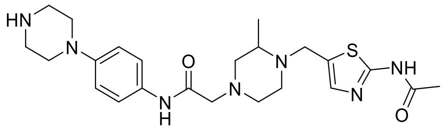


**7b**

##### 2-(4-((2-acetamidothiazol-5-yl)methyl)-3-methylpiperazin-1-yl)-N-(4-(piperazin-1-yl)phenyl)acetamide (7b)

OGA ligand **7b** was synthesized following the same procedure for preparing OGA ligand **7a** from **6b** (200 mg, 0.57 mmol). Column chromatography (silica gel, Methanol: DCM =1:100 to 1:30) and further purified via reverse-ISCO to give the **Boc-7b** (white solid, 64.6 mg, 20%). $^1$H NMR (700 MHz, $CDCl_3$) δ 11.60 (s, 1H), 8.92 (s, 1H), 7.46 – 7.42 (m, 2H), 7.21 (s, 1H), 6.91 – 6.88 (m, 2H), 4.04 (d, *J* = 14.5 Hz, 1H), 3.70 (d, *J* = 14.5 Hz, 1H), 3.57 (t, *J* = 5.1 Hz, 4H), 3.12 – 3.08 (m, 2H), 3.06 (d, *J* = 5.9 Hz, 4H), 2.81 (dt, *J* = 11.5, 3.2 Hz, 1H), 2.76 – 2.70 (m, 2H), 2.61 (s, 1H), 2.49 (t, *J* = 10.4 Hz, 1H), 2.41 (dd, *J* = 20.4, 10.2 Hz, 1H), 2.31 (s, 3H), 2.26 (s, 1H), 1.47 (s, 9H), 1.18 (d, *J* = 6.2 Hz, 3H).

**7b** was synthesized from **Boc-7b** (60 mg, 105 μmol) using **General Method A**.

**Scheme S4.** Synthesis of **A1**

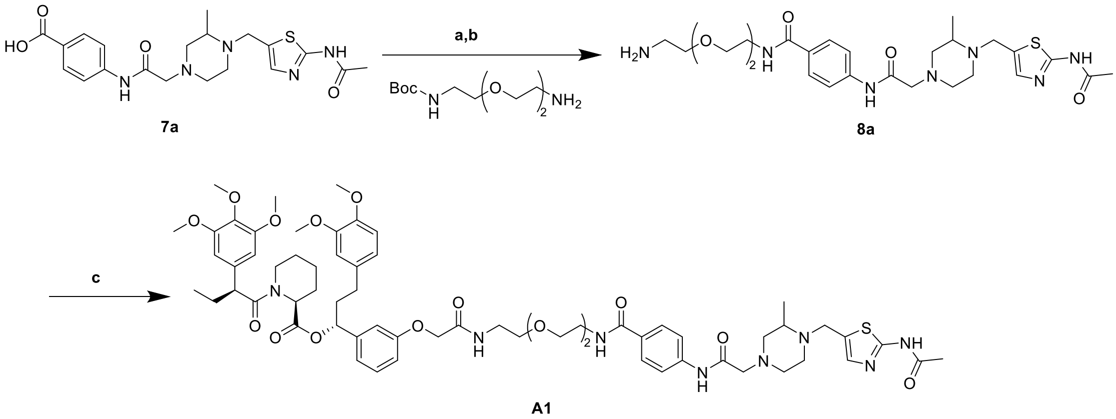

Reaction and conditions: a) HATU, DIPEA, DCM, rt, 17 h, 65%; b) TFA, DCM, rt, 1 h; c) AP1867, HATU, DIPEA, DCM, 15h, 40%.

**4-(2-(4-((2-acetamidothiazol-5-yl)methyl)-3-methylpiperazin-1-yl)acetamido)-N-(2-(2-(2-aminoethoxy)ethoxy)ethyl)benzamide (8a)**

Intermediate **8a** was synthesized from **7a** (40 mg, 93 μmol) and BocNH-PEG2-$NH_2$ (23 mg, 93 μmol) using **General Method B**. Flash column chromatography (silica gel, MeOH in DCM: 0-8%) to give the **Boc-8a** (white solid, 39.6 mg, 65%). $^1$H NMR (700 MHz, $CDCl_3$) δ 11.23 (s, 1H), 9.24 (s, 1H), 7.78 (d, *J* = 8.2 Hz, 2H), 7.62 (d, *J* = 8.2 Hz, 2H), 7.22 (s, 1H), 6.75 (s, 1H), 4.04 (d, *J* = 14.2 Hz, 1H), 3.72 (d, *J* = 14.7 Hz, 1H), 3.67 (s, 4H), 3.65 – 3.62 (m, 4H), 3.54 (t, *J* = 5.2 Hz, 2H), 3.31 (q, *J* = 5.6 Hz, 2H), 3.11 (d, *J* = 7.3 Hz, 2H), 2.82 (dd, *J* = 11.7, 3.6 Hz, 1H), 2.73 (t, *J* = 11.4 Hz, 2H), 2.62 (s, 1H), 2.51 (t, *J* = 10.6 Hz, 1H), 2.44 (t, *J* = 10.4 Hz, 1H), 2.31 (s, 3H), 2.28 (d, *J* = 10.1 Hz, 1H), 1.42 (s, 9H), 1.19 (d, *J* = 6.4 Hz, 3H).

**8a** was synthesized from **Boc-8a** (30 mg, 45 μmol) using **General Method A**.

**(1R)-1-(3-((1-(4-(2-(4-((2-acetamidothiazol-5-yl)methyl)-3-methylpiperazin-1-yl)acetamido)phenyl)-1,12-dioxo-5,8-dioxa-2,11-diazatridecan-13-yl)oxy)phenyl)-3-(3,4-dimethoxyphenyl)propyl (2S)-1-((S)-2-(3,4,5-trimethoxyphenyl)butanoyl)piperidine-2-carboxylate (A1)**

**A1** was synthesized from **8a** (20 mg, 30 μmol) and 2-(3-((R)-3-(3,4-dimethoxyphenyl)-1-(((S)-1-((S)-2-(3,4,5-trimethoxyphenyl)butanoyl)piperidine-2-carbonyl)oxy)propyl)phenoxy)acetic acid (hereafter **AP1867**, 21 mg, 30 μmol) using **General Method B**. pTLC (DCM: MeOH=15:1) to give the product **A1** (white solid, 14.9 mg, 40%). HPLC purity (254 nM)= 95.22%, RT= 15.371 min. $^1$H NMR (700 MHz, $CDCl_3$) δ 11.58 (s, 1H), 9.22 (d, *J* = 3.0 Hz, 1H), 7.78 (d, *J* = 8.3 Hz, 2H), 7.60 (d, *J* = 8.5 Hz, 2H), 7.19 (d, *J* = 6.5 Hz, 1H), 7.17 (d, *J* = 7.9 Hz, 1H), 7.12 (q, *J* = 7.9, 5.9 Hz, 1H), 6.78 – 6.74 (m, 3H), 6.64 (dt, *J* = 9.0, 2.1 Hz, 3H), 6.39 (s, 2H), 6.30 (s, 1H), 5.60 (dd, *J* = 8.3, 5.4 Hz, 1H), 5.46 – 5.43 (m, 1H), 4.47 (t, *J* = 3.3 Hz, 2H), 4.02 (d, *J* = 14.5 Hz, 1H), 3.84 (p, *J* = 2.8 Hz, 9H), 3.76 (d, *J* = 10.3 Hz, 3H), 3.70 (d, *J* = 14.6 Hz, 1H), 3.67 (s, 5H), 3.62 (d, *J* = 2.4 Hz, 4H), 3.59 (dd, *J* = 13.1, 4.7 Hz, 8H), 3.54 – 3.52 (m, 1H), 3.10 (d, *J* = 6.8 Hz, 2H), 2.82 – 2.76 (m, 2H), 2.72 (t, *J* = 10.2 Hz, 2H),

2.64 – 2.60 (m, 1H), 2.58 – 2.54 (m, 1H), 2.54 – 2.47 (m, 2H), 2.47 – 2.41 (m, 2H), 2.29 (d, *J* = 5.1 Hz, 3H), 2.24 (dd, *J* = 19.1, 11.1 Hz, 1H), 2.08 – 2.03 (m, 2H), 1.91 (dtd, *J* = 12.2, 10.1, 5.3 Hz, 1H), 1.72 – 1.67 (m, 3H), 1.61 – 1.58 (m, 1H), 1.44 – 1.39 (m, 1H), 1.17 (d, *J* = 6.2 Hz, 3H), 0.88 (t, *J* = 3.6 Hz, 3H). $^{13}$C NMR (176 MHz, $CDCl_3$) δ 172.7, 170.6, 168.6, 168.4, 168.1, 166.8, 159.7, 157.2, 153.2, 148.8, 147.3, 142.3, 140.3, 136.5, 135.3, 134.2, 133.3, 129.9, 128.2, 120.2, 120.2, 119.7, 119.0, 118.8, 113.7, 113.0, 111.6, 111.2, 104.9, 75.7, 70.2, 69.9, 69.7, 67.3, 61.8, 60.9, 60.8, 60.7, 56.3, 55.9, 55.9, 55.9, 55.8, 53.6, 52.1, 50.8, 49.1, 43.5, 39.7, 38.8, 38.8, 38.3, 31.3, 29.7, 26.8, 25.3, 23.2, 20.9, 12.6. HRMS (ESI, m/z): calcd for $C_{64}H_{84}N_8O_{15}SNa^+$ $[M+Na]^+$: 1259.5669, found 1259.5671.

**A2-A18** were synthesized following the same procedure for preparing **A1**.

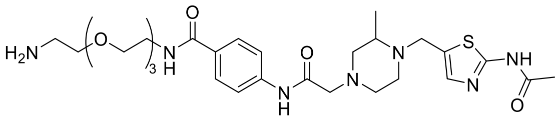

**8b**

**4-(2-(4-((2-acetamidothiazol-5-yl)methyl)-3-methylpiperazin-1-yl)acetamido)-N-(2-(2-(2-(2-aminoethoxy)ethoxy)ethoxy)ethyl)benzamide (8b)**

Intermediate **8b** was synthesized from **7a** (40 mg, 93 μmol) and BocNH-PEG3-$NH_2$ (27 mg, 93 μmol) using **General Method B**. Flash column chromatography (silica gel, MeOH in DCM: 0-8%) to give the **Boc-8b** (white solid, 34.5 mg, 53%). $^1$H NMR (700 MHz, $CDCl_3$) δ 10.90 (s, 1H), 9.23 (s, 1H), 7.80 (d, *J* = 8.2 Hz, 2H), 7.62 (d, *J* = 8.2 Hz, 2H), 7.22 (s, 1H), 6.90 (s, 1H), 4.03 (d, *J* = 14.6 Hz, 1H), 3.73 (d, *J* = 14.4 Hz, 1H), 3.67 (d, *J* = 5.3 Hz, 8H), 3.62 (dd, *J* = 18.4, 5.0 Hz, 4H), 3.51 (t, *J* = 5.3 Hz, 2H), 3.30 – 3.27 (m, 2H), 3.11 (d, *J* = 8.2 Hz, 2H), 2.83 (d, *J* = 11.5 Hz, 1H), 2.74 (t, *J* = 12.0 Hz, 2H), 2.62 (s, 1H), 2.51 (s, 1H), 2.44 (t, *J* = 10.7 Hz, 1H), 2.30 (s, 3H), 2.28 (d, *J* = 9.4 Hz, 1H), 1.42 (s, 9H), 1.19 (d, *J* = 6.4 Hz, 3H).

**8b** was synthesized from **Boc-8b** (30 mg, 43 μmol) using **General Method A**.

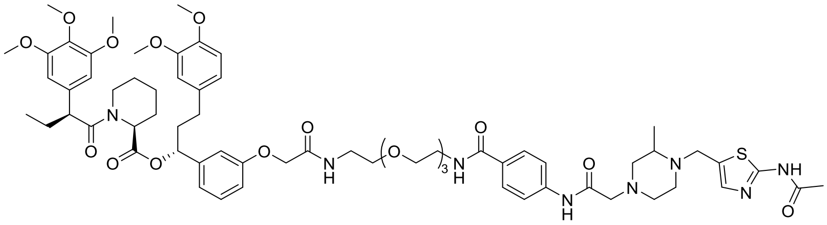

**A2**

**(1R)-1-(3-((1-(4-(2-(4-((2-acetamidothiazol-5-yl)methyl)-3-methylpiperazin-1-yl)acetamido)phenyl)-1,15-dioxo-5,8,11-trioxa-2,14-diazahexadecan-16-yl)oxy)phenyl)-3-(3,4-dimethoxyphenyl)propyl (2S)-1-((S)-2-(3,4,5-trimethoxyphenyl)butanoyl)piperidine-2-carboxylate (A2)**

**A2** was synthesized from **8b** (20 mg, 28 μmol) and **AP1867** (20 mg, 28 μmol) using **General Method B**. pTLC (DCM: MeOH=15:1) to give the product **A2** (white solid, 15.0 mg, 41%). HPLC purity (254 nM)= 95.48%, RT= 15.381 min. $^1$H NMR (700 MHz, $CDCl_3$) δ 11.45 (s, 1H), 9.23 (s, 1H), 7.79 (d, *J* = 8.7 Hz, 2H), 7.61 (dd, *J* = 8.8, 2.9 Hz, 2H), 7.20 (s, 1H), 7.17 (t, *J* = 7.7 Hz, 1H), 7.13 (t, *J* = 5.9 Hz, 1H), 6.79 – 6.75 (m, 3H), 6.66 – 6.62 (m, 3H), 6.40 (s, 2H), 5.61 (dd, *J* = 8.3, 5.4 Hz, 1H), 5.47 – 5.44 (m,

1H), 4.49 – 4.46 (m, 2H), 4.02 (d, *J* = 14.6 Hz, 1H), 3.86 – 3.82 (m, 9H), 3.77 (s, 3H), 3.73 (d, *J* = 14.6 Hz, 1H), 3.68 (s, 5H), 3.63 (d, *J* = 6.3 Hz, 9H), 3.62 – 3.58 (m, 5H), 3.58 – 3.55 (m, 2H), 3.53 – 3.51 (m, 1H), 3.13 – 3.06 (m, 2H), 2.83 – 2.76 (m, 2H), 2.72 (t, *J* = 11.5 Hz, 2H), 2.65 – 2.60 (m, 1H), 2.58 – 2.53 (m, 1H), 2.51 (dd, *J* = 23.6, 7.4 Hz, 2H), 2.47 – 2.41 (m, 2H), 2.29 (s, 3H), 2.26 (s, 1H), 2.07 – 2.03 (m, 2H), 1.93 – 1.90 (m, 1H), 1.72 – 1.67 (m, 3H), 1.62 – 1.59 (m, 1H), 1.43 (dd, *J* = 13.4, 4.3 Hz, 1H), 1.18 (d, *J* = 6.2 Hz, 3H), 0.89 (d, *J* = 4.0 Hz, 3H). $^{13}$C NMR (176 MHz, $CDCl_3$) δ 172.7, 170.6, 168.6, 168.3, 168.0, 166.8, 159.5, 157.3, 153.2, 148.8, 147.3, 142.3, 140.3, 136.5, 135.3, 134.7, 133.3, 129.9, 128.2, 120.2, 120.1, 119.7, 118.8, 113.8, 112.9, 111.6, 111.2, 104.9, 75.7, 70.5, 70.3, 70.2, 69.9, 69.7, 67.3, 61.8, 60.8, 56.3, 55.9, 55.9, 55.8, 53.7, 52.1, 50.8, 49.1, 43.5, 39.7, 38.8, 38.3, 31.3, 29.7, 26.8, 25.3, 23.2, 20.9, 12.6. HRMS (ESI, m/z): calcd for $C_{66}H_{88}N_8O_{16}SNa^+$ $[M+Na]^+$: 1303.5931, found 1303.5930.

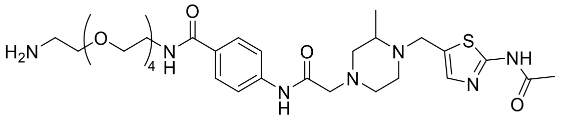

**8c**

**4-(2-(4-((2-acetamidothiazol-5-yl)methyl)-3-methylpiperazin-1-yl)acetamido)-N-(14-amino-3,6,9,12-tetraoxatetradecyl)benzamide (8c)**

Intermediate **8c** was synthesized from **7a** (40 mg, 93 µmol) and BocNH-PEG4-$NH_2$ (31 mg, 93 µmol) using **General Method B**. Flash column chromatography (silica gel, MeOH in DCM: 0-8%) to give the **Boc-8c** (white solid, 65.6 mg, 94%).$^1$H NMR (700 MHz, Chloroform-*d*) δ 10.91 (s, 1H), 9.23 (s, 1H), 7.84 (d, *J* = 8.4 Hz, 2H), 7.61 (d, *J* = 8.2 Hz, 2H), 7.22 (s, 1H), 4.03 (d, *J* = 14.5 Hz, 1H), 3.73 (d, *J* = 14.5 Hz, 1H), 3.66 (d, *J* = 8.2 Hz, 8H), 3.63 (t, *J* = 7.2 Hz, 4H), 3.60 (s, 4H), 3.51 (t, *J* = 5.1 Hz, 2H), 3.28 (q, *J* = 5.4 Hz, 2H), 3.11 (d, *J* = 8.5 Hz, 2H), 2.83 (d, *J* = 11.5 Hz, 1H), 2.73 (t, *J* = 9.8 Hz, 2H), 2.62 (s, 1H), 2.51 (d, *J* = 11.8 Hz, 1H), 2.44 (t, *J* = 10.4 Hz, 1H), 2.30 (s, 3H), 2.28 (d, *J* = 10.8 Hz, 1H), 1.41 (s, 9H), 1.19 (d, *J* = 6.4 Hz, 3H).

**8c** was synthesized from **Boc-8c** (30 mg, 40 µmol) using **General Method A**.

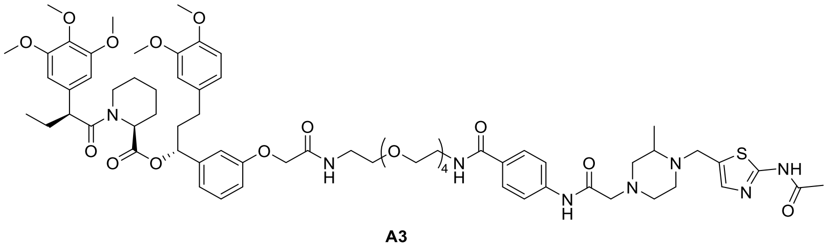

**A3**

**(1R)-1-(3-((1-(4-(2-(4-((2-acetamidothiazol-5-yl)methyl)-3-methylpiperazin-1-yl)acetamido)phenyl)-1,18-dioxo-5,8,11,14-tetraoxa-2,17-diazanonadecan-19-yl)oxy)phenyl)-3-(3,4-dimethoxyphenyl)propyl (2S)-1-((S)-2-(3,4,5-trimethoxyphenyl)butanoyl)piperidine-2-carboxylate (A3)**

**A3** was synthesized from **8c** (20 mg, 27 µmol) and **AP1867** (19 mg, 27 µmol) using **General Method B**. pTLC (DCM: MeOH=15:1) to give the product **A3** (white solid, 20.6 mg, 58%). HPLC purity (254 nM)= 96.75%, RT= 15.411 min. $^1$H NMR (700 MHz, $CDCl_3$) δ 11.70 (s, 1H), 9.22 (s, 1H), 7.82 (d, *J* = 8.7 Hz, 2H), 7.60 (dd, *J* = 8.7, 3.2 Hz, 2H), 7.23 (d, *J* = 5.6 Hz, 1H), 7.20 (s, 1H), 7.18 – 7.14 (m, 1H), 6.79 – 6.75 (m,

3H), 6.66 – 6.62 (m, 3H), 6.40 (d, *J* = 4.6 Hz, 2H), 5.61 (dd, *J* = 8.2, 5.4 Hz, 1H), 5.47 – 5.44 (m, 1H), 4.49 – 4.45 (m, 2H), 4.02 (d, *J* = 14.5 Hz, 1H), 3.85 – 3.82 (m, 9H), 3.77 (s, 3H), 3.71 (d, *J* = 14.4 Hz, 1H), 3.67 (s, 5H), 3.64 – 3.60 (m, 14H), 3.59 (d, *J* = 4.8 Hz, 4H), 3.57 – 3.55 (m, 2H), 3.50 (dd, *J* = 5.8, 2.8 Hz, 1H), 3.13 – 3.06 (m, 2H), 2.80 (ddd, *J* = 16.9, 9.9, 3.0 Hz, 2H), 2.72 (t, *J* = 10.5 Hz, 2H), 2.64 – 2.60 (m, 1H), 2.53 (d, *J* = 5.1 Hz, 1H), 2.52 – 2.47 (m, 2H), 2.45 – 2.41 (m, 2H), 2.29 (s, 3H), 2.27 – 2.23 (m, 1H), 2.08 – 2.03 (m, 2H), 1.95 – 1.88 (m, 1H), 1.72 – 1.66 (m, 3H), 1.62 – 1.58 (m, 1H), 1.43 (ddd, *J* = 17.0, 13.7, 9.8 Hz, 1H), 1.17 (d, *J* = 6.1 Hz, 3H), 0.88 (t, *J* = 3.5 Hz, 3H). $^{13}$C NMR (176 MHz, $CDCl_3$) δ 172.7, 170.6, 168.6, 168.4, 168.1, 166.9, 159.7, 157.3, 153.2, 148.8, 147.3, 142.3, 140.3, 136.5, 135.3, 134.5, 133.3, 129.9, 129.8, 128.3, 120.2, 120.1, 119.7, 118.7, 113.9, 112.9, 111.6, 111.2, 104.9, 75.7, 70.4, 70.4, 70.4, 70.1, 70.1, 70.1, 69.9, 67.3, 67.3, 61.8, 60.9, 60.8, 60.8, 60.7, 56.3, 55.9, 55.9, 55.9, 55.9, 55.8, 53.6, 52.1, 50.8, 49.1, 43.5, 39.8, 38.8, 38.3, 31.3, 29.7, 26.8, 25.3, 23.2, 20.9, 12.6. HRMS (ESI, m/z): calcd for $C_{68}H_{92}N_8O_{17}SNa^+$ [M+Na]$^+$: 1347.6193, found 1347.6183.

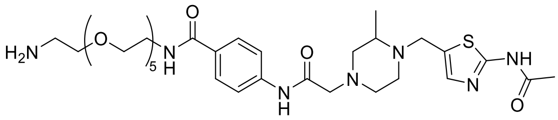

**8d**

**4-(2-(4-((2-acetamidothiazol-5-yl)methyl)-3-methylpiperazin-1-yl)acetamido)-N-(17-amino-3,6,9,12,15-pentaoxaheptadecyl)benzamide (8d)**

Intermediate **8d** was synthesized from **7a** (40 mg, 93 µmol) and BocNH-PEG5-$NH_2$ (35 mg, 93 µmol) using **General Method B**. Flash column chromatography (silica gel, MeOH in DCM: 0-8%) to give the **Boc-8d** (white solid, 68.5 mg, 93%). LC-MS (ESI$^+$) m/z= 794.4.

**8d** was synthesized from **Boc-8d** (30 mg, 38 µmol) using **General Method A**.

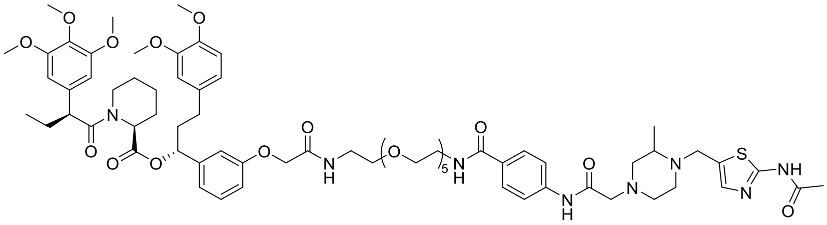

**A4**

**(1R)-1-(3-((1-(4-(2-(4-((2-acetamidothiazol-5-yl)methyl)-3-methylpiperazin-1-yl)acetamido)phenyl)-1,21-dioxo-5,8,11,14,17-pentaoxa-2,20-diazadocosan-22-yl)oxy)phenyl)-3-(3,4-dimethoxyphenyl)propyl (2S)-1-((S)-2-(3,4,5-trimethoxyphenyl)butanoyl)piperidine-2-carboxylate (A4)**

**A4** was synthesized from **8d** (20 mg, 27 µmol) and **AP1867** (19 mg, 27 µmol) using **General Method B**. pTLC (DCM: MeOH=15:1) to give the product **A4** (white solid, 10.3 mg, 27%). HPLC purity (254 nM)= 96.92%, RT= 15.410 min. $^1$H NMR (700 MHz, $CDCl_3$) δ 11.27 (s, 1H), 9.23 (s, 1H), 7.83 (d, *J* = 8.2 Hz, 2H), 7.60 (d, *J* = 8.2 Hz, 2H), 7.32 (t, *J* = 6.4 Hz, 1H), 7.24 (d, *J* = 6.1 Hz, 1H), 7.21 (s, 1H), 7.17 (t, *J* = 8.4 Hz, 1H), 6.77 (d, *J* = 14.0 Hz, 3H), 6.64 (d, *J* = 9.8 Hz, 3H), 6.40 (dd, *J* = 7.2, 1.9 Hz, 2H), 5.61 (t, *J* = 7.0 Hz, 1H), 5.45 (d, *J* = 5.7 Hz, 1H), 4.48 (d, *J* = 10.6 Hz, 2H), 4.03 (d, *J* = 14.6 Hz, 1H), 3.84 (q, *J* = 8.3, 6.6 Hz, 9H), 3.77 (d, *J* = 2.0 Hz, 3H), 3.74 (d, *J* =

14.4 Hz, 1H), 3.67 (d, $J$ = 1.9 Hz, 5H), 3.61 (dt, $J$ = 20.5, 13.1 Hz, 22H), 3.54 (d, $J$ = 15.5 Hz, 2H), 3.51 (d, $J$ = 5.8 Hz, 1H), 3.10 (d, $J$ = 8.0 Hz, 2H), 2.84 – 2.76 (m, 2H), 2.73 (s, 2H), 2.67 – 2.62 (m, 1H), 2.62 – 2.56 (m, 1H), 2.55 – 2.48 (m, 2H), 2.45 (dt, $J$ = 14.9, 8.6 Hz, 2H), 2.29 (d, $J$ = 6.3 Hz, 3H), 2.25 – 2.19 (m, 1H), 2.07 (dd, $J$ = 14.1, 7.0 Hz, 2H), 1.91 (d, $J$ = 7.0 Hz, 1H), 1.71 (dd, $J$ = 13.2, 6.5 Hz, 3H), 1.60 (d, $J$ = 13.7 Hz, 1H), 1.43 (d, $J$ = 11.5 Hz, 1H), 1.18 (d, $J$ = 6.3 Hz, 3H), 0.90 – 0.88 (m, 3H). $^{13}$C NMR (176 MHz, $CDCl_3$) δ 172.7, 170.6, 168.6, 168.4, 168.0, 166.9, 159.4, 157.3, 153.2, 148.8, 147.3, 142.3, 140.2, 136.5, 135.3, 133.3, 129.8, 128.3, 120.2, 120.1, 119.7, 118.8, 114.0, 112.8, 111.6, 111.2, 104.9, 104.5, 75.7, 70.4, 70.1, 70.1, 69.8, 67.3, 67.3, 61.8, 60.9, 60.8, 56.3, 55.9, 55.9, 55.8, 52.1, 50.8, 49.1, 43.5, 39.8, 38.8, 38.3, 31.3, 29.7, 26.8, 25.3, 23.2, 20.9, 12.6. HRMS (ESI, m/z): calcd for $C_{70}H_{96}N_8O_{18}SNa^+$ $[M+Na]^+$: 1391.6456, found 1391.6449.

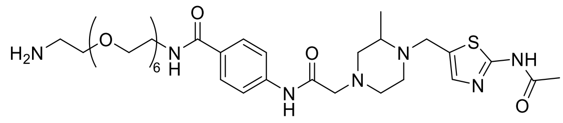

**8e**

**4-(2-(4-((2-acetamidothiazol-5-yl)methyl)-3-methylpiperazin-1-yl)acetamido)-N-(20-amino-3,6,9,12,15,18-hexaoxaicosyl)benzamide (8e)**

Intermediate **8e** was synthesized from **7a** (40 mg, 93 μmol) and BocNH-PEG6-$NH_2$ (39 mg, 93 μmol) using **General Method B**. Flash column chromatography (silica gel, MeOH in DCM: 0-8%) to give the **Boc-8e** (white solid, 75.3 mg, 97%). LC-MS (ESI$^+$) m/z= 838.4.

**8e** was synthesized from **Boc-8e** (30 mg, 36 μmol) using **General Method A**.

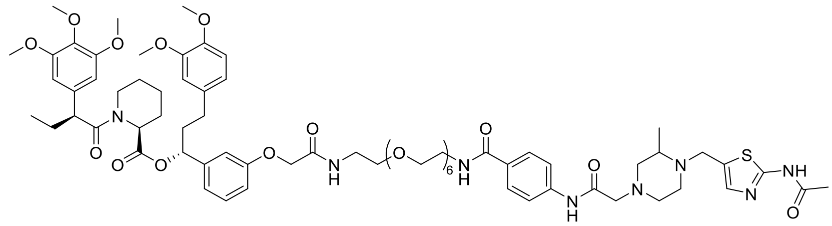

**A5**

**(1R)-1-(3-((1-(4-(2-(4-((2-acetamidothiazol-5-yl)methyl)-3-methylpiperazin-1-yl)acetamido)phenyl)-1,24-dioxo-5,8,11,14,17,20-hexaoxa-2,23-diazapentacosan-25-yl)oxy)phenyl)-3-(3,4-dimethoxyphenyl)propyl (2S)-1-((S)-2-(3,4,5-trimethoxyphenyl)butanoyl)piperidine-2-carboxylate (A5)**

**A5** was synthesized from **8e** (20 mg, 26 μmol) and **AP1867** (18 mg, 26 μmol) using **General Method B.** pTLC (DCM: MeOH=15:1) to give the product **A5** (white solid, 8.8 mg, 24%). HPLC purity (254 nM)= 96.91%, RT= 15.422 min. $^1$H NMR (700 MHz, $CDCl_3$) δ 10.93 (s, 1H), 9.22 (s, 1H), 7.83 (d, $J$ = 8.4 Hz, 2H), 7.61 (d, $J$ = 8.3 Hz, 2H), 7.24 (d, $J$ = 6.1 Hz, 1H), 7.21 (s, 1H), 7.18 (d, $J$ = 8.3 Hz, 1H), 6.80 – 6.76 (m, 3H), 6.64 (dt, $J$ = 9.5, 1.9 Hz, 3H), 6.41 (d, $J$ = 4.8 Hz, 2H), 5.61 (dd, $J$ = 8.2, 5.5 Hz, 1H), 5.47 – 5.45 (m, 1H), 4.51 – 4.48 (m, 2H), 4.02 (d, $J$ = 14.7 Hz, 1H), 3.86 – 3.81 (m, 9H), 3.78 (s, 3H), 3.76 – 3.72 (m, 1H), 3.68 (s, 5H), 3.66 – 3.62 (m, 10H), 3.62 – 3.59 (m, 11H), 3.57 (q, $J$ = 5.5, 4.7 Hz, 7H), 3.53 (s, 1H), 3.10 (d, $J$ = 8.0 Hz, 2H), 2.84 – 2.78 (m, 2H), 2.73 (t, $J$ = 11.3 Hz, 2H), 2.61 (d, $J$ = 6.7 Hz, 1H), 2.55 (d, $J$ = 8.9 Hz, 1H), 2.51 (dd, $J$ = 16.7, 7.9 Hz, 2H), 2.45 (ddd, $J$ = 13.6, 9.4, 6.3 Hz, 2H),

2.29 (d, $J$ = 7.3 Hz, 3H), 2.26 (d, $J$ = 11.1 Hz, 1H), 2.09 – 2.04 (m, 2H), 1.94 – 1.90 (m, 1H), 1.70 (dq, $J$ = 6.7, 3.4, 2.9 Hz, 3H), 1.60 (d, $J$ = 13.4 Hz, 1H), 1.43 (dq, $J$ = 13.2, 5.2, 4.5 Hz, 1H), 1.18 (d, $J$ = 6.2 Hz, 3H), 0.91 – 0.88 (m, 3H). $^{13}$C NMR (176 MHz, $CDCl_3$) δ 172.7, 170.6, 168.6, 168.5, 167.9, 166.9, 166.0, 157.3, 153.1, 148.8, 147.3, 142.3, 140.3, 136.5, 135.3, 134.2, 133.3, 129.5, 128.3, 120.2, 120.1, 119.9, 119.7, 118.8, 113.9, 112.8, 111.6, 111.2, 104.9, 75.7, 70.3, 70.2, 70.0, 70.0, 69.8, 67.8, 67.3, 61.8, 61.7, 60.9, 60.8, 60.7, 56.3, 55.9, 55.9, 55.8, 52.1, 50.8, 49.1, 43.5, 39.7, 38.8, 38.3, 31.3, 30.6, 29.7, 26.8, 25.3, 24.0, 23.2, 20.9, 12.6. HRMS (ESI, m/z): calcd for $C_{72}H_{100}N_8O_{19}SNa^+$ [M+Na]$^+$: 1435.6718, found 1435.6710.

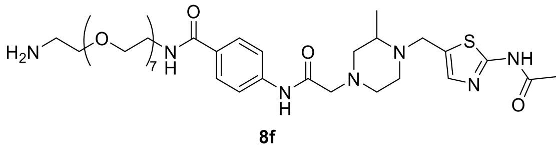

**8f**

**4-(2-(4-((2-acetamidothiazol-5-yl)methyl)-3-methylpiperazin-1-yl)acetamido)-N-(23-amino-3,6,9,12,15,18,21-heptaoxatricosyl)benzamide (8f)**

Intermediate **8f** was synthesized from **7a** (40 mg, 93 μmol) and BocNH-PEG7-$NH_2$ (45 mg, 93 μmol) using **General Method B**. Flash column chromatography (silica gel, MeOH in DCM: 0-8%) to give the **Boc-8f** (white solid, 77.2 mg, 94%). LC-MS (ESI$^+$) m/z= 882.5.

**8f** was synthesized from **Boc-8f** (30 mg, 36 μmol) using **General Method A**.

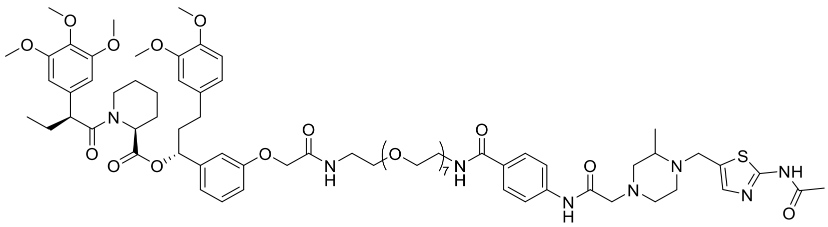

**A6**

**(1R)-1-(3-((1-(4-(2-(4-((2-acetamidothiazol-5-yl)methyl)-3-methylpiperazin-1-yl)acetamido)phenyl)-1,27-dioxo-5,8,11,14,17,20,23-heptaoxa-2,26-diazaoctacosan-28-yl)oxy)phenyl)-3-(3,4-dimethoxyphenyl)propyl (2S)-1-((S)-2-(3,4,5-trimethoxyphenyl)butanoyl)piperidine-2-carboxylate (A6)**

**A6** was synthesized from **8f** (20 mg, 24 μmol) and **AP1867** (17 mg, 24 μmol) using **General Method B.** pTLC (DCM: MeOH=15:1) to give the product **A6** (white solid, 7.7 mg, 21%). HPLC purity (254 nM)= 95.92%, RT= 15.446 min. $^1$H NMR (700 MHz, $CDCl_3$) δ 10.95 (s, 1H), 9.22 (s, 1H), 7.84 (d, $J$ = 8.4 Hz, 2H), 7.60 (d, $J$ = 8.4 Hz, 2H), 7.30 (s, 1H), 7.21 (s, 1H), 7.17 (t, $J$ = 7.8 Hz, 1H), 6.81 – 6.75 (m, 3H), 6.64 (dt, $J$ = 8.9, 1.9 Hz, 3H), 6.41 (d, $J$ = 5.8 Hz, 2H), 5.61 (dd, $J$ = 8.2, 5.5 Hz, 1H), 5.46 (d, $J$ = 5.5 Hz, 1H), 4.51 – 4.47 (m, 2H), 4.02 (d, $J$ = 14.6 Hz, 1H), 3.86 – 3.81 (m, 9H), 3.78 (s, 3H), 3.74 (d, $J$ = 14.4 Hz, 1H), 3.68 (s, 5H), 3.66 – 3.62 (m, 10H), 3.62 – 3.57 (m, 22H), 3.53 (d, $J$ = 6.6 Hz, 1H), 3.14 – 3.07 (m, 2H), 2.82 – 2.76 (m, 2H), 2.73 (s, 2H), 2.61 (d, $J$ = 15.9 Hz, 1H), 2.55 (d, $J$ = 8.8 Hz, 1H), 2.53 – 2.47 (m, 2H), 2.45 (ddd, $J$ = 14.0, 9.2, 6.3 Hz, 2H), 2.28 (s, 3H), 2.22 (t, $J$ = 7.6 Hz, 1H), 2.09 – 2.04 (m, 2H), 1.94 – 1.91 (m, 1H), 1.71 (dt, $J$ = 13.7, 6.8 Hz, 3H), 1.60 (d, $J$ = 16.1 Hz, 1H), 1.44 (dt, $J$ = 9.5, 5.2 Hz, 1H), 1.18 (d, $J$ = 6.2 Hz, 3H), 0.89 (t, $J$ = 7.2 Hz, 3H). $^{13}$C NMR (176 MHz, $CDCl_3$) δ 172.7, 170.6, 168.6, 168.3, 167.9, 166.8, 159.1,

157.3, 153.2, 148.8, 147.3, 142.3, 140.2, 136.5, 135.3, 133.3, 129.5, 128.4, 120.2, 120.1, 119.9, 119.7, 118.7, 114.0, 112.7, 111.6, 111.2, 104.9, 75.7, 70.5, 70.5, 70.4, 70.2, 70.1, 70.1, 69.7, 67.8, 67.3, 61.8, 60.9, 60.8, 56.3, 56.0, 55.9, 55.9, 55.8, 52.1, 50.8, 43.5, 39.8, 38.8, 38.3, 31.3, 30.6, 29.7, 26.8, 25.3, 24.0, 23.2, 20.9. HRMS (ESI, m/z): calcd for $C_{74}H_{104}N_8O_{20}SNa^+$ $[M+Na]^+$: 1479.6980, found 1479.6985.

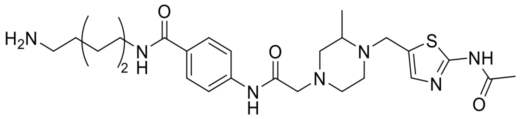

**8g**

**4-(2-(4-((2-acetamidothiazol-5-yl)methyl)-3-methylpiperazin-1-yl)acetamido)-N-(4-aminobutyl)benzamide (8g)**

Intermediate **8g** was synthesized from **7a** (20 mg, 46 μmol) and BocNH-C4-$NH_2$ (9 mg, 46 μmol) using **General Method B**. Flash column chromatography (silica gel, MeOH in DCM: 0-8%) to give the **Boc-8g** (white solid, 13.0 mg, 47%). $^1H$ NMR (700 MHz, $CDCl_3$) δ 10.39 (s, 1H), 9.23 (s, 1H), 7.78 (d, *J* = 8.3 Hz, 2H), 7.62 (d, *J* = 8.6 Hz, 2H), 7.22 (s, 1H), 6.47 (s, 1H), 4.03 (d, *J* = 14.5 Hz, 1H), 3.73 (d, *J* = 14.5 Hz, 1H), 3.49 – 3.46 (m, 2H), 3.17 (d, *J* = 6.8 Hz, 2H), 3.11 (d, *J* = 7.3 Hz, 2H), 2.82 (d, *J* = 11.9 Hz, 1H), 2.73 (t, *J* = 12.0 Hz, 2H), 2.62 (s, 1H), 2.51 (d, *J* = 10.5 Hz, 1H), 2.44 (t, *J* = 10.5 Hz, 1H), 2.30 (s, 3H), 2.27 (s, 1H), 1.58 (s, 4H), 1.44 (s, 9H), 1.19 (d, *J* = 6.2 Hz, 3H).

**8g** was synthesized from **Boc-8g** (13 mg, 22 μmol) using **General Method A**.

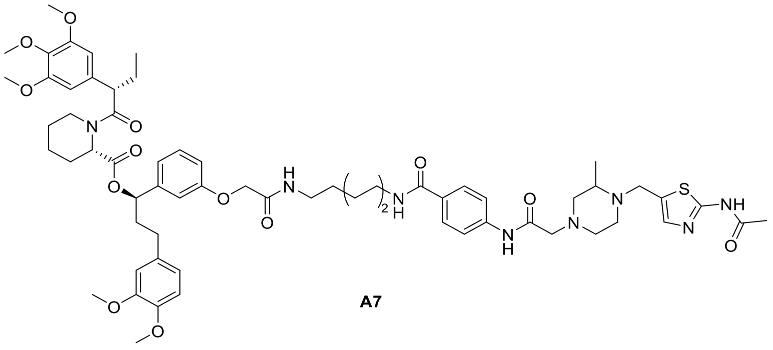

**A7**

**(1R)-1-(3-(2-((4-(4-(2-(4-((2-acetamidothiazol-5-yl)methyl)-3-methylpiperazin-1-yl)acetamido)benzamido)butyl)amino)-2-oxoethoxy)phenyl)-3-(3,4-dimethoxyphenyl)propyl (2S)-1-((S)-2-(3,4,5-trimethoxyphenyl)butanoyl)piperidine-2-carboxylate (A7)**

**A7** was synthesized from **8g** (13 mg, 22 μmol) and **AP1867** (15 mg, 22 μmol) using **General Method B.** pTLC (DCM: MeOH=15:1) to give the product **A7** (white solid, 15.4 mg, 60%). HPLC purity (254 nM)= 97.06%, RT= 15.583 min. $^1H$ NMR (700 MHz, $CDCl_3$) δ 11.78 (s, 1H), 9.22 (s, 1H), 7.78 (d, *J* = 8.3 Hz, 2H), 7.60 (d, *J* = 8.3 Hz, 2H), 7.21 (s, 1H), 7.18 (t, *J* = 7.8 Hz, 1H), 6.93 – 6.90 (m, 1H), 6.77 (h, *J* = 5.8, 3.9 Hz, 3H), 6.69 – 6.62 (m, 3H), 6.39 (d, *J* = 16.1 Hz, 2H), 5.60 (dd, *J* = 8.4, 5.4 Hz, 1H), 5.44 (d, *J* = 5.2 Hz, 1H), 4.49 (d, *J* = 2.9 Hz, 2H), 4.03 (d, *J* = 14.5 Hz, 1H), 3.87 – 3.80 (m, 9H), 3.77 (s, 3H), 3.72 (d, *J* = 14.8 Hz, 1H), 3.65 (s, 5H), 3.59 (t, *J* = 7.2 Hz, 1H), 3.40 (dp, *J* = 25.8, 6.7 Hz, 4H), 3.13 – 3.07 (m, 2H), 2.84 – 2.77 (m, 2H), 2.72 (t, *J* = 11.0 Hz, 2H), 2.62 (s, 1H), 2.59 – 2.54 (m, 1H), 2.54 – 2.46 (m, 2H), 2.47 – 2.40

(m, 2H), 2.30 (d, *J* = 3.1 Hz, 3H), 2.27 (d, *J* = 12.8 Hz, 1H), 2.06 (dp, *J* = 21.1, 7.2, 6.0 Hz, 2H), 1.94 – 1.91 (m, 1H), 1.70 (dt, *J* = 10.1, 6.8 Hz, 3H), 1.66 – 1.58 (m, 5H), 1.43 (ddd, *J* = 17.3, 13.4, 9.3 Hz, 1H), 1.18 (d, *J* = 6.1 Hz, 3H), 0.91 – 0.88 (m, 3H). $^{13}$C NMR (176 MHz, $CDCl_3$) δ 172.8, 170.6, 168.6, 168.5, 168.0, 166.9, 159.7, 157.3, 153.1, 148.8, 147.3, 142.4, 140.2, 136.5, 135.3, 134.6, 133.3, 130.0, 129.8, 128.1, 120.2, 119.8, 118.8, 113.5, 112.9, 111.6, 111.2, 104.9, 75.7, 67.3, 61.8, 60.9, 60.8, 60.7, 56.3, 55.9, 55.9, 55.8, 54.2, 53.7, 52.1, 50.8, 49.2, 43.5, 39.6, 38.6, 38.2, 31.3, 29.7, 27.2, 26.8, 26.6, 25.3, 23.2, 20.9, 12.6. HRMS (ESI, m/z): calcd for $C_{62}H_{80}N_8O_{13}SNa^+$ $[M+Na]^+$: 1199.5458, found 1199.5450.

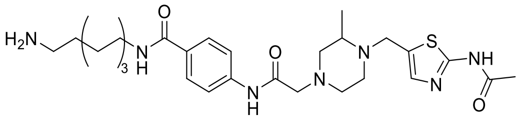

**8h**

**4-(2-(4-((2-acetamidothiazol-5-yl)methyl)-3-methylpiperazin-1-yl)acetamido)-N-(5-aminopentyl)benzamide (8h)**

Intermediate **8h** was synthesized from **7a** (20 mg, 46 µmol) and BocNH-C5-$NH_2$ (9 mg, 46 µmol) using **General Method B**. Flash column chromatography (silica gel, MeOH in DCM: 0-9%) to give the **Boc-8h** (white solid, 13.0 mg, 46%).$^1$H NMR (700 MHz, $CDCl_3$) δ 10.43 (s, 1H), 9.23 (s, 1H), 7.75 (d, *J* = 8.2 Hz, 2H), 7.62 (d, *J* = 8.7 Hz, 2H), 7.22 (s, 1H), 4.03 (d, *J* = 14.5 Hz, 1H), 3.73 (d, *J* = 14.6 Hz, 1H), 3.44 (q, *J* = 6.7 Hz, 2H), 3.12 (dd, *J* = 13.9, 9.9 Hz, 4H), 2.99 (s, 2H), 2.82 (dd, *J* = 11.5, 3.5 Hz, 1H), 2.73 (t, *J* = 12.2 Hz, 2H), 2.62 (s, 1H), 2.52 (d, *J* = 10.5 Hz, 1H), 2.44 (t, *J* = 10.5 Hz, 1H), 2.30 (s, 3H), 2.27 (s, 1H), 1.64 (d, *J* = 7.4 Hz, 2H), 1.53 (t, *J* = 7.4 Hz, 2H), 1.42 (s, 9H), 1.28 (s, 2H), 1.19 (d, *J* = 6.2 Hz, 3H).

**8h** was synthesized from **Boc-8h** (13 mg, 21 µmol) using **General Method A**.

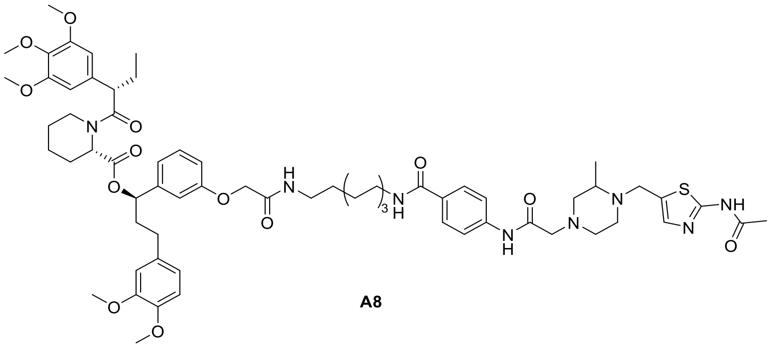

**A8**

**(1R)-1-(3-(2-((5-(4-(2-(4-((2-acetamidothiazol-5-yl)methyl)-3-methylpiperazin-1-yl)acetamido)benzamido)pentyl)amino)-2-oxoethoxy)phenyl)-3-(3,4-dimethoxyphenyl)propyl (2S)-1-((S)-2-(3,4,5-trimethoxyphenyl)butanoyl)piperidine-2-carboxylate (A8)**

**A8** was synthesized from **8h** (13 mg, 21 µmol) and **AP1867** (15 mg, 22 µmol) using **General Method B.** pTLC (DCM: MeOH=15:1) to give the product **A8** (white solid, 16.5 mg, 65%). HPLC purity (254 nM)= 96.42%, RT= 15.787 min. $^1$H NMR (700 MHz, $CDCl_3$) δ 11.86 (s, 1H), 9.22 (s, 1H), 7.77 (d, *J* = 8.2 Hz, 2H), 7.62 – 7.57 (m, 2H), 7.22 (s, 1H), 7.17 (t, *J* = 7.9 Hz, 1H), 6.81 (d, *J* = 6.3 Hz, 1H), 6.77 – 6.73 (m, 3H), 6.68 – 6.62 (m, 3H), 6.51 (s, 1H), 6.39 (d, *J* = 15.2 Hz, 2H), 5.60 (dd, *J* = 8.3, 5.3 Hz,

1H), 5.45 (d, *J* = 5.6 Hz, 1H), 4.47 – 4.44 (m, 2H), 4.04 (d, *J* = 14.5 Hz, 1H), 3.85 – 3.81 (m, 9H), 3.77 (s, 3H), 3.73 (d, *J* = 14.6 Hz, 1H), 3.65 (s, 5H), 3.59 (d, *J* = 7.2 Hz, 1H), 3.39 (dq, *J* = 14.1, 7.5, 6.9 Hz, 2H), 3.34 (p, *J* = 7.0 Hz, 2H), 3.10 (d, *J* = 7.5 Hz, 2H), 2.84 – 2.77 (m, 2H), 2.72 (d, *J* = 11.1 Hz, 2H), 2.63 (s, 1H), 2.59 – 2.54 (m, 1H), 2.54 – 2.47 (m, 2H), 2.47 – 2.40 (m, 2H), 2.32 – 2.29 (m, 3H), 2.29 – 2.25 (m, 1H), 2.06 (ddd, *J* = 21.3, 10.3, 4.4 Hz, 2H), 1.91 (dd, *J* = 14.7, 6.7 Hz, 1H), 1.70 (d, *J* = 14.0 Hz, 3H), 1.59 (tt, *J* = 14.6, 7.0 Hz, 5H), 1.44 – 1.40 (m, 1H), 1.36 (q, *J* = 8.0 Hz, 2H), 1.18 (d, *J* = 6.1 Hz, 3H), 0.90 – 0.88 (m, 3H). $^{13}$C NMR (176 MHz, $CDCl_3$) δ 172.8, 170.6, 168.6, 168.4, 168.0, 166.9, 159.8, 157.3, 153.1, 148.8, 147.3, 142.4, 140.2, 136.4, 135.3, 134.7, 133.3, 130.0, 129.8, 128.1, 120.2, 119.8, 118.8, 113.4, 113.0, 111.6, 111.2, 104.8, 75.7, 67.3, 61.8, 60.9, 60.8, 60.7, 56.3, 55.9, 55.8, 54.2, 53.6, 52.1, 50.8, 49.2, 43.5, 39.8, 38.7, 38.3, 31.3, 29.3, 29.0, 26.8, 25.3, 24.0, 23.2, 20.9, 12.6. HRMS (ESI, m/z): calcd for $C_{63}H_{82}N_8O_{13}SNa^+$ $[M+Na]^+$: 1213.5614, found 1213.5596.

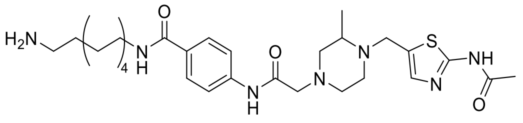

8i

**4-(2-(4-((2-acetamidothiazol-5-yl)methyl)-3-methylpiperazin-1-yl)acetamido)-N-(6-aminohexyl)benzamide (8i)**

Intermediate **8i** was synthesized from **7a** (40 mg, 93 μmol) and BocNH-C6-$NH_2$ (20 mg, 93 μmol) using **General Method B**. Flash column chromatography (silica gel, MeOH in DCM: 0-10%) to give the **Boc-8i** (white solid, 36.9 mg, 63%).$^1$H NMR (700 MHz, $CDCl_3$) δ 11.17 (s, 1H), 9.23 (s, 1H), 7.78 (d, *J* = 8.2 Hz, 2H), 7.62 (d, *J* = 8.3 Hz, 2H), 7.22 (s, 1H), 6.38 (d, *J* = 6.1 Hz, 1H), 4.04 (d, *J* = 14.5 Hz, 1H), 3.73 (d, *J* = 14.5 Hz, 1H), 3.43 (q, *J* = 6.7 Hz, 2H), 3.12 (dd, *J* = 15.6, 8.2 Hz, 4H), 2.83 (dt, *J* = 11.4, 3.2 Hz, 1H), 2.74 (d, *J* = 11.9 Hz, 2H), 2.62 (s, 1H), 2.51 (t, *J* = 10.4 Hz, 1H), 2.44 (t, *J* = 10.4 Hz, 1H), 2.31 (s, 3H), 2.27 (t, *J* = 10.4 Hz, 1H), 1.62 – 1.59 (m, 2H), 1.48 (q, *J* = 7.2 Hz, 4H), 1.43 (s, 9H), 1.36 (q, *J* = 8.3, 7.8 Hz, 3H), 1.19 (d, *J* = 6.2 Hz, 3H).

**8i** was synthesized from **Boc-8i** (20 mg, 32 μmol) using **General Method A**.

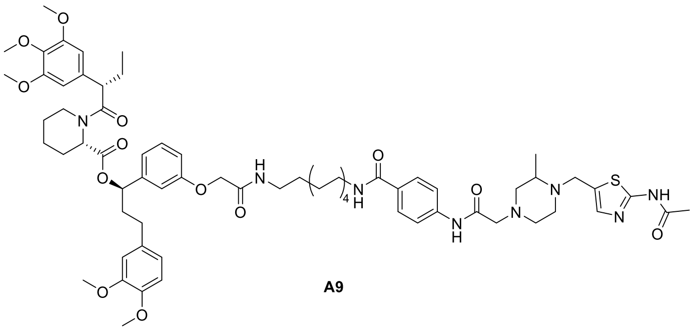

A9

**(1R)-1-(3-(2-((6-(4-(2-(4-((2-acetamidothiazol-5-yl)methyl)-3-methylpiperazin-1-yl)acetamido)benzamido)hexyl)amino)-2-oxoethoxy)phenyl)-3-(3,4-dimethoxyphenyl)propyl (2S)-1-((S)-2-(3,4,5-trimethoxyphenyl)butanoyl)piperidine-2-carboxylate (A9)**

**A9** was synthesized from **8i** (20 mg, 32 μmol) and **AP1867** (22 mg, 32 μmol) using **General Method B.** pTLC (DCM: MeOH=15:1) to give the product **A9** (white solid,

15.3 mg, 40%). HPLC purity (254 nM)= 98.75%, RT= 16.021 min. $^{1}$H NMR (700 MHz, $CDCl_3$) δ 11.76 (s, 1H), 9.22 (s, 1H), 7.77 (dd, *J* = 8.8, 2.1 Hz, 2H), 7.61 (dd, *J* = 8.7, 1.8 Hz, 2H), 7.21 (s, 1H), 7.18 (dd, *J* = 9.0, 7.6 Hz, 1H), 6.78 – 6.75 (m, 3H), 6.67 (dd, *J* = 7.9, 1.5 Hz, 1H), 6.65 – 6.62 (m, 2H), 6.47 (t, *J* = 5.6 Hz, 1H), 6.40 (d, *J* = 12.9 Hz, 2H), 5.61 (dd, *J* = 8.3, 5.3 Hz, 1H), 5.46 – 5.44 (m, 1H), 4.48 (d, *J* = 4.5 Hz, 2H), 4.03 (d, *J* = 14.5 Hz, 1H), 3.86 – 3.81 (m, 9H), 3.77 (s, 3H), 3.71 (d, *J* = 14.5 Hz, 1H), 3.66 (s, 5H), 3.60 – 3.57 (m, 1H), 3.38 (q, *J* = 6.7 Hz, 2H), 3.34 (dq, *J* = 13.1, 6.8 Hz, 2H), 3.10 (d, *J* = 7.0 Hz, 2H), 2.84 – 2.77 (m, 2H), 2.72 (t, *J* = 11.1 Hz, 2H), 2.65 – 2.60 (m, 1H), 2.59 – 2.54 (m, 1H), 2.54 – 2.47 (m, 2H), 2.47 – 2.40 (m, 2H), 2.30 (d, *J* = 3.0 Hz, 3H), 2.28 – 2.24 (m, 1H), 2.10 – 2.03 (m, 2H), 1.95 – 1.89 (m, 1H), 1.71 (dh, *J* = 14.7, 4.2, 2.9 Hz, 3H), 1.62 – 1.59 (m, 1H), 1.55 (dq, *J* = 14.1, 6.9 Hz, 4H), 1.46 – 1.41 (m, 1H), 1.39 – 1.35 (m, 2H), 1.34 (dd, *J* = 8.1, 5.7 Hz, 2H), 1.18 (d, *J* = 6.2 Hz, 3H), 0.89 (d, *J* = 6.5 Hz, 3H). $^{13}$C NMR (176 MHz, $CDCl_3$) δ 172.7, 170.6, 168.6, 168.2, 168.0, 166.8, 159.7, 157.3, 153.2, 148.8, 147.3, 142.3, 140.2, 136.5, 135.3, 134.6, 133.3, 130.1, 129.8, 128.1, 128.0, 120.2, 120.1, 119.8, 118.8, 113.5, 113.0, 111.6, 111.2, 104.8, 75.6, 67.3, 61.8, 60.9, 60.8, 60.7, 56.3, 55.9, 55.9, 55.8, 53.7, 52.1, 50.8, 49.2, 43.5, 39.7, 38.7, 38.3, 31.3, 29.5, 26.8, 26.2, 26.1, 25.3, 23.2, 20.9, 12.6. HRMS (ESI, m/z): calcd for $C_{64}H_{84}N_8O_{13}SNa^+$ $[M+Na]^+$: 1227.5771, found 1227.5774.

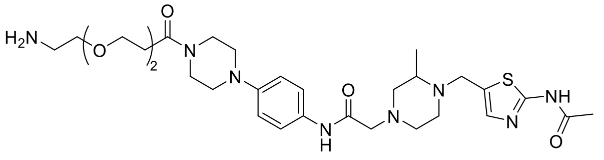


**8j**

**2-(4-((2-acetamidothiazol-5-yl)methyl)-3-methylpiperazin-1-yl)-N-(4-(piperazin-1-yl)phenyl)acetamide (8j)**

Intermediate **8j** was synthesized from **7b** (50 mg, 88 μmol) and BocNH-PEG2-$CH_2CH_2COOH$ (24 mg, 88 μmol) using **General Method B**. Flash column chromatography (silica gel, MeOH in DCM: 0-10%) to give **Boc-8j** (white solid, 60.6 mg, 94%). $^{1}$H NMR (700 MHz, $CDCl_3$) δ 11.17 (s, 1H), 8.93 (s, 1H), 7.45 (d, *J* = 8.4 Hz, 2H), 7.21 (s, 1H), 6.89 (d, *J* = 8.5 Hz, 2H), 4.03 (d, *J* = 14.6 Hz, 1H), 3.82 (t, *J* = 6.6 Hz, 2H), 3.78 (t, *J* = 5.2 Hz, 2H), 3.71 (d, *J* = 14.3 Hz, 1H), 3.64 (t, *J* = 5.0 Hz, 2H), 3.62 (d, *J* = 4.9 Hz, 2H), 3.59 (d, *J* = 4.9 Hz, 2H), 3.51 (d, *J* = 5.3 Hz, 2H), 3.48 (q, *J* = 7.0 Hz, 2H), 3.30 (q, *J* = 5.4 Hz, 2H), 3.12 (d, *J* = 5.4 Hz, 2H), 3.10 – 3.06 (m, 4H), 2.81 (d, *J* = 11.5 Hz, 1H), 2.72 (d, *J* = 10.8 Hz, 2H), 2.68 (t, *J* = 6.6 Hz, 2H), 2.61 (s, 1H), 2.49 (t, *J* = 10.5 Hz, 1H), 2.42 (t, *J* = 10.5 Hz, 1H), 2.30 (s, 3H), 2.26 (d, *J* = 10.3 Hz, 1H), 1.43 (s, 9H), 1.18 (d, *J* = 6.2 Hz, 3H).

**8j** was synthesized from **Boc-8j** (20 mg, 27 μmol) using **General Method A**.

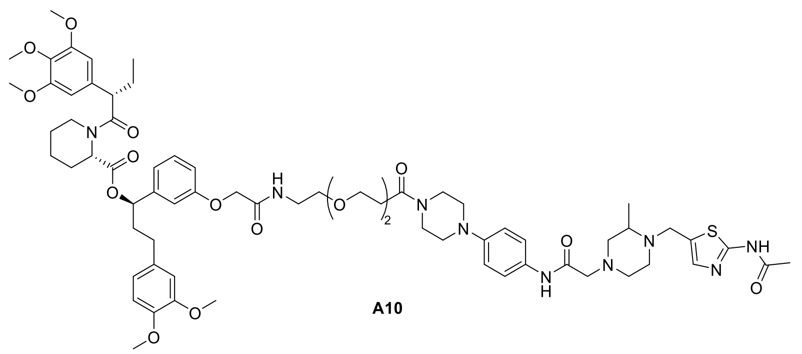

**A10**

**(1R)-1-(3-(2-((2-(2-(3-(4-(4-(2-(4-((2-acetamidothiazol-5-yl)methyl)-3-methylpiperazin-1-yl)acetamido)phenyl)piperazin-1-yl)-3-oxopropoxy)ethoxy)ethyl)amino)-2-oxoethoxy)phenyl)-3-(3,4-dimethoxyphenyl)propyl (2S)-1-((S)-2-(3,4,5-trimethoxyphenyl)butanoyl)piperidine-2-carboxylate (A10)**

**A10** was synthesized from **8j** (20 mg, 27 μmol) and **AP1867** (19 mg, 27 μmol) using **General Method B.** pTLC (DCM: MeOH=15:1) to give the product **A10** (white solid, 15.0 mg, 42%). HPLC purity (254 nM)= 96.14%, RT= 15.277 min. $^{1}$H NMR (700 MHz, $CDCl_3$) δ 11.75 (s, 1H), 8.92 (s, 1H), 7.46 – 7.42 (m, 2H), 7.20 (s, 1H), 7.19 – 7.16 (m, 1H), 7.11 (t, *J* = 5.8 Hz, 1H), 6.88 – 6.86 (m, 2H), 6.80 – 6.75 (m, 3H), 6.66 – 6.62 (m, 3H), 6.40 (s, 2H), 5.61 (dd, *J* = 8.2, 5.4 Hz, 1H), 5.47 – 5.45 (m, 1H), 4.50 – 4.47 (m, 2H), 4.03 (d, *J* = 14.5 Hz, 1H), 3.85 – 3.82 (m, 9H), 3.80 (t, *J* = 6.7 Hz, 3H), 3.77 (s, 3H), 3.74 (t, *J* = 5.3 Hz, 2H), 3.67 (s, 5H), 3.60 (p, *J* = 3.6, 2.9 Hz, 6H), 3.59 – 3.55 (m, 4H), 3.54 – 3.53 (m, 1H), 3.08 (t, *J* = 5.6 Hz, 4H), 3.06 (t, *J* = 5.2 Hz, 2H), 2.81 (dt, *J* = 9.6, 2.8 Hz, 1H), 2.77 (dd, *J* = 13.4, 3.0 Hz, 1H), 2.74 – 2.70 (m, 2H), 2.65 (t, *J* = 6.6 Hz, 2H), 2.60 (dt, *J* = 9.3, 5.5 Hz, 1H), 2.54 (ddd, *J* = 14.7, 9.8, 5.5 Hz, 1H), 2.48 (dd, *J* = 13.5, 5.1 Hz, 2H), 2.45 – 2.37 (m, 2H), 2.31 – 2.30 (m, 3H), 2.25 (s, 1H), 2.08 (ddd, *J* = 17.5, 12.4, 6.7 Hz, 2H), 1.94 – 1.89 (m, 1H), 1.73 – 1.67 (m, 3H), 1.59 (d, *J* = 14.1 Hz, 1H), 1.42 (tt, *J* = 13.4, 4.1 Hz, 1H), 1.17 (d, *J* = 6.2 Hz, 3H), 0.91 – 0.88 (m, 3H). $^{13}$C NMR (176 MHz, $CDCl_3$) δ 172.7, 170.6, 169.4, 168.2, 168.1, 167.9, 159.7, 157.3, 153.2, 148.8, 147.8, 147.3, 142.3, 136.5, 135.3, 134.5, 133.3, 130.9, 129.8, 120.8, 120.2, 120.1, 119.7, 117.3, 117.3, 113.9, 112.8, 111.6, 111.2, 104.9, 75.7, 70.4, 70.3, 69.7, 67.3, 67.3, 61.8, 60.9, 60.8, 60.7, 56.3, 56.0, 55.9, 55.9, 55.9, 55.8, 53.7, 52.0, 50.8, 50.2, 49.8, 49.2, 45.5, 43.5, 41.4, 38.8, 38.3, 33.5, 31.3, 29.7, 26.8, 25.3, 23.2, 21.0, 12.6. HRMS (ESI, m/z): calcd for $C_{68}H_{91}N_9O_{15}SNa^+$ $[M+Na]^+$: 1328.6248, found 1328.6255.

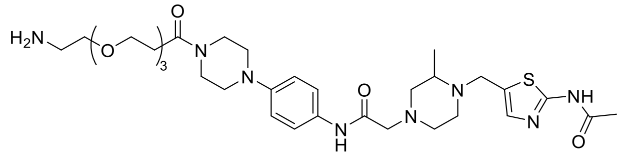

**8k**

**2-(4-((2-acetamidothiazol-5-yl)methyl)-3-methylpiperazin-1-yl)-N-(4-(4-(3-(2-(2-(2-aminoethoxy)ethoxy)ethoxy)propanoyl)piperazin-1-yl)phenyl)acetamide (8k)**

Intermediate **8k** was synthesized from **7b** (40 mg, 70 μmol) and BocNH-PEG3-$CH_2CH_2COOH$ (23 mg, 70 μmol) using **General Method B**. Flash column chromatography (silica gel, MeOH in DCM: 0-10%) to give **Boc-8k** (white solid, 42.0 mg, 77%). $^{1}$H NMR (700 MHz, $CDCl_3$) δ 11.13 (s, 1H), 8.92 (s, 1H), 7.45 (d, *J* = 8.5 Hz, 2H), 7.26 – 7.21 (m, 1H), 6.89 (d, *J* = 8.6 Hz, 2H), 4.07 (s, 1H), 3.81 (t, *J* = 6.7 Hz, 2H),

3.77 (t, *J* = 5.2 Hz, 2H), 3.67 (dt, *J* = 6.5, 3.2 Hz, 1H), 3.64 (t, *J* = 4.1 Hz, 5H), 3.60 (q, *J* = 5.0, 4.4 Hz, 4H), 3.52 (t, *J* = 5.2 Hz, 2H), 3.48 (q, *J* = 7.0 Hz, 1H), 3.30 (q, *J* = 5.4 Hz, 2H), 3.12 (q, *J* = 6.7, 5.0 Hz, 3H), 3.09 (q, *J* = 6.6, 5.2 Hz, 3H), 2.85 (s, 1H), 2.76 (d, *J* = 17.5 Hz, 2H), 2.68 (t, *J* = 6.7 Hz, 3H), 2.51 (d, *J* = 51.4 Hz, 3H), 2.31 (s, 3H), 1.43 (d, *J* = 2.4 Hz, 9H), 1.24 – 1.18 (m, 3H).

**8k** was synthesized from **Boc-8k** (20 mg, 26 μmol) using **General Method A**.

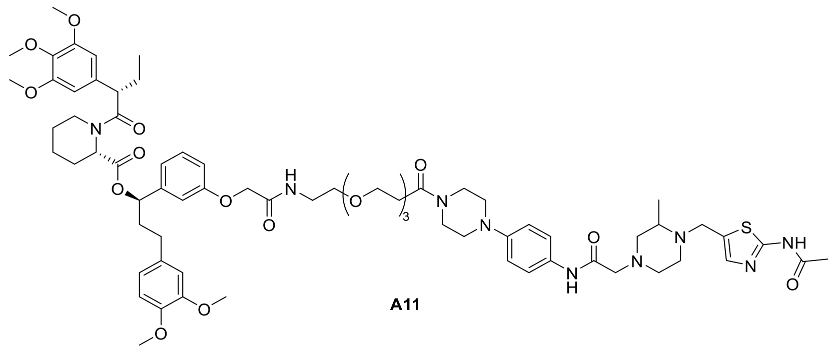

**A11**

**(1R)-1-(3-((15-(4-(4-(2-(4-((2-acetamidothiazol-5-yl)methyl)-3-methylpiperazin-1-yl)acetamido)phenyl)piperazin-1-yl)-2,15-dioxo-6,9,12-trioxa-3-azapentadecyl)oxy)phenyl)-3-(3,4-dimethoxyphenyl)propyl (2S)-1-((S)-2-(3,4,5-trimethoxyphenyl)butanoyl)piperidine-2-carboxylate (A11)**

**A11** was synthesized from **8k** (20 mg, 26 μmol) and **AP1867** (18 mg, 26 μmol) using **General Method B.** pTLC (DCM: MeOH=15:1) to give the product **A11** (white solid, 17.2 mg, 49%). HPLC purity (254 nM)= 95.04%, RT= 15.305 min. $^1$H NMR (700 MHz, $CDCl_3$) δ 11.77 (s, 1H), 8.92 (s, 1H), 7.44 (d, *J* = 8.9 Hz, 2H), 7.20 (s, 1H), 7.19 – 7.16 (m, 1H), 7.13 (t, *J* = 5.7 Hz, 1H), 6.87 (d, *J* = 9.0 Hz, 2H), 6.80 – 6.74 (m, 3H), 6.66 – 6.62 (m, 3H), 6.40 (d, *J* = 3.8 Hz, 2H), 5.61 (dd, *J* = 8.2, 5.5 Hz, 1H), 5.47 – 5.44 (m, 1H), 4.51 – 4.47 (m, 2H), 4.02 (d, *J* = 14.6 Hz, 1H), 3.86 – 3.81 (m, 9H), 3.80 – 3.78 (m, 2H), 3.77 (s, 3H), 3.75 (t, *J* = 5.3 Hz, 2H), 3.67 (s, 5H), 3.60 (dd, *J* = 11.1, 3.6 Hz, 10H), 3.57 (t, *J* = 5.6 Hz, 3H), 3.54 (dd, *J* = 4.1, 2.1 Hz, 1H), 3.11 – 3.08 (m, 3H), 3.08 – 3.04 (m, 3H), 2.82 – 2.75 (m, 2H), 2.74 – 2.69 (m, 2H), 2.65 (t, *J* = 6.7 Hz, 2H), 2.62 – 2.57 (m, 1H), 2.54 (td, *J* = 9.5, 4.8 Hz, 1H), 2.47 (td, *J* = 10.4, 9.5, 6.0 Hz, 2H), 2.45 – 2.37 (m, 2H), 2.30 (s, 3H), 2.23 (dd, *J* = 16.0, 5.5 Hz, 1H), 2.07 (dtd, *J* = 14.6, 9.3, 4.8 Hz, 2H), 1.92 – 1.89 (m, 1H), 1.73 – 1.66 (m, 3H), 1.62 – 1.56 (m, 1H), 1.43 (dtd, *J* = 12.9, 8.8, 4.4 Hz, 1H), 1.17 (d, *J* = 6.2 Hz, 3H), 0.90 – 0.88 (m, 3H). $^{13}$C NMR (176 MHz, $CDCl_3$) δ 172.6, 170.6, 169.4, 168.2, 168.1, 168.0, 159.6, 157.3, 153.2, 148.8, 147.8, 147.3, 142.3, 136.5, 135.3, 134.6, 133.3, 130.8, 129.8, 120.8, 120.2, 120.1, 119.7, 117.3, 113.9, 112.8, 111.6, 111.2, 104.9, 75.7, 70.5, 70.5, 70.4, 70.3, 70.2, 69.7, 67.4, 67.3, 61.8, 60.9, 60.8, 60.8, 60.7, 56.3, 56.0, 55.9, 55.9, 55.8, 53.7, 52.0, 50.8, 50.2, 49.8, 49.2, 45.5, 43.5, 41.4, 38.8, 38.3, 33.5, 31.3, 29.7, 26.8, 25.3, 23.2, 21.0, 12.6. HRMS (ESI, m/z): calcd for $C_{70}H_{95}N_9O_{16}SNa^+$ $[M+Na]^+$: 1372.6510, found 1372.6516.

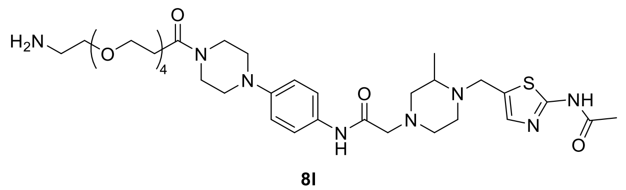

**8l**

**2-(4-((2-acetamidothiazol-5-yl)methyl)-3-methylpiperazin-1-yl)-N-(4-(4-(1-amino-3,6,9,12-tetraoxapentadecan-15-oyl)piperazin-1-yl)phenyl)acetamide (8l)**

Intermediate **8l** was synthesized from **7b** (50 mg, 88 μmol) and BocNH-PEG4-$CH_2CH_2COOH$ (32 mg, 88 μmol) using **General Method B**. Flash column chromatography (silica gel, MeOH in DCM: 0-10%) to give **Boc-8l** (white solid, 65.9 mg, 92%). $^1H$ NMR (700 MHz, $CDCl_3$) δ 11.06 (s, 1H), 8.87 (s, 1H), 7.46 (d, *J* = 8.4 Hz, 2H), 7.24 (d, *J* = 2.1 Hz, 1H), 6.88 (d, *J* = 8.5 Hz, 2H), 4.11 (d, *J* = 8.1 Hz, 1H), 3.81 (t, *J* = 6.7 Hz, 2H), 3.77 (t, *J* = 5.2 Hz, 2H), 3.68 (t, *J* = 6.6 Hz, 1H), 3.62 (d, *J* = 18.1 Hz, 14H), 3.52 (t, *J* = 5.2 Hz, 2H), 3.31 (q, *J* = 5.4 Hz, 2H), 3.09 (dq, *J* = 18.0, 4.9 Hz, 6H), 2.91 – 2.75 (m, 4H), 2.72 (s, 1H), 2.68 (t, *J* = 6.7 Hz, 2H), 2.62 – 2.51 (m, 2H), 2.31 (s, 3H), 1.45 (s, 3H), 1.43 (d, *J* = 4.1 Hz, 9H).

**8l** was synthesized from **Boc-8l** (20 mg, 24 μmol) using **General Method A**.

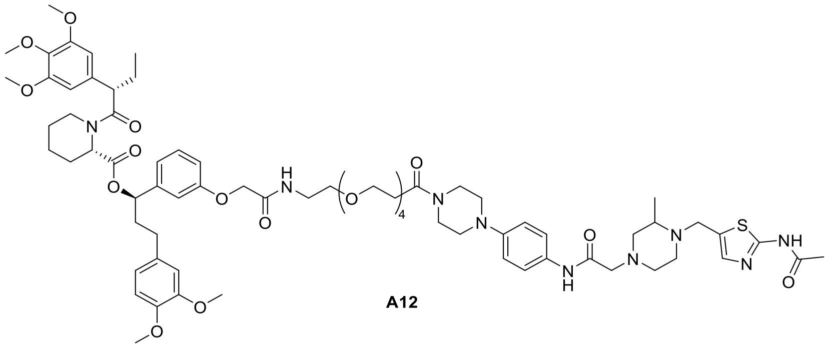


A12

**(1R)-1-(3-((18-(4-(4-(2-(4-((2-acetamidothiazol-5-yl)methyl)-3-methylpiperazin-1-yl)acetamido)phenyl)piperazin-1-yl)-2,18-dioxo-6,9,12,15-tetraoxa-3-azaoctadecyl)oxy)phenyl)-3-(3,4-dimethoxyphenyl)propyl (2S)-1-((S)-2-(3,4,5-trimethoxyphenyl)butanoyl)piperidine-2-carboxylate (A12)**

**A12** was synthesized from **8l** (20 mg, 24 μmol) and **AP1867** (17 mg, 24 μmol) using **General Method B.** pTLC (DCM: MeOH=15:1) to give the product **A12** (white solid, 25.7 mg, 75%). HPLC purity (254 nM)= 95.80%, RT= 15.309 min. $^1H$ NMR (700 MHz, $CDCl_3$) δ 11.97 (s, 1H), 8.92 (s, 1H), 7.46 – 7.42 (m, 2H), 7.20 (s, 1H), 7.17 (d, *J* = 1.4 Hz, 1H), 7.16 – 7.14 (m, 1H), 6.88 – 6.85 (m, 2H), 6.79 – 6.75 (m, 3H), 6.65 – 6.62 (m, 3H), 6.40 (d, *J* = 4.2 Hz, 2H), 4.49 – 4.47 (m, 2H), 4.02 (d, *J* = 14.5 Hz, 1H), 3.85 – 3.82 (m, 8H), 3.79 (d, *J* = 6.6 Hz, 2H), 3.77 (d, *J* = 2.4 Hz, 3H), 3.75 (d, *J* = 5.3 Hz, 1H), 3.70 (s, 1H), 3.67 (s, 5H), 3.62 – 3.56 (m, 17H), 3.54 (dd, *J* = 4.9, 3.0 Hz, 1H), 3.09 (dd, *J* = 10.6, 5.4 Hz, 3H), 3.06 (q, *J* = 4.4, 4.0 Hz, 3H), 2.82 – 2.75 (m, 2H), 2.71 (t, *J* = 10.2 Hz, 2H), 2.66 (t, *J* = 6.7 Hz, 2H), 2.61 – 2.57 (m, 1H), 2.54 (td, *J* = 9.6, 4.8 Hz, 1H), 2.51 – 2.44 (m, 2H), 2.45 – 2.35 (m, 2H), 2.30 (d, *J* = 2.0 Hz, 3H), 2.26 – 2.21 (m, 1H), 2.08 – 2.04 (m, 2H), 1.94 – 1.88 (m, 1H), 1.73 – 1.66 (m, 3H), 1.59 (d, *J* = 14.1 Hz, 1H), 1.42 (dddd, *J* = 17.0, 13.0, 8.2, 3.9 Hz, 1H), 1.16 (d, *J* = 6.2 Hz, 3H), 0.90 – 0.87 (m, 3H). $^{13}C$ NMR (176 MHz, $CDCl_3$) δ 172.6, 170.6, 169.5, 168.3, 168.1, 168.0, 159.8, 157.3, 153.4, 153.2, 148.8, 147.8, 147.3, 142.3, 136.5, 135.3, 134.5, 133.3, 130.8, 129.8, 120.8, 120.2, 119.7, 117.3, 113.9, 112.8, 111.6, 111.2, 104.9, 75.7, 70.5, 70.4, 70.4, 70.4, 70.2, 69.8, 67.3, 67.3, 61.8, 60.9, 60.8, 60.7, 56.3, 55.9, 55.9, 55.9, 55.9, 55.8, 53.7, 52.0, 50.8, 50.2, 49.8, 49.2, 45.5, 43.5, 41.4, 38.8, 38.3, 33.5, 31.3, 29.7, 26.8, 25.3, 23.2, 20.9, 12.6. HRMS (ESI, m/z): calcd for $C_{72}H_{99}N_9O_{17}SNa^+$ $[M+Na]^+$: 1416.6772, found 1416.6769.

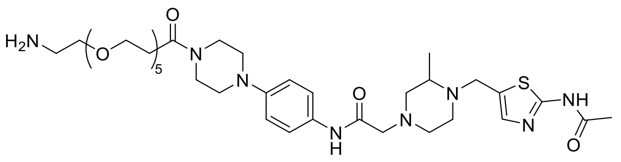

**8m**

**2-(4-((2-acetamidothiazol-5-yl)methyl)-3-methylpiperazin-1-yl)-N-(4-(4-(1-amino-3,6,9,12,15-pentaoxaoctadecan-18-oyl)piperazin-1-yl)phenyl)acetamide (8m)**

Intermediate **8m** was synthesized from **7b** (20 mg, 35 μmol) and BocNH-PEG5-$CH_2CH_2COOH$ (14 mg, 35 μmol) using **General Method B**. Flash column chromatography (silica gel, MeOH in DCM: 0-12%) to give **Boc-8m** (white solid, 14.2 mg, 47%). $^1H$ NMR (700 MHz, $CDCl_3$) δ 10.48 (s, 1H), 8.90 (s, 1H), 7.45 (d, *J* = 8.4 Hz, 2H), 7.24 (s, 1H), 6.89 (d, *J* = 8.5 Hz, 2H), 4.07 (s, 1H), 3.79 (dt, *J* = 26.3, 5.8 Hz, 4H), 3.74 – 3.71 (m, 1H), 3.64 (q, *J* = 14.7, 12.3 Hz, 18H), 3.53 (t, *J* = 5.0 Hz, 2H), 3.31 (d, *J* = 5.4 Hz, 2H), 3.16 – 3.10 (m, 4H), 3.08 (t, *J* = 5.1 Hz, 2H), 2.87 (s, 1H), 2.78 (s, 1H), 2.68 (t, *J* = 6.6 Hz, 2H), 2.57 (d, *J* = 37.2 Hz, 2H), 2.49 (t, *J* = 5.5 Hz, 1H), 2.45 (s, 1H), 2.29 (s, 3H), 2.26 (d, *J* = 9.6 Hz, 1H), 1.43 (s, 9H), 1.25 (s, 3H).

**8m** was synthesized from **Boc-8m** (14.2 mg, 16 μmol) using **General Method A**.

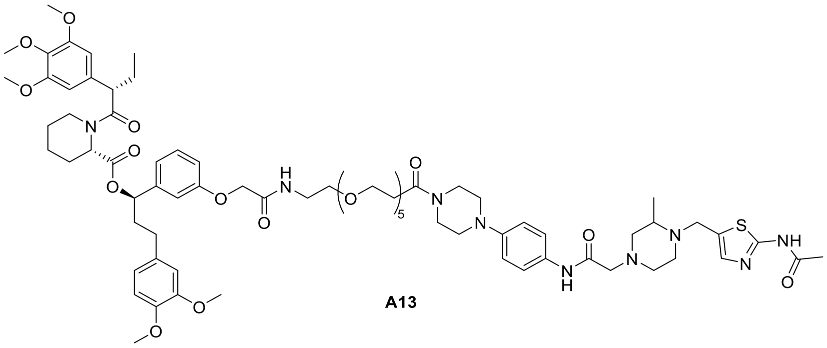

**A13**

**(1R)-1-(3-((21-(4-(4-(2-(4-((2-acetamidothiazol-5-yl)methyl)-3-methylpiperazin-1-yl)acetamido)phenyl)piperazin-1-yl)-2,21-dioxo-6,9,12,15,18-pentaoxa-3-azahenicosyl)oxy)phenyl)-3-(3,4-dimethoxyphenyl)propyl (2S)-1-((S)-2-(3,4,5-trimethoxyphenyl)butanoyl)piperidine-2-carboxylate (A13)**

**A13** was synthesized from **8m** (13 mg, 15 μmol) and **AP1867** (10 mg, 15 μmol) using **General Method B.** pTLC (DCM: MeOH=15:1) to give the product **A13** (white solid, 8.3 mg, 38%). HPLC purity (254 nM)= 96.80%, RT= 15.948 min. $^1H$ NMR (700 MHz, $CDCl_3$) δ 10.97 (s, 1H), 8.92 (s, 1H), 7.45 (d, *J* = 7.2 Hz, 2H), 7.21 (s, 1H), 7.18 (s, 1H), 7.15 (d, *J* = 5.7 Hz, 1H), 6.89 – 6.88 (m, 2H), 6.77 (d, *J* = 13.5 Hz, 3H), 6.65 (dt, *J* = 10.2, 2.2 Hz, 3H), 6.41 (d, *J* = 4.4 Hz, 2H), 5.62 (dd, *J* = 8.2, 5.3 Hz, 1H), 5.47 – 5.45 (m, 1H), 4.51 – 4.48 (m, 2H), 4.02 (d, *J* = 14.7 Hz, 1H), 3.84 (dd, *J* = 11.9, 3.4 Hz, 9H), 3.80 (d, *J* = 6.2 Hz, 2H), 3.78 (s, 3H), 3.76 (d, *J* = 5.0 Hz, 2H), 3.71 (s, 1H), 3.68 (s, 5H), 3.64 (s, 2H), 3.62 (d, *J* = 6.7 Hz, 16H), 3.56 – 3.54 (m, 2H), 3.54 (d, *J* = 5.2 Hz, 1H), 3.44 (q, *J* = 5.2 Hz, 2H), 3.11 (d, *J* = 4.8 Hz, 4H), 3.08 (s, 2H), 2.81 (d, *J* = 10.5 Hz, 2H), 2.73 (d, *J* = 11.3 Hz, 2H), 2.67 (q, *J* = 3.3 Hz, 2H), 2.60 (s, 1H), 2.56 – 2.53 (m, 1H), 2.51 – 2.47 (m, 2H), 2.43 (dd, *J* = 14.4, 6.9 Hz, 2H), 2.29 (d, *J* = 4.5 Hz, 3H), 2.25 (s, 1H), 2.06 (td, *J* = 14.9, 12.4, 5.6 Hz, 2H), 1.92 (td, *J* = 8.9, 8.4, 4.2 Hz, 1H), 1.60 (q, *J* = 6.1 Hz, 3H), 1.56 (s, 1H), 1.44 – 1.41 (m, 1H), 1.17 (d, *J* = 6.1 Hz, 3H), 0.90 (d, *J* = 7.4 Hz, 3H). $^{13}C$ NMR (176 MHz, $CDCl_3$) δ 172.7, 170.6, 169.5, 168.0, 167.8, 159.2, 157.3, 153.2, 148.8, 147.8, 147.3, 142.3, 136.5, 135.3, 133.3,

130.9, 129.8, 120.8, 120.2, 117.4, 114.0, 112.8, 111.6, 111.2, 104.9, 75.7, 70.5, 70.5, 70.2, 70.1, 69.8, 67.3, 61.7, 60.8, 56.3, 56.0, 55.9, 55.9, 55.8, 54.1, 53.6, 52.1, 50.8, 50.2, 49.8, 49.2, 49.1, 45.6, 43.5, 41.4, 39.1, 38.8, 38.3, 36.7, 33.5, 31.9, 31.3, 29.7, 29.7, 29.4, 26.8, 25.8, 25.3, 23.2, 21.0, 12.6. HRMS (ESI, m/z): calcd for $C_{74}H_{103}N_9O_{18}SNa^+$ $[M+Na]^+$: 1460.7034, found 1460.7012.

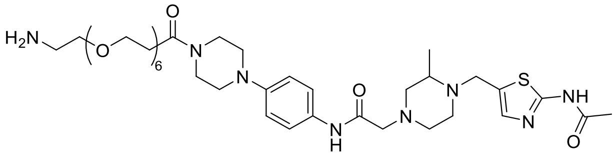

**8n**

**2-(4-((2-acetamidothiazol-5-yl)methyl)-3-methylpiperazin-1-yl)-N-(4-(4-(1-amino-3,6,9,12,15,18-hexaoxahenicosan-21-oyl)piperazin-1-yl)phenyl)acetamide (8n)**

Intermediate **8n** was synthesized from **7b** (20 mg, 35 μmol) and BocNH-PEG6-$CH_2CH_2COOH$ (16 mg, 35 μmol) using **General Method B**. Flash column chromatography (silica gel, MeOH in DCM: 0-12%) to give **Boc-8n** (white solid, 14.6 mg, 46%). $^1H$ NMR (700 MHz, CDCl3) δ 10.57 (s, 1H), 8.90 (s, 1H), 7.45 (d, *J* = 8.3 Hz, 2H), 7.23 (s, 1H), 6.89 (d, *J* = 8.5 Hz, 2H), 4.10 – 4.04 (m, 1H), 3.81 (t, *J* = 6.7 Hz, 2H), 3.77 (t, *J* = 5.1 Hz, 2H), 3.69 (d, *J* = 6.4 Hz, 1H), 3.63 (q, *J* = 10.4, 8.6 Hz, 22H), 3.53 (t, *J* = 5.0 Hz, 2H), 3.31 (q, *J* = 5.3 Hz, 2H), 3.12 (t, *J* = 5.0 Hz, 4H), 3.08 (t, *J* = 5.1 Hz, 2H), 2.86 (s, 1H), 2.78 (s, 2H), 2.73 (s, 1H), 2.68 (t, *J* = 6.6 Hz, 2H), 2.61 (s, 1H), 2.49 (t, *J* = 5.6 Hz, 1H), 2.30 (s, 3H), 2.24 (s, 1H), 1.43 (s, 9H), 1.25 (s, 3H).

**8n** was synthesized from **Boc-8n** (14.6 mg, 16 μmol) using **General Method A**.

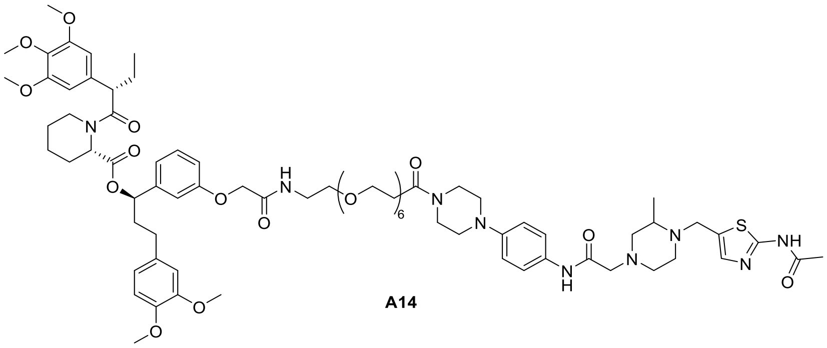

**A14**

**(1R)-1-(3-((24-(4-(4-(2-(4-((2-acetamidothiazol-5-yl)methyl)-3-methylpiperazin-1-yl)acetamido)phenyl)piperazin-1-yl)-2,24-dioxo-6,9,12,15,18,21-hexaoxa-3-azatetracosyl)oxy)phenyl)-3-(3,4-dimethoxyphenyl)propyl (2S)-1-((S)-2-(3,4,5-trimethoxyphenyl)butanoyl)piperidine-2-carboxylate (A14)**

**A14** was synthesized from **8n** (13 mg, 14 μmol) and **AP1867** (10 mg, 15 μmol) using **General Method B.** pTLC (DCM: MeOH=15:1) to give the product **A14** (white solid, 10.4 mg, 49%). HPLC purity (254 nM)= 95.12%, RT= 15.317 min. $^1H$ NMR (700 MHz, $CDCl_3$) δ 11.29 (s, 1H), 8.92 (s, 1H), 7.44 (d, *J* = 8.9 Hz, 2H), 7.20 (s, 1H), 7.18 (d, *J* = 7.9 Hz, 1H), 7.16 (d, *J* = 7.2 Hz, 1H), 6.88 (d, *J* = 8.9 Hz, 2H), 6.80 – 6.75 (m, 3H), 6.66 – 6.62 (m, 3H), 6.41 (d, *J* = 4.4 Hz, 2H), 5.61 (dd, *J* = 8.2, 5.4 Hz, 1H), 5.47 – 5.45 (m, 1H), 4.51 – 4.48 (m, 2H), 4.02 (d, *J* = 14.6 Hz, 1H), 3.84 (dd, *J* = 12.0, 3.5 Hz, 9H), 3.80 (d, *J* = 6.5 Hz, 2H), 3.78 (s, 3H), 3.76 (t, *J* = 5.0 Hz, 2H), 3.71 (d, *J* = 14.3 Hz, 1H), 3.68 (s, 5H), 3.64 – 3.63 (m, 2H), 3.62 (dd, *J* = 7.9, 2.5 Hz, 22H), 3.58

(d, *J* = 6.0 Hz, 2H), 3.55 – 3.54 (m, 1H), 3.12 – 3.10 (m, 2H), 3.08 (d, *J* = 6.4 Hz, 4H), 2.80 (t, *J* = 10.5 Hz, 2H), 2.72 (s, 2H), 2.67 (t, *J* = 6.7 Hz, 2H), 2.60 (s, 1H), 2.56 – 2.52 (m, 1H), 2.48 (dd, *J* = 13.9, 6.5 Hz, 2H), 2.43 (dd, *J* = 13.4, 8.2 Hz, 2H), 2.29 (s, 3H), 2.25 (s, 1H), 2.06 (dt, *J* = 14.3, 7.5 Hz, 2H), 1.92 (dd, *J* = 9.6, 6.2 Hz, 1H), 1.70 (dd, *J* = 13.9, 7.0 Hz, 3H), 1.60 (d, *J* = 15.3 Hz, 1H), 1.43 (dt, *J* = 12.8, 3.5 Hz, 1H), 1.17 (d, *J* = 6.2 Hz, 3H), 0.89 (s, 3H). $^{13}$C NMR (176 MHz, $CDCl_3$) δ 175.8, 172.7, 170.6, 169.5, 168.2, 168.0, 167.9, 159.4, 157.3, 153.2, 148.8, 147.8, 147.3, 142.3, 136.5, 135.3, 133.3, 130.8, 129.8, 120.8, 120.2, 119.7, 117.3, 114.0, 112.7, 111.6, 111.2, 104.9, 75.7, 70.5, 70.5, 70.4, 70.3, 69.7, 67.4, 67.3, 61.7, 60.9, 60.8, 60.7, 56.3, 56.0, 55.9, 55.8, 53.6, 52.1, 50.8, 50.2, 49.8, 49.2, 45.6, 43.5, 41.4, 38.8, 38.3, 36.0, 33.6, 31.3, 29.3, 26.8, 25.3, 23.2, 21.0, 12.6. HRMS (ESI, m/z): calcd for $C_{76}H_{107}N_9O_{19}SNa^+$ $[M+Na]^+$: 1504.7296, found 1504.7281.

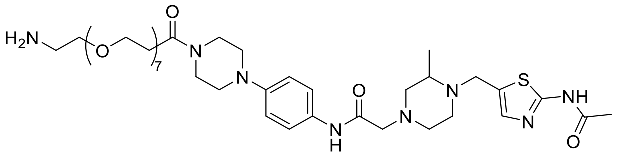

**8o**

**2-(4-((2-acetamidothiazol-5-yl)methyl)-3-methylpiperazin-1-yl)-N-(4-(4-(1-amino-3,6,9,12,15,18,21-heptaoxatetracosan-24-oyl)piperazin-1-yl)phenyl)acetamide (8o)**

Intermediate **8o** was synthesized from **7b** (20 mg, 35 μmol) and BocNH-PEG7-$CH_2CH_2COOH$ (17 mg, 35 μmol) using **General Method B**. Flash column chromatography (silica gel, MeOH in DCM: 0-12%) to give **Boc-8o** (white solid, 13.6 mg, 41%). $^{1}$H NMR (700 MHz, $CDCl_3$) δ 10.42 (s, 1H), 8.91 (s, 1H), 7.45 (d, *J* = 8.4 Hz, 2H), 7.23 (d, *J* = 7.1 Hz, 0H), 6.89 (d, *J* = 8.8 Hz, 2H), 4.05 (s, 1H), 3.80 (d, *J* = 6.7 Hz, 2H), 3.77 (t, *J* = 5.1 Hz, 2H), 3.72 – 3.69 (m, 0H), 3.67 – 3.59 (m, 28H), 3.53 (t, *J* = 5.1 Hz, 2H), 3.31 (q, *J* = 5.2 Hz, 2H), 3.14 – 3.09 (m, 4H), 3.08 (d, *J* = 5.2 Hz, 2H), 2.84 (s, 1H), 2.76 (s, 2H), 2.68 (t, *J* = 6.7 Hz, 2H), 2.62 (d, *J* = 17.4 Hz, 1H), 2.55 (s, 1H), 2.49 (t, *J* = 5.6 Hz, 0H), 2.29 (s, 3H), 2.26 (s, 1H), 1.43 (s, 9H), 1.20 (s, 3H).

**8o** was synthesized from **Boc-8o** (13.6 mg, 14 μmol) using **General Method A**.

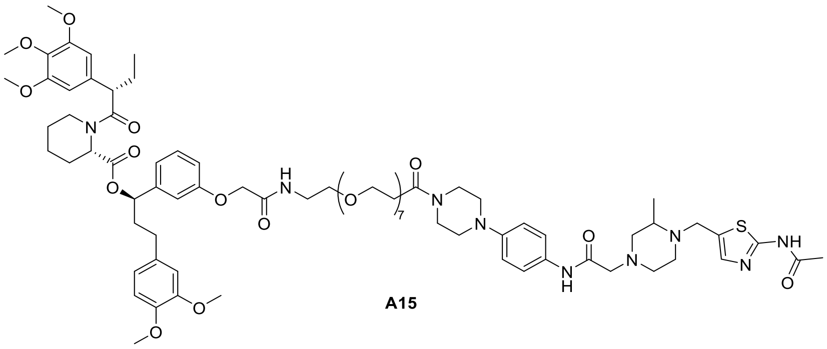

**A15**

**(1R)-1-(3-((27-(4-(4-(2-(4-((2-acetamidothiazol-5-yl)methyl)-3-methylpiperazin-1-yl)acetamido)phenyl)piperazin-1-yl)-2,27-dioxo-6,9,12,15,18,21,24-heptaoxa-3-azaheptacosyl)oxy)phenyl)-3-(3,4-dimethoxyphenyl)propyl (2S)-1-((S)-2-(3,4,5-trimethoxyphenyl)butanoyl)piperidine-2-carboxylate (A15)**

**A15** was synthesized from **8o** (12 mg, 13 μmol) and **AP1867** (9 mg, 13 μmol) using

**General Method B.** pTLC (DCM: MeOH=15:1) to give the product **A15** (white solid, 10.8 mg, 56%). HPLC purity (254 nM)= 95.17%, RT= 15.361 min. $^{1}$H NMR (700 MHz, $CDCl_3$) δ 8.93 (s, 1H), 7.44 (d, *J* = 8.5 Hz, 2H), 7.28 (s, 1H), 7.19 (s, 1H), 7.16 (dd, *J* = 15.2, 6.3 Hz, 1H), 6.88 (d, *J* = 8.8 Hz, 2H), 6.83 – 6.73 (m, 3H), 6.67 – 6.62 (m, 3H), 6.55 (t, *J* = 5.8 Hz, 1H), 6.40 (d, *J* = 4.6 Hz, 1H), 6.33 (s, 1H), 5.56 (dd, *J* = 8.3, 5.4 Hz, 1H), 5.46 (d, *J* = 4.6 Hz, 1H), 4.57 (d, *J* = 11.7 Hz, 2H), 4.02 (d, *J* = 14.6 Hz, 1H), 3.83 (dt, *J* = 12.1, 6.2 Hz, 9H), 3.80 (q, *J* = 5.2, 3.9 Hz, 2H), 3.77 (s, 2H), 3.76 (s, 3H), 3.71 (d, *J* = 3.4 Hz, 1H), 3.68 (d, *J* = 7.1 Hz, 2H), 3.64 (s, 5H), 3.62 (d, *J* = 8.2 Hz, 24H), 3.59 – 3.54 (m, 2H), 3.53 (d, *J* = 5.1 Hz, 1H), 3.44 (q, *J* = 5.3 Hz, 2H), 3.10 (d, *J* = 5.0 Hz, 2H), 3.08 (d, *J* = 6.5 Hz, 4H), 2.81 (d, *J* = 11.2 Hz, 2H), 2.71 (d, *J* = 10.4 Hz, 2H), 2.67 (t, *J* = 6.6 Hz, 2H), 2.63 – 2.59 (m, 1H), 2.57 – 2.54 (m, 1H), 2.50 – 2.46 (m, 2H), 2.41 (t, *J* = 10.3 Hz, 2H), 2.28 (d, *J* = 3.0 Hz, 3H), 2.25 (s, 1H), 2.08 – 2.02 (m, 2H), 1.93 (dt, *J* = 15.2, 7.9 Hz, 1H), 1.73 – 1.64 (m, 3H), 1.54 (dt, *J* = 14.7, 7.8 Hz, 1H), 1.42 (ddt, *J* = 13.1, 9.1, 3.6 Hz, 1H), 1.17 (d, *J* = 6.2 Hz, 3H), 0.89 (d, *J* = 5.8 Hz, 3H). $^{13}$C NMR (176 MHz, $CDCl_3$) δ 173.5, 172.7, 170.4, 169.5, 168.0, 159.6, 158.2, 153.1, 148.8, 147.8, 147.3, 142.1, 136.5, 136.4, 135.1, 134.2, 133.4, 130.8, 130.2, 129.8, 129.6, 125.5, 120.8, 120.2, 117.3, 114.0, 112.7, 111.6, 111.2, 104.8, 76.1, 75.7, 70.5, 70.5, 70.4, 70.1, 67.4, 61.7, 60.8, 56.3, 56.0, 55.9, 55.8, 53.6, 52.1, 50.9, 50.3, 49.8, 49.1, 45.6, 43.5, 41.4, 39.1, 38.3, 36.7, 33.5, 31.9, 31.3, 29.7, 26.8, 25.8, 25.3, 23.2, 22.7, 20.9, 12.6. HRMS (ESI, m/z): calcd for $C_{78}H_{111}N_9O_{20}SNa^+$ $[M+Na]^+$: 1548.7558, found 1548.7550.

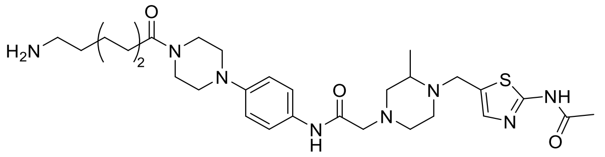

**8p**

**2-(4-((2-acetamidothiazol-5-yl)methyl)-3-methylpiperazin-1-yl)-N-(4-(4-(5-aminopentanoyl)piperazin-1-yl)phenyl)acetamide (8p)**

Intermediate **8p** was synthesized from **7b** (20 mg, 35 μmol) and BocNH-C2-$CH_2CH_2COOH$ (8 mg, 35 μmol) using **General Method B**. Flash column chromatography (silica gel, MeOH in DCM: 0-11%) to give **Boc-8p** (white solid, 12.8 mg, 54%). $^{1}$H NMR (700 MHz, CDCl3) δ 9.77 (s, 1H), 8.92 (s, 1H), 7.45 (d, *J* = 9.0 Hz, 2H), 7.21 (s, 1H), 6.89 (d, *J* = 9.0 Hz, 2H), 4.02 (d, *J* = 14.7 Hz, 1H), 3.77 (t, *J* = 5.1 Hz, 2H), 3.72 (d, *J* = 14.3 Hz, 1H), 3.62 (s, 2H), 3.17 – 3.11 (m, 4H), 3.09 (q, *J* = 5.3, 4.5 Hz, 4H), 2.81 (d, *J* = 11.4 Hz, 1H), 2.72 (d, *J* = 11.8 Hz, 2H), 2.60 (s, 1H), 2.50 (s, 1H), 2.42 (s, 1H), 2.39 (t, *J* = 7.6 Hz, 2H), 2.28 (s, 3H), 2.26 (d, *J* = 7.7 Hz, 1H), 1.69 (t, *J* = 7.4 Hz, 2H), 1.53 (d, *J* = 7.4 Hz, 2H), 1.43 (s, 9H), 1.18 (d, *J* = 6.1 Hz, 3H).

**8p** was synthesized from **Boc-8p** (12.8 mg, 19 μmol) using **General Method A**.

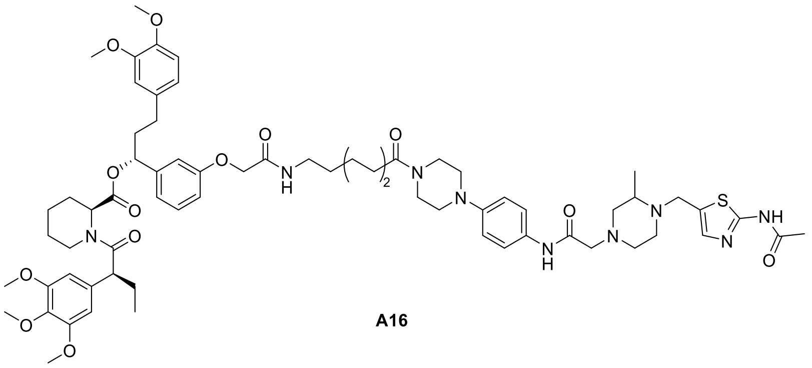

A16

**(1R)-1-(3-(2-((5-(4-(4-(2-(4-((2-acetamidothiazol-5-yl)methyl)-3-methylpiperazin-1-yl)acetamido)phenyl)piperazin-1-yl)-5-oxopentyl)amino)-2-oxoethoxy)phenyl)-3-(3,4-dimethoxyphenyl)propyl (2S)-1-((S)-2-(3,4,5-trimethoxyphenyl)butanoyl)piperidine-2-carboxylate (A16)**

**A16** was synthesized from **8p** (12 mg, 18 μmol) and **AP1867** (12 mg, 18 μmol) using **General Method B.** pTLC (DCM: MeOH=15:1) to give the product **A16** (white solid, 9.5 mg, 42%). HPLC purity (254 nM)= 95.25%, RT= 15.532 min. $^{1}$H NMR (700 MHz, $CDCl_3$) δ 11.40 (s, 1H), 8.92 (s, 1H), 7.44 (d, *J* = 8.7 Hz, 2H), 7.22 (s, 1H), 7.18 (t, *J* = 8.0 Hz, 1H), 6.97 (t, *J* = 5.9 Hz, 1H), 6.89 – 6.86 (m, 2H), 6.80 – 6.75 (m, 3H), 6.67 – 6.63 (m, 3H), 6.40 (d, *J* = 9.3 Hz, 2H), 5.61 (dd, *J* = 8.4, 5.2 Hz, 1H), 5.47 – 5.44 (m, 1H), 4.48 (d, *J* = 10.1 Hz, 2H), 4.04 (d, *J* = 14.5 Hz, 1H), 3.86 – 3.82 (m, 9H), 3.78 (d, *J* = 2.2 Hz, 3H), 3.76 – 3.73 (m, 2H), 3.71 (q, *J* = 5.8, 5.4 Hz, 1H), 3.67 (s, 5H), 3.60 (dd, *J* = 6.6, 3.7 Hz, 2H), 3.58 (d, *J* = 7.1 Hz, 1H), 3.38 (dt, *J* = 13.9, 7.0 Hz, 2H), 3.09 (d, *J* = 6.3 Hz, 4H), 3.06 (d, *J* = 4.7 Hz, 2H), 2.82 (s, 2H), 2.74 (s, 2H), 2.62 (d, *J* = 7.6 Hz, 1H), 2.56 (d, *J* = 8.8 Hz, 1H), 2.55 – 2.47 (m, 2H), 2.47 – 2.41 (m, 2H), 2.39 (t, *J* = 7.2 Hz, 2H), 2.29 (s, 3H), 2.26 (s, 1H), 2.07 (tt, *J* = 14.6, 5.8 Hz, 2H), 1.93 (ddd, *J* = 15.0, 9.4, 6.2 Hz, 1H), 1.71 (dt, *J* = 6.9, 3.4 Hz, 3H), 1.66 – 1.55 (m, 5H), 1.47 – 1.43 (m, 1H), 1.18 (d, *J* = 6.1 Hz, 3H), 0.91 – 0.88 (m, 3H). $^{13}$C NMR (176 MHz, $CDCl_3$) δ 175.8, 172.7, 171.1, 170.6, 168.4, 168.2, 167.9, 159.5, 157.3, 153.2, 148.8, 147.8, 147.3, 142.3, 136.5, 136.0, 135.3, 133.3, 130.8, 129.8, 120.9, 120.2, 119.8, 117.4, 113.5, 113.1, 111.6, 111.2, 104.9, 75.6, 67.3, 61.7, 60.9, 60.8, 56.3, 55.9, 55.9, 55.8, 53.5, 52.1, 50.8, 50.2, 49.9, 49.2, 45.4, 43.5, 41.5, 38.7, 38.2, 36.0, 34.9, 32.5, 31.9, 31.3, 29.3, 29.1, 27.2, 26.8, 25.3, 23.2, 22.2, 20.9, 12.6. HRMS (ESI, m/z): calcd for $C_{66}H_{87}N_9O_{13}SNa^+$ $[M+Na]^+$: 1268.6036, found 1268.6024.

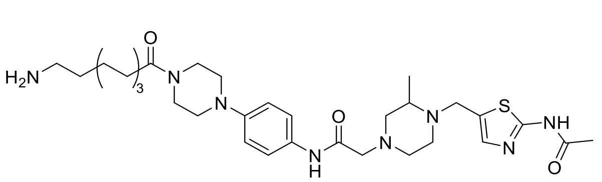

8q

**2-(4-((2-acetamidothiazol-5-yl)methyl)-3-methylpiperazin-1-yl)-N-(4-(4-(6-aminohexanoyl)piperazin-1-yl)phenyl)acetamide (8q)**

Intermediate **8q** was synthesized from **7b** (20 mg, 35 μmol) and BocNH-C3-$CH_2CH_2COOH$ (8 mg, 35 μmol) using **General Method B**. Flash column chromatography (silica gel, MeOH in DCM: 0-9%) to give **Boc-8q** (white solid, 15.6 mg, 65%). $^{1}$H NMR (700 MHz, $CDCl_3$) δ 8.92 (s, 1H), 7.45 (d, *J* = 9.1 Hz, 2H), 7.22 (s, 1H), 6.89 (d, *J* = 9.1 Hz, 2H), 4.05 (s, 1H), 3.76 (t, *J* = 5.1 Hz, 2H), 3.70 (dt, *J* = 6.8, 3.6 Hz, 1H), 3.61 (t, *J* = 5.0 Hz, 2H), 3.13 (dq, *J* = 16.3, 6.5, 4.6 Hz, 6H), 3.08 (d, *J* =

4.7 Hz, 2H), 2.84 (s, 1H), 2.76 (s, 2H), 2.65 (s, 1H), 2.53 (s, 1H), 2.47 (s, 1H), 2.36 (t, *J* = 7.6 Hz, 2H), 2.29 (s, 3H), 2.25 (s, 1H), 1.67 (d, *J* = 7.6 Hz, 2H), 1.43 (s, 9H), 1.38 – 1.36 (m, 2H), 1.20 (s, 3H).

**8q** was synthesized from **Boc-8q** (15.6 mg, 23 µmol) using **General Method A**.

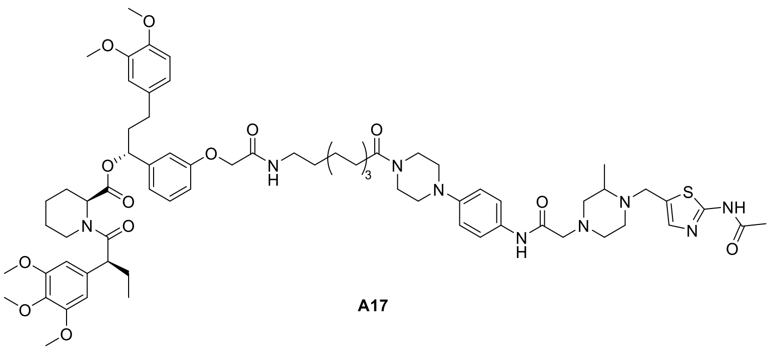

**A17**

**(1R)-1-(3-(2-((6-(4-(4-(2-(4-((2-acetamidothiazol-5-yl)methyl)-3-methylpiperazin-1-yl)acetamido)phenyl)piperazin-1-yl)-6-oxohexyl)amino)-2-oxoethoxy)phenyl)-3-(3,4-dimethoxyphenyl)propyl (2S)-1-((S)-2-(3,4,5-trimethoxyphenyl)butanoyl)piperidine-2-carboxylate (A17)**

**A17** was synthesized from **8q** (14 mg, 21 µmol) and **AP1867** (14 mg, 21 µmol) using **General Method B.** pTLC (DCM: MeOH=15:1) to give the product **A17** (white solid, 11.0 mg, 43%). HPLC purity (254 nM)= 96.29%, RT= 15.694 min. $^{1}$H NMR (700 MHz, $CDCl_3$) δ 11.58 (s, 1H), 8.93 (s, 1H), 7.44 (d, *J* = 8.4 Hz, 2H), 7.22 (s, 1H), 7.18 (t, *J* = 8.4 Hz, 1H), 6.88 (d, *J* = 8.5 Hz, 2H), 6.82 (t, *J* = 6.2 Hz, 1H), 6.80 – 6.75 (m, 3H), 6.69 – 6.62 (m, 3H), 6.40 (d, *J* = 10.3 Hz, 2H), 5.61 (dd, *J* = 8.3, 5.4 Hz, 1H), 5.46 (d, *J* = 5.5 Hz, 1H), 4.47 (s, 2H), 4.04 (d, *J* = 14.5 Hz, 1H), 3.84 (q, *J* = 6.2, 5.4 Hz, 9H), 3.77 (s, 3H), 3.75 (t, *J* = 5.0 Hz, 2H), 3.71 (d, *J* = 13.9 Hz, 1H), 3.67 (s, 5H), 3.60 (d, *J* = 4.9 Hz, 2H), 3.58 (d, *J* = 7.1 Hz, 1H), 3.38 – 3.33 (m, 2H), 3.10 (t, *J* = 5.3 Hz, 4H), 3.07 (d, *J* = 5.3 Hz, 2H), 2.84 – 2.78 (m, 2H), 2.73 (d, *J* = 10.5 Hz, 2H), 2.61 (d, *J* = 15.1 Hz, 1H), 2.56 (q, *J* = 4.4 Hz, 1H), 2.54 – 2.46 (m, 2H), 2.45 (dt, *J* = 14.4, 5.2 Hz, 2H), 2.36 (t, *J* = 7.5 Hz, 2H), 2.30 (s, 3H), 2.27 (dd, *J* = 16.7, 7.6 Hz, 1H), 2.07 (dt, *J* = 13.5, 7.4 Hz, 2H), 1.95 – 1.91 (m, 1H), 1.72 – 1.66 (m, 5H), 1.60 (p, *J* = 7.5 Hz, 3H), 1.43 (d, *J* = 12.4 Hz, 1H), 1.41 – 1.37 (m, 2H), 1.18 (d, *J* = 6.2 Hz, 3H), 0.89 (s, 3H). $^{13}$C NMR (176 MHz, $CDCl_3$) δ 171.4, 170.6, 168.1, 167.9, 159.6, 157.3, 153.2, 148.8, 147.9, 147.3, 142.4, 136.5, 135.3, 133.3, 130.8, 129.8, 129.5, 120.8, 120.2, 119.8, 117.4, 113.5, 113.0, 111.6, 111.2, 104.9, 75.6, 67.3, 61.7, 60.8, 56.3, 55.9, 55.9, 55.8, 53.6, 52.0, 50.8, 50.3, 49.9, 49.2, 45.5, 43.5, 41.4, 38.9, 38.3, 33.1, 31.3, 29.5, 26.7, 24.8, 23.2, 20.9, 12.6. HRMS (ESI, m/z): calcd for $C_{67}H_{89}N_9O_{13}SNa^+$ $[M+Na]^+$: 1282.6193, found 1282.6190.

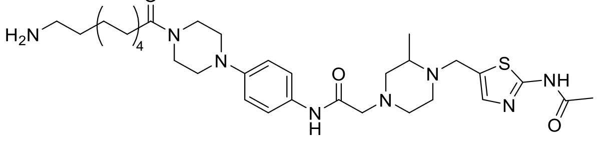

**8r**

**2-(4-((2-acetamidothiazol-5-yl)methyl)-3-methylpiperazin-1-yl)-N-(4-(4-(7-aminoheptanoyl)piperazin-1-yl)phenyl)acetamide (8r)**

Intermediate **8r** was synthesized from **7b** (40 mg, 70 μmol) and BocNH-C4-$CH_2CH_2COOH$ (17 mg, 70 μmol) using **General Method B**. Flash column chromatography (silica gel, MeOH in DCM: 0-10%) to give **Boc-8r** (white solid, 40.0 mg, 81%). $^1$H NMR (700 MHz, $CDCl_3$) δ 11.46 (s, 1H), 8.93 (s, 1H), 7.45 (d, *J* = 8.4 Hz, 2H), 7.21 (s, 1H), 6.89 (d, *J* = 8.4 Hz, 2H), 4.53 (s, 1H), 4.03 (d, *J* = 14.6 Hz, 1H), 3.76 (t, *J* = 5.1 Hz, 2H), 3.70 (d, *J* = 14.6 Hz, 1H), 3.61 (t, *J* = 5.0 Hz, 2H), 3.49 – 3.46 (m, 1H), 3.11 (t, *J* = 5.5 Hz, 3H), 3.08 (d, *J* = 6.9 Hz, 3H), 2.81 (d, *J* = 11.7 Hz, 1H), 2.73 (t, *J* = 10.6 Hz, 2H), 2.60 (s, 1H), 2.49 (t, *J* = 10.6 Hz, 1H), 2.41 (t, *J* = 10.9 Hz, 1H), 2.35 (t, *J* = 7.7 Hz, 2H), 2.31 (s, 3H), 2.28 – 2.21 (m, 1H), 1.64 (t, *J* = 7.4 Hz, 2H), 1.51 – 1.48 (m, 2H), 1.43 (s, 9H), 1.35 (dq, *J* = 12.4, 6.5 Hz, 4H), 1.17 (d, *J* = 6.2 Hz, 3H).

**8r** was synthesized from **Boc-8r** (20.0 mg, 29 μmol) using **General Method A**.

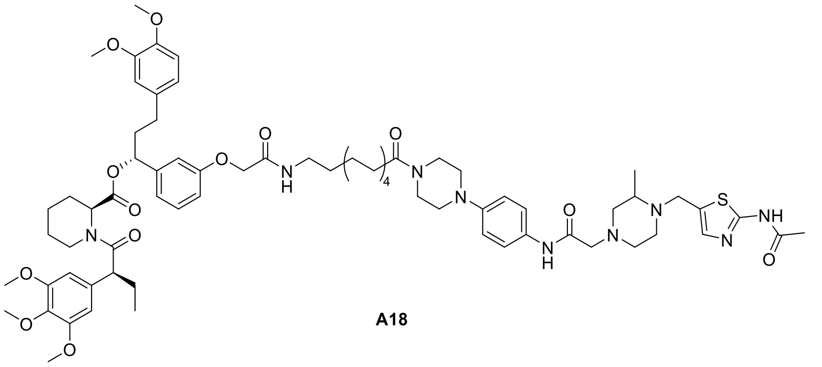

A18

**(1R)-1-(3-(2-((7-(4-(4-(2-(4-((2-acetamidothiazol-5-yl)methyl)-3-methylpiperazin-1-yl)acetamido)phenyl)piperazin-1-yl)-7-oxoheptyl)amino)-2-oxoethoxy)phenyl)-3-(3,4-dimethoxyphenyl)propyl (2S)-1-((S)-2-(3,4,5-trimethoxyphenyl)butanoyl)piperidine-2-carboxylate (A18)**

**A18** was synthesized from **8r** (20 mg, 29 μmol) and **AP1867** (20 mg, 29 μmol) using **General Method B.** pTLC (DCM: MeOH=15:1) to give the product **A18** (white solid, 16.7 mg, 46%). HPLC purity (254 nM)= 95.25%, RT= 15.939 min. $^1$H NMR (700 MHz, $CDCl_3$) δ 11.81 (s, 1H), 8.92 (s, 1H), 7.45 (d, *J* = 8.9 Hz, 2H), 7.20 (s, 1H), 7.20 – 7.16 (m, 1H), 6.88 (d, *J* = 8.9 Hz, 2H), 6.78 – 6.74 (m, 3H), 6.67 – 6.62 (m, 3H), 6.40 (d, *J* = 8.7 Hz, 2H), 5.61 (dd, *J* = 8.3, 5.4 Hz, 1H), 5.47 – 5.45 (m, 1H), 4.47 (s, 2H), 4.03 (d, *J* = 14.5 Hz, 1H), 3.86 – 3.81 (m, 9H), 3.77 (s, 3H), 3.75 (dd, *J* = 6.2, 4.1 Hz, 2H), 3.69 (d, *J* = 14.6 Hz, 1H), 3.67 (s, 5H), 3.61 – 3.59 (m, 2H), 3.59 – 3.56 (m, 1H), 3.33 (dt, *J* = 16.3, 7.1 Hz, 2H), 3.10 (t, *J* = 5.3 Hz, 2H), 3.09 – 3.05 (m, 4H), 2.82 – 2.76 (m, 2H), 2.74 – 2.70 (m, 2H), 2.64 – 2.58 (m, 1H), 2.55 (ddd, *J* = 14.6, 9.7, 5.5 Hz, 1H), 2.52 – 2.45 (m, 2H), 2.45 – 2.37 (m, 2H), 2.34 (dd, *J* = 8.9, 6.3 Hz, 2H), 2.30 (s, 3H), 2.26 – 2.20 (m, 1H), 2.10 – 2.04 (m, 2H), 1.94 – 1.90 (m, 1H), 1.70 (tt, *J* = 11.3, 6.0 Hz, 3H), 1.66 – 1.62 (m, 2H), 1.62 – 1.58 (m, 1H), 1.58 – 1.53 (m, 2H), 1.43 (ddt, *J* = 18.0, 9.0, 4.5 Hz, 1H), 1.37 (dt, *J* = 7.5, 4.7 Hz, 4H), 1.17 (d, *J* = 6.2 Hz, 3H), 0.90 – 0.88 (m, 3H). $^{13}$C NMR (176 MHz, $CDCl_3$) δ 172.7, 171.5, 170.6, 168.1, 168.0, 168.0, 159.7, 157.3, 153.2, 148.8, 147.8, 147.3, 142.3, 136.5, 135.3, 134.5, 133.3, 130.9, 129.8, 120.8, 120.2, 119.8, 117.4, 113.5, 113.0, 111.6, 111.2, 104.9, 75.6, 67.3, 61.8, 60.9, 60.8, 60.7, 56.3, 55.9, 55.9, 55.8, 53.7, 52.0, 50.8, 50.3, 49.9, 49.2, 45.5, 43.5, 41.4, 39.0, 38.2, 33.1, 31.3, 29.5, 29.1, 26.7, 25.1, 23.2, 20.9, 12.6. HRMS (ESI, m/z): calcd for $C_{68}H_{91}N_9O_{13}SNa^+$ $[M+Na]^+$: 1296.6349, found 1296.6348.

**Scheme S5.** Synthesis of **B1-B2**

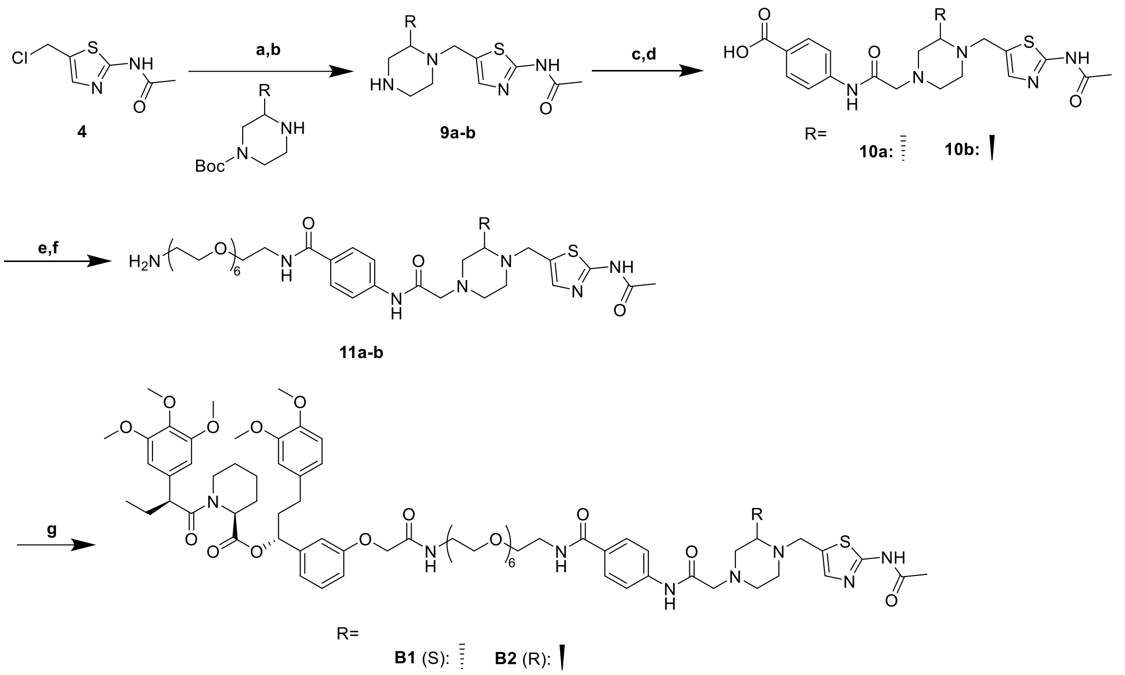


Reaction and conditions: a) TEA, MeCN, rt, 12 h, 63%; b) TFA, DCM, rt, 1.5 h; c) $K_2CO_3$, MeCN, rt, 17 h, 21%; d) TFA, DCM, rt, 2 h; e) HATU, DIPEA, DCM, rt, 14 h, 86%; f) TFA, DCM, rt, 1 h; g) AP1867, HATU, DIPEA, DCM, 15 h, 62%.

**B1** and **B2** were synthesized following the same procedure for preparing **A1**.

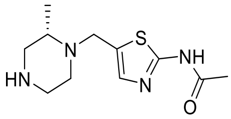

**9a**

##### (S)-N-(5-((2-methylpiperazin-1-yl)methyl)thiazol-2-yl)acetamide (9a) (LE1)

OGA ligand **9a** was synthesized following the same procedure for preparing intermediate **5** from intermediate **4** (500 mg, 2.62 mmol) and tert-butyl (S)-3-methylpiperazine-1-carboxylate (630 mg, 3.15 mmol). Flash column chromatography (silica gel, MeOH in DCM: 0-5%) to give **Boc-9a** (white solid, 582.3 mg, 63%. $^1$H NMR (700 MHz, $CDCl_3$) δ 11.72 (s, 2H), 7.19 (s, 1H), 3.99 (d, $J$ = 14.5 Hz, 1H), 3.78 (s, 1H), 3.70 (d, $J$ = 12.9 Hz, 1H), 3.10 (s, 1H), 2.88 (s, 1H), 2.77 (s, 1H), 2.70 (dt, $J$ = 11.6, 3.7 Hz, 1H), 2.46 (s, 1H), 2.31 (s, 3H), 2.23 (t, $J$ = 10.3 Hz, 1H), 1.44 (s, 9H), 1.12 (d, $J$ = 6.2 Hz, 3H).

**9a** was synthesized from **Boc-9a** (500 mg, 1.41 mmol) using **General Method A**.

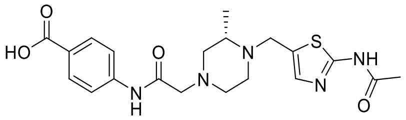

**10a**

##### (S)-4-(2-(4-((2-acetamidothiazol-5-yl)methyl)-3-methylpiperazin-1-yl)acetamido)benzoic acid (10a) (LB1)

OGA ligand **10a** was synthesized following the same procedure for preparing OGA ligand **7a** from **6a** (150 mg, 0.56 mmol) and **9a** (196 mg, 0.57 mmol). Column chromatography (silica gel, Methanol: DCM =1:100 to 1:30) and further purified via reverse-ISCO to give the **t-Bu-10a** (white solid, 57.6 mg, 21%). LC-MS ($ESI^+$) m/z= 488.1.

The **t-Bu-10a** (50 mg. 0.11 mmol) was dissolved in TFA/DCM (1:4, 8 mL). The mixture was stirred at rt for 2h until the reaction was complete (monitored by TLC). The mixture was concentrated under reduced pressure to give the intermediate compound **10a** without further purification (white solid).

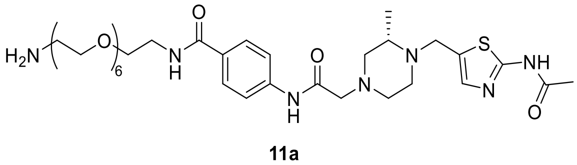

**11a**

**(S)-4-(2-(4-((2-acetamidothiazol-5-yl)methyl)-3-methylpiperazin-1-yl)acetamido)-N-(20-amino-3,6,9,12,15,18-hexaoxaicosyl)benzamide (11a)**

Intermediate **11a** was synthesized from **10a** (28 mg, 65 μmol) and BocNH-PEG6-$NH_2$ (28 mg, 65 μmol) using **General Method B**. Flash column chromatography (silica gel, MeOH in DCM: 0-8%) to give the **Boc-11a** (white solid, 46.7 mg, 86%). $^1H$ NMR (700 MHz, $CDCl_3$) δ 9.22 (s, 1H), 7.83 (d, $J$ = 8.4 Hz, 2H), 7.62 (d, $J$ = 8.4 Hz, 2H), 7.18 (s, 1H), 7.17 (s, 1H), 4.02 (d, $J$ = 14.6 Hz, 1H), 3.72 (d, $J$ = 13.5 Hz, 1H), 3.69 – 3.56 (m, 24H), 3.51 (t, $J$ = 5.1 Hz, 2H), 3.30 (q, $J$ = 5.4 Hz, 2H), 3.15 – 3.08 (m, 2H), 2.82 (d, $J$ = 11.3 Hz, 1H), 2.74 (t, $J$ = 10.8 Hz, 2H), 2.63 (s, 1H), 2.52 (s, 1H), 2.44 (s, 1H), 2.28 (s, 3H), 2.27 (s, 1H), 1.43 (d, $J$ = 5.1 Hz, 9H), 1.19 (d, $J$ = 6.2 Hz, 3H).

**11a** was synthesized from **Boc-11a** (20 mg, 24 μmol) using **General Method A**.

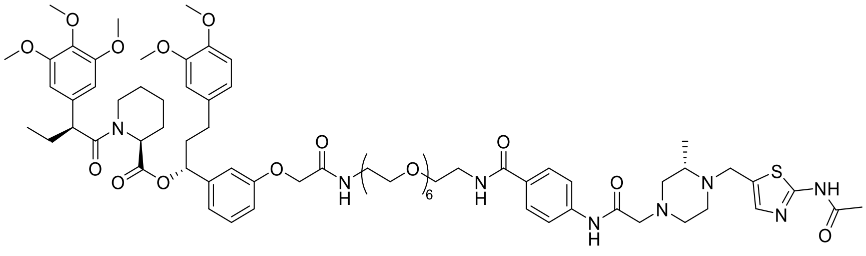

**B1**

**(R)-1-(3-((1-(4-(2-((S)-4-((2-acetamidothiazol-5-yl)methyl)-3-methylpiperazin-1-yl)acetamido)phenyl)-1,24-dioxo-5,8,11,14,17,20-hexaoxa-2,23-diazapentacosan-25-yl)oxy)phenyl)-3-(3,4-dimethoxyphenyl)propyl (S)-1-((S)-2-(3,4,5-trimethoxyphenyl)butanoyl)piperidine-2-carboxylate (B1)**

**B1** was synthesized from **11a** (20 mg, 24 μmol) and **AP1867** (17 mg, 24 μmol) using **General Method B.** pTLC (DCM: MeOH=15:1) to give the product **B1** (white solid, 21.0 mg, 62%). HPLC purity (254 nM)= 97.60%, RT= 15.410 min. $^1H$ NMR (700 MHz, $CDCl_3$) δ 9.22 (s, 1H), 7.83 (d, $J$ = 8.2 Hz, 2H), 7.60 (d, $J$ = 8.2 Hz, 2H), 7.27 (d, $J$ = 5.2 Hz, 1H), 7.26 (s, 1H), 7.16 (t, $J$ = 7.8 Hz, 1H), 6.77 (q, $J$ = 8.0 Hz, 3H), 6.65 – 6.60 (m, 3H), 6.40 (d, $J$ = 5.1 Hz, 2H), 4.48 (d, $J$ = 9.8 Hz, 2H), 4.02 (d, $J$ = 14.5 Hz, 1H), 3.83 (q, $J$ = 8.4, 6.6 Hz, 9H), 3.77 (s, 3H), 3.70 (d, $J$ = 14.6 Hz, 1H), 3.67 (s,

5H), 3.65 – 3.59 (m, 22H), 3.56 (d, $J$ = 7.5 Hz, 6H), 3.52 (d, $J$ = 6.3 Hz, 1H), 3.09 (d, $J$ = 7.4 Hz, 2H), 2.83 – 2.75 (m, 2H), 2.71 (d, $J$ = 10.9 Hz, 2H), 2.61 (s, 1H), 2.53 (dd, $J$ = 11.1, 6.0 Hz, 1H), 2.49 (t, $J$ = 11.1 Hz, 2H), 2.43 (d, $J$ = 11.3 Hz, 2H), 2.29 (s, 3H), 2.25 (d, $J$ = 10.0 Hz, 1H), 2.05 (td, $J$ = 14.2, 7.2 Hz, 2H), 1.91 (p, $J$ = 7.8, 7.1 Hz, 1H), 1.69 (tt, $J$ = 10.9, 5.7 Hz, 3H), 1.59 (d, $J$ = 13.7 Hz, 1H), 1.46 – 1.38 (m, 1H), 1.17 (d, $J$ = 6.2 Hz, 3H), 0.90 – 0.87 (m, 3H). $^{13}$C NMR (176 MHz, $CDCl_3$) δ 176.5, 172.7, 170.6, 168.6, 168.3, 168.1, 166.8, 159.7, 157.3, 153.1, 148.8, 147.3, 142.2, 140.2, 136.5, 135.3, 134.5, 133.3, 129.9, 129.8, 128.3, 120.2, 119.7, 118.7, 114.0, 112.7, 111.6, 111.2, 104.9, 75.7, 70.5, 70.4, 70.2, 70.1, 70.0, 69.7, 67.3, 61.8, 60.9, 60.8, 60.7, 56.3, 55.9, 55.9, 55.8, 54.2, 53.7, 52.1, 50.8, 49.2, 43.5, 39.8, 38.8, 38.3, 31.3, 29.7, 26.8, 25.3, 23.2, 20.9, 12.6. HRMS (ESI, m/z): calcd for $C_{72}H_{100}N_8O_{19}SNa^+$ $[M+Na]^+$: 1435.6718, found 1435.6709.

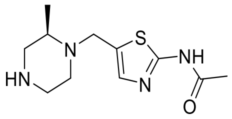


**9b**

**(R)-N-(5-((2-methylpiperazin-1-yl)methyl)thiazol-2-yl)acetamide (9b) (LE2)**

OGA ligand **9b** was synthesized following the same procedure for preparing intermediate **5** from intermediate **4** (500 mg, 2.62 mmol) and tert-butyl (R)-3-methylpiperazine-1-carboxylate (630 mg, 3.15 mmol). Flash column chromatography (silica gel, MeOH in DCM: 0-5%) to give **Boc-9b** (white solid, 582.1 mg, 63%. $^{1}$H NMR (700 MHz, $CDCl_3$) δ 11.81 (s, 1H), 7.19 (s, 1H), 3.99 (d, $J$ = 14.5 Hz, 1H), 3.78 (s, 1H), 3.70 (d, $J$ = 13.2 Hz, 1H), 3.10 (s, 1H), 2.88 (s, 1H), 2.77 (s, 1H), 2.70 (dt, $J$ = 11.6, 3.7 Hz, 1H), 2.46 (s, 1H), 2.31 (s, 3H), 2.23 (s, 1H), 1.44 (s, 9H), 1.12 (d, $J$ = 6.2 Hz, 3H).

**9b** was synthesized from **Boc-9b** (500 mg, 1.41 mmol) using **General Method A**.

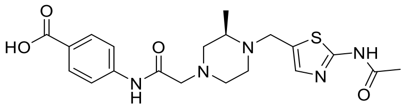


**10b**

**(R)-4-(2-(4-((2-acetamidothiazol-5-yl)methyl)-3-methylpiperazin-1-yl)acetamido)benzoic acid (10b) (LB2)**

OGA ligand **10b** was synthesized following the same procedure for preparing OGA ligand **7a** from **6a** (150 mg, 0.56 mmol) and **9b** (196 mg, 0.57 mmol). Column chromatography (silica gel, Methanol: DCM =1:100 to 1:30) and further purified via reverse-ISCO to give the **t-Bu-10b** (white solid, 53.7 mg, 20%). LC-MS (ESI$^+$) m/z= 488.1.

The **t-Bu-10b** (50 mg. 0.11 mmol) was dissolved in TFA/DCM (1:4, 8 mL). The mixture was stirred at rt for 2h until the reaction was complete (monitored by TLC). The mixture was concentrated under reduced pressure to give the intermediate compound **10b** without further purification (white solid).

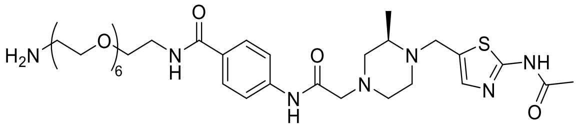

**11b**

**(R)-4-(2-(4-((2-acetamidothiazol-5-yl)methyl)-3-methylpiperazin-1-yl)acetamido)-N-(20-amino-3,6,9,12,15,18-hexaoxaicosyl)benzamide (11b)**

Intermediate **11b** was synthesized from **10b** (26 mg, 60 μmol) and BocNH-PEG6-$NH_2$ (26 mg, 60 μmol) using **General Method B**. Flash column chromatography (silica gel, MeOH in DCM: 0-8%) to give the **Boc-11b** (white solid, 30.0 mg, 59%). $^1H$ NMR (700 MHz, $CDCl_3$) δ 9.24 (s, 1H), 7.84 (d, *J* = 8.6 Hz, 2H), 7.62 (d, *J* = 8.3 Hz, 2H), 7.22 (s, 1H), 4.04 (d, *J* = 14.6 Hz, 1H), 3.76 (d, *J* = 14.7 Hz, 1H), 3.69 (t, *J* = 5.0 Hz, 2H), 3.67 – 3.61 (m, 18H), 3.58 (s, 4H), 3.51 (t, *J* = 5.1 Hz, 2H), 3.30 (q, *J* = 5.5 Hz, 2H), 3.11 (d, *J* = 8.3 Hz, 2H), 2.83 (d, *J* = 11.1 Hz, 1H), 2.74 (t, *J* = 12.0 Hz, 2H), 2.64 (s, 1H), 2.51 (d, *J* = 10.7 Hz, 1H), 2.46 (s, 1H), 2.30 (s, 3H), 2.28 – 2.24 (m, 1H), 1.42 (s, 9H), 1.19 (d, *J* = 6.2 Hz, 3H).

**11b** was synthesized from **Boc-11b** (20 mg, 24 μmol) using **General Method A**.

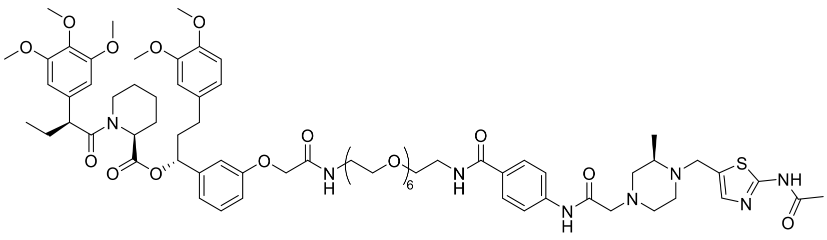

**B2**

**(R)-1-(3-((1-(4-(2-((R)-4-((2-acetamidothiazol-5-yl)methyl)-3-methylpiperazin-1-yl)acetamido)phenyl)-1,24-dioxo-5,8,11,14,17,20-hexaoxa-2,23-diazapentacosan-25-yl)oxy)phenyl)-3-(3,4-dimethoxyphenyl)propyl (S)-1-((S)-2-(3,4,5-trimethoxyphenyl)butanoyl)piperidine-2-carboxylate (B2)**

**B2** was synthesized from **11b** (20 mg, 24 μmol) and **AP1867** (17 mg, 24 μmol) using **General Method B.** pTLC (DCM: MeOH=15:1) to give the product **B2** (white solid, 9.5 mg, 28%). HPLC purity (254 nM)= 97.79%, RT= 15.424 min. $^1H$ NMR (700 MHz, $CDCl_3$) δ 9.22 (s, 1H), 7.83 (d, *J* = 8.2 Hz, 2H), 7.61 (d, *J* = 8.1 Hz, 2H), 7.23 (t, *J* = 6.9 Hz, 1H), 7.20 (d, *J* = 8.0 Hz, 1H), 7.17 (t, *J* = 8.1 Hz, 1H), 6.79 – 6.74 (m, 3H), 6.66 – 6.62 (m, 3H), 6.41 (d, *J* = 7.1 Hz, 2H), 5.61 (dd, *J* = 8.2, 5.5 Hz, 1H), 5.45 (d, *J* = 5.4 Hz, 1H), 4.48 (d, *J* = 11.2 Hz, 2H), 4.03 (s, 1H), 3.84 (q, *J* = 7.7, 6.4 Hz, 9H), 3.77 (s, 3H), 3.73 (s, 1H), 3.68 (s, 5H), 3.66 – 3.60 (m, 20H), 3.58 (d, *J* = 4.0 Hz, 8H), 3.53 (d, *J* = 5.7 Hz, 1H), 3.11 (d, *J* = 8.8 Hz, 2H), 2.86 – 2.77 (m, 2H), 2.73 (d, *J* = 11.3 Hz, 2H), 2.65 (s, 1H), 2.59 (dd, *J* = 16.3, 7.3 Hz, 1H), 2.53 (dt, *J* = 13.6, 8.5 Hz, 2H), 2.45 (td, *J* = 9.1, 8.2, 4.7 Hz, 2H), 2.31 (s, 1H), 2.28 (s, 3H), 2.06 (dd, *J* = 13.8, 7.1 Hz, 2H), 1.93 (s, 1H), 1.71 (dd, *J* = 12.9, 6.3 Hz, 3H), 1.61 (d, *J* = 13.4 Hz, 1H), 1.47 (d, *J* = 6.6 Hz, 1H), 1.19 (s, 3H), 0.91 – 0.89 (m, 3H). $^{13}C$ NMR (176 MHz, $CDCl_3$) δ 176.2, 172.7, 170.6, 168.5, 168.0, 166.0, 157.3, 153.2, 148.8, 147.3, 142.3, 140.3, 136.5, 135.3, 134.2, 133.3, 129.8, 129.5, 128.3, 120.2, 119.7, 118.8, 113.9, 112.8, 111.6, 111.2, 104.9, 75.7, 70.3, 70.0, 69.9, 67.8, 67.3, 61.8, 60.8, 56.3, 55.9, 55.9, 55.8, 54.8, 52.1, 50.7, 49.1, 43.5, 39.7, 38.9, 38.8, 38.3, 31.3, 30.6, 29.7, 29.0, 26.8, 25.3, 24.0, 23.2, 23.0, 20.9, 12.6. HRMS (ESI, m/z): calcd for $C_{72}H_{100}N_8O_{19}SNa^+$ $[M+Na]^+$: 1435.6718, found 1435.6713.

**Scheme S6.** Synthesis of **C1-C4**

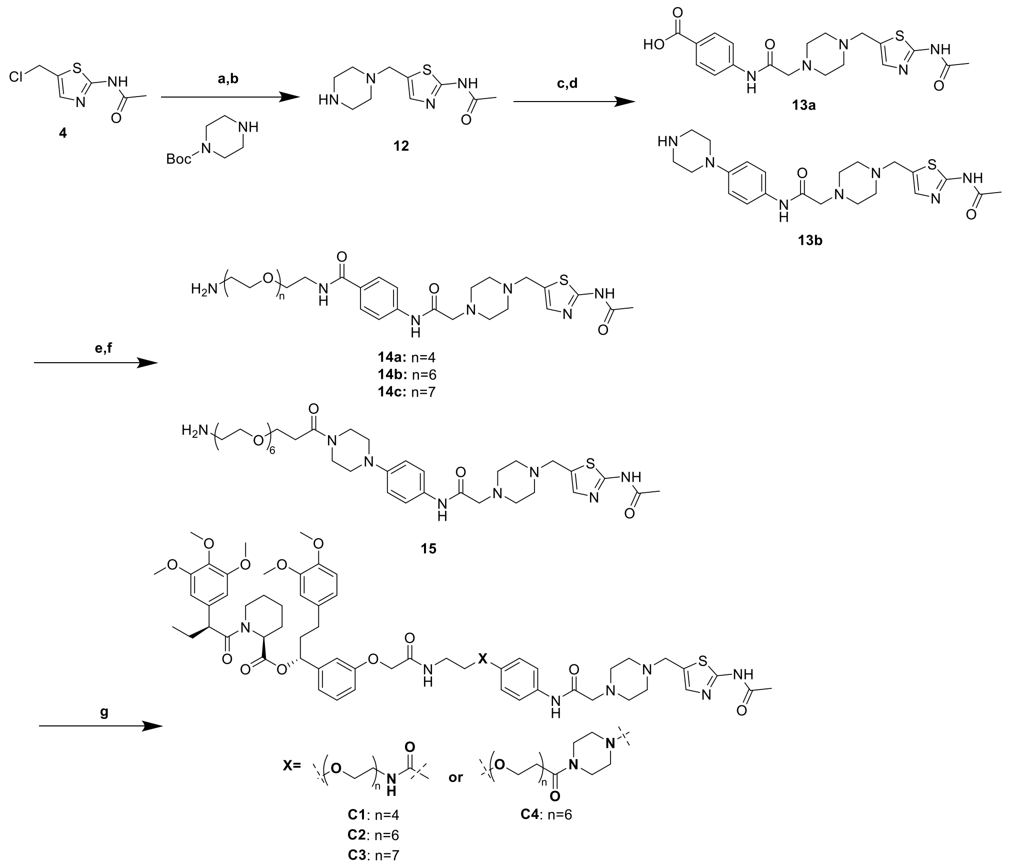


Reaction and conditions: a) TEA, MeCN, rt, 12 h, 40%; b) TFA, DCM, rt, 1.5 h; c) $K_2CO_3$, MeCN, rt, 17 h, 20%; d) TFA, DCM, rt, 2 h; e) HATU, DIPEA, DCM, rt, 14 h, 50%; f) TFA, DCM, rt, 1 h; g) AP1867, HATU, DIPEA, DCM, 15 h, 38%.

**C1-C4** were synthesized following the same procedure for preparing **A1**.

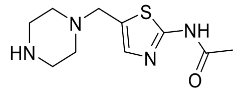

**12**

**N-(5-(piperazin-1-ylmethyl)thiazol-2-yl)acetamide (12)**

Intermediate **12** was synthesized following the same procedure for preparing intermediate **5** from intermediate **4** (1.0 g, 5.52 mmol) and tert-butyl piperazine-1-carboxylate (1.2 g, 6.29 mmol). Flash column chromatography (silica gel, MeOH in DCM: 0-1%) to give **Boc-12** (white solid, 720.7 mg, 40%). LCMS ($ESI^+$) m/z = 341.0.

**12** was synthesized from **Boc-12** (700 mg, 2.06 mmol) using **General Method A**. $^1H$ NMR (700 MHz, $CDCl_3$) δ 7.18 (s, 1H), 3.73 (s, 2H), 3.15 (t, *J* = 4.9 Hz, 4H), 2.73 (s, 4H), 2.19 (d, *J* = 1.9 Hz, 3H).

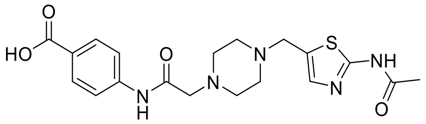

**13a**

**4-(2-(4-((2-acetamidothiazol-5-yl)methyl)piperazin-1-yl)acetamido)benzoic acid (13a) (L9)**

OGA ligand **13a** was synthesized following the same procedure for preparing OGA ligand **7a** from **6a** (300 mg, 1.11 mmol) and **12** (376 mg, 1.11 mmol). Flash column chromatography (silica gel, MeOH in DCM: 0-5%) to give the **t-Bu-13a** (white solid, 107.4 mg, 20%). $^{1}$H NMR (700 MHz, $CDCl_3$) δ 11.06 (s, 1H), 9.28 (s, 1H), 7.95 (d, *J* = 8.7 Hz, 2H), 7.62 – 7.60 (m, 2H), 7.22 (s, 1H), 3.74 – 3.71 (m, 2H), 3.15 (s, 2H), 2.66 (s, 4H), 2.59 (s, 4H), 2.30 (s, 3H), 1.58 (s, 9H).

The **t-Bu-13a** (100 mg. 0.21 mmol) was dissolved in TFA/DCM (1:4, 8 mL). The mixture was stirred at rt for 2h until the reaction was complete (monitored by TLC). The mixture was concentrated under reduced pressure to give the intermediate compound **13a** without further purification (white solid).

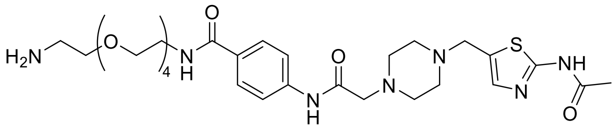

**14a**

**4-(2-(4-((2-acetamidothiazol-5-yl)methyl)piperazin-1-yl)acetamido)-N-(14-amino-3,6,9,12-tetraoxatetradecyl)benzamide (14a)**

Intermediate **14a** was synthesized from **13a** (20 mg, 48 μmol) and BocNH-PEG4-$NH_2$ (16 mg, 48 μmol) using **General Method B**. Flash column chromatography (silica gel, MeOH in DCM: 0-8%) to give the **Boc-14a** (white solid, 17.7 mg, 50%). LCMS ($ESI^+$) m/z= 736.2.

**14a** was synthesized from **Boc-14a** (17.7 mg, 24 μmol) using **General Method A**.

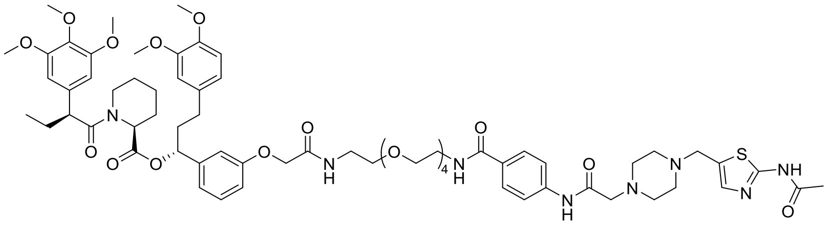

**C1**

**(R)-1-(3-((1-(4-(2-(4-((2-acetamidothiazol-5-yl)methyl)piperazin-1-yl)acetamido)phenyl)-1,18-dioxo-5,8,11,14-tetraoxa-2,17-diazanonadecan-19-yl)oxy)phenyl)-3-(3,4-dimethoxyphenyl)propyl (S)-1-((S)-2-(3,4,5-trimethoxyphenyl)butanoyl)piperidine-2-carboxylate (C1)**

**C1** was synthesized from **14a** (18 mg, 25 μmol) and **AP1867** (17 mg, 25 μmol) using **General Method B.** pTLC (DCM: MeOH=15:1) to give the product **C1** (white solid, 12.3 mg, 38%). HPLC purity (254 nM)= 96.86%, RT= 15.343 min. $^{1}$H NMR (700 MHz, $CDCl_3$) δ 11.60 (s, 1H), 9.25 (s, 1H), 7.83 (d, *J* = 8.6 Hz, 2H), 7.61 (d, *J* = 8.6 Hz,

2H), 7.23 (d, *J* = 6.0 Hz, 1H), 7.20 (s, 1H), 7.16 (s, 1H), 6.77 (q, *J* = 6.9, 5.7 Hz, 3H), 6.66 – 6.61 (m, 3H), 6.40 (s, 2H), 5.61 (t, *J* = 6.9 Hz, 1H), 5.45 (d, *J* = 5.5 Hz, 1H), 4.47 (d, *J* = 8.5 Hz, 2H), 3.85 – 3.82 (m, 9H), 3.77 (d, *J* = 1.7 Hz, 3H), 3.71 (s, 2H), 3.67 (d, *J* = 1.6 Hz, 5H), 3.65 – 3.59 (m, 13H), 3.59 (d, *J* = 1.7 Hz, 5H), 3.56 (t, *J* = 5.4 Hz, 2H), 3.51 (s, 1H), 3.13 (s, 2H), 2.78 (t, *J* = 13.0 Hz, 1H), 2.64 (s, 4H), 2.60 – 2.56 (m, 2H), 2.55 – 2.52 (m, 1H), 2.44 (dt, *J* = 14.6, 7.8 Hz, 1H), 2.29 (s, 3H), 2.23 (d, *J* = 13.4 Hz, 1H), 2.20 (s, 2H), 2.08 – 2.04 (m, 2H), 1.91 (dt, *J* = 18.0, 7.2 Hz, 1H), 1.70 (dq, *J* = 12.8, 6.6 Hz, 3H), 1.59 (s, 1H), 1.45 – 1.41 (m, 1H), 0.90 – 0.88 (m, 3H). $^{13}$C NMR (176 MHz, $CDCl_3$) δ 175.9, 172.7, 170.6, 168.6, 168.0, 166.8, 159.7, 157.3, 153.2, 148.8, 147.3, 142.2, 140.3, 136.5, 135.3, 134.5, 133.3, 129.9, 129.8, 129.5, 129.3, 128.3, 120.2, 119.7, 118.7, 114.0, 112.8, 111.6, 111.2, 104.9, 75.7, 70.4, 70.4, 70.4, 70.1, 70.1, 69.9, 67.8, 67.3, 61.8, 60.8, 56.3, 55.9, 55.9, 55.9, 55.8, 54.1, 53.4, 52.7, 52.1, 50.8, 43.5, 39.8, 38.8, 38.3, 36.0, 31.9, 31.3, 30.6, 29.3, 26.8, 25.3, 24.0, 23.2, 20.9, 12.6. HRMS (ESI, m/z): calcd for $C_{67}H_{90}N_8O_{17}SNa^+$ $[M+Na]^+$: 1333.6037, found 1333.6029.

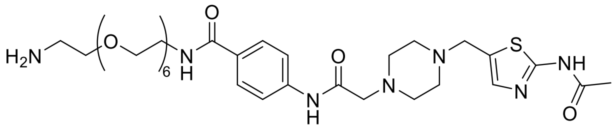


**14b**

**4-(2-(4-((2-acetamidothiazol-5-yl)methyl)piperazin-1-yl)acetamido)-N-(20-amino-3,6,9,12,15,18-hexaoxaicosyl)benzamide (14b)**

Intermediate **14b** was synthesized from **13a** (20 mg, 48 μmol) and BocNH-PEG6-$NH_2$ (20 mg, 48 μmol) using **General Method B**. Flash column chromatography (silica gel, MeOH in DCM: 0-8%) to give the **Boc-14b** (white solid, 19.0 mg, 49%). $^{1}$H NMR (700 MHz, $CDCl_3$) δ 10.61 (s, 1H), 9.26 (s, 1H), 7.84 (d, *J* = 8.3 Hz, 2H), 7.62 (d, *J* = 8.6 Hz, 2H), 7.22 (s, 1H), 7.20 (s, 1H), 3.73 (s, 2H), 3.68 – 3.64 (m, 10H), 3.62 (d, *J* = 7.2 Hz, 10H), 3.59 – 3.58 (m, 2H), 3.51 (t, *J* = 5.1 Hz, 2H), 3.30 (q, *J* = 5.4 Hz, 2H), 3.15 (s, 2H), 3.00 (d, *J* = 36.3 Hz, 2H), 2.66 (s, 4H), 2.59 (s, 4H), 2.29 (s, 3H), 1.43 (s, 9H).

**14b** was synthesized from **Boc-14b** (19.0 mg, 23 μmol) using **General Method A**.

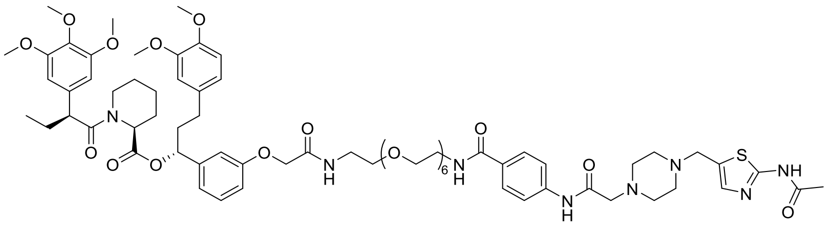

**C2**

**(R)-1-(3-((1-(4-(2-(4-((2-acetamidothiazol-5-yl)methyl)piperazin-1-yl)acetamido)phenyl)-1,24-dioxo-5,8,11,14,17,20-hexaoxa-2,23-diazapentacosan-25-yl)oxy)phenyl)-3-(3,4-dimethoxyphenyl)propyl (S)-1-((S)-2-(3,4,5-trimethoxyphenyl)butanoyl)piperidine-2-carboxylate (C2)**

**C2** was synthesized from **14b** (18 mg, 22 μmol) and **AP1867** (15 mg, 22 μmol) using **General Method B.** pTLC (DCM: MeOH=15:1) to give the product **C2** (white solid, 15.3 mg, 50%). HPLC purity (254 nM)= 98.02%, RT= 15.326 min. $^{1}$H NMR (700 MHz,

$CDCl_3$) δ 11.72 (s, 1H), 9.25 (s, 1H), 7.85 (d, *J* = 8.3 Hz, 2H), 7.61 (d, *J* = 8.3 Hz, 2H), 7.35 (s, 1H), 7.21 (s, 1H), 7.16 (t, *J* = 8.3 Hz, 1H), 6.77 (td, *J* = 10.9, 9.2, 6.2 Hz, 3H), 6.64 (dd, *J* = 10.2, 2.7 Hz, 3H), 6.40 (d, *J* = 6.1 Hz, 2H), 5.61 (dd, *J* = 8.3, 5.4 Hz, 1H), 5.47 – 5.43 (m, 1H), 4.47 (s, 2H), 3.86 – 3.81 (m, 9H), 3.77 (s, 3H), 3.72 (s, 2H), 3.67 (s, 5H), 3.65 – 3.62 (m, 8H), 3.60 (t, *J* = 7.0 Hz, 14H), 3.57 (d, *J* = 4.5 Hz, 6H), 3.53 (s, 1H), 3.13 (s, 2H), 2.78 (td, *J* = 13.4, 3.1 Hz, 1H), 2.65 (s, 4H), 2.62 – 2.53 (m, 4H), 2.52 (dd, *J* = 11.6, 6.3 Hz, 1H), 2.44 (ddd, *J* = 13.9, 9.6, 6.7 Hz, 1H), 2.33 (s, 1H), 2.29 (s, 3H), 2.07 – 2.03 (m, 2H), 1.91 (ddt, *J* = 12.7, 9.8, 6.1 Hz, 1H), 1.70 (qd, *J* = 9.1, 8.2, 3.5 Hz, 3H), 1.60 (d, *J* = 13.4 Hz, 1H), 1.46 – 1.38 (m, 1H), 0.89 (t, *J* = 6.1 Hz, 3H). $^{13}$C NMR (176 MHz, $CDCl_3$) δ 172.7, 170.6, 168.6, 168.4, 168.0, 166.9, 159.8, 157.3, 153.1, 148.8, 147.3, 142.3, 140.3, 136.5, 135.3, 134.5, 133.3, 129.9, 129.8, 128.4, 120.2, 120.1, 119.7, 118.7, 113.9, 112.8, 111.6, 111.2, 104.9, 75.7, 70.3, 70.3, 70.3, 70.3, 70.2, 70.2, 70.1, 70.0, 69.8, 67.3, 61.8, 60.9, 60.8, 56.3, 55.9, 55.9, 55.9, 55.9, 55.8, 54.1, 53.4, 52.7, 52.1, 50.8, 43.5, 39.8, 38.8, 38.3, 31.3, 29.7, 26.8, 25.3, 23.2, 20.9, 12.6. HRMS (ESI, m/z): calcd for $C_{71}H_{98}N_8O_{19}SNa^+$ $[M+Na]^+$: 1421.6561, found 1421.6563.

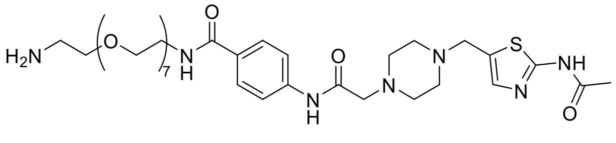


**14c**

**4-(2-(4-((2-acetamidothiazol-5-yl)methyl)piperazin-1-yl)acetamido)-N-(23-amino-3,6,9,12,15,18,21-heptaoxatricosyl)benzamide (14c)**

Intermediate **14c** was synthesized from **13a** (20 mg, 48 μmol) and BocNH-PEG7-$NH_2$ (22 mg, 48 μmol) using **General Method B**. Flash column chromatography (silica gel, MeOH in DCM: 0-9%) to give the **Boc-14c** (white solid, 16.8 mg, 40%). LCMS ($ESI^+$) m/z= 868.2.

**14c** was synthesized from **Boc-14c** (16.8 mg, 19 μmol) using **General Method A**.

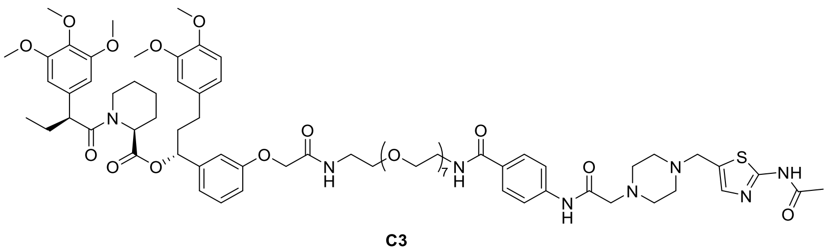

**C3**

**(R)-1-(3-((1-(4-(2-(4-((2-acetamidothiazol-5-yl)methyl)piperazin-1-yl)acetamido)phenyl)-1,27-dioxo-5,8,11,14,17,20,23-heptaoxa-2,26-diazaoctacosan-28-yl)oxy)phenyl)-3-(3,4-dimethoxyphenyl)propyl (S)-1-((S)-2-(3,4,5-trimethoxyphenyl)butanoyl)piperidine-2-carboxylate (C3)**

**C3** was synthesized from **14c** (17 mg, 20 μmol) and **AP1867** (14 mg, 20 μmol) using **General Method B.** pTLC (DCM: MeOH=15:1) to give the product **C3** (white solid, 16.4 mg, 58%). HPLC purity (254 nM)= 97.32%, RT= 15.343 min. $^1$H NMR (700 MHz, $CDCl_3$) δ 9.25 (s, 1H), 7.83 (d, *J* = 8.1 Hz, 2H), 7.61 (d, *J* = 8.2 Hz, 2H), 7.20 (d, *J* = 6.7 Hz, 2H), 7.16 (t, *J* = 7.4 Hz, 2H), 6.79 – 6.75 (m, 3H), 6.65 – 6.61 (m, 3H), 6.40 (d, *J* = 4.3 Hz, 2H), 5.61 (t, *J* = 6.9 Hz, 1H), 5.45 (s, 1H), 4.48 (d, *J* = 9.9 Hz, 2H),

3.84 (q, $J$ = 8.8, 7.0 Hz, 9H), 3.77 (s, 3H), 3.71 (s, 2H), 3.67 (s, 5H), 3.64 (d, $J$ = 8.9 Hz, 11H), 3.59 (d, $J$ = 7.5 Hz, 24H), 3.53 (s, 1H), 3.13 (s, 2H), 2.78 (t, $J$ = 11.9 Hz, 1H), 2.64 (s, 4H), 2.59 – 2.55 (m, 2H), 2.55 – 2.50 (m, 2H), 2.43 (dd, $J$ = 14.7, 7.9 Hz, 1H), 2.30 (s, 1H), 2.29 (s, 3H), 2.23 (dd, $J$ = 34.9, 10.5 Hz, 2H), 2.06 (dd, $J$ = 14.1, 7.0 Hz, 2H), 1.91 (p, $J$ = 8.0, 7.2 Hz, 1H), 1.70 (dq, $J$ = 12.6, 6.4 Hz, 3H), 1.60 (d, $J$ = 14.9 Hz, 1H), 1.42 (dt, $J$ = 17.3, 11.1 Hz, 1H), 0.89 (t, $J$ = 7.1 Hz, 3H). $^{13}$C NMR (176 MHz, $CDCl_3$) δ 172.7, 170.6, 168.6, 168.3, 168.0, 166.8, 159.8, 157.3, 153.2, 148.8, 147.3, 142.3, 140.3, 136.5, 135.3, 134.6, 133.3, 130.0, 129.8, 129.3, 128.3, 120.2, 119.7, 118.7, 114.0, 112.7, 111.6, 111.2, 104.9, 75.7, 70.5, 70.4, 70.2, 70.2, 70.0, 69.7, 67.3, 61.9, 60.9, 60.8, 56.3, 55.9, 55.9, 55.8, 54.2, 53.4, 52.8, 52.1, 50.8, 43.5, 39.8, 38.8, 38.3, 31.3, 29.3, 26.8, 25.3, 23.2, 20.9, 12.6. HRMS (ESI, m/z): calcd for $C_{73}H_{102}N_8O_{20}SNa^+$ [M+Na]$^+$: 1465.6823, found 1465.6804.

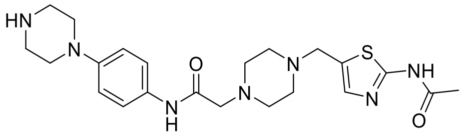


**13b**

**2-(4-((2-acetamidothiazol-5-yl)methyl)piperazin-1-yl)-N-(4-(piperazin-1-yl)phenyl)acetamide (13b) (LC4)**

OGA ligand **13b** was synthesized following the same procedure for preparing OGA ligand **7a** from **6b** (150 mg, 0.42 mmol) and **12** (158 mg, 0.47 mmol). Flash column chromatography (silica gel, MeOH in DCM: 0-7%) to give the **Boc-13b** (white solid, 148.1 mg, 63%). LCMS (ESI$^+$) m/z= 558.1.

**13b** was synthesized from **Boc-13a** (100 mg, 179 μmol) using **General Method A**.

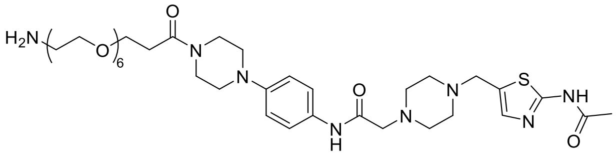


**15**

**2-(4-((2-acetamidothiazol-5-yl)methyl)piperazin-1-yl)-N-(4-(4-(1-amino-3,6,9,12,15,18-hexaoxahenicosan-21-oyl)piperazin-1-yl)phenyl)acetamide (15)**

Intermediate **15** was synthesized from **13b** (20 mg, 36 μmol) and BocNH-PEG6-$CH_2CH_2COOH$ (16 mg, 36 μmol) using **General Method B**. Flash column chromatography (silica gel, MeOH in DCM: 0-10%) to give the **Boc-15** (white solid, 20.0 mg, 62%). LCMS (ESI$^+$) m/z= 736.2. $^1$H NMR (700 MHz, $CDCl_3$) δ 10.84 (s, 1H), 8.98 (s, 1H), 7.46 (d, $J$ = 8.8 Hz, 2H), 7.24 (s, 1H), 6.89 (d, $J$ = 9.0 Hz, 2H), 3.82 – 3.79 (m, 2H), 3.77 (t, $J$ = 5.3 Hz, 2H), 3.73 (dd, $J$ = 12.7, 7.2 Hz, 2H), 3.63 (td, $J$ = 12.5, 5.5 Hz, 22H), 3.53 (t, $J$ = 5.1 Hz, 2H), 3.31 (q, $J$ = 5.4 Hz, 2H), 3.16 (s, 2H), 3.12 (t, $J$ = 5.1 Hz, 2H), 3.09 (t, $J$ = 5.2 Hz, 2H), 2.76 – 2.67 (m, 6H), 2.67 – 2.56 (m, 4H), 2.29 (s, 3H), 1.43 (s, 9H).

**15** was synthesized from **Boc-15** (20.0 mg, 22 μmol) using **General Method A**.

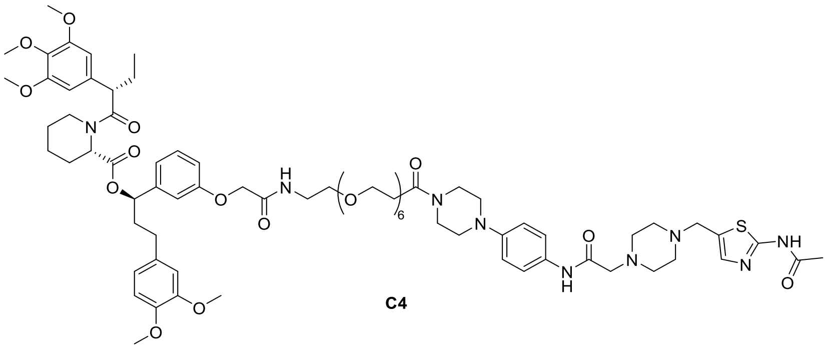


**(R)-1-(3-((24-(4-(4-(2-(4-((2-acetamidothiazol-5-yl)methyl)piperazin-1-yl)acetamido)phenyl)piperazin-1-yl)-2,24-dioxo-6,9,12,15,18,21-hexaoxa-3-azatetracosyl)oxy)phenyl)-3-(3,4-dimethoxyphenyl)propyl (S)-1-((S)-2-(3,4,5-trimethoxyphenyl)butanoyl)piperidine-2-carboxylate (C4)**

**C4** was synthesized from **15** (20 mg, 22 μmol) and **AP1867** (16 mg, 22 μmol) using **General Method B.** pTLC (DCM: MeOH=15:1) to give the product **C4** (white solid, 18.0 mg, 55%). HPLC purity (254 nM)= 95.30%, RT= 15.232 min. $^{1}$H NMR (700 MHz, $CDCl_3$) δ 11.78 (s, 1H), 8.95 (s, 1H), 7.45 (d, *J* = 8.9 Hz, 2H), 7.21 (s, 1H), 7.17 (t, *J* = 7.7 Hz, 2H), 6.88 (d, *J* = 2.4 Hz, 2H), 6.80 – 6.74 (m, 3H), 6.65 – 6.62 (m, 3H), 6.40 (s, 2H), 5.61 (dd, *J* = 8.2, 5.5 Hz, 1H), 5.47 – 5.43 (m, 1H), 4.51 – 4.47 (m, 2H), 3.85 – 3.82 (m, 9H), 3.81 – 3.78 (m, 3H), 3.77 (s, 3H), 3.75 (t, *J* = 5.2 Hz, 2H), 3.70 (s, 2H), 3.67 (s, 5H), 3.61 (dd, *J* = 9.6, 5.4 Hz, 25H), 3.60 – 3.57 (m, 4H), 3.55 – 3.54 (m, 1H), 3.12 (s, 2H), 3.10 (d, *J* = 5.3 Hz, 2H), 3.08 (t, *J* = 5.2 Hz, 2H), 2.78 (td, *J* = 13.4, 2.7 Hz, 1H), 2.68 (d, *J* = 6.6 Hz, 2H), 2.64 (s, 2H), 2.53 (dd, *J* = 9.5, 5.2 Hz, 1H), 2.47 – 2.42 (m, 1H), 2.30 (s, 1H), 2.29 (s, 3H), 2.09 – 2.05 (m, 2H), 1.91 (ddt, *J* = 13.9, 9.9, 6.1 Hz, 1H), 1.74 – 1.65 (m, 3H), 1.58 (t, *J* = 16.4 Hz, 1H), 1.42 (ddt, *J* = 12.9, 9.0, 4.0 Hz, 1H), 0.90 – 0.88 (m, 3H). $^{13}$C NMR (176 MHz, $CDCl_3$) δ 172.7, 170.6, 169.5, 168.3, 168.0, 159.8, 157.3, 153.2, 148.8, 147.8, 147.3, 142.3, 136.5, 135.3, 134.5, 133.3, 130.8, 129.8, 120.8, 120.2, 119.7, 117.4, 114.0, 112.8, 111.6, 111.2, 104.9, 75.7, 70.5, 70.5, 70.4, 70.2, 69.7, 67.4, 67.3, 61.8, 60.9, 60.8, 56.3, 55.9, 55.9, 55.8, 54.1, 53.4, 52.8, 52.0, 50.8, 50.2, 49.8, 45.6, 43.5, 41.5, 38.8, 38.3, 33.5, 31.3, 29.7, 26.8, 25.3, 23.2, 20.9, 12.6. HRMS (ESI, m/z): calcd for $C_{75}H_{105}N_9O_{19}SNa^+$ $[M+Na]^+$: 1490.7140, found 1490.7117.

**Scheme S7.** Synthesis of **D1-D6**

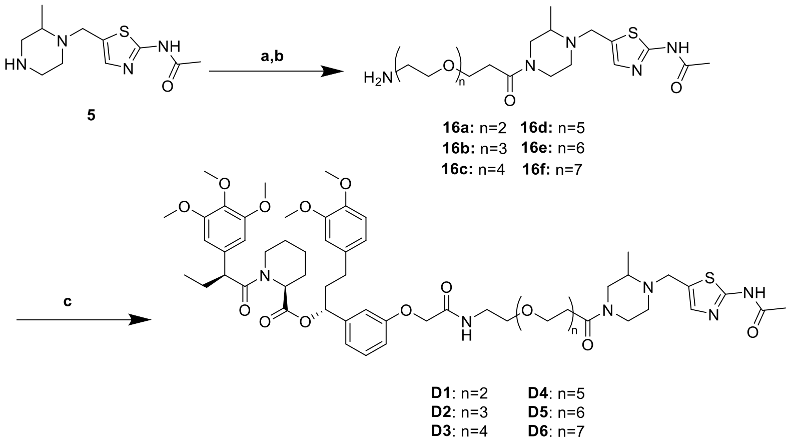


Reaction and conditions: a) HATU, DIPEA, DCM, rt, 13 h, 51%; b) TFA, DCM, rt, 1 h;

c) AP1867, HATU, DIPEA, DCM, 15 h, 67%.

**D1-D6** were synthesized following the same procedure for preparing **A1**.

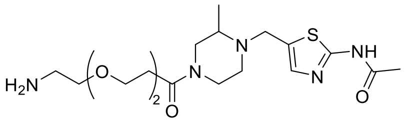

**16a**

**N-(5-((4-(3-(2-(2-aminoethoxy)ethoxy)propanoyl)-2-methylpiperazin-1-yl)methyl)thiazol-2-yl)acetamide (16a)**

Intermediate **16a** was synthesized from **5** (20 mg, 79 µmol) and BocNH-PEG2-$CH_2CH_2COOH$ (22 mg, 79 µmol) using **General Method B**. Flash column chromatography (silica gel, MeOH in DCM: 0-7%) to give the **Boc-16a** (white solid, 20.4 mg, 51%). $^1H$ NMR (700 MHz, $CDCl_3$) δ 11.06 (s, 1H), 7.19 (s, 1H), 4.12 (d, *J* = 12.9 Hz, 1H), 4.05 (d, *J* = 12.7 Hz, 1H), 3.98 (d, *J* = 20.4 Hz, 1H), 3.77 (q, *J* = 6.3 Hz, 2H), 3.66 (dd, *J* = 14.3, 5.8 Hz, 1H), 3.61 – 3.57 (m, 4H), 3.52 (q, *J* = 4.9 Hz, 3H), 3.30 (q, *J* = 5.0 Hz, 2H), 3.14 (t, *J* = 11.0 Hz, 1H), 3.05 (dd, *J* = 13.0, 8.5 Hz, 1H), 2.77 (d, *J* = 11.2 Hz, 1H), 2.73 (dt, *J* = 11.5, 3.7 Hz, 1H), 2.61 (dt, *J* = 13.1, 8.3 Hz, 2H), 2.29 (s, 3H), 1.44 (s, 9H), 1.14 (d, *J* = 6.2 Hz, 3H).

**16a** was synthesized from **Boc-16a** (20.0 mg, 39 µmol) using **General Method A**.

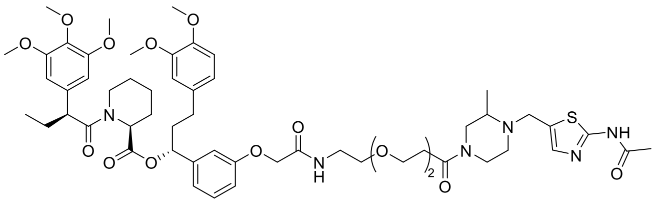

**D1**

**(1R)-1-(3-(2-((2-(2-(3-(4-((2-acetamidothiazol-5-yl)methyl)-3-methylpiperazin-1-yl)-3-oxopropoxy)ethoxy)ethyl)amino)-2-oxoethoxy)phenyl)-3-(3,4-dimethoxyphenyl)propyl (2S)-1-((S)-2-(3,4,5-trimethoxyphenyl)butanoyl)piperidine-2-carboxylate (D1)**

**D1** was synthesized from **16a** (18 mg, 35 µmol) and **AP1867** (24 mg, 35 µmol) using **General Method B.** pTLC (DCM: MeOH=15:1) to give the product **D1** (white solid, 25.7 mg, 67%). HPLC purity (254 nM)= 97.21%, RT= 15.426 min. $^1H$ NMR (700 MHz, $CDCl_3$) δ 11.92 (s, 1H), 7.17 (s, 1H), 7.16 – 7.14 (m, 1H), 7.14 – 7.12 (m, 1H), 6.79 – 6.75 (m, 3H), 6.65 – 6.62 (m, 3H), 6.39 (s, 2H), 5.61 (dd, *J* = 8.2, 5.5 Hz, 1H), 5.46 (d, *J* = 5.6 Hz, 1H), 4.50 – 4.47 (m, 2H), 4.09 (d, *J* = 13.1 Hz, 1H), 4.03 – 3.99 (m, 1H), 3.98 – 3.93 (m, 1H), 3.85 – 3.82 (m, 9H), 3.77 (d, *J* = 1.3 Hz, 3H), 3.75 – 3.74 (m, 1H), 3.67 (d, *J* = 1.2 Hz, 5H), 3.63 (dd, *J* = 14.5, 6.0 Hz, 1H), 3.59 – 3.56 (m, 8H), 3.55 (d, *J* = 5.0 Hz, 1H), 3.53 (d, *J* = 5.6 Hz, 1H), 3.26 (tt, *J* = 9.9, 3.3 Hz, 1H), 3.16 – 3.10 (m, 1H), 3.01 (dd, *J* = 13.1, 8.5 Hz, 1H), 2.79 (dd, *J* = 13.4, 3.0 Hz, 1H), 2.77 – 2.73 (m, 1H), 2.70 (dt, *J* = 11.7, 3.8 Hz, 1H), 2.58 (qd, *J* = 7.0, 6.1, 3.8 Hz, 2H), 2.50 – 2.45 (m, 1H), 2.45 – 2.40 (m, 1H), 2.28 (d, *J* = 3.6 Hz, 3H), 2.22 (ddd, *J* = 18.0, 10.9, 3.8 Hz, 1H), 2.05 (dt, *J* = 14.6, 7.5 Hz, 2H), 1.94 – 1.90 (m, 1H), 1.73 – 1.66 (m, 3H), 1.62 – 1.55 (m, 1H), 1.47 – 1.40 (m, 1H), 1.11 (d, *J* = 6.2 Hz, 3H), 0.90 – 0.87 (m, 3H). $^{13}C$ NMR (176 MHz, $CDCl_3$) δ 172.7, 170.6, 169.1, 168.3, 168.1, 159.7,

157.3, 153.1, 148.8, 147.3, 142.3, 136.5, 135.3, 134.5, 133.3, 129.8, 129.0, 120.2, 119.7, 113.9, 112.7, 111.6, 111.2, 104.9, 75.7, 70.3, 70.2, 69.7, 67.3, 60.8, 56.3, 55.9, 55.9, 55.8, 54.3, 54.1, 52.1, 50.8, 49.4, 49.3, 47.8, 45.7, 43.5, 41.6, 38.8, 38.3, 33.3, 31.3, 26.8, 25.3, 23.1, 20.9, 12.6. HRMS (ESI, m/z): calcd for $C_{56}H_{76}N_6O_{14}SNa^+$ $[M+Na]^+$: 1111.5032, found 1111.5033.

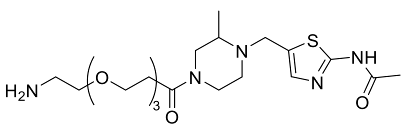


**16b**

**N-(5-((4-(3-(2-(2-(2-aminoethoxy)ethoxy)ethoxy)propanoyl)-2-methylpiperazin-1-yl)methyl)thiazol-2-yl)acetamide (16b)**

Intermediate **16b** was synthesized from **5** (20 mg, 79 μmol) and BocNH-PEG3-$CH_2CH_2COOH$ (25 mg, 79 μmol) using **General Method B**. Flash column chromatography (silica gel, MeOH in DCM: 0-7%) to give the **Boc-16b** (white solid, 23.7 mg, 54%). $^1H$ NMR (700 MHz $CDCl_3$) δ 11.07 (s, 1H), 7.19 (s, 1H), 4.12 (d, *J* = 12.9 Hz, 1H), 4.02 (d, *J* = 11.4 Hz, 1H), 4.01 – 3.96 (m, 1H), 3.76 (s, 1H), 3.61 (s, 10H), 3.54 – 3.52 (m, 2H), 3.32 – 3.28 (m, 3H), 3.14 (s, 1H), 3.03 (d, *J* = 11.5 Hz, 1H), 2.76 (d, *J* = 13.3 Hz, 1H), 2.73 (d, *J* = 11.3 Hz, 1H), 2.61 (s, 2H), 2.29 (s, 3H), 1.43 (s, 9H), 1.14 (s, 3H).

**16b** was synthesized from **Boc-16b** (20.0 mg, 36 μmol) using **General Method A**.

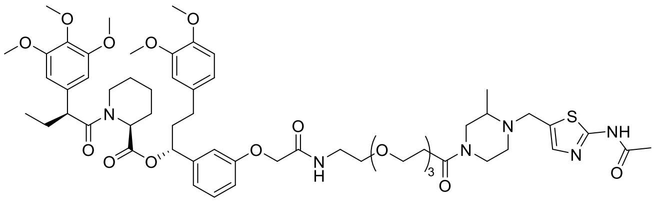


**D2**

**(1R)-1-(3-((15-(4-((2-acetamidothiazol-5-yl)methyl)-3-methylpiperazin-1-yl)-2,15-dioxo-6,9,12-trioxa-3-azapentadecyl)oxy)phenyl)-3-(3,4-dimethoxyphenyl)propyl (2S)-1-((S)-2-(3,4,5-trimethoxyphenyl)butanoyl)piperidine-2-carboxylate (D2)**

**D2** was synthesized from **16b** (20 mg, 36 μmol) and **AP1867** (25 mg, 36 μmol) using **General Method B.** pTLC (DCM: MeOH=15:1) to give the product **D2** (white solid, 16.9 mg, 41%). HPLC purity (254 nM)= 98.57%, RT= 15.438 min. $^1H$ NMR (700 MHz, $CDCl_3$) δ 11.72 (s, 1H), 7.18 (d, *J* = 1.5 Hz, 1H), 7.17 – 7.17 (m, 1H), 7.16 – 7.15 (m, 1H), 6.79 – 6.75 (m, 3H), 6.64 (dt, *J* = 9.6, 1.9 Hz, 3H), 6.40 (s, 2H), 5.61 (dd, *J* = 8.2, 5.5 Hz, 1H), 5.46 (d, *J* = 5.6 Hz, 1H), 4.51 – 4.48 (m, 2H), 4.11 – 4.07 (m, 1H), 4.01 (d, *J* = 15.7 Hz, 1H), 3.98 – 3.94 (m, 1H), 3.83 (dd, *J* = 11.9, 3.7 Hz, 9H), 3.77 (d, *J* = 0.9 Hz, 3H), 3.74 – 3.73 (m, 1H), 3.67 (d, *J* = 0.9 Hz, 5H), 3.59 (dq, *J* = 9.9, 5.5, 4.9 Hz, 14H), 3.55 – 3.53 (m, 1H), 3.26 (ddt, *J* = 12.7, 10.1, 2.7 Hz, 1H), 3.12 (ddd, *J* = 13.1, 9.6, 3.2 Hz, 1H), 3.02 (dd, *J* = 13.1, 8.5 Hz, 1H), 2.79 (dd, *J* = 13.4, 2.9 Hz, 1H), 2.75 (ddd, *J* = 11.7, 7.1, 3.3 Hz, 1H), 2.71 (dt, *J* = 11.6, 3.8 Hz, 1H), 2.58 (dd, *J* = 8.6, 5.7 Hz, 2H), 2.56 – 2.53 (m, 1H), 2.44 (ddd, *J* = 13.4, 8.3, 4.6 Hz, 1H), 2.28 (d, *J* = 2.5 Hz, 3H), 2.24 – 2.19 (m, 1H), 2.09 – 2.03 (m, 2H), 1.94 (s, 1H), 1.70 (dq, *J* = 11.6, 6.1, 4.8 Hz, 3H), 1.62 – 1.56 (m, 1H), 1.42 (tdt, *J* = 13.0, 8.1, 4.1 Hz,

1H), 1.12 (d, *J* = 6.2 Hz, 3H), 0.90 – 0.88 (m, 3H). $^{13}$C NMR (176 MHz, $CDCl_3$) δ 172.7, 170.6, 169.1, 168.3, 168.0, 159.6, 157.3, 153.2, 148.8, 147.3, 142.3, 136.5, 135.3, 134.6, 133.3, 129.8, 129.1, 120.2, 119.7, 114.0, 112.8, 111.6, 111.2, 104.9, 75.7, 70.5, 70.5, 70.4, 70.4, 70.4, 70.3, 70.2, 69.7, 67.3, 67.3, 60.8, 56.3, 55.9, 55.9, 55.9, 55.8, 54.3, 54.1, 52.1, 50.8, 50.5, 49.4, 49.3, 47.8, 45.8, 43.5, 41.6, 39.7, 38.8, 38.3, 33.4, 31.3, 26.8, 25.3, 23.2, 20.9, 12.6. HRMS (ESI, m/z): calcd for $C_{58}H_{80}N_6O_{15}SNa^+$ $[M+Na]^+$: 1155.5295, found 1155.5291.

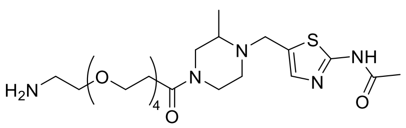


**16c**

**N-(5-((4-(1-amino-3,6,9,12-tetraoxapentadecan-15-oyl)-2-methylpiperazin-1-yl)methyl)thiazol-2-yl)acetamide (16c)**

Intermediate **16c** was synthesized from **5** (20 mg, 79 μmol) and BocNH-PEG4-$CH_2CH_2COOH$ (29 mg, 79 μmol) using **General Method B**. Flash column chromatography (silica gel, MeOH in DCM: 0-8%) to give the **Boc-16c** (white solid, 26.9 mg, 57%). $^1$H NMR (700 MHz, $CDCl_3$) δ 10.88 (s, 1H), 7.19 (s, 1H), 4.06 (d, *J* = 13.7 Hz, 1H), 4.00 (d, *J* = 14.6 Hz, 1H), 3.97 (d, *J* = 14.5 Hz, 1H), 3.77 – 3.75 (m, 1H), 3.64 – 3.59 (m, 14H), 3.53 (t, *J* = 4.7 Hz, 2H), 3.31 (t, *J* = 5.3 Hz, 2H), 3.27 (d, *J* = 9.3 Hz, 1H), 3.11 (d, *J* = 12.0 Hz, 1H), 3.03 (dd, *J* = 13.2, 8.6 Hz, 1H), 2.77 (d, *J* = 9.6 Hz, 1H), 2.74 (s, 1H), 2.65 – 2.56 (m, 2H), 2.28 (s, 2H), 1.43 (s, 9H), 1.14 (d, *J* = 6.2 Hz, 3H).

**16c** was synthesized from **Boc-16c** (20.0 mg, 33 μmol) using **General Method A**.

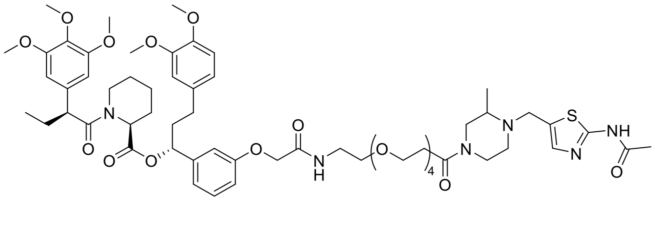

**D3**

**(1R)-1-(3-((18-(4-((2-acetamidothiazol-5-yl)methyl)-3-methylpiperazin-1-yl)-2,18-dioxo-6,9,12,15-tetraoxa-3-azaoctadecyl)oxy)phenyl)-3-(3,4-dimethoxyphenyl)propyl (2S)-1-((S)-2-(3,4,5-trimethoxyphenyl)butanoyl)piperidine-2-carboxylate (D3)**

**D3** was synthesized from **16c** (20 mg, 33 μmol) and **AP1867** (23 mg, 33 μmol) using **General Method B.** pTLC (DCM: MeOH=15:1) to give the product **D3** (white solid, 20.2 mg, 51%). HPLC purity (254 nM)= 97.88%, RT= 15.452 min. $^1$H NMR (700 MHz, $CDCl_3$) δ 11.75 (s, 1H), 7.17 (q, *J* = 7.6, 6.1 Hz, 3H), 6.79 – 6.75 (m, 3H), 6.65 – 6.62 (m, 3H), 6.40 (d, *J* = 5.0 Hz, 2H), 5.61 (dd, *J* = 8.2, 5.4 Hz, 1H), 5.47 – 5.44 (m, 1H), 4.49 (d, *J* = 8.8 Hz, 2H), 4.11 – 4.06 (m, 1H), 4.04 – 3.99 (m, 1H), 3.98 – 3.94 (m, 1H), 3.83 (dd, *J* = 12.0, 3.8 Hz, 9H), 3.77 (s, 3H), 3.75 – 3.73 (m, 1H), 3.67 (s, 5H), 3.62 – 3.57 (m, 18H), 3.55 – 3.54 (m, 1H), 3.29 – 3.22 (m, 1H), 3.12 (ddd, *J* = 13.0, 9.7, 3.2 Hz, 1H), 3.02 (dd, *J* = 13.2, 8.5 Hz, 1H), 2.81 – 2.78 (m, 1H), 2.77 – 2.74 (m, 1H), 2.71 (dt, *J* = 11.6, 3.8 Hz, 1H), 2.59 (dtd, *J* = 9.8, 6.6, 6.1, 3.5 Hz, 2H), 2.55 (dt,

*J* = 13.9, 3.9 Hz, 1H), 2.44 (td, *J* = 9.6, 8.0, 5.5 Hz, 1H), 2.28 (d, *J* = 2.1 Hz, 3H), 2.23 – 2.19 (m, 1H), 2.08 – 2.03 (m, 2H), 1.90 (dt, *J* = 15.5, 4.7 Hz, 1H), 1.70 (ddd, *J* = 13.6, 8.1, 4.9 Hz, 3H), 1.62 – 1.58 (m, 1H), 1.43 (ddt, *J* = 17.1, 13.1, 6.6 Hz, 1H), 1.12 (d, *J* = 6.2 Hz, 3H), 0.90 – 0.88 (m, 3H). $^{13}C$ NMR (176 MHz, $CDCl_3$) δ 172.7, 170.6, 169.2, 168.3, 168.0, 166.0, 159.6, 157.3, 153.2, 148.8, 147.3, 142.3, 136.5, 135.3, 134.6, 133.3, 129.8, 129.5, 129.1, 120.2, 119.7, 114.0, 112.8, 111.6, 111.2, 104.9, 75.7, 70.5, 70.5, 70.4, 70.4, 70.2, 69.8, 67.4, 67.3, 60.8, 56.3, 55.9, 55.9, 55.8, 54.3, 54.1, 52.1, 52.0, 51.2, 50.8, 50.5, 49.4, 49.3, 47.8, 45.8, 43.5, 41.6, 38.8, 38.3, 33.3, 31.3, 30.6, 29.3, 26.8, 25.3, 23.2, 20.9, 12.6. HRMS (ESI, m/z): calcd for $C_{60}H_{84}N_6O_{16}SNa^+$ $[M+Na]^+$: 1199.5557, found 1199.5553.

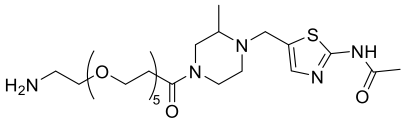


**16d**

**N-(5-((4-(1-amino-3,6,9,12,15-pentaoxaoctadecan-18-oyl)-2-methylpiperazin-1-yl)methyl)thiazol-2-yl)acetamide (16d)**

Intermediate **16d** was synthesized from **5** (20 mg, 79 μmol) and BocNH-PEG5-$CH_2CH_2COOH$ (32 mg, 79 μmol) using **General Method B**. Flash column chromatography (silica gel, MeOH in DCM: 0-8%) to give the **Boc-16d** (white solid, 29.0 mg, 57%). $^1H$ NMR (700 MHz, $CDCl_3$) δ 10.84 (s, 1H), 7.19 (s, 1H), 4.13 (d, *J* = 12.9 Hz, 1H), 4.07 (d, *J* = 12.8 Hz, 1H), 4.00 – 3.96 (m, 1H), 3.75 (d, *J* = 5.9 Hz, 1H), 3.65 – 3.61 (m, 18H), 3.53 (d, *J* = 5.0 Hz, 2H), 3.31 (q, *J* = 5.4 Hz, 3H), 3.11 (d, *J* = 11.9 Hz, 1H), 3.03 (dd, *J* = 13.1, 8.6 Hz, 1H), 2.76 (d, *J* = 13.1 Hz, 1H), 2.73 (d, *J* = 11.6 Hz, 1H), 2.64 – 2.56 (m, 2H), 2.29 (s, 2H), 1.43 (s, 9H), 1.14 (d, *J* = 6.1 Hz, 3H).

**16d** was synthesized from **Boc-16d** (20.0 mg, 31 μmol) using **General Method A**.

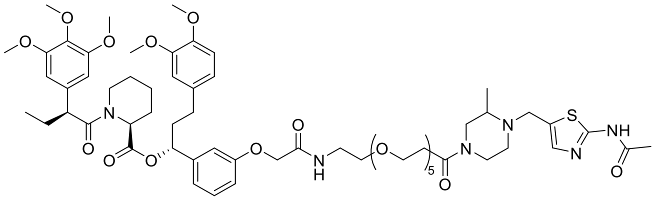

**D4**

**(1R)-1-(3-((21-(4-((2-acetamidothiazol-5-yl)methyl)-3-methylpiperazin-1-yl)-2,21-dioxo-6,9,12,15,18-pentaoxa-3-azahenicosyl)oxy)phenyl)-3-(3,4-dimethoxyphenyl)propyl (2S)-1-((S)-2-(3,4,5-trimethoxyphenyl)butanoyl)piperidine-2-carboxylate (D4)**

**D4** was synthesized from **16d** (20 mg, 31 μmol) and **AP1867** (22 mg, 31 μmol) using **General Method B.** pTLC (DCM: MeOH=15:1) to give the product **D4** (white solid, 16.9 mg, 45%). HPLC purity (254 nM)= 97.55%, RT= 15.454 min. $^1H$ NMR (700 MHz, $CDCl_3$) δ 11.64 (s, 1H), 7.18 (s, 1H), 7.17 – 7.15 (m, 1H), 6.80 – 6.75 (m, 3H), 6.64 (dt, *J* = 9.8, 2.2 Hz, 3H), 6.41 (d, *J* = 6.2 Hz, 2H), 5.61 (dd, *J* = 8.2, 5.4 Hz, 1H), 5.47 – 5.45 (m, 1H), 4.48 (s, 2H), 4.10 (d, *J* = 12.4 Hz, 1H), 4.06 – 4.00 (m, 1H), 3.99 – 3.95 (m, 1H), 3.86 – 3.82 (m, 9H), 3.77 (s, 3H), 3.75 – 3.73 (m, 1H), 3.68 (s, 5H), 3.63 – 3.57 (m, 22H), 3.54 (q, *J* = 2.9 Hz, 1H), 3.30 – 3.24 (m, 1H), 3.11 (td, *J* = 10.6,

9.7, 5.7 Hz, 1H), 3.02 (dd, *J* = 13.2, 8.6 Hz, 1H), 2.79 (dd, *J* = 13.4, 3.0 Hz, 1H), 2.78 – 2.74 (m, 1H), 2.72 (dq, *J* = 11.7, 4.6, 3.8 Hz, 1H), 2.62 – 2.57 (m, 2H), 2.56 – 2.52 (m, 1H), 2.47 – 2.43 (m, 1H), 2.29 – 2.27 (m, 3H), 2.25 – 2.21 (m, 1H), 2.08 – 2.04 (m, 2H), 1.94 – 1.92 (m, 1H), 1.74 – 1.67 (m, 3H), 1.60 (d, *J* = 13.7 Hz, 1H), 1.43 (dtd, *J* = 13.0, 9.0, 4.7 Hz, 1H), 1.13 (dd, *J* = 6.2, 3.5 Hz, 3H), 0.90 – 0.88 (m, 3H). $^{13}C$ NMR (176 MHz, $CDCl_3$) δ 172.7, 170.6, 169.3, 168.5, 168.0, 159.5, 157.3, 153.2, 148.8, 147.3, 142.3, 136.5, 135.3, 134.7, 133.3, 129.8, 128.9, 120.2, 119.7, 113.9, 112.8, 111.6, 111.2, 104.9, 75.7, 70.4, 70.2, 69.9, 67.3, 67.1, 60.8, 56.3, 56.0, 55.9, 55.8, 54.1, 52.1, 50.8, 50.5, 49.4, 49.2, 47.8, 45.8, 43.5, 41.7, 39.7, 38.8, 38.3, 33.2, 31.3, 29.3, 26.8, 25.3, 23.2, 21.0, 12.6. HRMS (ESI, m/z): calcd for $C_{62}H_{88}N_6O_{17}SNa^+$ $[M+Na]^+$: 1243.5819, found 1243.5813.

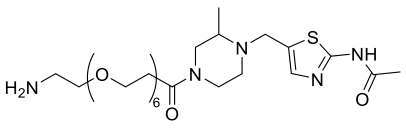


**16e**

**N-(5-((4-(1-amino-3,6,9,12,15,18-hexaoxahenicosan-21-oyl)-2-methylpiperazin-1-yl)methyl)thiazol-2-yl)acetamide (16e)**

Intermediate **16e** was synthesized from **5** (20 mg, 79 μmol) and BocNH-PEG6-$CH_2CH_2COOH$ (36 mg, 79 μmol) using **General Method B**. Flash column chromatography (silica gel, MeOH in DCM: 0-8%) to give the **Boc-16e** (white solid, 11.0 mg, 20%). $^1H$ NMR (700 MHz, $CDCl_3$) δ 7.21 (s, 1H), 4.07 (d, *J* = 12.3 Hz, 1H), 4.00 (s, 1H), 3.98 (d, *J* = 6.5 Hz, 1H), 3.75 (s, 3H), 3.64 (t, *J* = 15.2 Hz, 17H), 3.55 (s, 3H), 3.40 (d, *J* = 3.2 Hz, 1H), 3.30 (d, *J* = 9.5 Hz, 3H), 3.18 (dt, *J* = 7.4, 3.9 Hz, 1H), 3.03 (d, *J* = 2.3 Hz, 2H), 2.79 (d, *J* = 11.6 Hz, 1H), 2.76 – 2.74 (m, 1H), 2.61 (s, 2H), 2.28 (s, 3H), 1.48 (d, *J* = 6.7 Hz, 3H), 1.43 (s, 9H).

**16e** was synthesized from **Boc-16e** (20.0 mg, 29 μmol) using **General Method A**.

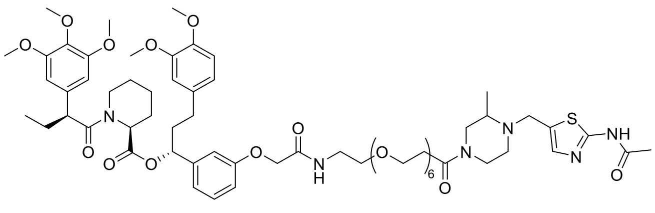


**D5**

**(1R)-1-(3-((24-(4-((2-acetamidothiazol-5-yl)methyl)-3-methylpiperazin-1-yl)-2,24-dioxo-6,9,12,15,18,21-hexaoxa-3-azatetracosyl)oxy)phenyl)-3-(3,4-dimethoxyphenyl)propyl (2S)-1-((S)-2-(3,4,5-trimethoxyphenyl)butanoyl)piperidine-2-carboxylate (D5)**

**D5** was synthesized from **16e** (10 mg, 15 μmol) and **AP1867** (10 mg, 15 μmol) using **General Method B.** pTLC (DCM: MeOH=15:1) to give the product **D5** (white solid, 9.8 mg, 53%). HPLC purity (254 nM)= 95.00%, RT= 15.425 min. $^1H$ NMR (700 MHz, $CDCl_3$) δ 11.22 (s, 1H), 8.09 (s, 1H), 7.22 (d, *J* = 5.5 Hz, 1H), 7.17 (s, 2H), 6.82 – 6.75 (m, 3H), 6.67 – 6.62 (m, 3H), 6.41 (d, *J* = 4.9 Hz, 2H), 5.62 (dd, *J* = 8.2, 5.4 Hz, 1H), 5.46 (d, *J* = 5.5 Hz, 1H), 4.50 (d, *J* = 10.8 Hz, 2H), 4.12 (d, *J* = 12.9 Hz, 1H), 4.05 (d, *J* = 13.2 Hz, 1H), 3.97 (dd, *J* = 20.5, 14.6 Hz, 1H), 3.86 – 3.82 (m, 9H), 3.77

(d, *J* = 8.5 Hz, 3H), 3.75 – 3.73 (m, 1H), 3.68 (s, 5H), 3.64 – 3.57 (m, 26H), 3.56 – 3.54 (m, 1H), 3.29 – 3.25 (m, 1H), 3.11 (t, *J* = 11.2 Hz, 1H), 3.02 (dd, *J* = 13.2, 8.6 Hz, 1H), 2.82 – 2.75 (m, 2H), 2.74 – 2.69 (m, 1H), 2.63 (dd, *J* = 15.9, 7.0 Hz, 1H), 2.58 – 2.52 (m, 2H), 2.46 – 2.42 (m, 1H), 2.27 (s, 3H), 2.25 – 2.21 (m, 1H), 2.07 (dt, *J* = 11.4, 4.9 Hz, 2H), 1.92 – 1.90 (m, 1H), 1.71 (dt, *J* = 13.7, 6.1 Hz, 3H), 1.61 (d, *J* = 13.6 Hz, 1H), 1.45 (dt, *J* = 6.8, 3.3 Hz, 1H), 1.13 (d, *J* = 6.2 Hz, 3H), 0.90 – 0.88 (m, 3H). $^{13}C$ NMR (176 MHz, $CDCl_3$) δ 172.7, 170.6, 169.3, 168.4, 168.0, 166.0, 159.3, 157.3, 153.2, 148.8, 147.3, 142.3, 136.5, 135.3, 134.2, 133.3, 129.5, 120.2, 119.7, 113.9, 112.8, 111.6, 111.2, 104.9, 75.7, 70.3, 70.3, 70.2, 70.1, 69.8, 67.8, 67.3, 60.9, 60.8, 56.3, 56.0, 55.9, 55.8, 54.1, 52.1, 50.8, 49.6, 49.3, 49.2, 47.8, 45.8, 43.5, 41.7, 38.9, 38.8, 38.3, 33.3, 31.3, 30.6, 29.7, 29.0, 26.8, 25.3, 24.0, 23.0, 21.0, 12.6. HRMS (ESI, m/z): calcd for $C_{64}H_{92}N_6O_{18}SNa^+$ $[M+Na]^+$: 1287.6081, found 1287.6082.

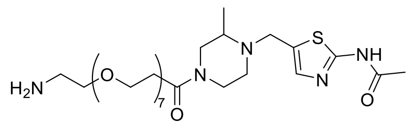

**16f**

##### N-(5-((4-(1-amino-3,6,9,12,15,18,21-heptaoxatetracosan-24-oyl)-2-methylpiperazin-1-yl)methyl)thiazol-2-yl)acetamide (16f)

Intermediate **16f** was synthesized from **5** (20 mg, 79 μmol) and BocNH-PEG7-$CH_2CH_2COOH$ (39 mg, 79 μmol) using **General Method B**. Flash column chromatography (silica gel, MeOH in DCM: 0-8%) to give the **Boc-16f** (white solid, 34.1 mg, 59%). $^1H$ NMR (700 MHz, $CDCl_3$) δ 10.83 (s, 1H), 7.19 (s, 1H), 4.14 – 4.11 (m, 1H), 4.06 (d, *J* = 13.2 Hz, 1H), 3.98 (dd, *J* = 23.0, 14.6 Hz, 1H), 3.76 – 3.74 (m, 1H), 3.64 (d, *J* = 3.1 Hz, 12H), 3.63 – 3.61 (m, 12H), 3.59 (d, *J* = 1.8 Hz, 2H), 3.54 (d, *J* = 5.1 Hz, 2H), 3.31 (t, *J* = 5.5 Hz, 2H), 3.29 – 3.24 (m, 1H), 3.14 – 3.10 (m, 1H), 3.03 (dd, *J* = 13.1, 8.6 Hz, 1H), 2.76 (td, *J* = 8.9, 8.1, 4.2 Hz, 1H), 2.75 – 2.71 (m, 1H), 2.65 – 2.57 (m, 2H), 2.28 (s, 3H), 1.43 (s, 9H), 1.14 (dd, *J* = 6.2, 2.0 Hz, 3H).

**16f** was synthesized from **Boc-16f** (20.0 mg, 27 μmol) using **General Method A**.

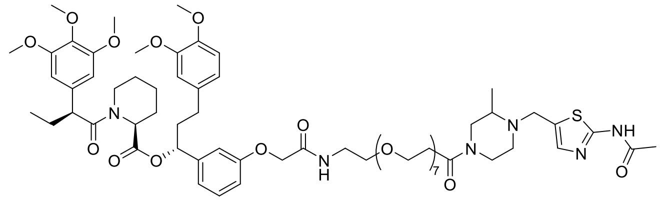

**D6**

##### (1R)-1-(3-((27-(4-((2-acetamidothiazol-5-yl)methyl)-3-methylpiperazin-1-yl)-2,27-dioxo-6,9,12,15,18,21,24-heptaoxa-3-azaheptacosyl)oxy)phenyl)-3-(3,4-dimethoxyphenyl)propyl (2S)-1-((S)-2-(3,4,5-trimethoxyphenyl)butanoyl)piperidine-2-carboxylate (D6)

**D6** was synthesized from **16f** (20 mg, 27 μmol) and **AP1867** (19 mg, 27 μmol) using **General Method B.** pTLC (DCM: MeOH=15:1) to give the product **D6** (white solid, 17.7 mg, 49%). HPLC purity (254 nM)= 95.05%, RT= 15.438 min. $^1H$ NMR (700 MHz, $CDCl_3$) δ 11.69 (s, 1H), 7.18 (s, 1H), 7.17 (s, 1H), 7.15 (d, *J* = 5.4 Hz, 1H), 6.80 – 6.75 (m, 3H), 6.66 – 6.62 (m, 3H), 6.40 (s, 2H), 5.61 (dd, *J* = 8.2, 5.4 Hz, 1H), 5.47 –

5.44 (m, 1H), 4.48 (s, 2H), 4.10 (d, *J* = 11.4 Hz, 1H), 4.03 (dd, *J* = 13.3, 3.9 Hz, 1H), 3.98 (dd, *J* = 17.5, 14.6 Hz, 1H), 3.83 (dd, *J* = 12.2, 4.3 Hz, 9H), 3.77 (s, 3H), 3.74 (dd, *J* = 4.4, 2.5 Hz, 1H), 3.67 (s, 5H), 3.62 (dd, *J* = 9.3, 5.3 Hz, 26H), 3.59 (d, *J* = 4.9 Hz, 4H), 3.54 (d, *J* = 5.0 Hz, 1H), 3.27 (ddd, *J* = 13.1, 9.9, 3.2 Hz, 1H), 3.11 (ddd, *J* = 13.1, 9.6, 3.3 Hz, 1H), 3.05 – 3.00 (m, 1H), 2.80 – 2.78 (m, 1H), 2.78 – 2.75 (m, 1H), 2.75 – 2.70 (m, 1H), 2.62 – 2.56 (m, 2H), 2.53 (td, *J* = 9.4, 5.0 Hz, 1H), 2.44 (ddd, *J* = 11.5, 7.1, 3.6 Hz, 1H), 2.28 (d, *J* = 4.0 Hz, 3H), 2.24 – 2.20 (m, 1H), 2.08 – 2.05 (m, 2H), 1.94 – 1.89 (m, 1H), 1.71 (ddt, *J* = 17.2, 13.2, 6.9 Hz, 3H), 1.62 – 1.59 (m, 1H), 1.42 (tdt, *J* = 13.0, 8.4, 4.2 Hz, 1H), 1.13 (dd, *J* = 6.2, 4.2 Hz, 3H), 0.90 – 0.88 (m, 3H). $^{13}$C NMR (176 MHz, $CDCl_3$) δ 172.7, 170.6, 169.4, 168.4, 168.1, 159.5, 157.3, 153.1, 148.8, 147.3, 142.3, 136.5, 135.3, 134.7, 133.3, 129.8, 128.9, 120.2, 119.7, 113.9, 112.8, 111.6, 111.2, 104.9, 75.7, 70.3, 70.2, 70.2, 70.1, 69.8, 67.3, 60.8, 56.3, 55.9, 55.9, 55.9, 55.8, 54.2, 54.1, 52.1, 50.8, 50.4, 49.4, 49.3, 47.8, 45.8, 43.5, 41.7, 38.8, 38.3, 36.7, 33.2, 31.9, 31.3, 29.7, 26.8, 25.3, 23.2, 20.9, 12.6. HRMS (ESI, m/z): calcd for $C_{66}H_{96}N_6O_{19}SNa^+$ $[M+Na]^+$: 1331.6343, found 1331.6331.

**Scheme S8.** Synthesis of **E1-E2**

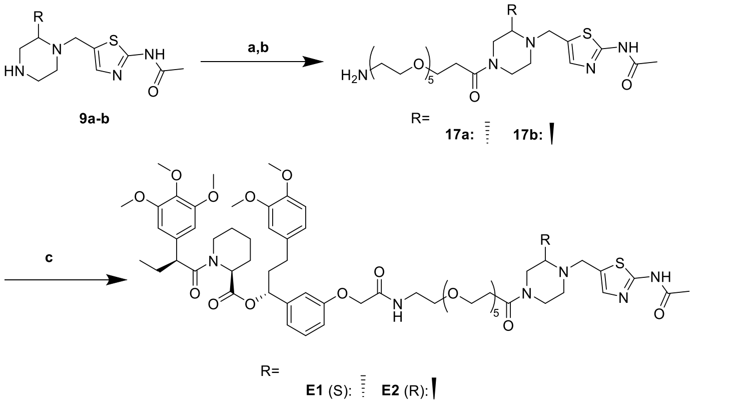


Reaction and conditions: a) HATU, DIPEA, DCM, rt, 13h, 28%; b) TFA, DCM, rt, 1h; c) AP1867, HATU, DIPEA, DCM, 15h, 44%.

**E1-E2** were synthesized following the same procedure for preparing **A1**.

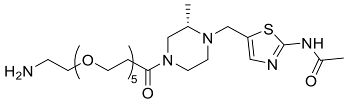


**17a**

##### (S)-N-(5-((4-(1-amino-3,6,9,12,15-pentaoxaoctadecan-18-oyl)-2-methylpiperazin-1-yl)methyl)thiazol-2-yl)acetamide (17a)

Intermediate **17a** was synthesized from **9a** (20 mg, 79 μmol) and BocNH-PEG5-$CH_2CH_2COOH$ (32 mg, 79 μmol) using **General Method B**. Flash column chromatography (silica gel, MeOH in DCM: 0-8%) to give the **Boc-17a** (white solid, 14.3 mg, 28%). $^1$H NMR (700 MHz, $CDCl_3$) δ 11.12 (s, 1H), 7.21 (s, 1H), 4.14 (d, *J* = 10.7 Hz, 1H), 4.05 (d, *J* = 19.0 Hz, 1H), 4.02 – 3.95 (m, 1H), 3.75 (d, *J* = 6.4 Hz, 1H), 3.65 – 3.61 (m, 16H), 3.59 (s, 2H), 3.53 (t, *J* = 5.1 Hz, 2H), 3.31 (d, *J* = 5.0 Hz, 2H), 3.30 (d, *J* = 5.4 Hz, 1H), 3.15 (s, 1H), 3.08 (s, 1H), 2.82 (s, 1H), 2.76 (s, 1H), 2.64 (s, 2H), 2.29 (s, 3H), 1.43 (s, 9H), 1.16 (s, 3H).

**17a** was synthesized from **Boc-17a** (14.3 mg, 22 μmol) using **General Method A**.

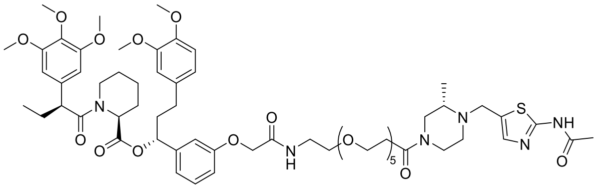

**E1**

##### (R)-1-(3-((21-((S)-4-((2-acetamidothiazol-5-yl)methyl)-3-methylpiperazin-1-yl)-2,21-dioxo-6,9,12,15,18-pentaoxa-3-azahenicosyl)oxy)phenyl)-3-(3,4-dimethoxyphenyl)propyl (S)-1-((S)-2-(3,4,5-trimethoxyphenyl)butanoyl)piperidine-2-carboxylate (E1)

**E1** was synthesized from **17a** (12 mg, 19 µmol) and **AP1867** (13 mg, 19 µmol) using **General Method B.** pTLC (DCM: MeOH=15:1) to give the product **E1** (white solid, 10.1 mg, 44%). HPLC purity (254 nM)= 96.25%, RT= 15.291 min. $^{1}$H NMR (700 MHz, $CDCl_3$) δ 11.39 (s, 1H), 7.19 – 7.16 (m, 2H), 7.13 (q, *J* = 6.3 Hz, 1H), 6.80 – 6.75 (m, 3H), 6.66 – 6.63 (m, 3H), 6.41 (d, *J* = 2.7 Hz, 2H), 5.62 (dd, *J* = 8.2, 5.5 Hz, 1H), 5.48 – 5.45 (m, 1H), 4.51 – 4.48 (m, 2H), 4.13 – 4.09 (m, 1H), 4.04 (d, *J* = 13.2 Hz, 1H), 3.98 (dd, *J* = 19.5, 14.5 Hz, 1H), 3.86 – 3.82 (m, 9H), 3.78 (s, 3H), 3.75 (d, *J* = 3.6 Hz, 1H), 3.68 (s, 5H), 3.63 – 3.58 (m, 22H), 3.54 (d, *J* = 5.1 Hz, 1H), 3.27 (t, *J* = 11.0 Hz, 1H), 3.15 – 3.10 (m, 1H), 3.03 (dd, *J* = 13.1, 8.5 Hz, 1H), 2.79 (dd, *J* = 13.5, 3.0 Hz, 1H), 2.77 – 2.74 (m, 1H), 2.74 – 2.70 (m, 1H), 2.59 (dt, *J* = 13.4, 6.9 Hz, 2H), 2.54 (td, *J* = 9.5, 5.0 Hz, 1H), 2.46 – 2.43 (m, 1H), 2.28 (s, 3H), 2.24 – 2.20 (m, 1H), 2.06 (dq, *J* = 15.3, 7.6, 7.2 Hz, 2H), 1.94 – 1.89 (m, 1H), 1.73 – 1.68 (m, 3H), 1.60 (d, *J* = 14.6 Hz, 1H), 1.46 – 1.39 (m, 1H), 1.13 (d, *J* = 6.2 Hz, 3H), 0.91 – 0.88 (m, 3H). $^{13}$C NMR (176 MHz, $CDCl_3$) δ 172.6, 170.6, 169.2, 168.3, 167.9, 159.3, 157.3, 153.2, 148.8, 147.3, 142.3, 136.5, 135.3, 134.8, 133.3, 129.8, 120.2, 119.7, 114.0, 112.8, 111.6, 111.2, 104.9, 75.7, 70.5, 70.5, 70.4, 70.4, 70.3, 69.7, 67.3, 60.9, 60.8, 56.3, 56.0, 55.9, 55.8, 54.2, 54.1, 52.1, 50.8, 49.4, 49.3, 47.8, 45.8, 43.5, 41.6, 38.8, 38.3, 33.4, 31.3, 29.3, 26.8, 25.3, 23.2, 21.0, 12.6. HRMS (ESI, m/z): calcd for $C_{62}H_{88}N_6O_{17}SNa^+$ $[M+Na]^+$: 1243.5819, found 1243.5816.

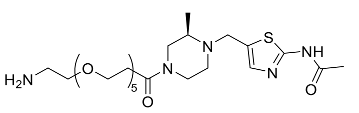

**17b**

##### (R)-N-(5-((4-(1-amino-3,6,9,12,15-pentaoxaoctadecan-18-oyl)-2-methylpiperazin-1-yl)methyl)thiazol-2-yl)acetamide (17b)

Intermediate **17b** was synthesized from **9b** (20 mg, 79 µmol) and BocNH-PEG5-$CH_2CH_2COOH$ (32 mg, 79 µmol) using **General Method B**. Flash column chromatography (silica gel, MeOH in DCM: 0-8%) to give the **Boc-17b** (white solid, 12.5 mg, 25%). $^{1}$H NMR (700 MHz, $CDCl_3$) δ 11.31 (s, 1H), 7.21 (s, 1H), 4.05 (s, 1H), 4.02 (s, 1H), 3.76 (d, *J* = 6.6 Hz, 1H), 3.66 – 3.60 (m, 18H), 3.53 (t, *J* = 5.1 Hz, 2H), 3.31 (t, *J* = 5.2 Hz, 2H), 3.29 (s, 1H), 3.15 (s, 1H), 3.04 (s, 1H), 2.85 (s, 1H), 2.77 (s, 1H), 2.66 – 2.56 (m, 2H), 2.29 (s, 3H), 1.43 (s, 9H), 1.16 (s, 3H).

**17b** was synthesized from **Boc-17b** (12.5 mg, 19 µmol) using **General Method A**.

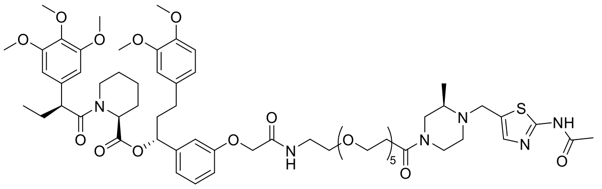

E2

**(R)-1-(3-((21-((R)-4-((2-acetamidothiazol-5-yl)methyl)-3-methylpiperazin-1-yl)-2,21-dioxo-6,9,12,15,18-pentaoxa-3-azahenicosyl)oxy)phenyl)-3-(3,4-dimethoxyphenyl)propyl (S)-1-((S)-2-(3,4,5-trimethoxyphenyl)butanoyl)piperidine-2-carboxylate (E2)**

**E2** was synthesized from **17b** (10 mg, 16 μmol) and **AP1867** (11 mg, 16 μmol) using **General Method B.** pTLC (DCM: MeOH=15:1) to give the product **E2** (white solid, 10.2 mg, 54%). HPLC purity (254 nM)= 97.72%, RT= 15.320 min. $^{1}$H NMR (700 MHz, $CDCl_3$) δ 11.38 (s, 1H), 7.19 – 7.16 (m, 2H), 7.12 (q, *J* = 6.1 Hz, 1H), 6.80 – 6.76 (m, 3H), 6.66 – 6.63 (m, 3H), 6.41 (d, *J* = 2.7 Hz, 2H), 5.62 (dd, *J* = 8.2, 5.5 Hz, 1H), 5.47 (d, *J* = 5.7 Hz, 1H), 4.51 – 4.48 (m, 2H), 4.11 (d, *J* = 12.9 Hz, 1H), 4.06 – 4.00 (m, 1H), 4.00 – 3.95 (m, 1H), 3.86 – 3.83 (m, 9H), 3.78 (s, 3H), 3.75 (d, *J* = 3.6 Hz, 1H), 3.68 (s, 5H), 3.63 – 3.58 (m, 22H), 3.55 (dd, *J* = 5.1, 3.1 Hz, 1H), 3.28 (s, 1H), 3.15 – 3.10 (m, 1H), 3.03 (dd, *J* = 13.1, 8.5 Hz, 1H), 2.79 (dd, *J* = 13.3, 3.0 Hz, 1H), 2.76 (d, *J* = 3.1 Hz, 1H), 2.72 (d, *J* = 11.8 Hz, 1H), 2.59 (dt, *J* = 13.7, 7.2 Hz, 2H), 2.54 (dt, *J* = 9.6, 4.8 Hz, 1H), 2.44 (ddd, *J* = 13.8, 8.4, 4.7 Hz, 1H), 2.28 (d, *J* = 2.4 Hz, 3H), 2.22 (dd, *J* = 10.5, 4.8 Hz, 1H), 2.10 – 2.04 (m, 2H), 1.95 – 1.89 (m, 1H), 1.71 (dt, *J* = 13.7, 6.7 Hz, 3H), 1.62 – 1.59 (m, 1H), 1.49 – 1.42 (m, 1H), 1.13 (d, *J* = 6.2 Hz, 3H), 0.90 (d, *J* = 7.3 Hz, 3H). $^{13}$C NMR (176 MHz, $CDCl_3$) δ 172.6, 170.6, 169.2, 168.2, 167.9, 159.4, 157.3, 153.2, 148.8, 147.3, 142.3, 136.5, 135.3, 134.8, 133.3, 129.8, 129.5, 120.2, 119.7, 114.0, 112.8, 111.6, 111.2, 104.9, 75.7, 70.5, 70.5, 70.4, 70.4, 70.3, 69.7, 67.4, 67.3, 60.9, 60.8, 56.3, 56.0, 55.9, 55.8, 54.1, 53.5, 52.1, 50.8, 49.6, 49.4, 43.5, 38.8, 38.3, 33.4, 31.3, 29.3, 26.8, 25.3, 23.2, 21.0, 12.6. HRMS (ESI, m/z): calcd for $C_{62}H_{88}N_6O_{17}SNa^+$ [M+Na]$^+$: 1243.5819, found 1243.5815.

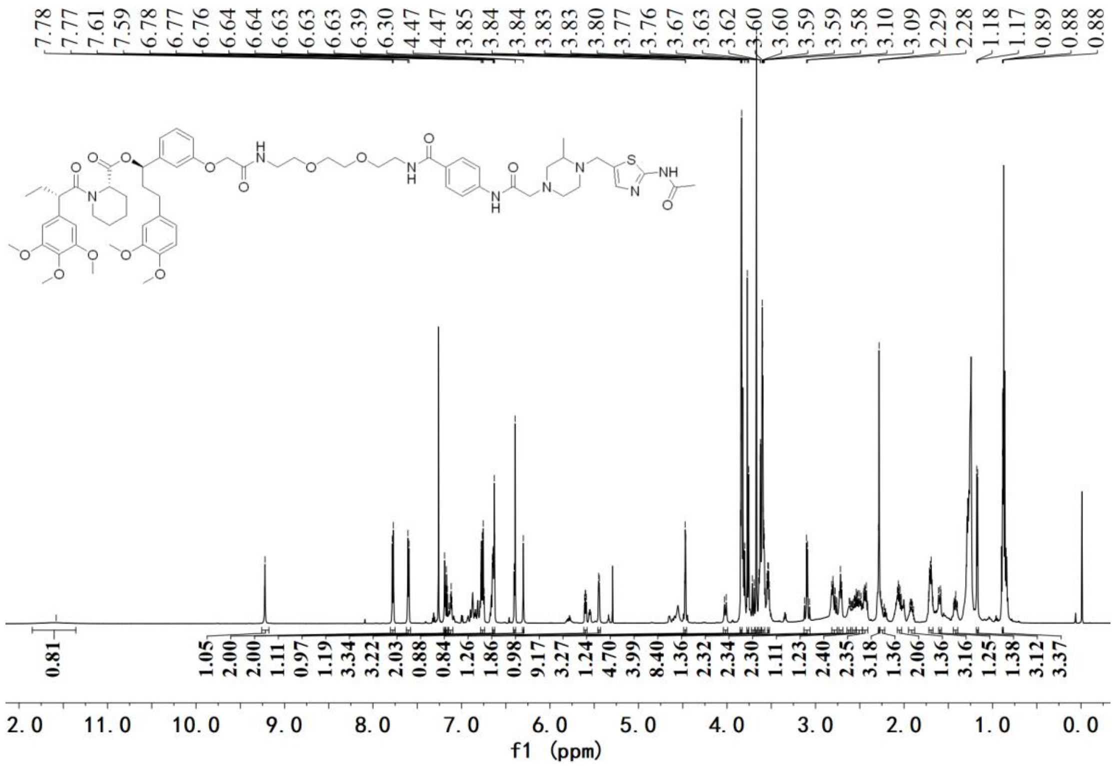

[1]H NMR spectrum of **A1**

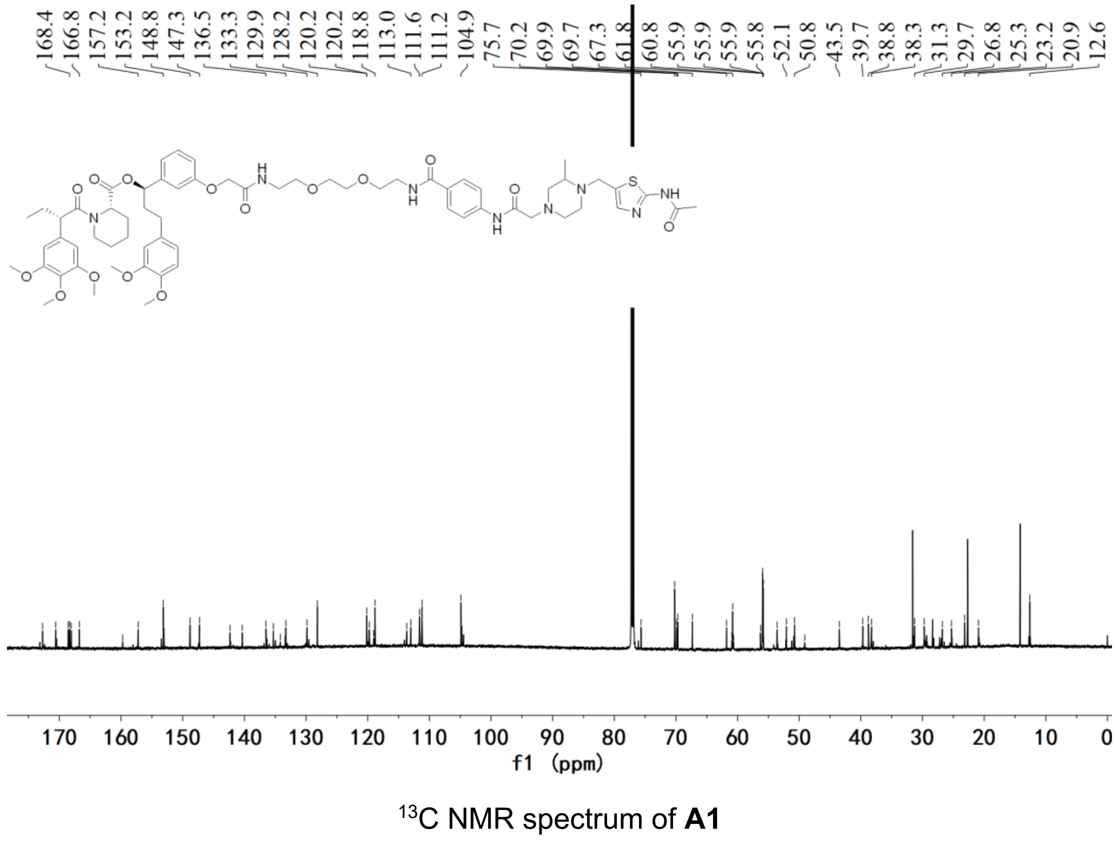

[13]C NMR spectrum of **A1**

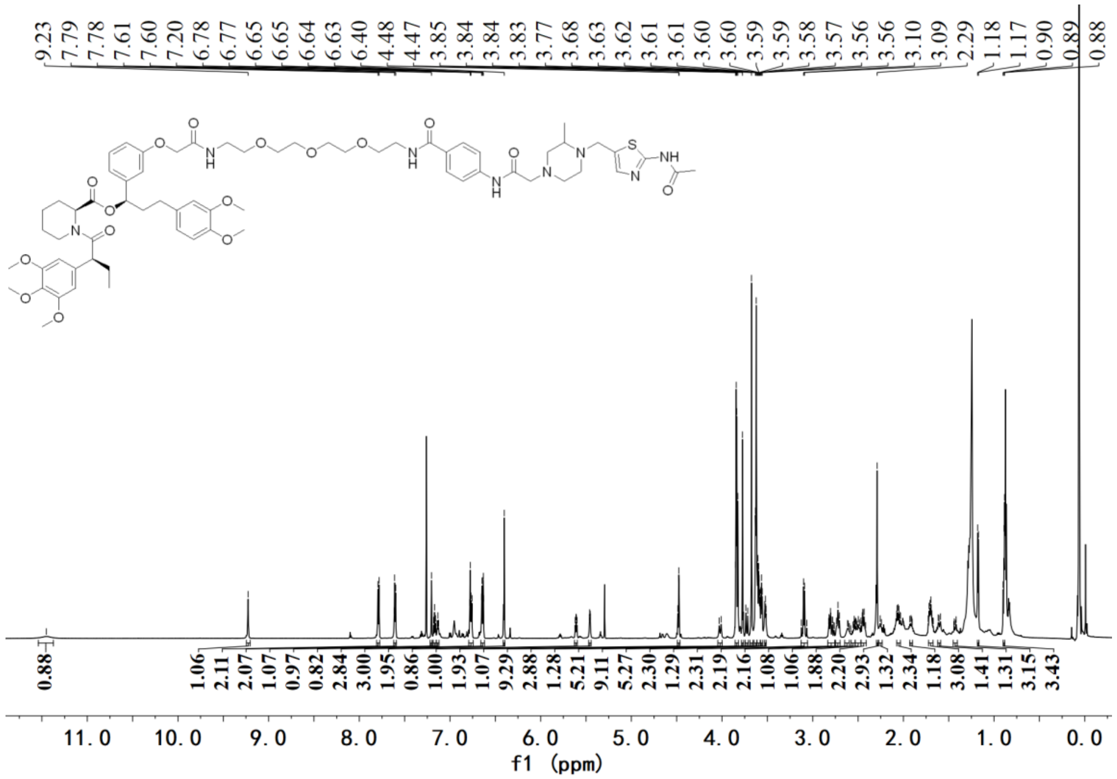


[1]H NMR spectrum of **A2**

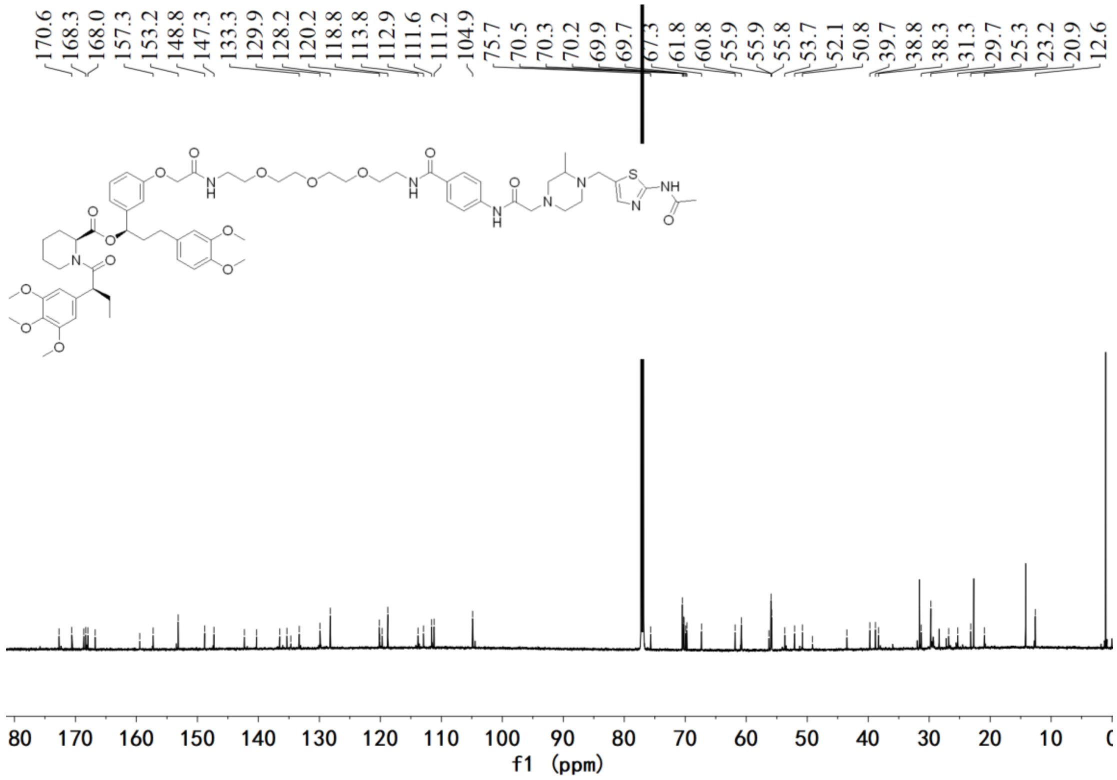


[13]C NMR spectrum of **A2**

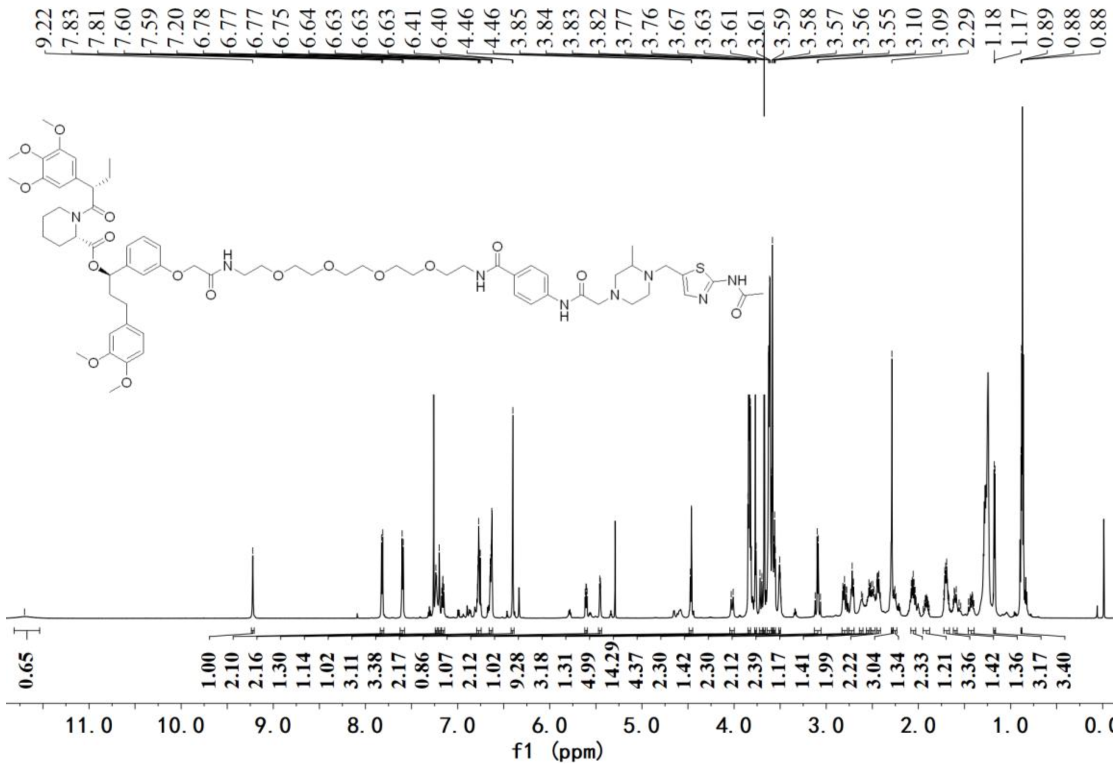


[1]H NMR spectrum of **A3**

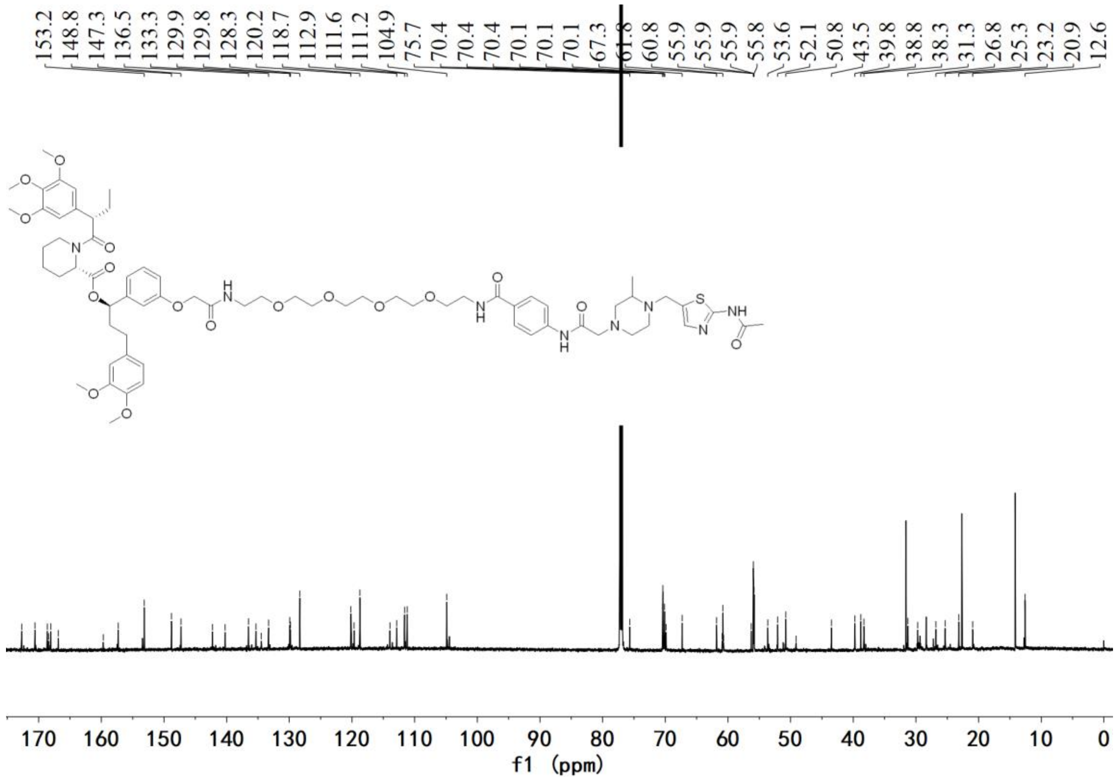


[13]C NMR spectrum of **A3**

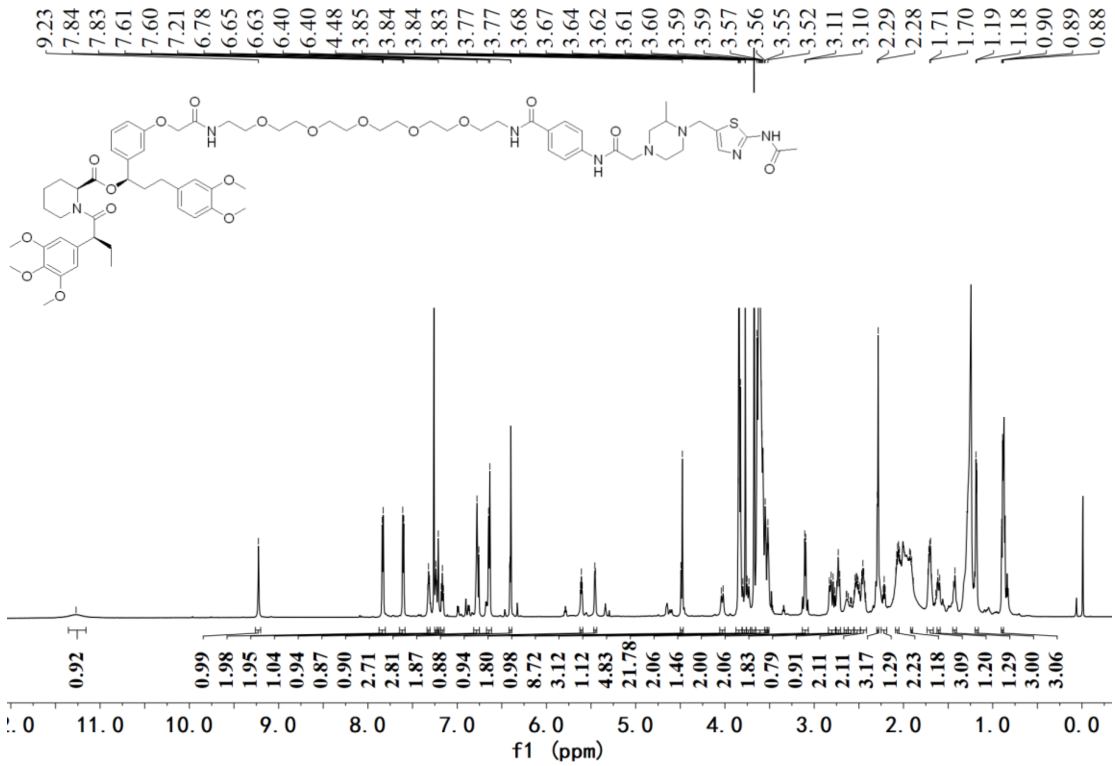

[1]H NMR spectrum of **A4**

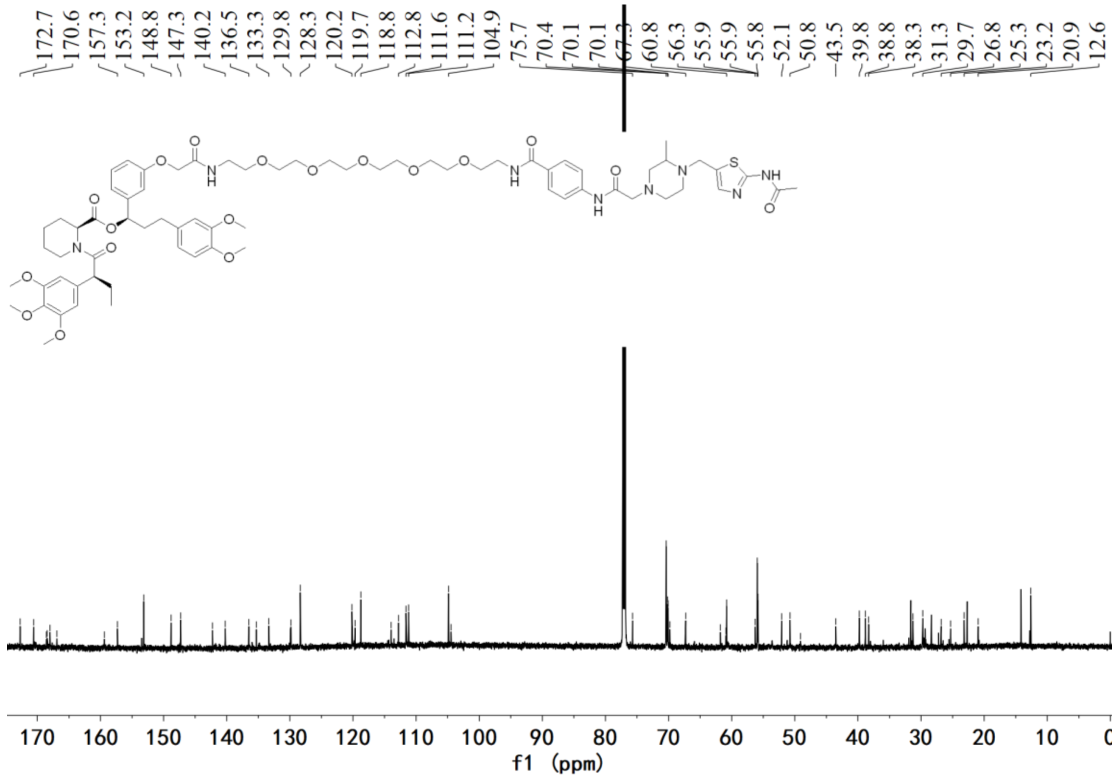

[13]C NMR spectrum of **A4**

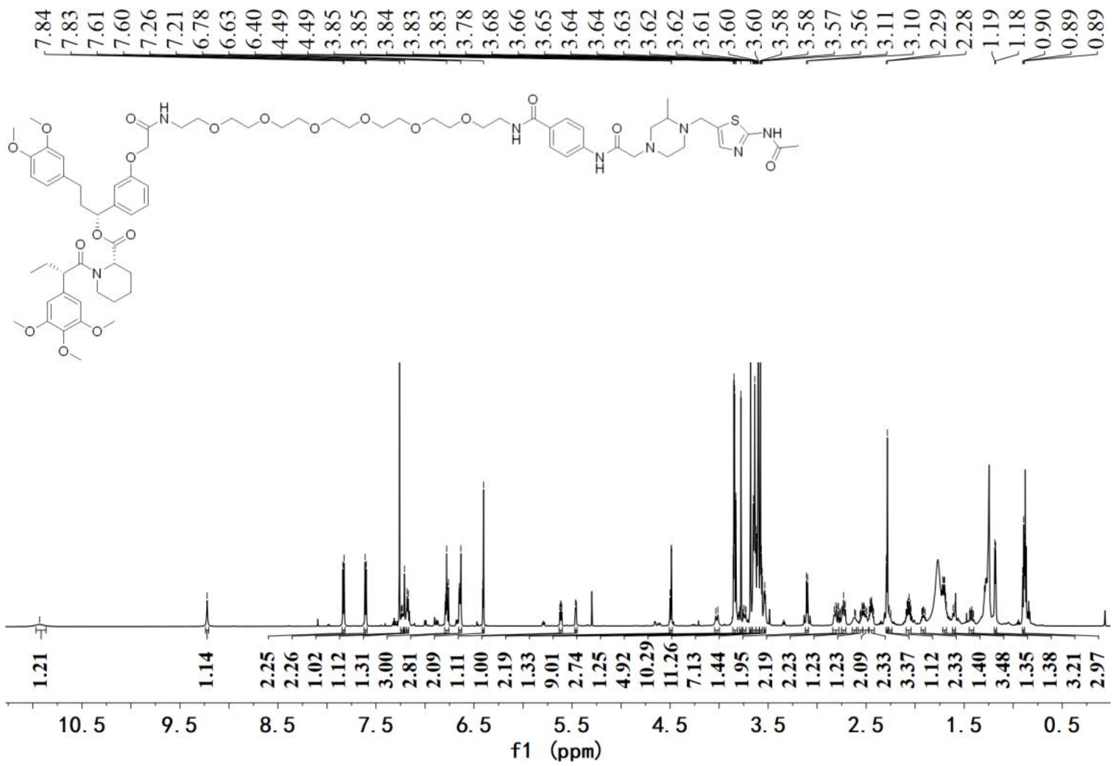

[1]H NMR spectrum of **A5**

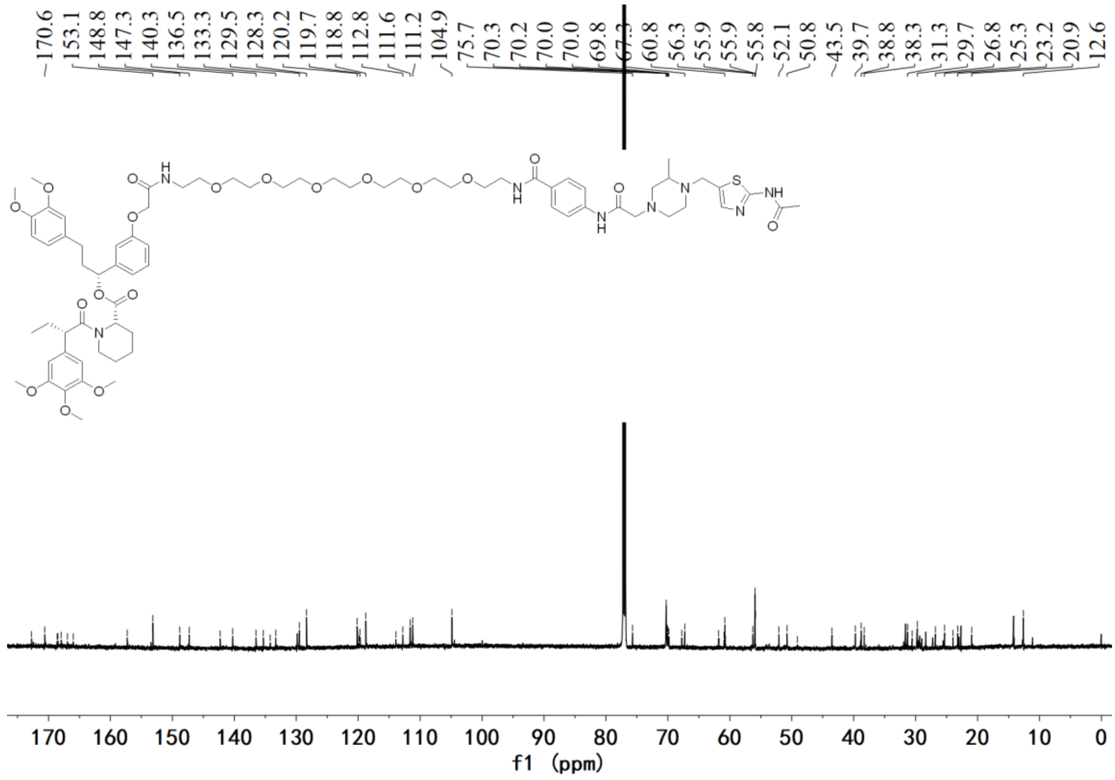

[13]C NMR spectrum of **A5**

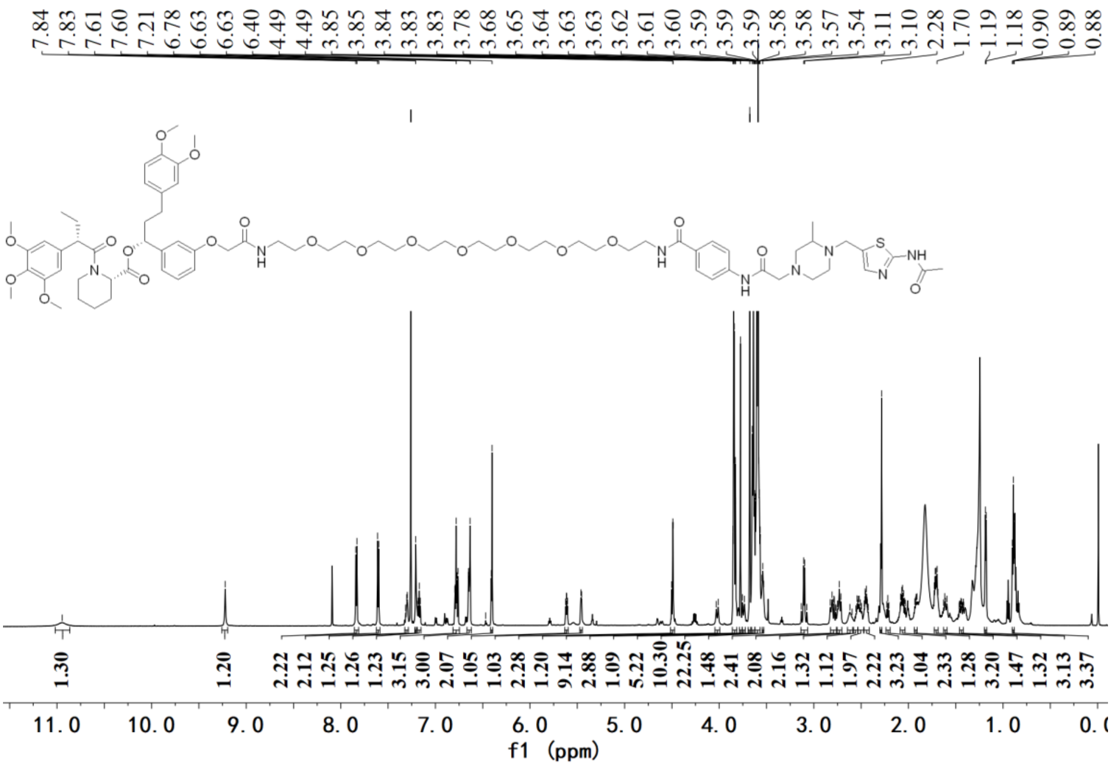


$^{1}H$ NMR spectrum of **A6**

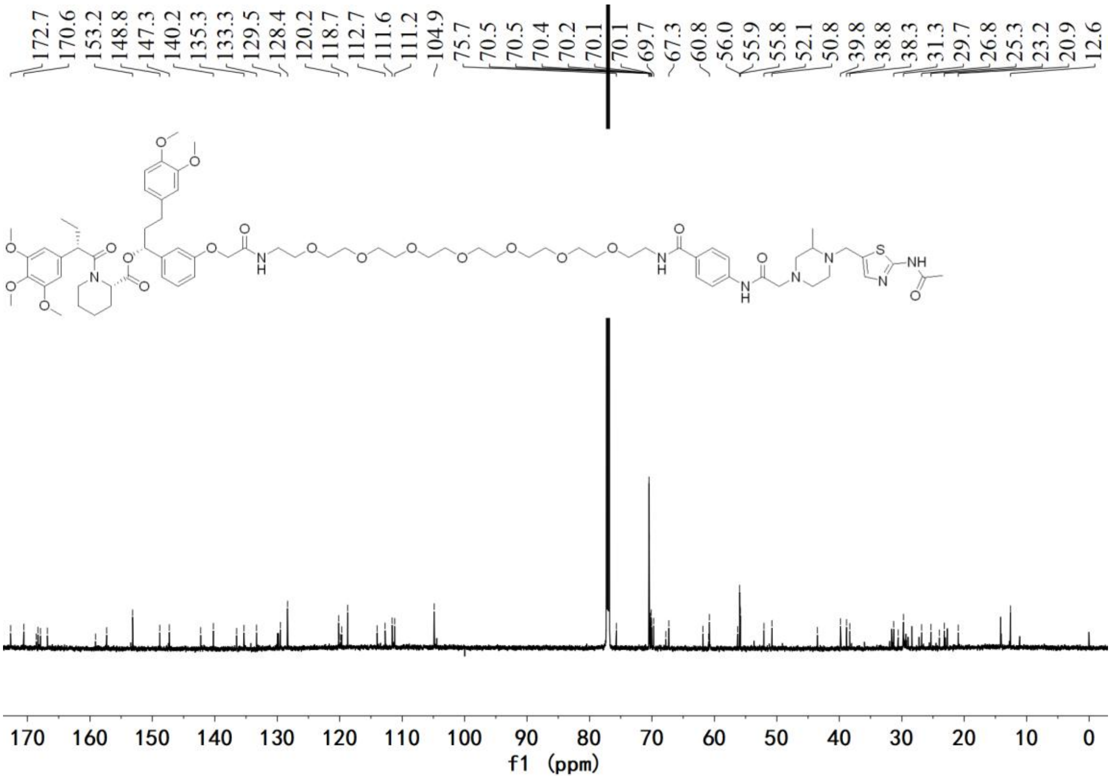


$^{13}C$ NMR spectrum of **A6**

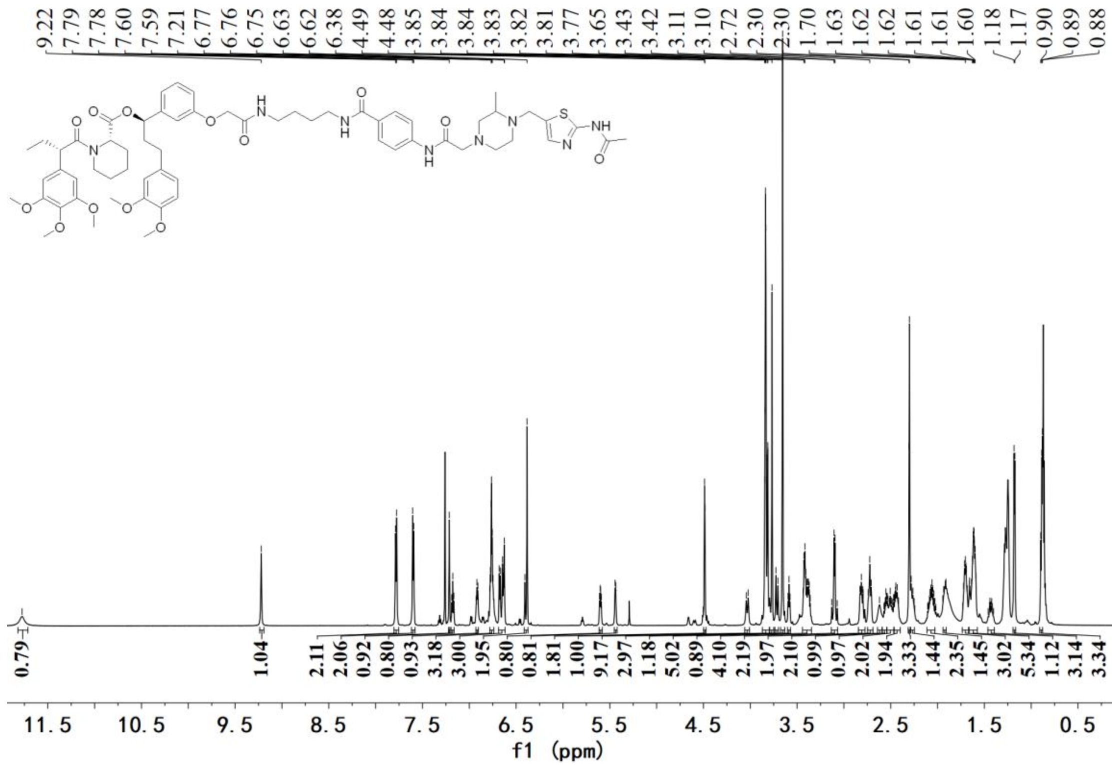

$^{1}$H NMR spectrum of **A7**

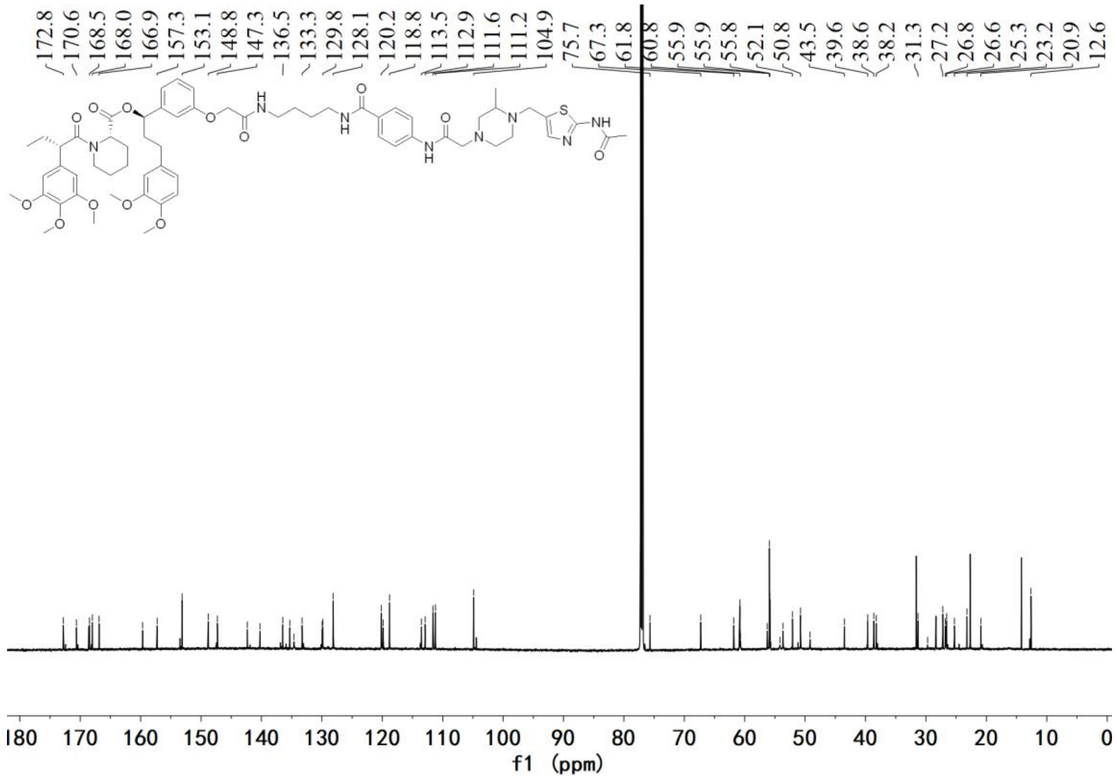

$^{13}$C NMR spectrum of **A7**

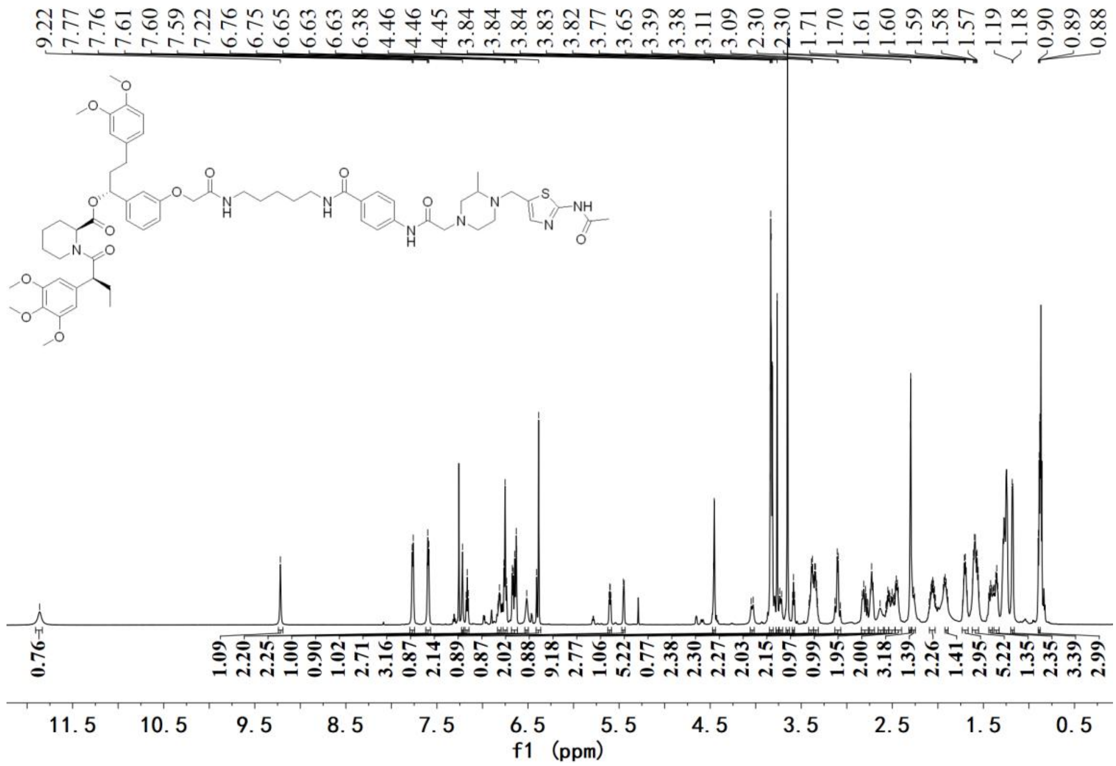


$^{1}$H NMR spectrum of **A8**

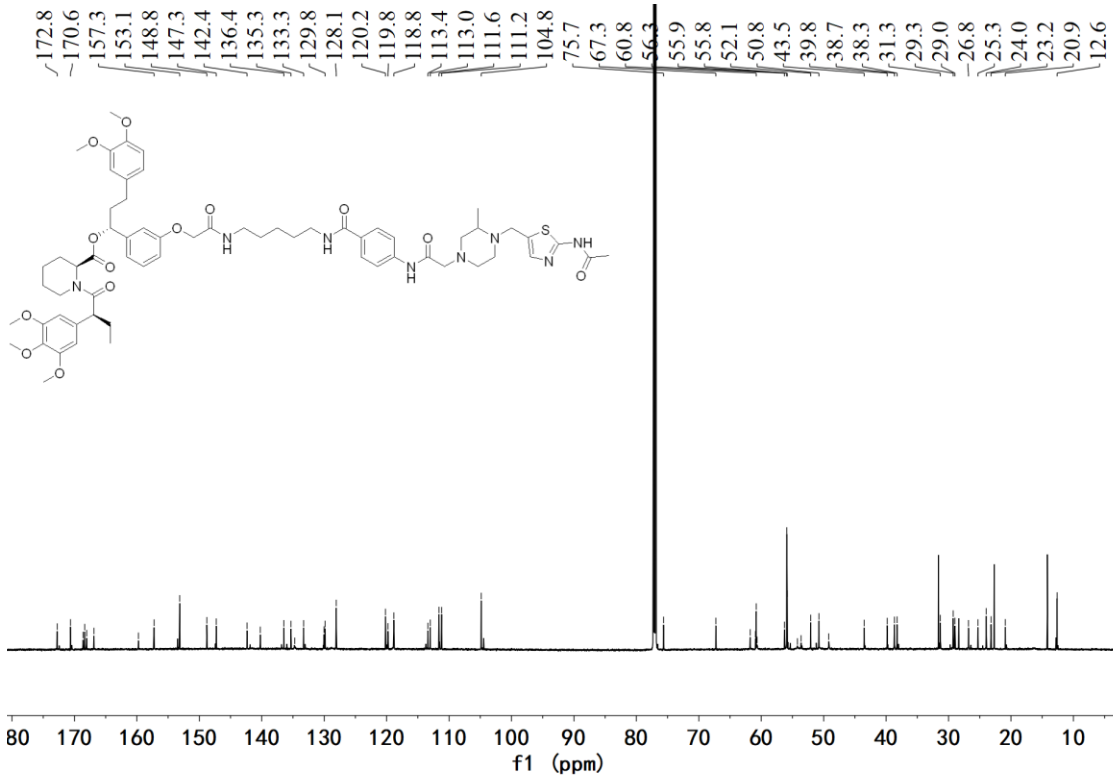


$^{13}$C NMR spectrum of **A8**

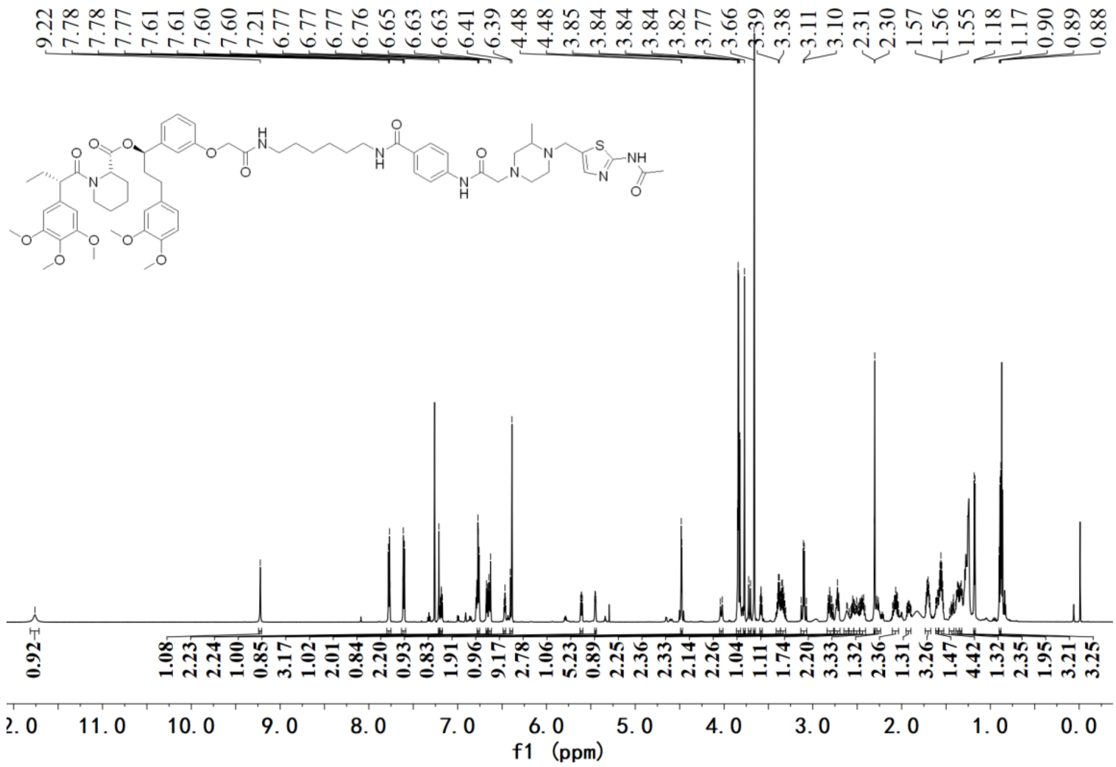


[1]H NMR spectrum of **A9**

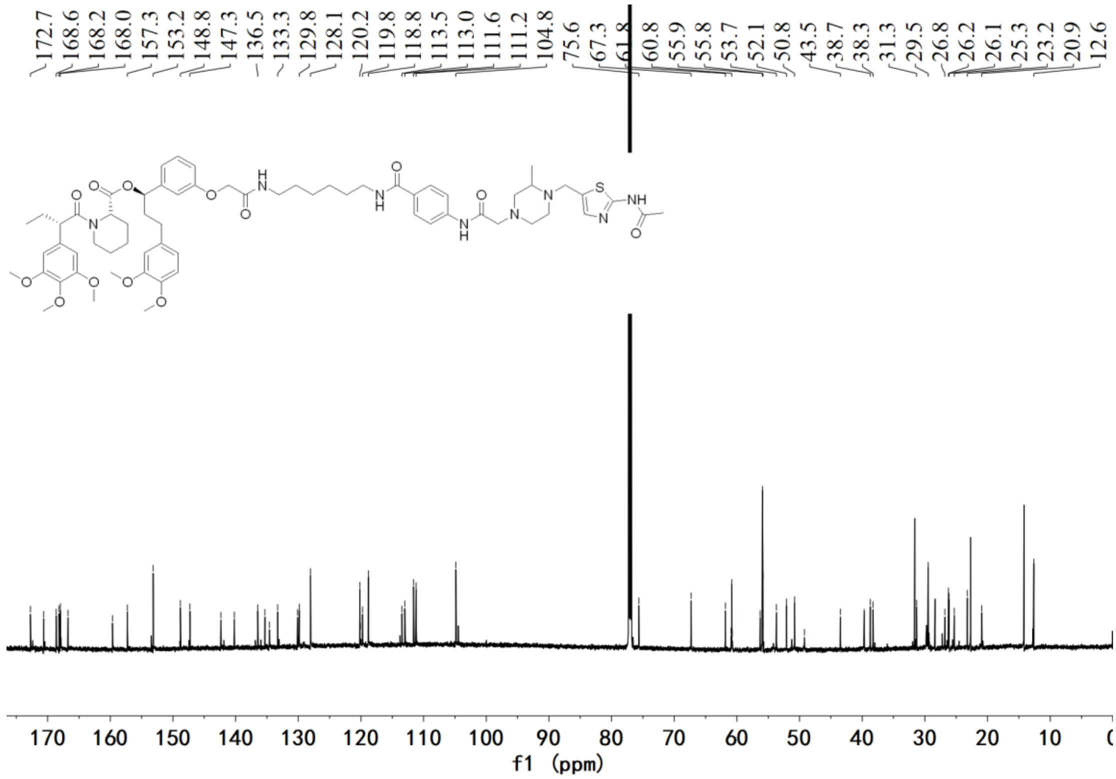


[13]C NMR spectrum of **A9**

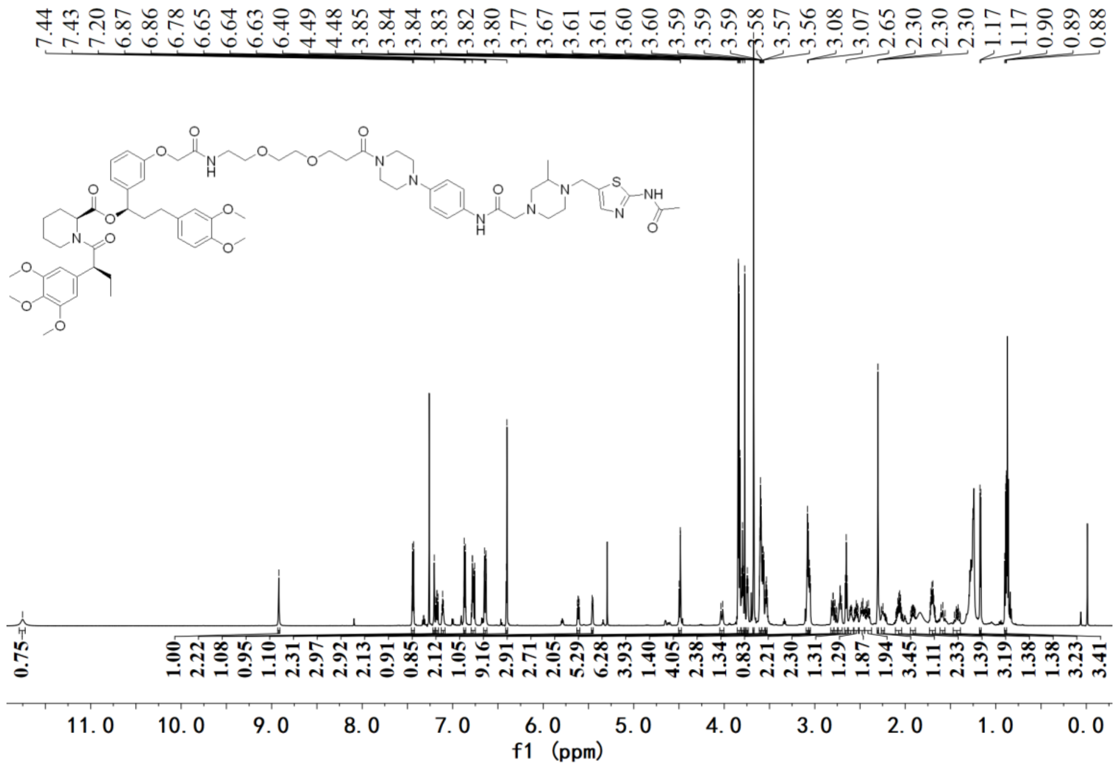

[1]H NMR spectrum of **A10**

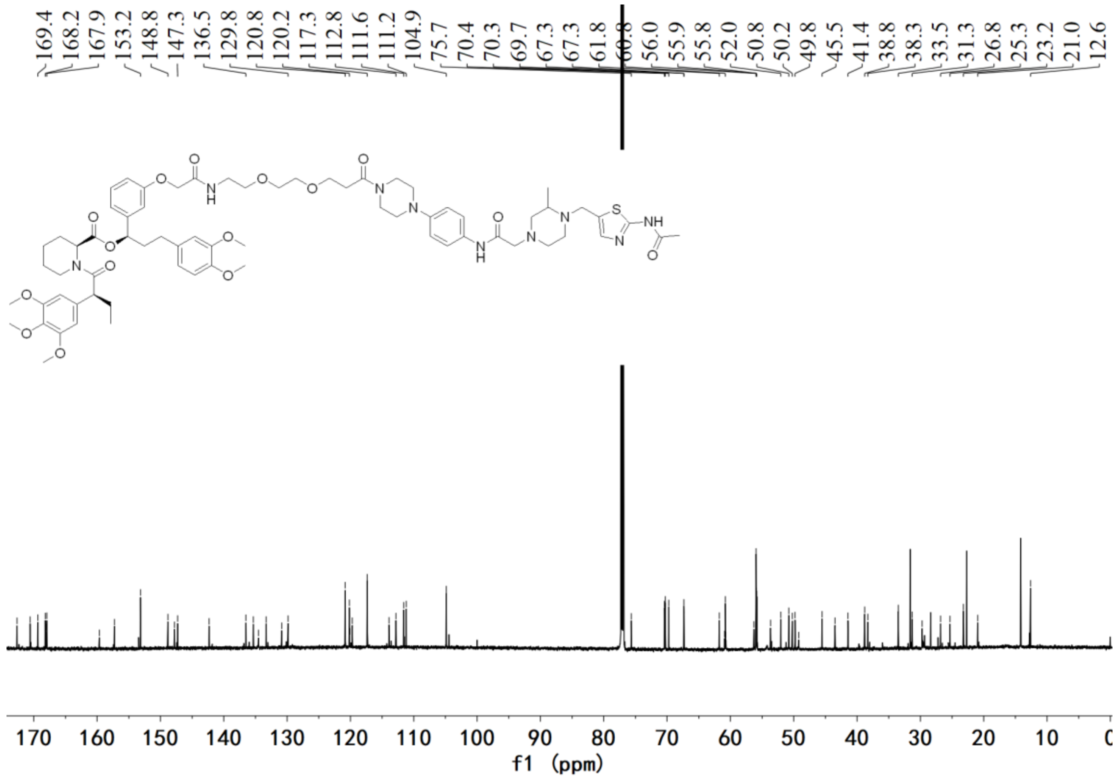

[13]C NMR spectrum of **A10**

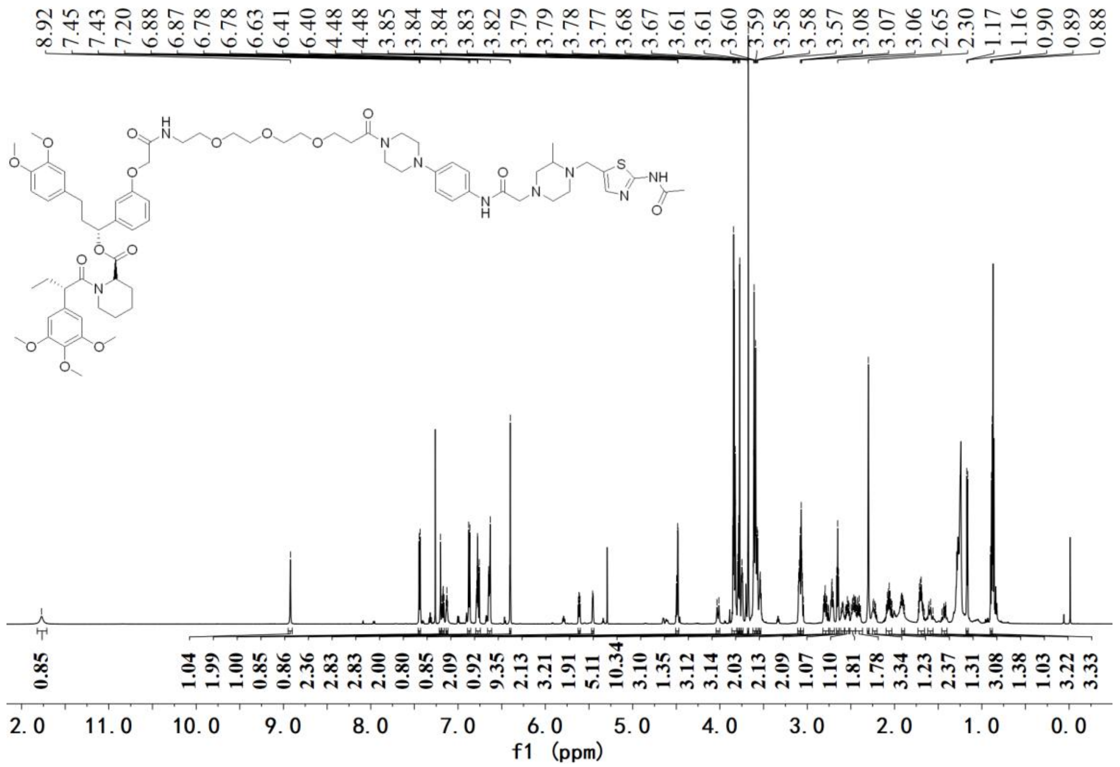


$^{1}H$ NMR spectrum of **A11**

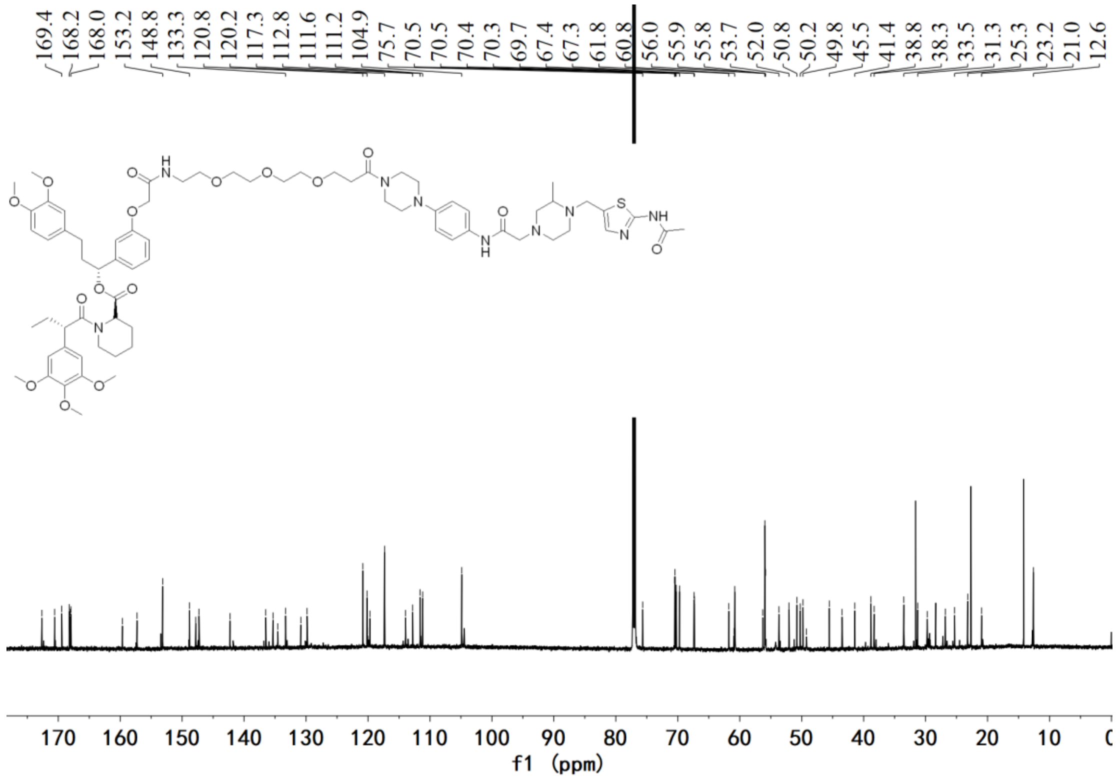


$^{13}C$ NMR spectrum of **A11**

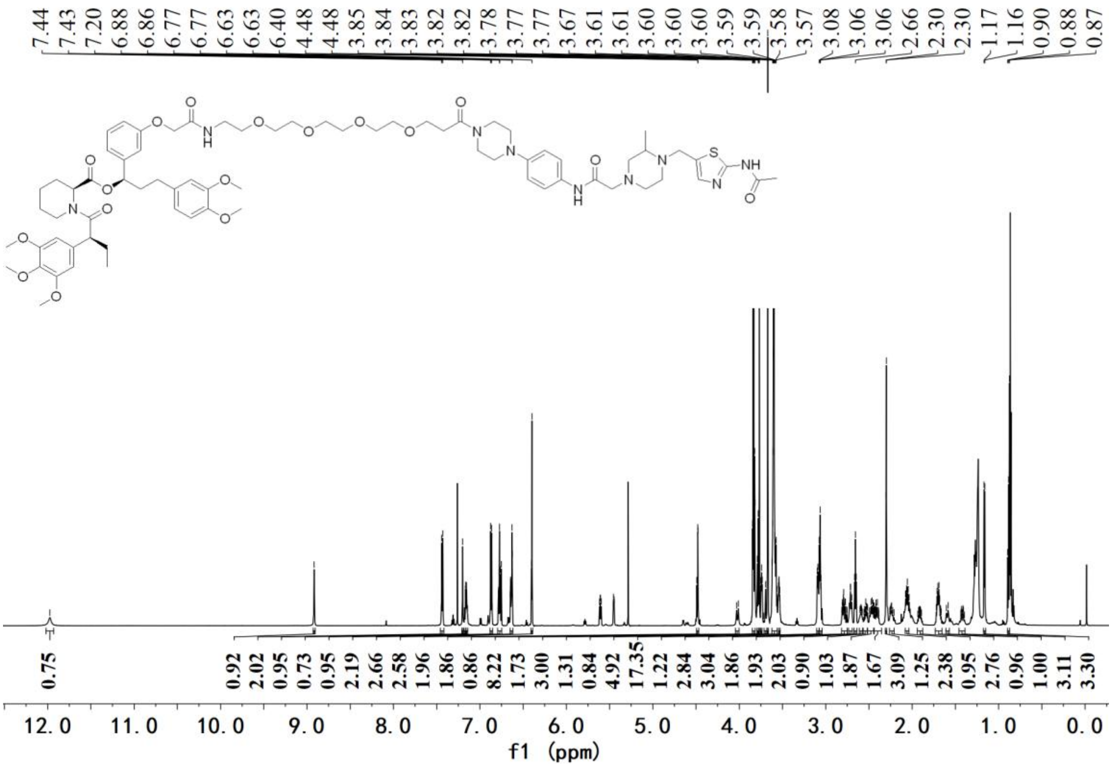


[1]H NMR spectrum of **A12**

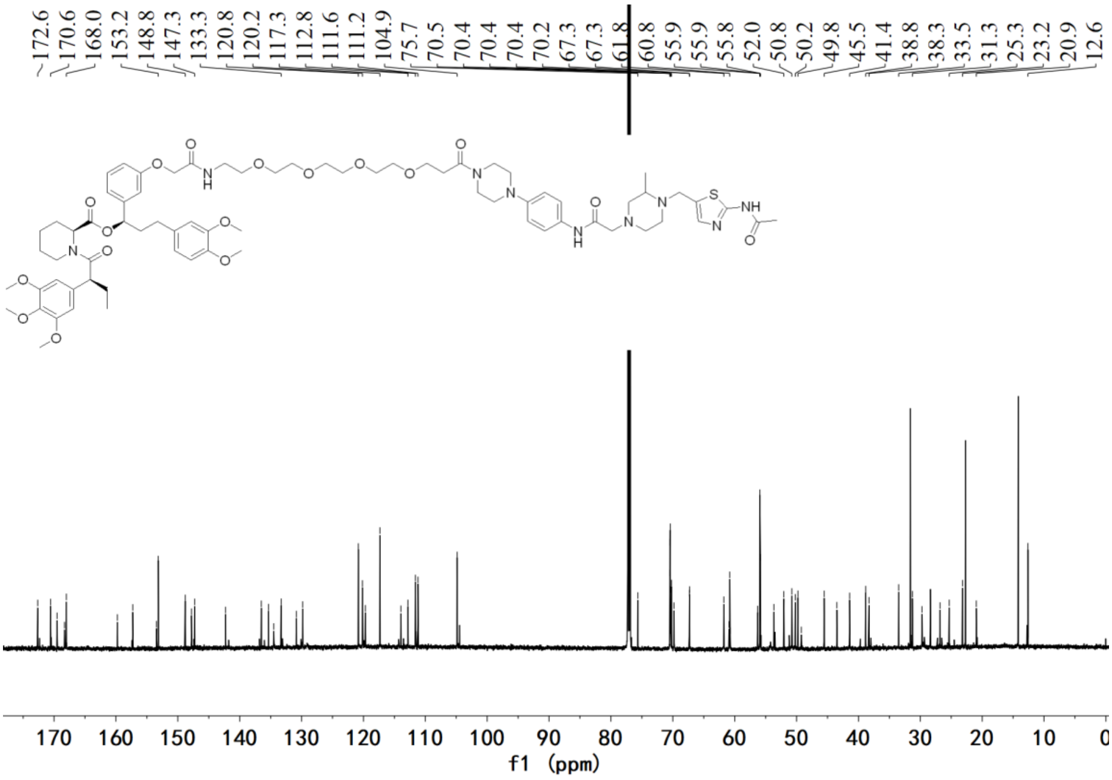


[13]C NMR spectrum of **A12**

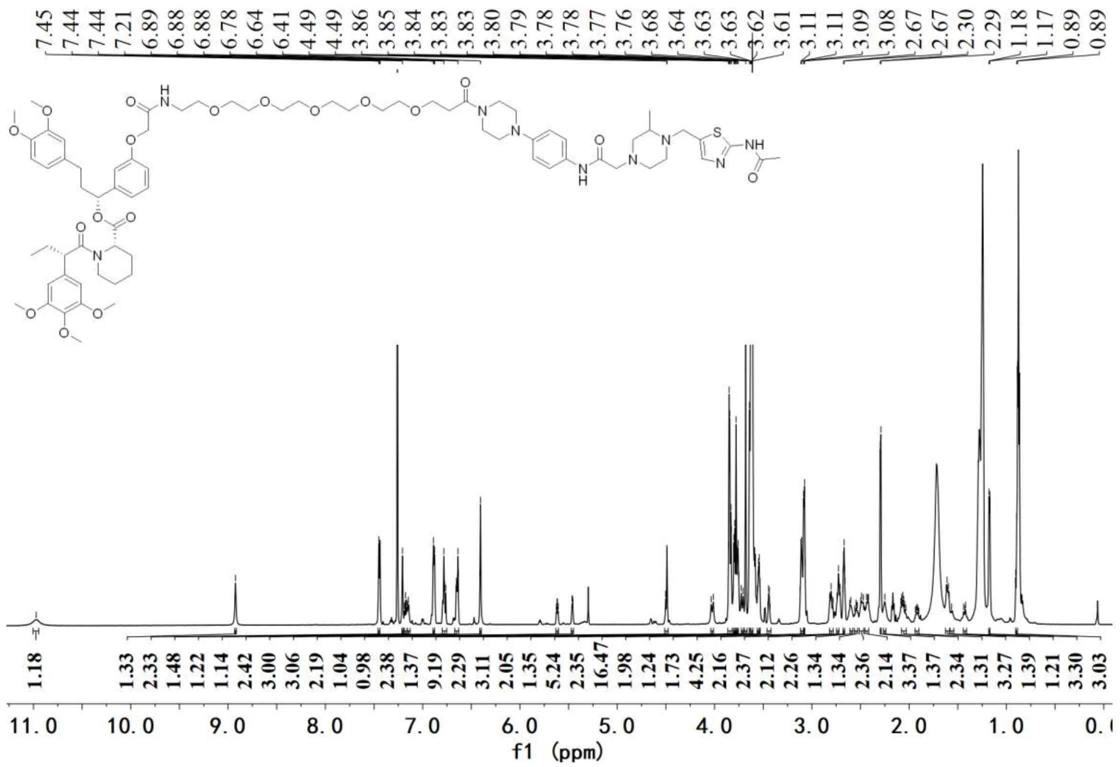

[1]H NMR spectrum of **A13**

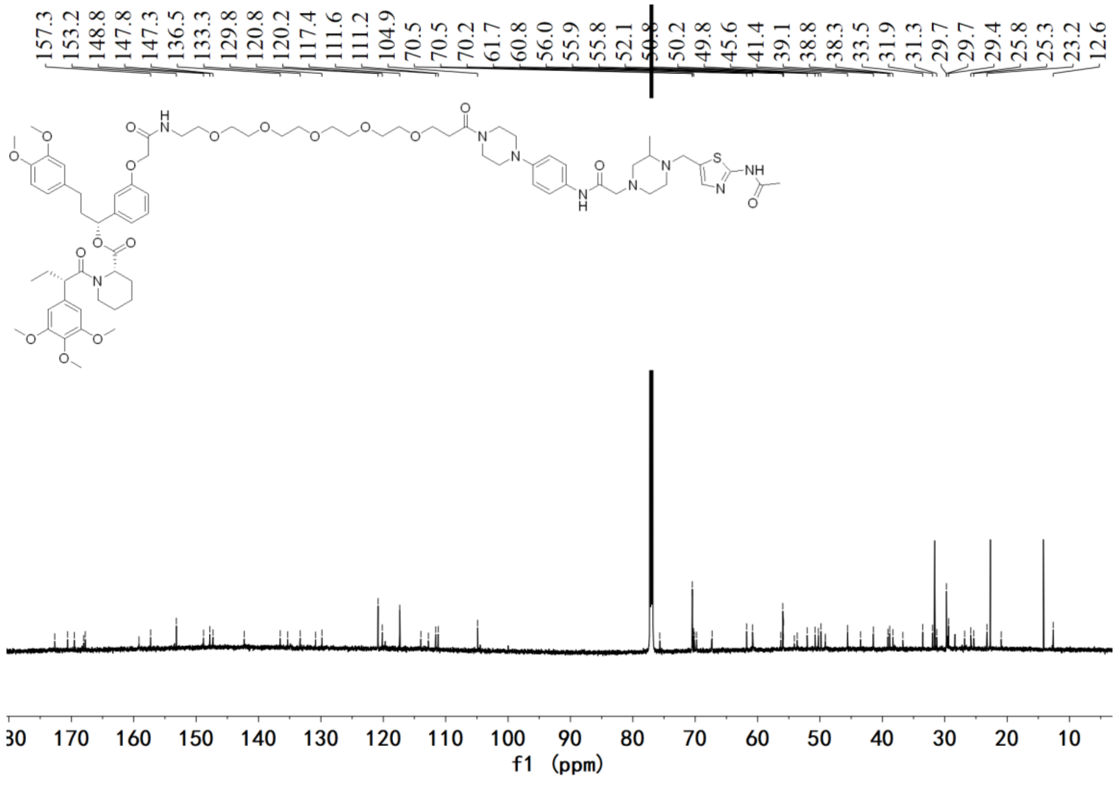

[13]C NMR spectrum of **A13**

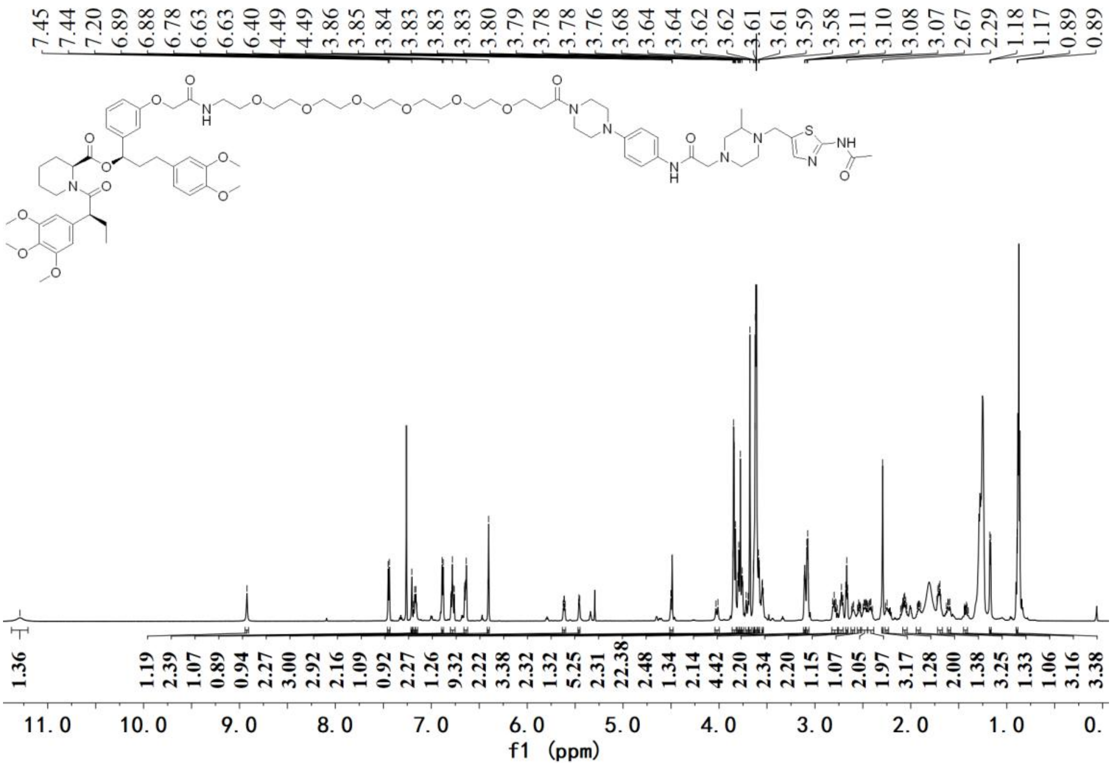


[1]H NMR spectrum of **A14**

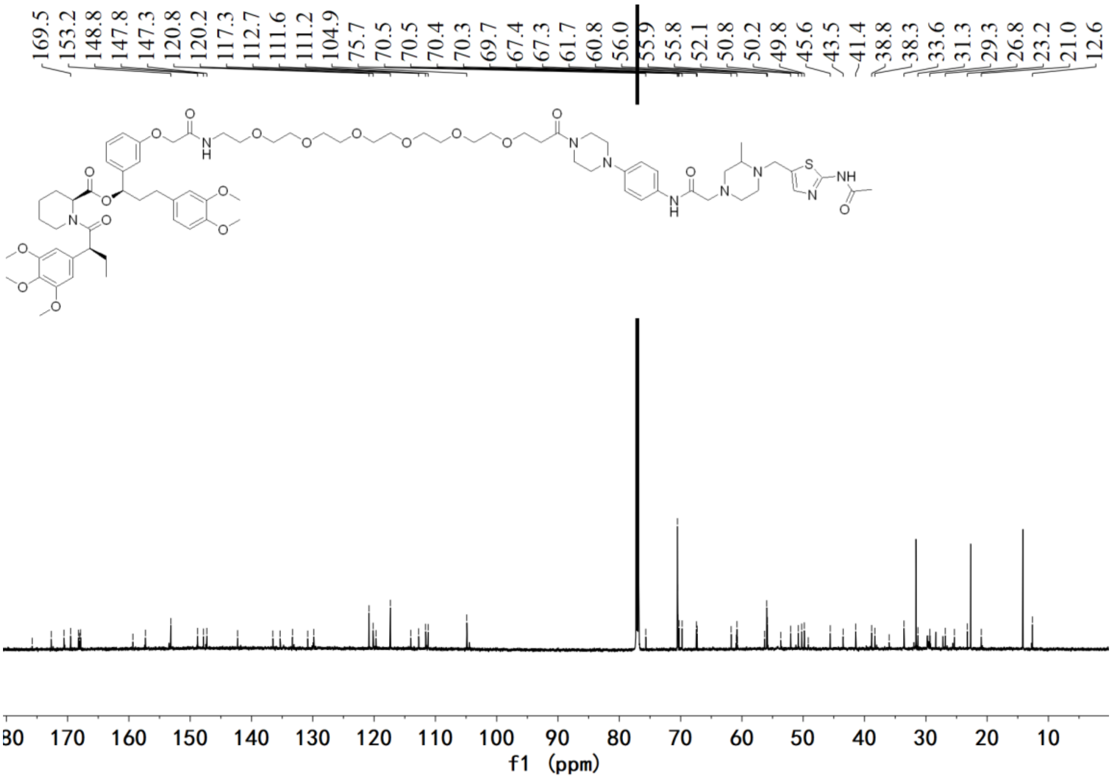


[13]C NMR spectrum of **A14**

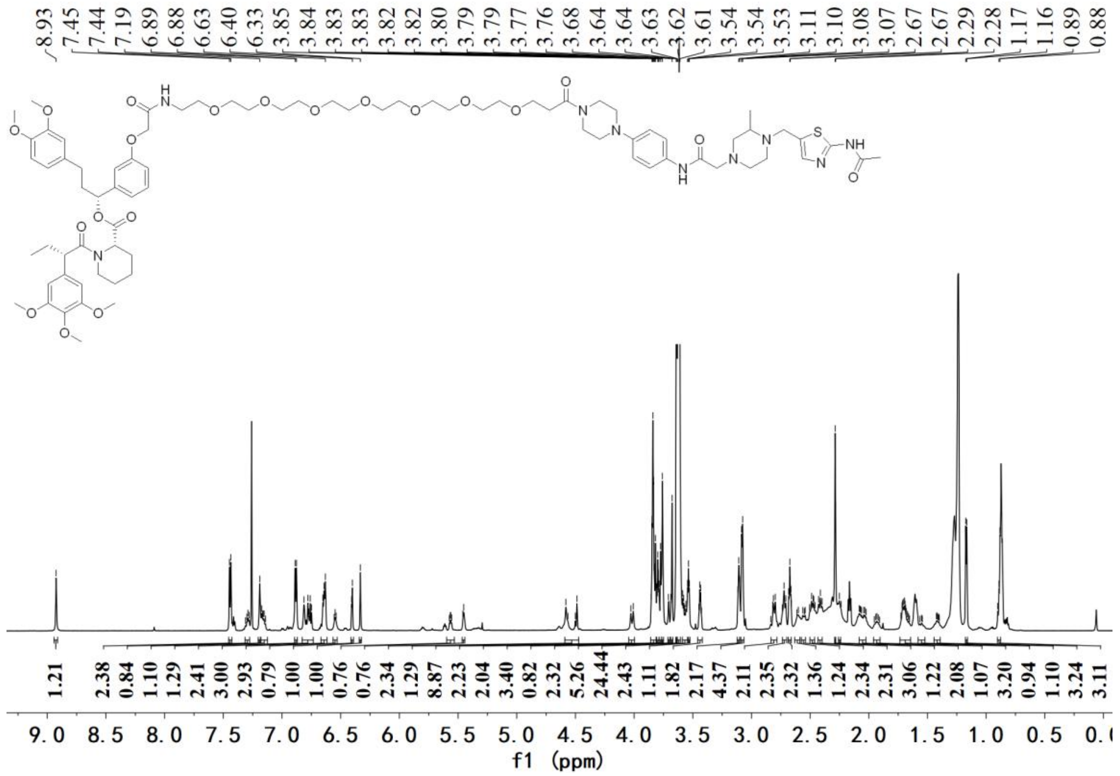

[1]H NMR spectrum of **A15**

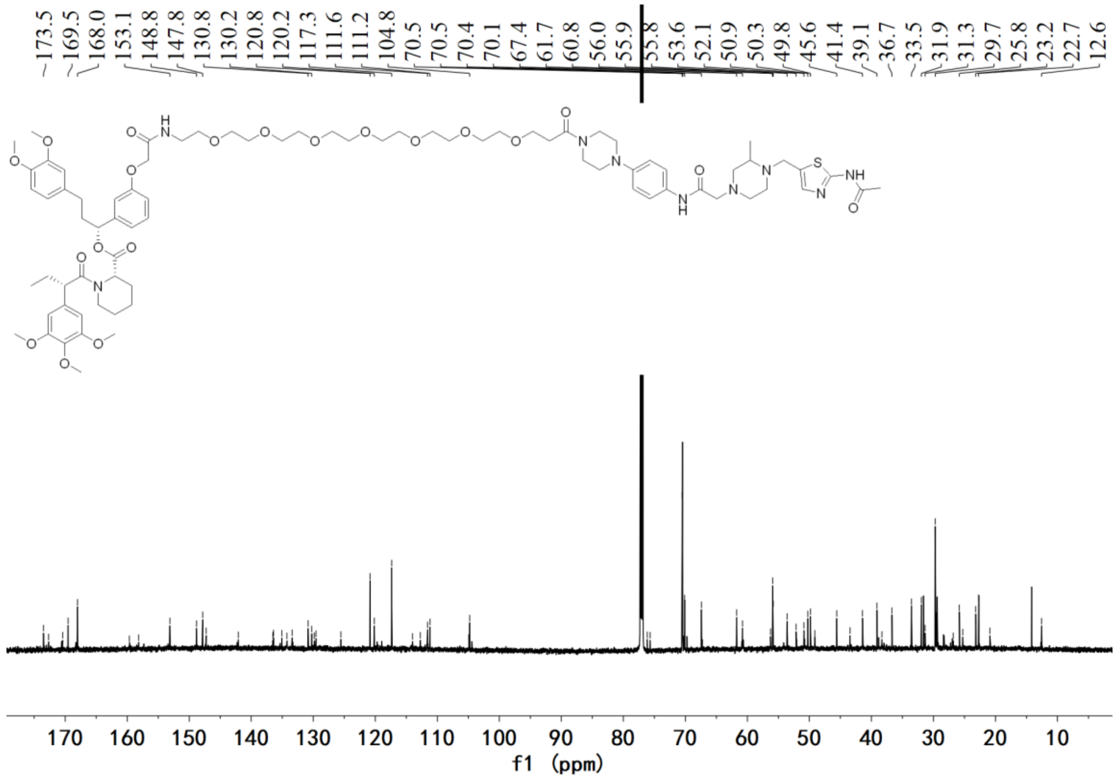

[13]C NMR spectrum of **A15**

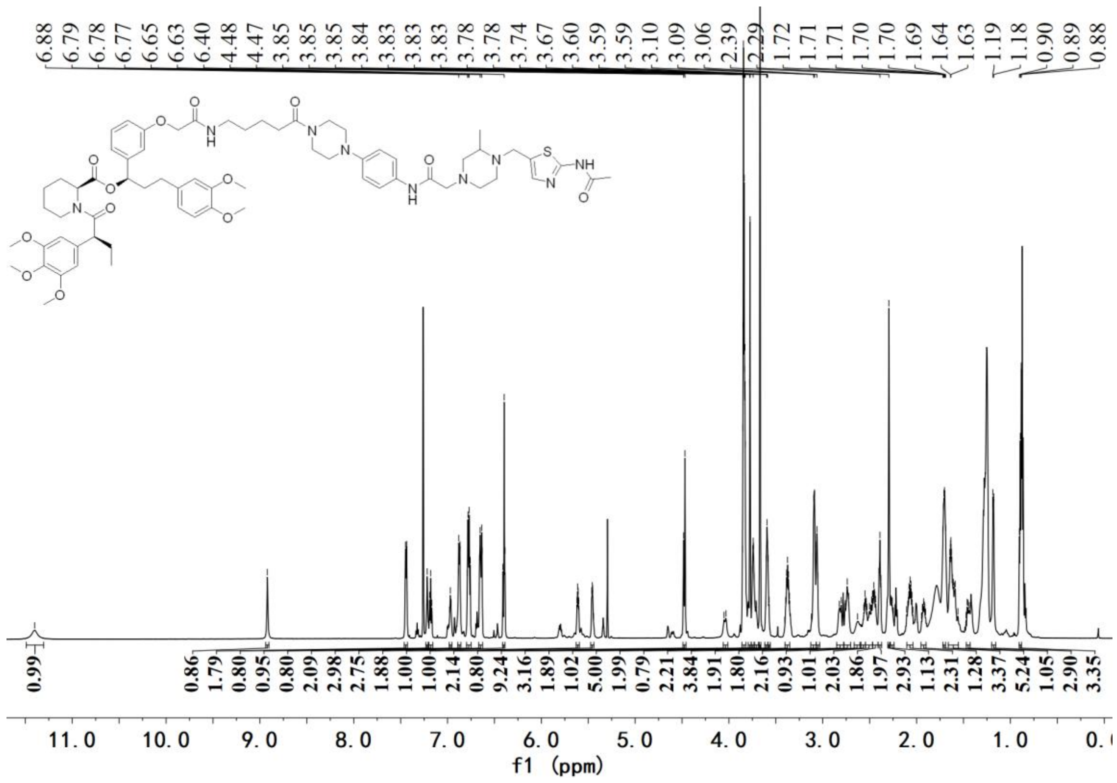


$^{1}H$ NMR spectrum of **A16**

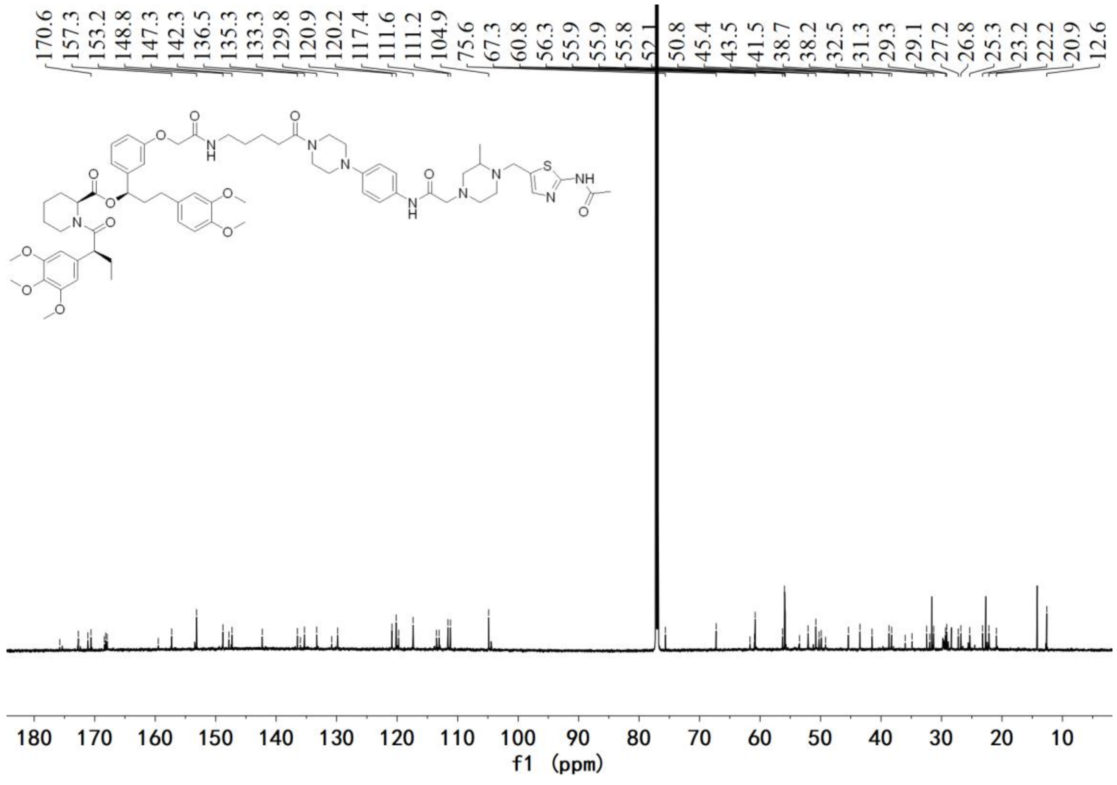


$^{13}C$ NMR spectrum of **A16**

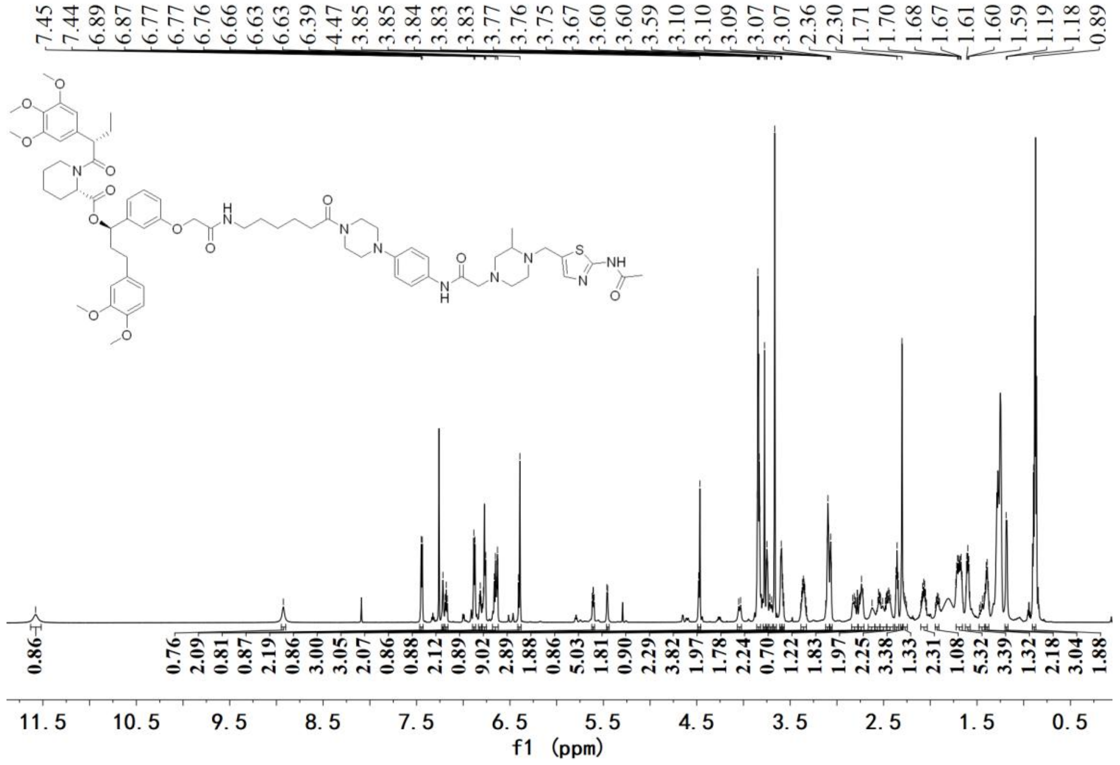


$^{1}H$ NMR spectrum of **A17**

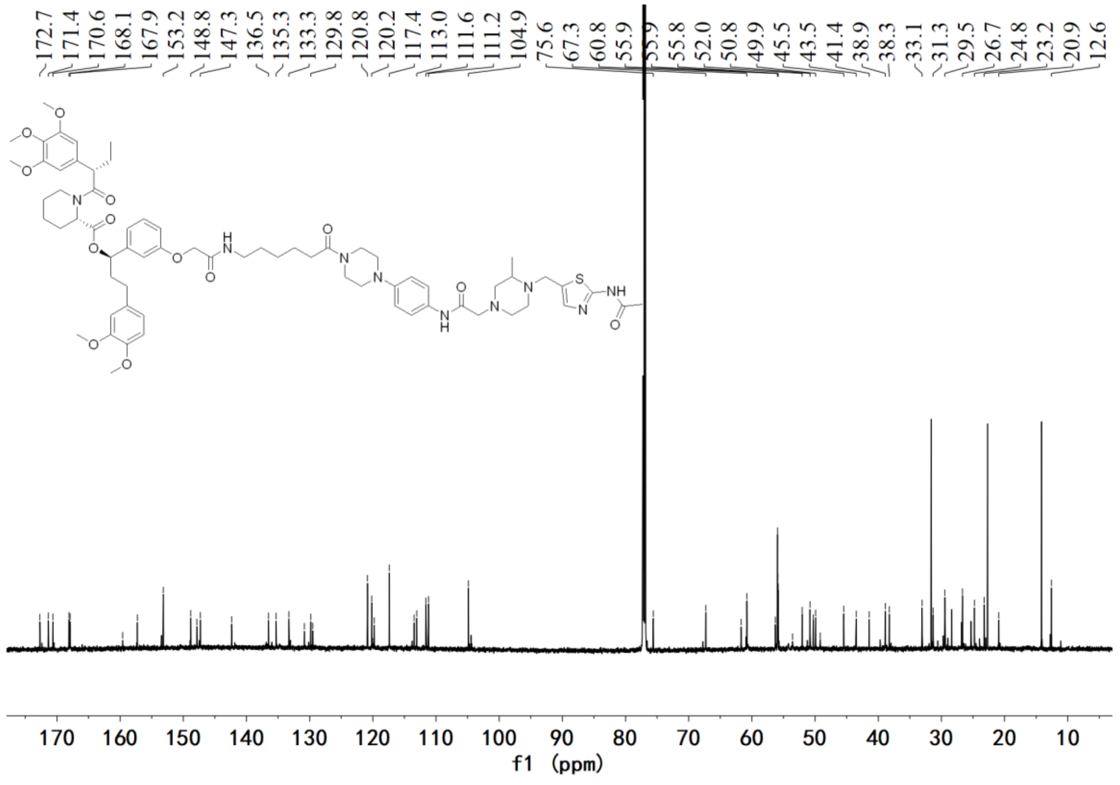


$^{13}C$ NMR spectrum of **A17**

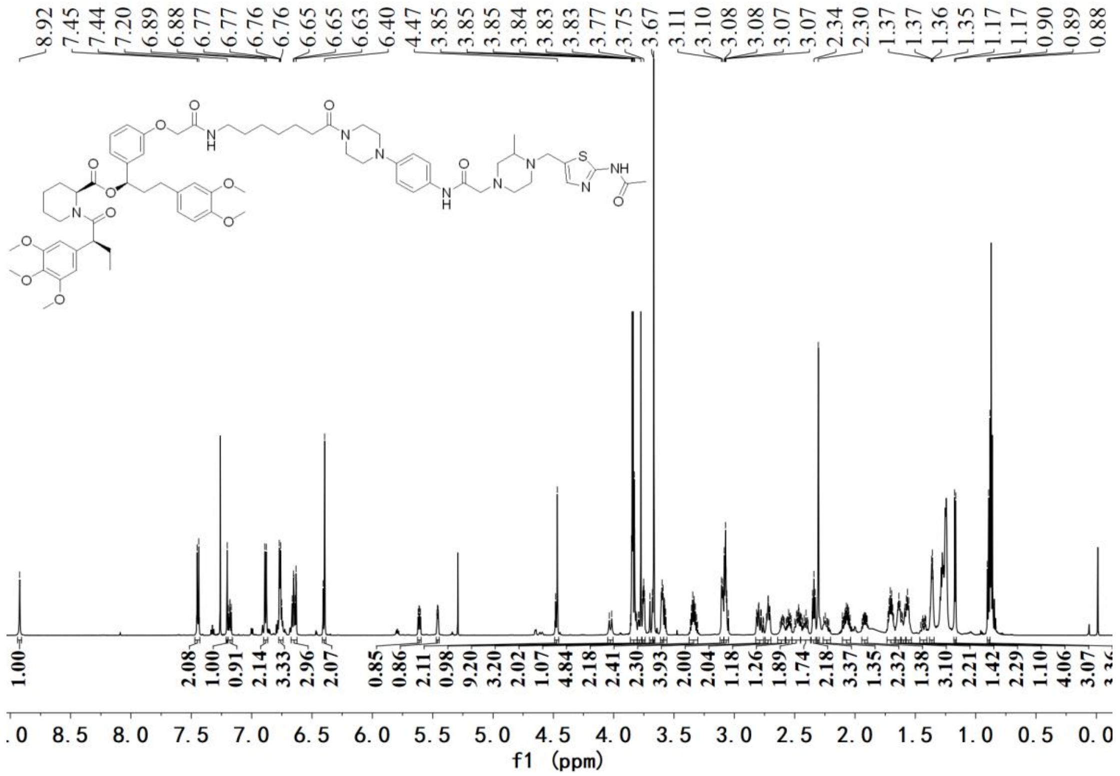

$^{1}H$ NMR spectrum of **A18**

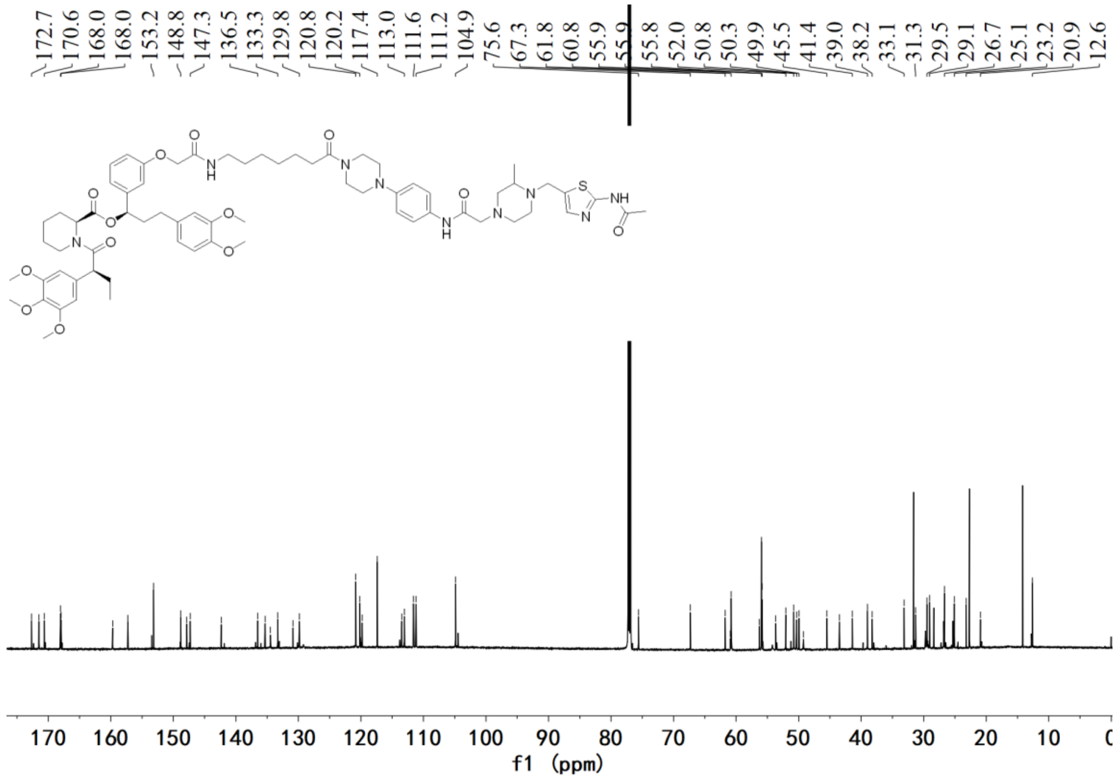

$^{13}C$ NMR spectrum of **A18**

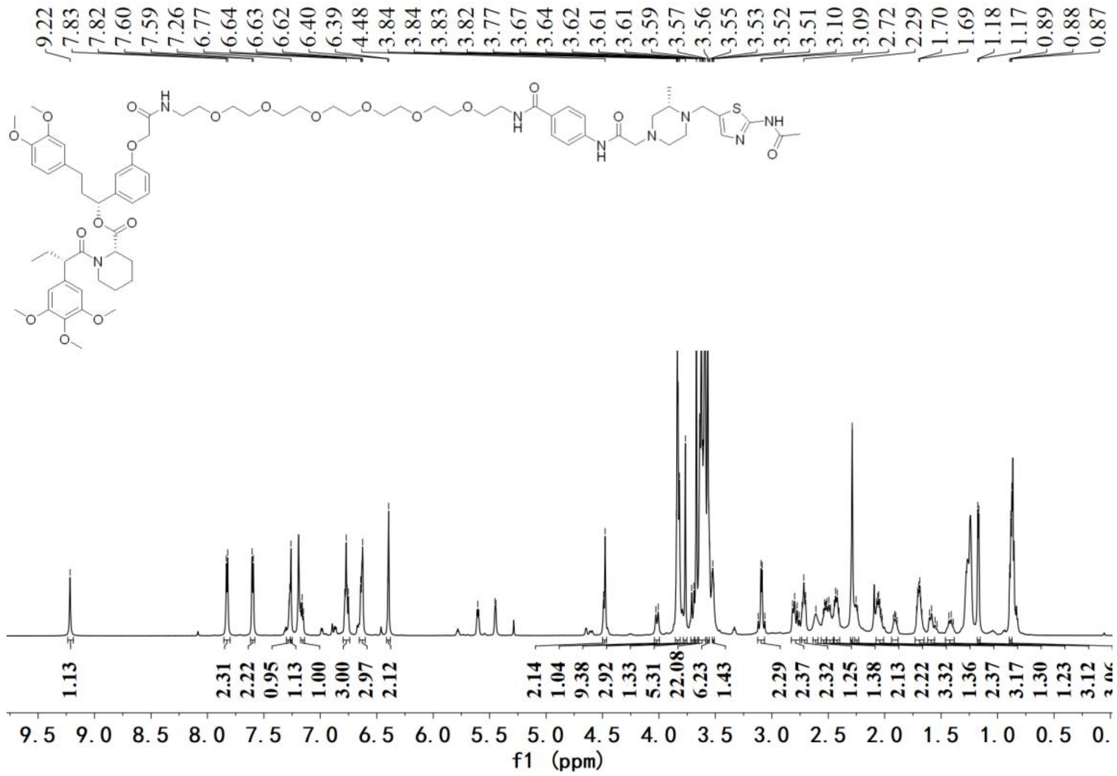

[1]H NMR spectrum of **B1**

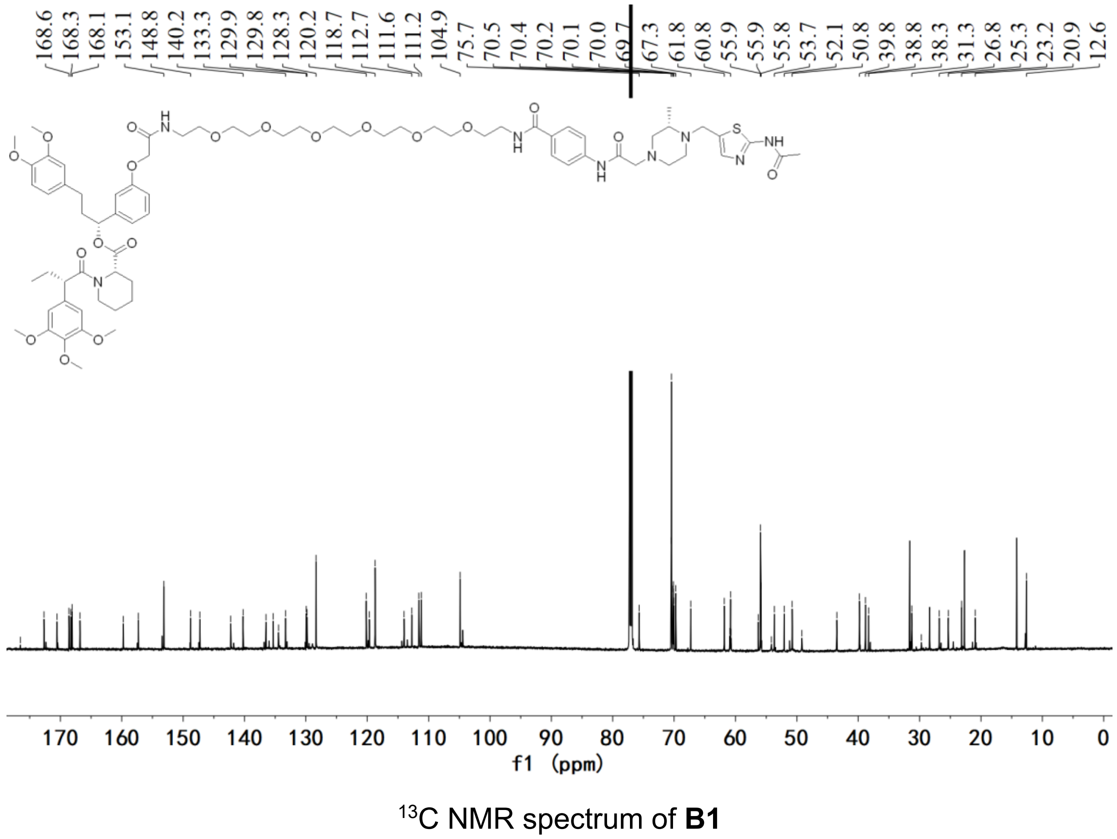

[13]C NMR spectrum of **B1**

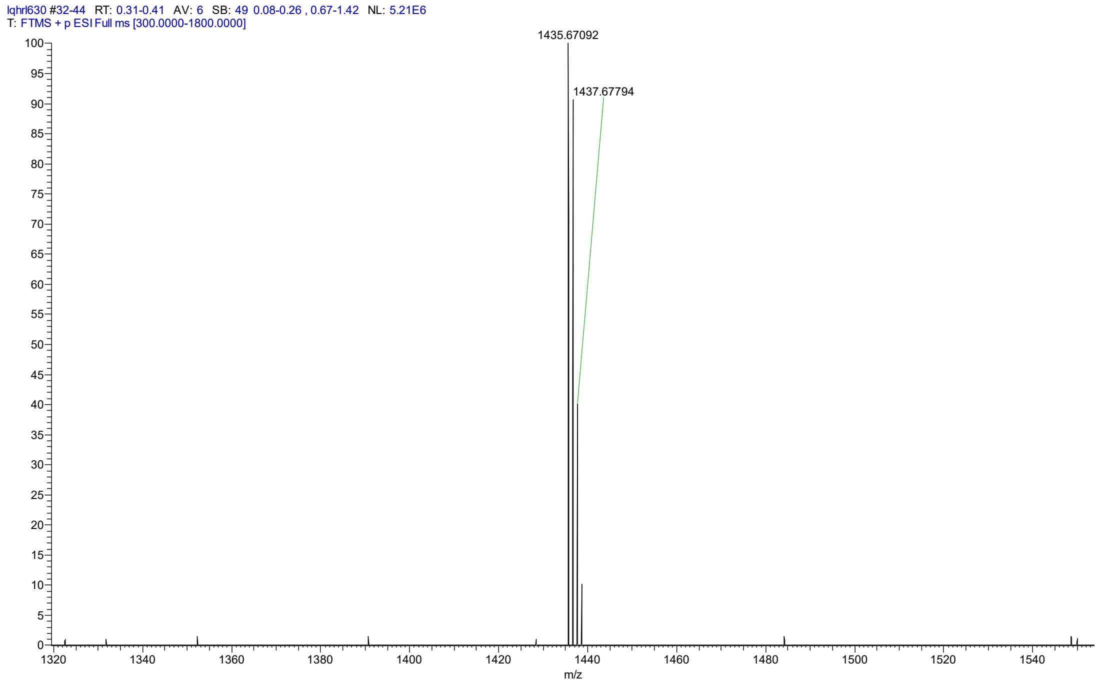


HRMS spectrum of **B1**

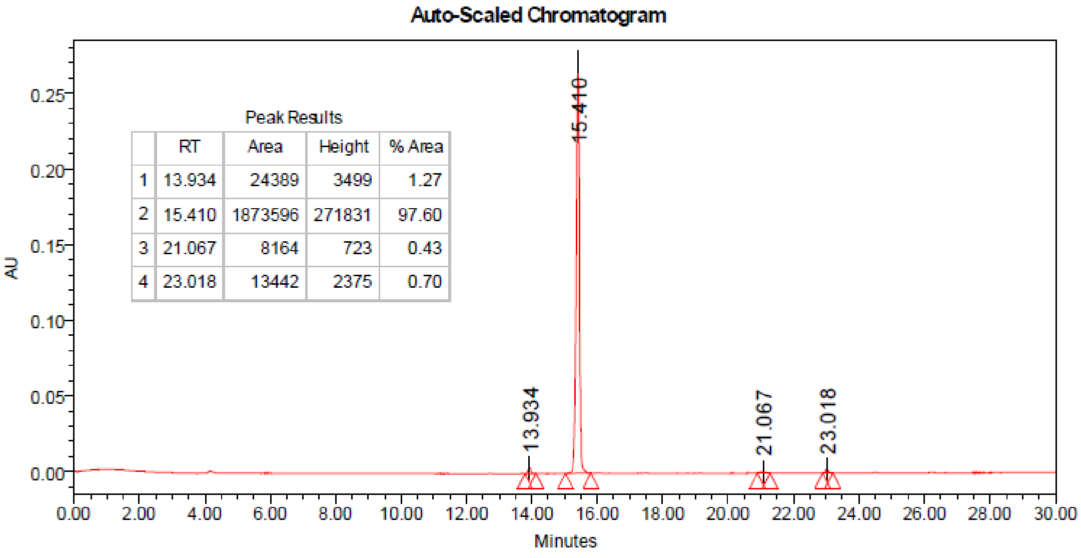


Peak Results

| | RT | Area | Height | % Area |
|---|---|---|---|---|
| 1 | 13.934 | 24389 | 3499 | 1.27 |
| 2 | 15.410 | 1873596 | 271831 | 97.60 |
| 3 | 21.067 | 8164 | 723 | 0.43 |
| 4 | 23.018 | 13442 | 2375 | 0.70 |

HPLC spectrum of **B1**

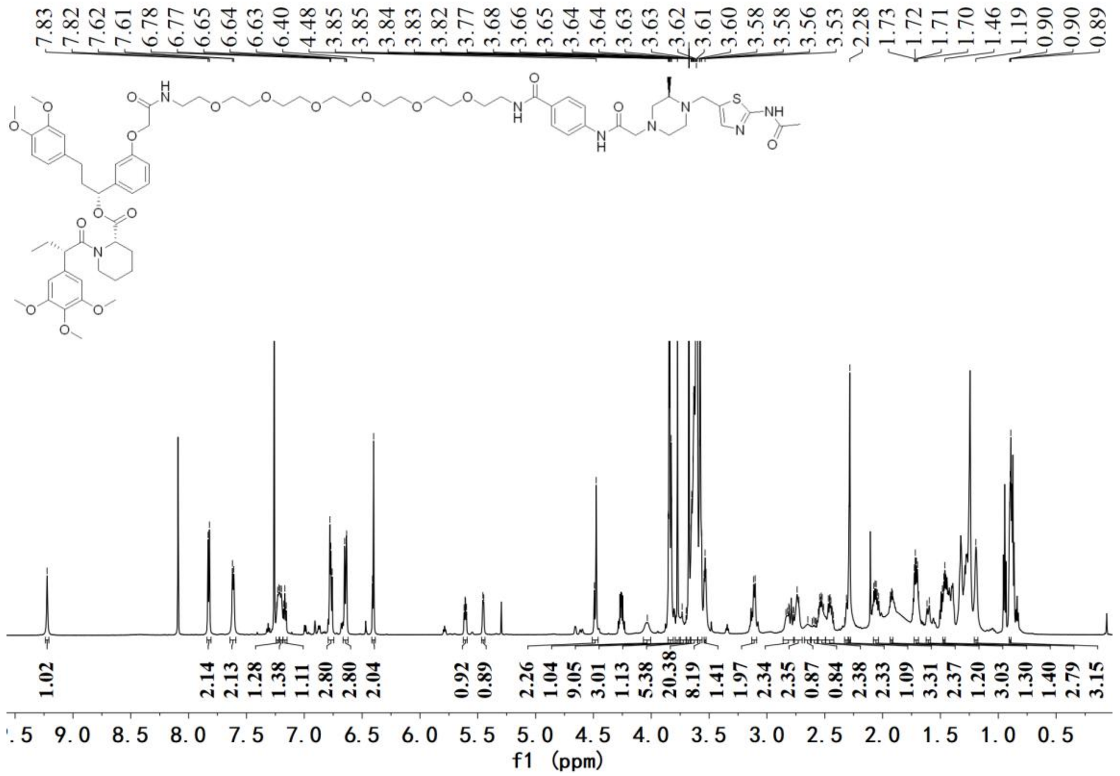


[1]H NMR spectrum of **B2**

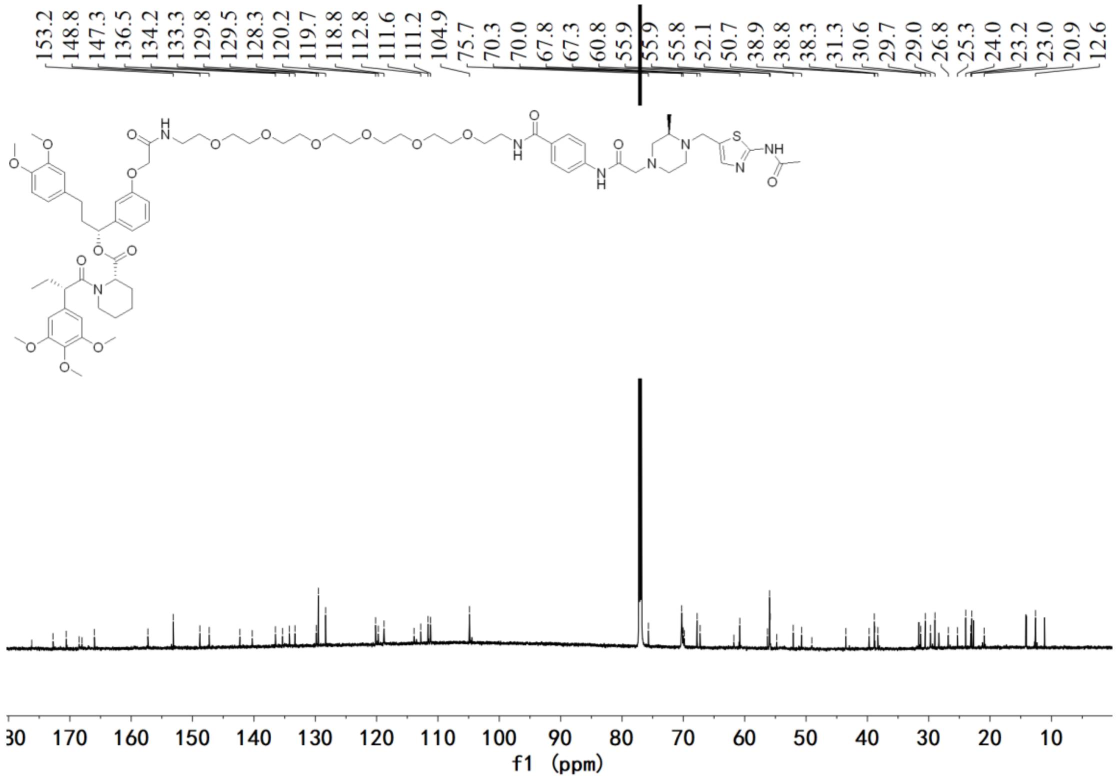


[13]C NMR spectrum of **B2**

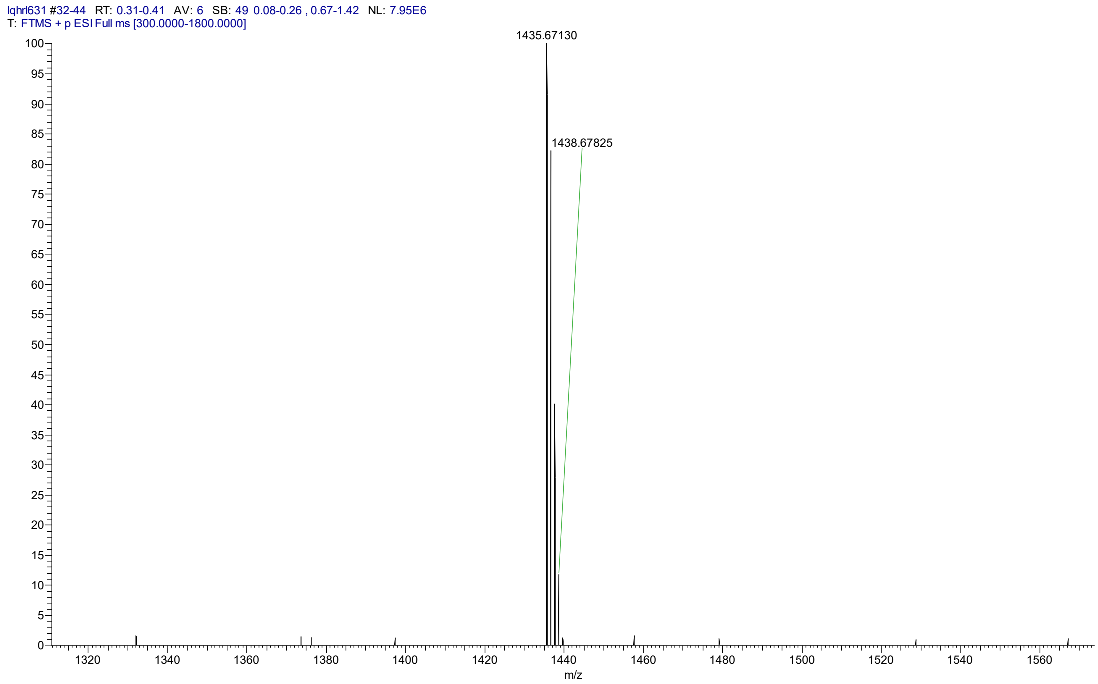


HRMS spectrum of **B2**

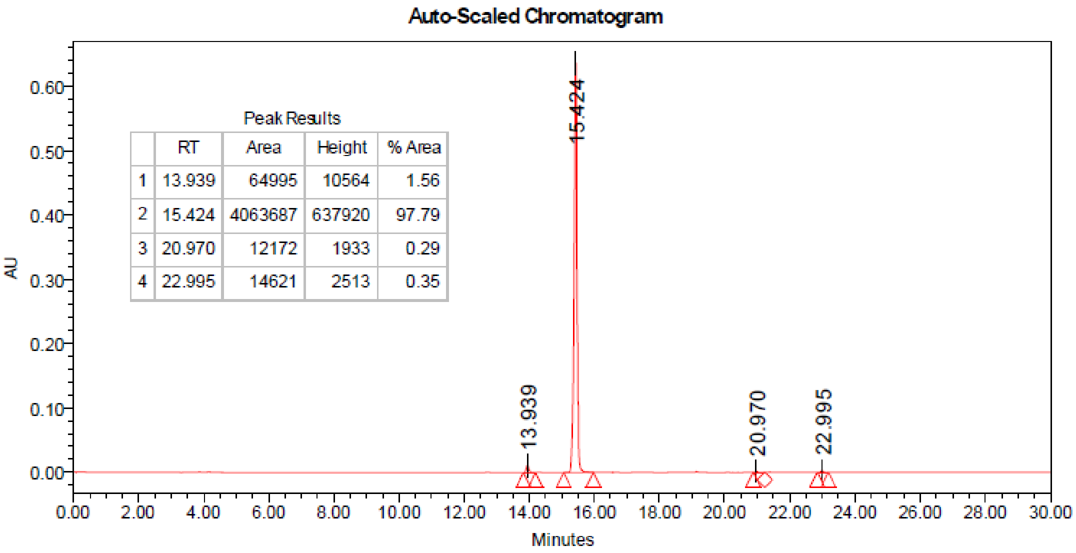


| | RT | Area | Height | % Area |
|---|---|---|---|---|
| 1 | 13.939 | 64995 | 10564 | 1.56 |
| 2 | 15.424 | 4063687 | 637920 | 97.79 |
| 3 | 20.970 | 12172 | 1933 | 0.29 |
| 4 | 22.995 | 14621 | 2513 | 0.35 |

HPLC spectrum of **B2**

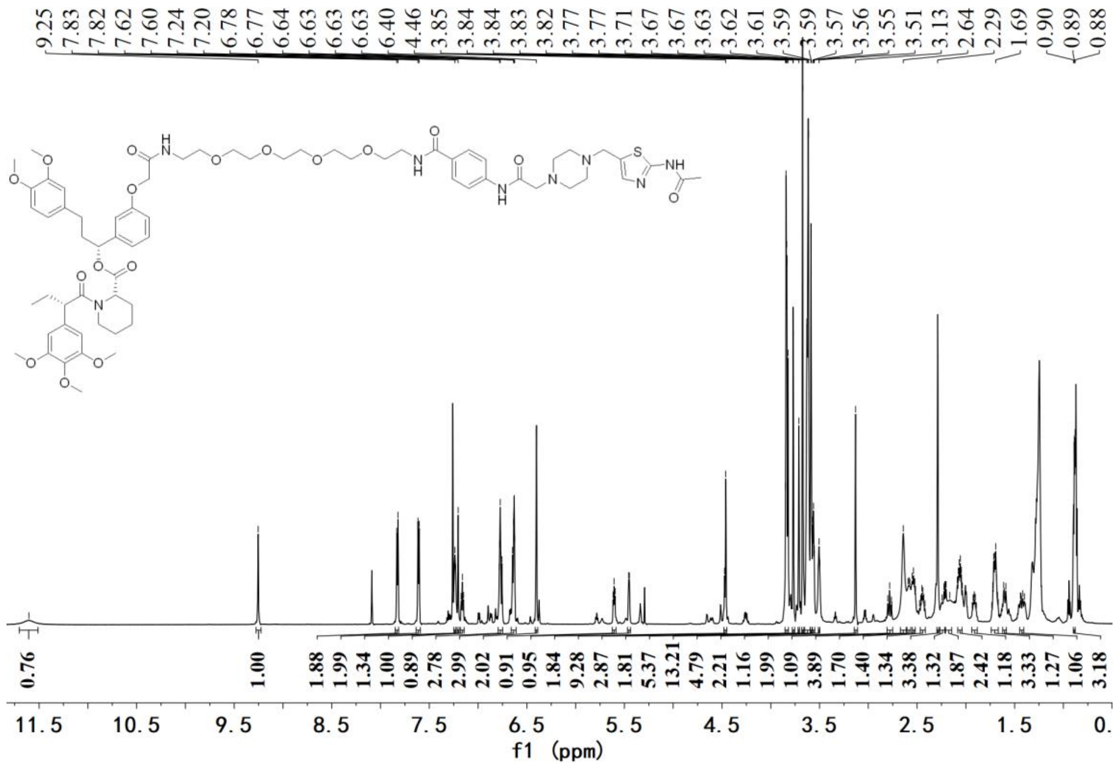


[1]H NMR spectrum of **C1**

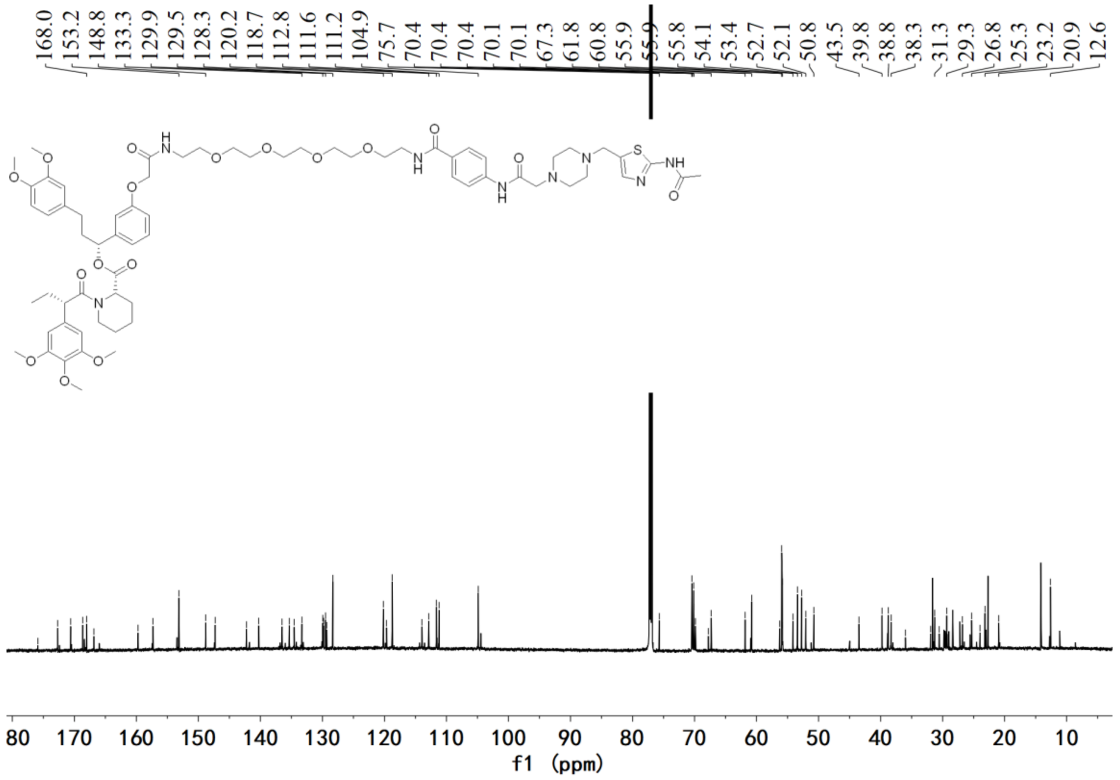


[13]C NMR spectrum of **C1**

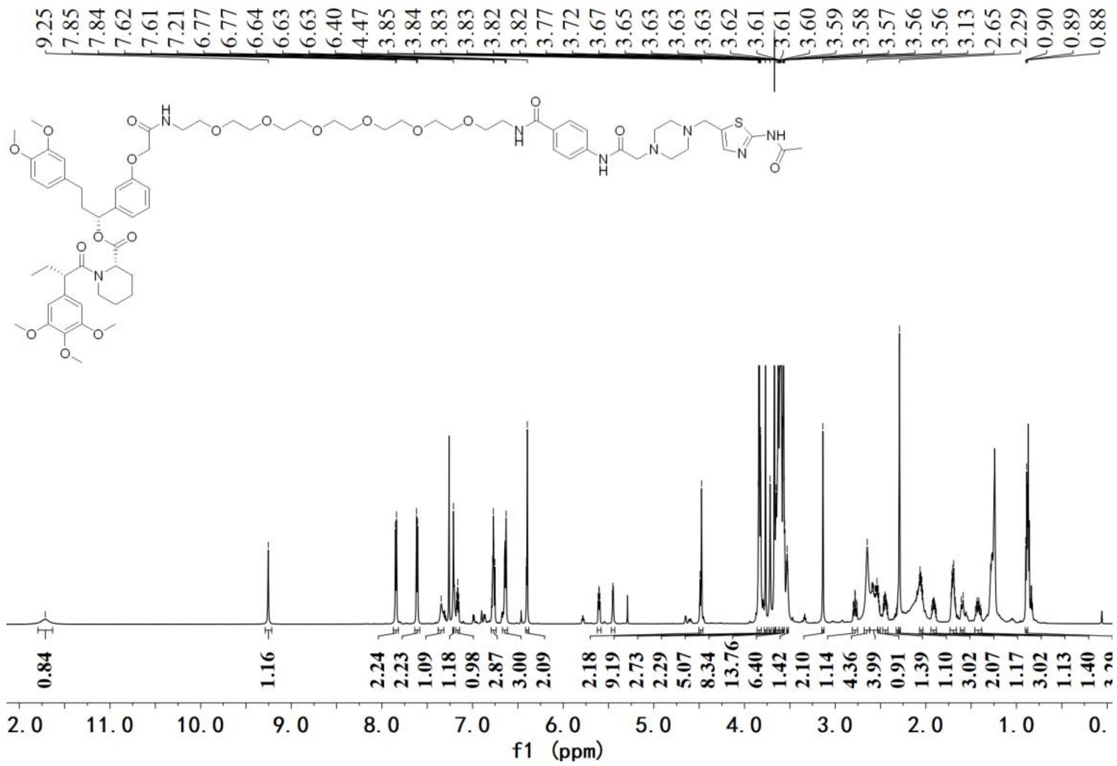


[1]H NMR spectrum of **C2**

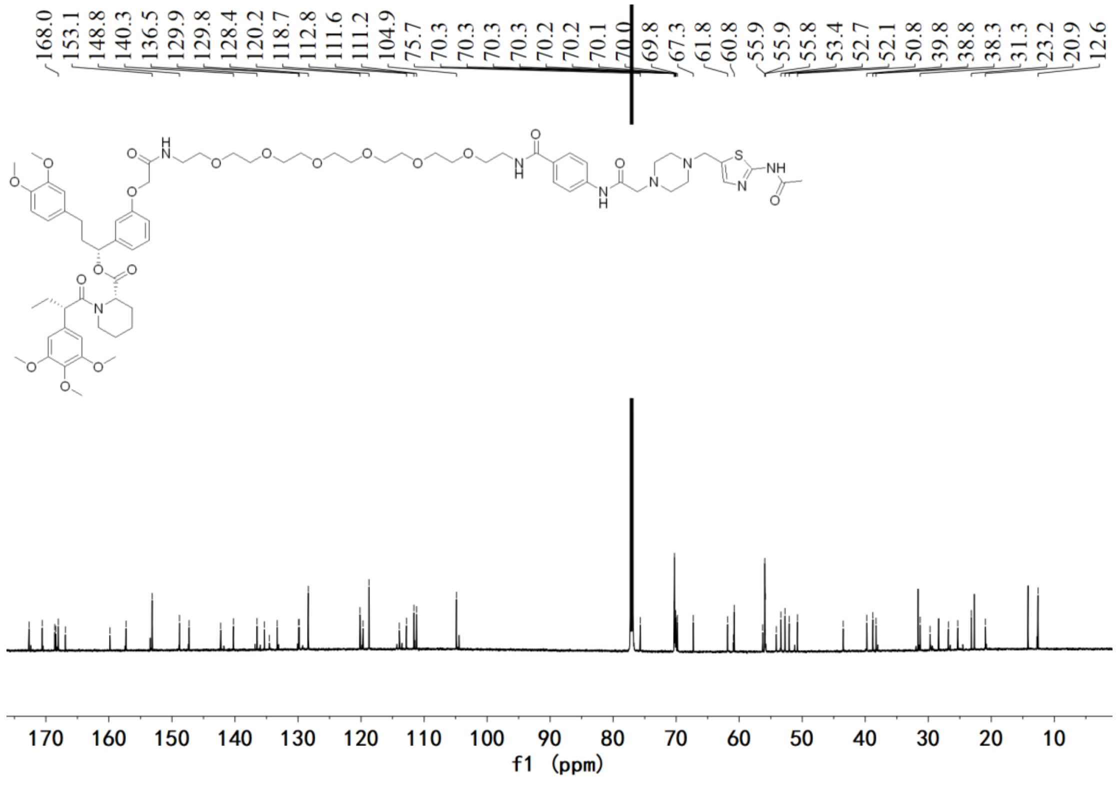


[13]C NMR spectrum of **C2**

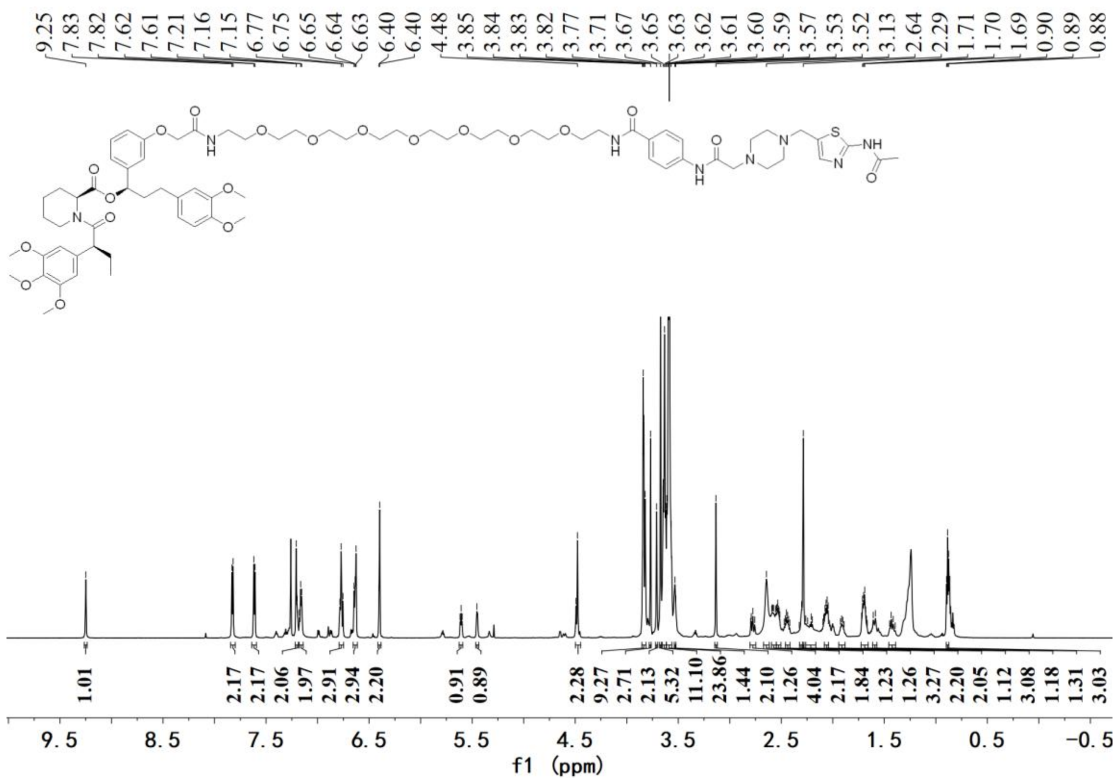

[1]H NMR spectrum of **C3**

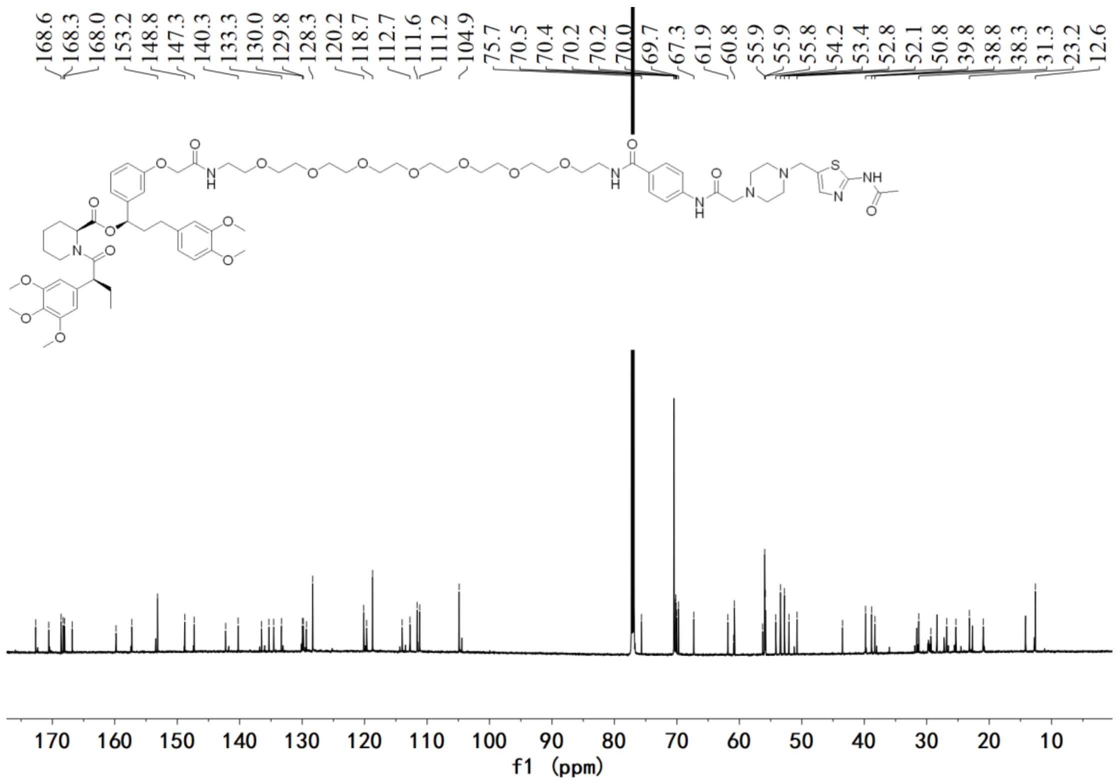

[13]C NMR spectrum of **C3**

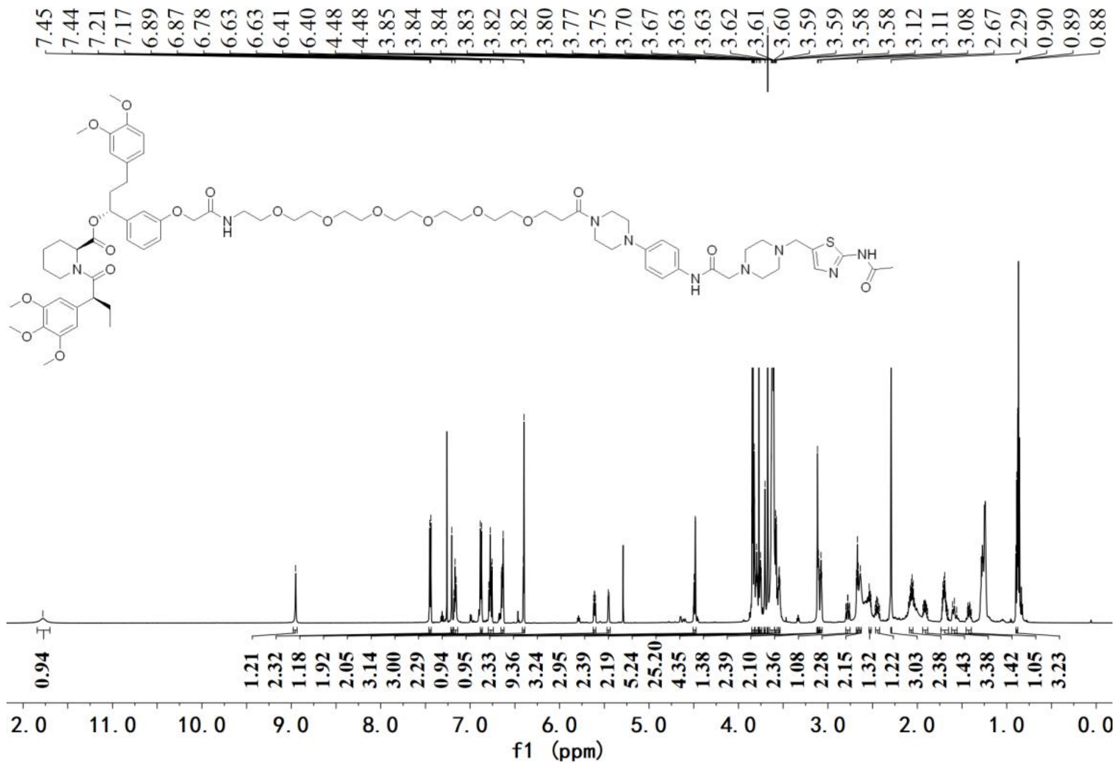

[1]H NMR spectrum of **C4**

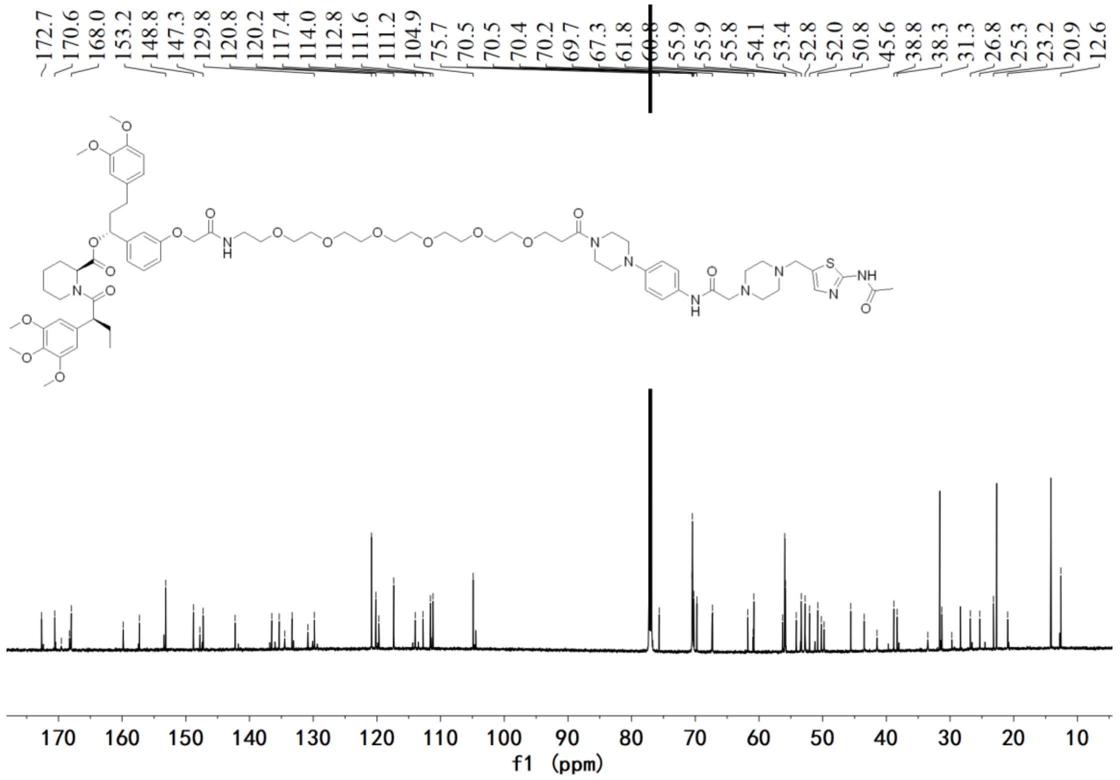

[13]C NMR spectrum of **C4**

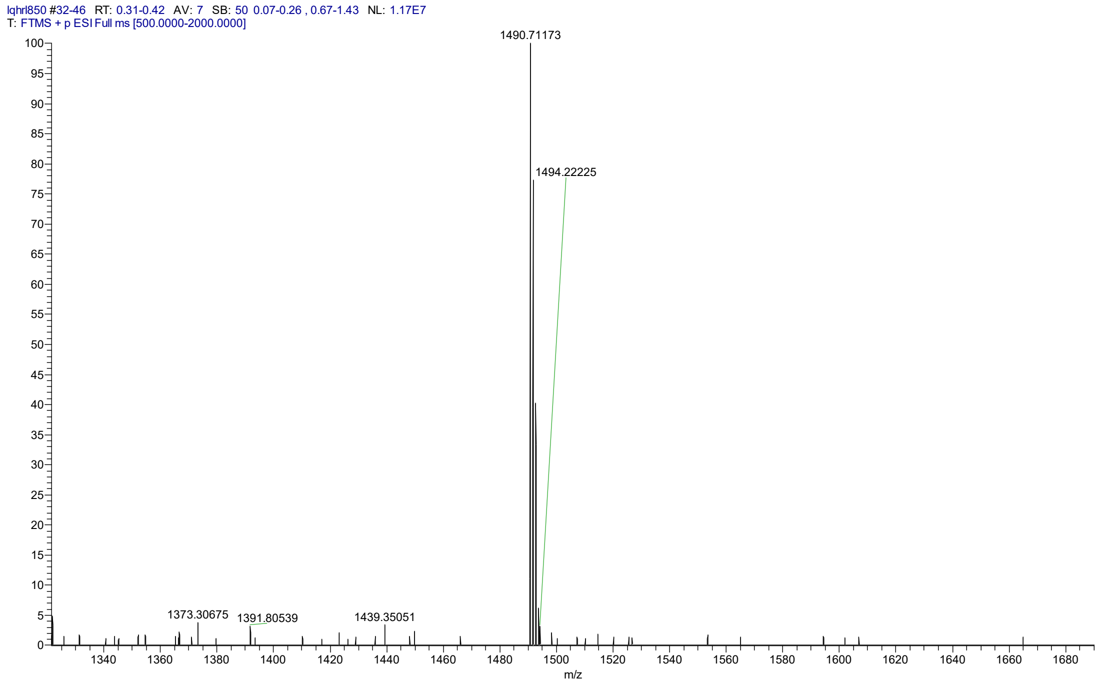


HRMS spectrum of **C4**

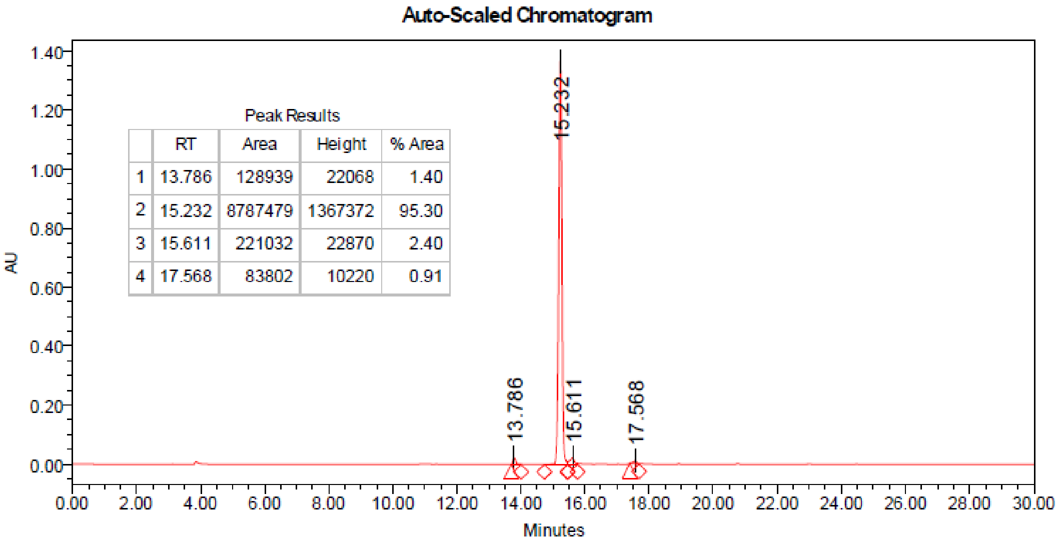


| | RT | Area | Height | % Area |
|---|---|---|---|---|
| 1 | 13.786 | 128939 | 22068 | 1.40 |
| 2 | 15.232 | 8787479 | 1367372 | 95.30 |
| 3 | 15.611 | 221032 | 22870 | 2.40 |
| 4 | 17.568 | 83802 | 10220 | 0.91 |

HPLC spectrum of **C4**

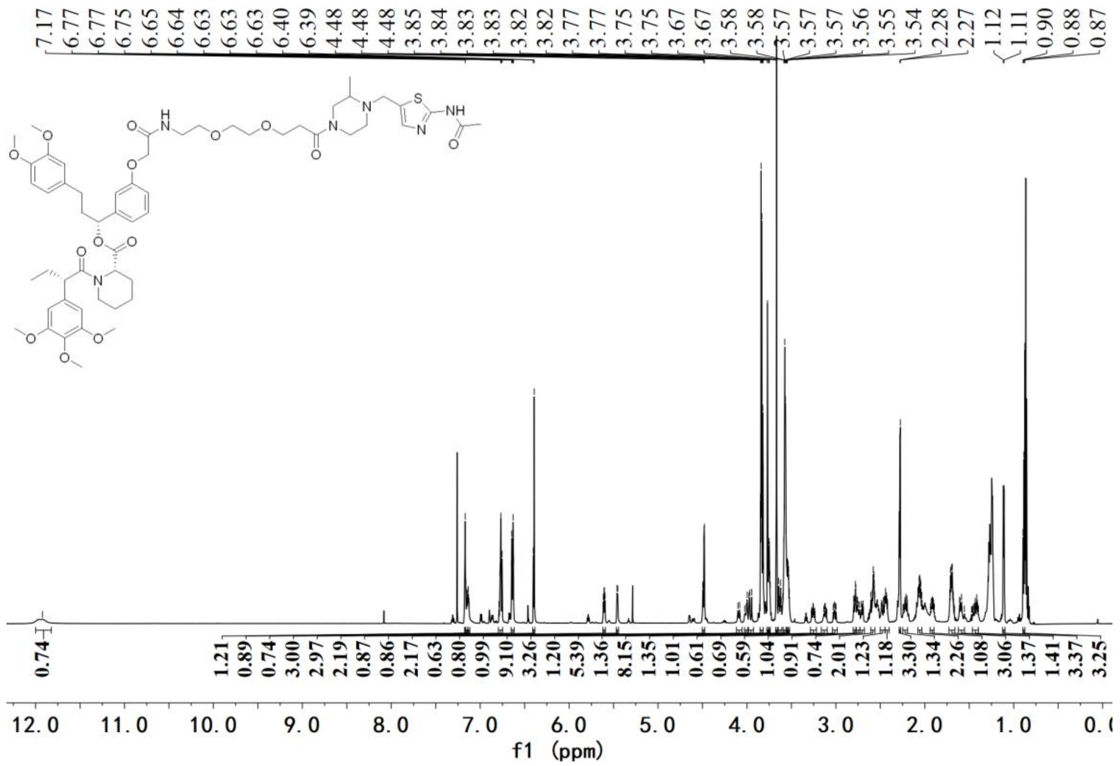


[1]H NMR spectrum of **D1**

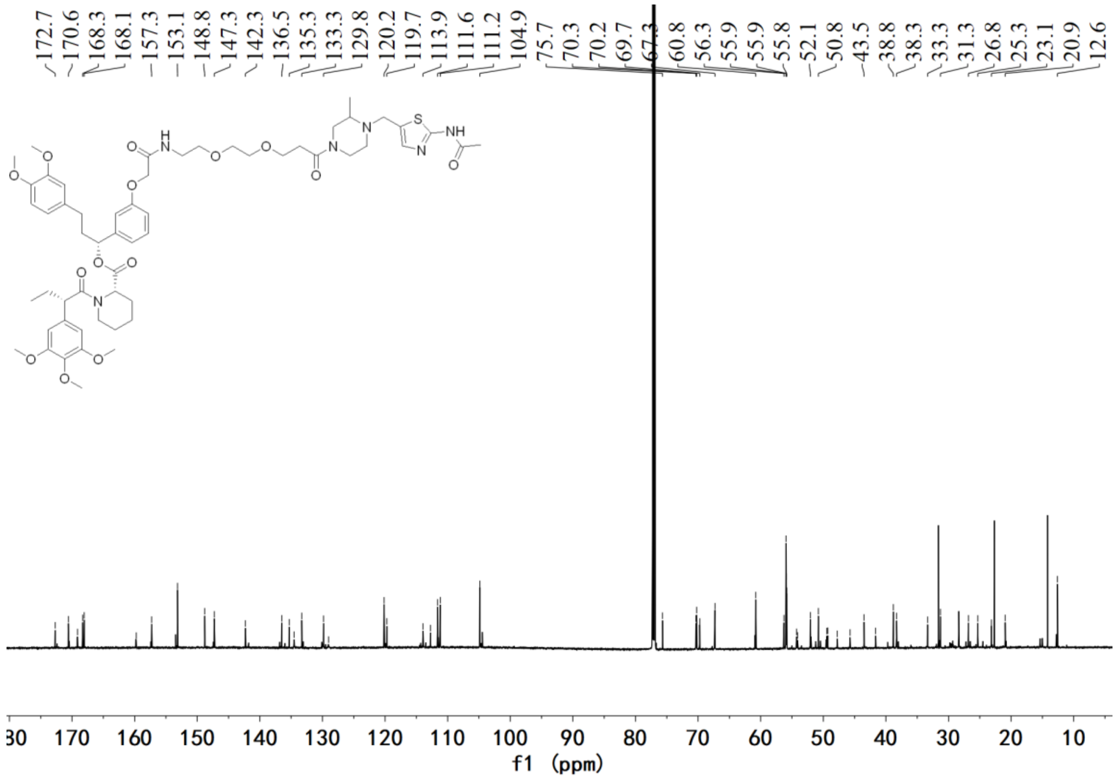


[13]C NMR spectrum of **D1**

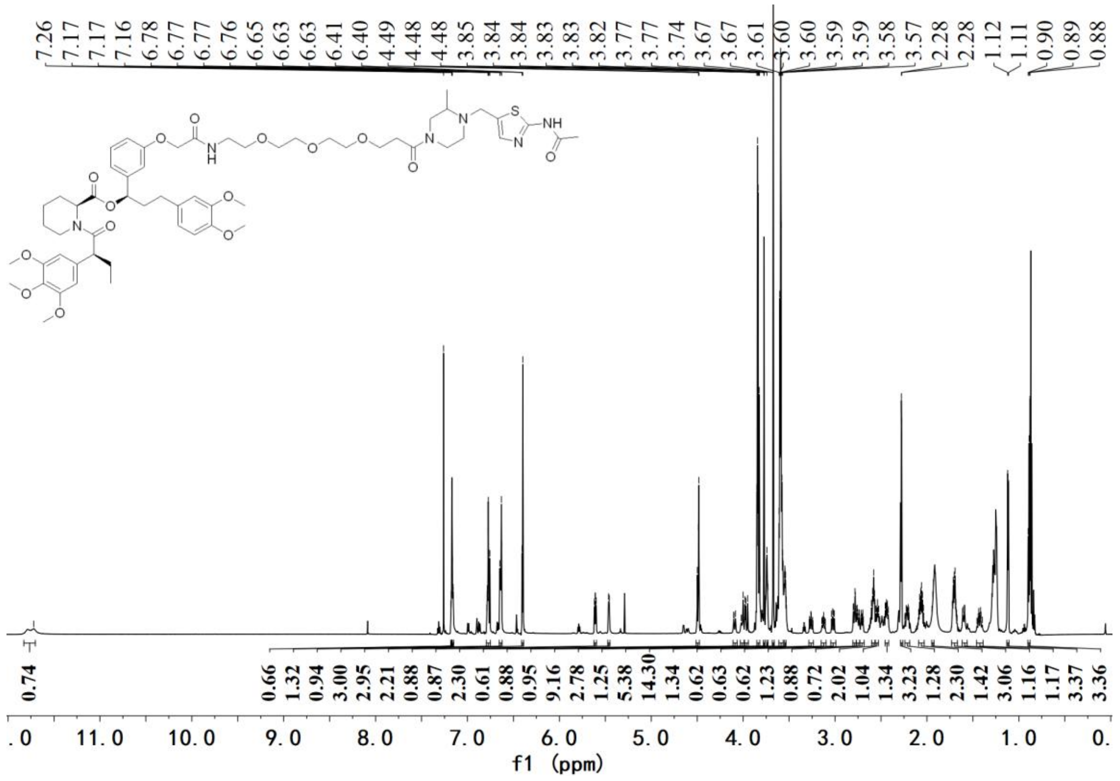

[1]H NMR spectrum of **D2**

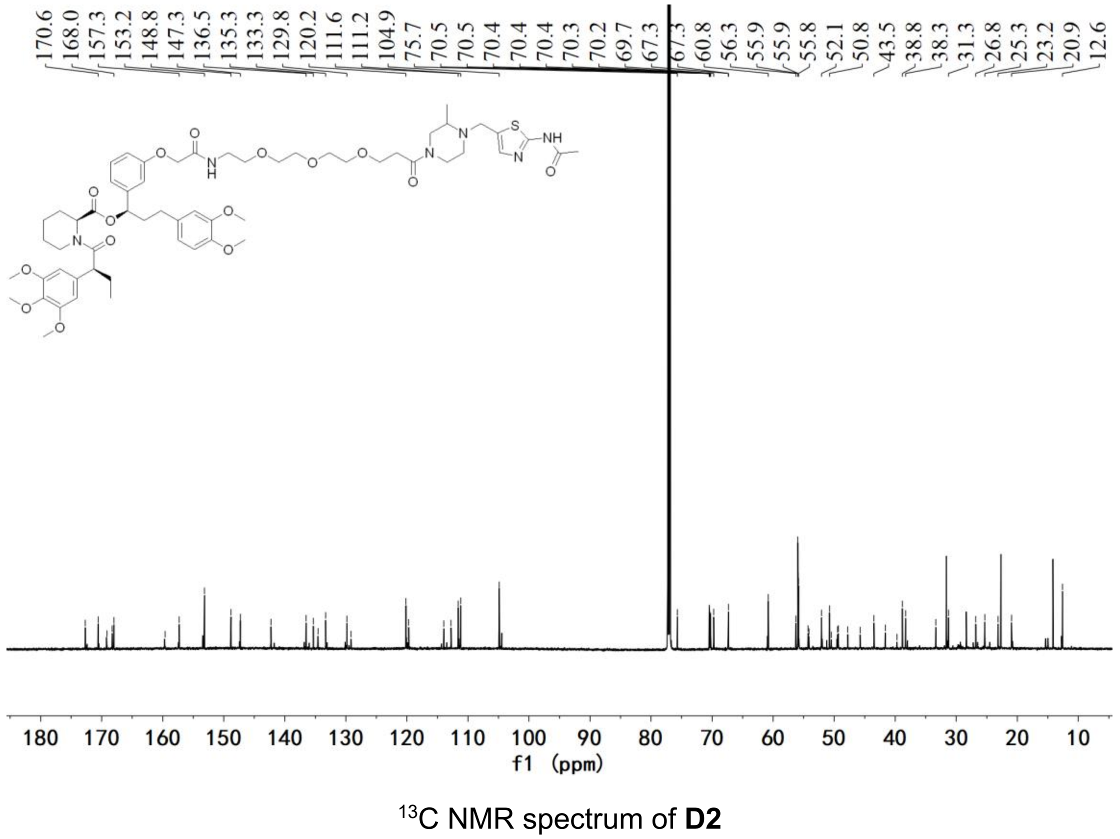

[13]C NMR spectrum of **D2**

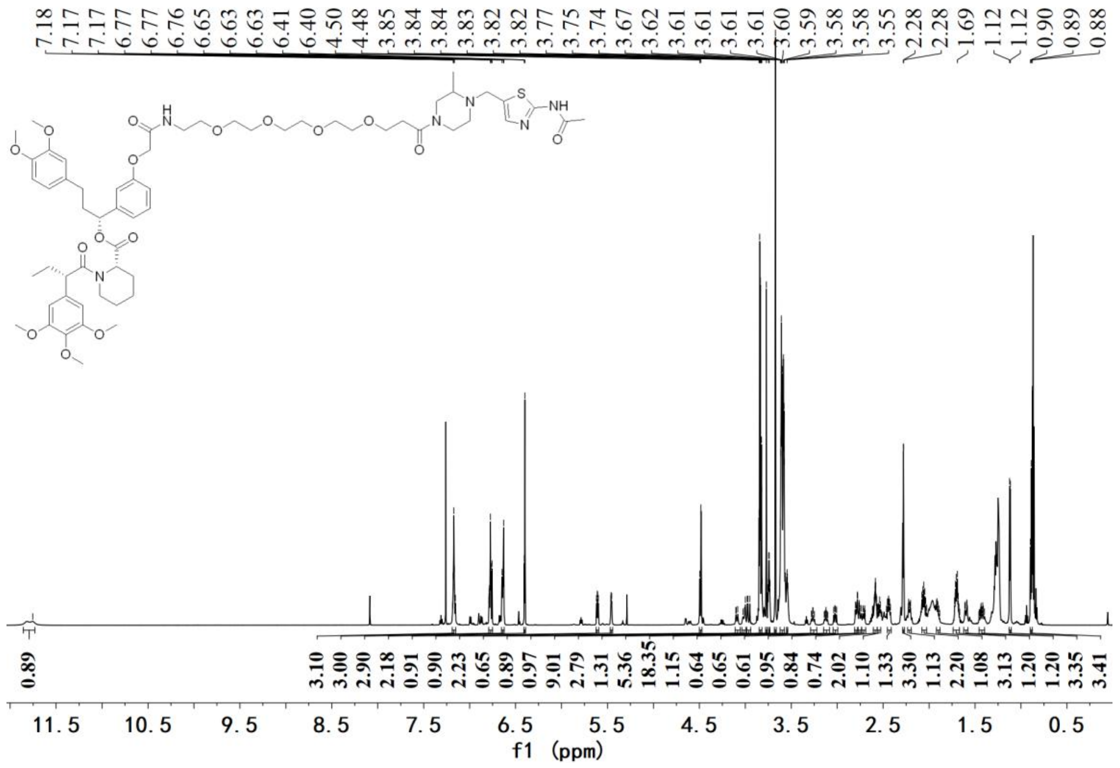

[1]H NMR spectrum of **D3**

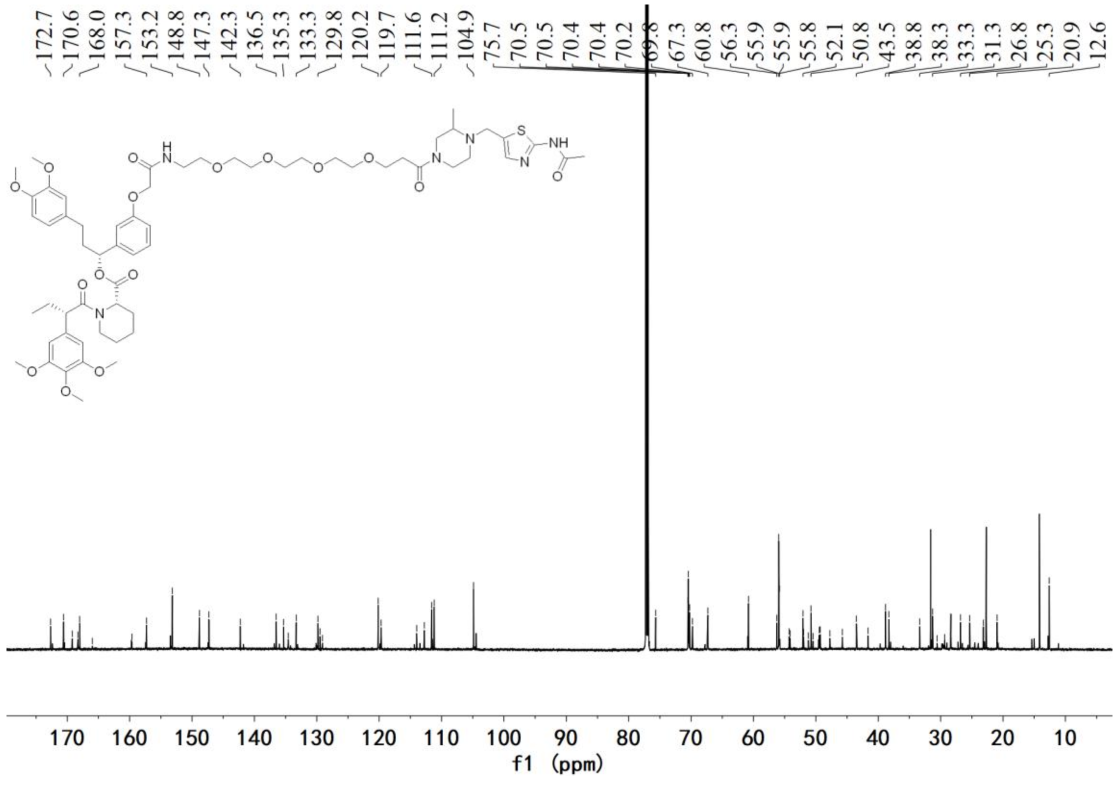

[13]C NMR spectrum of **D3**

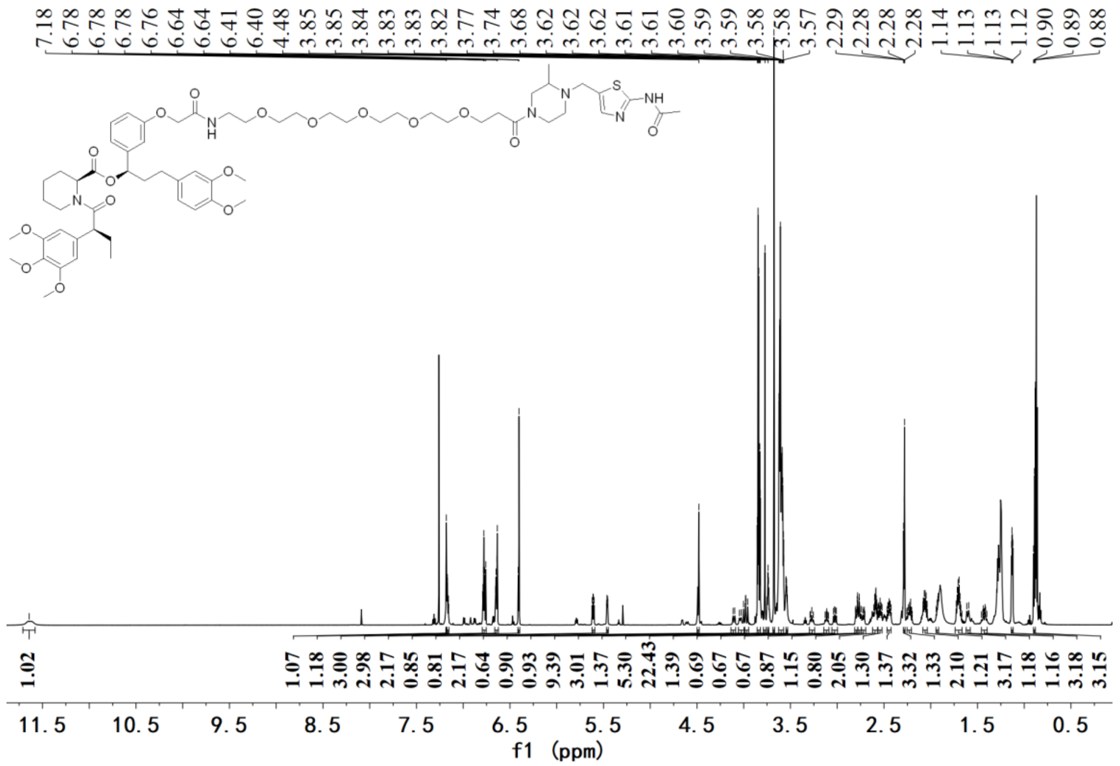

$^1H$ NMR spectrum of **D4**

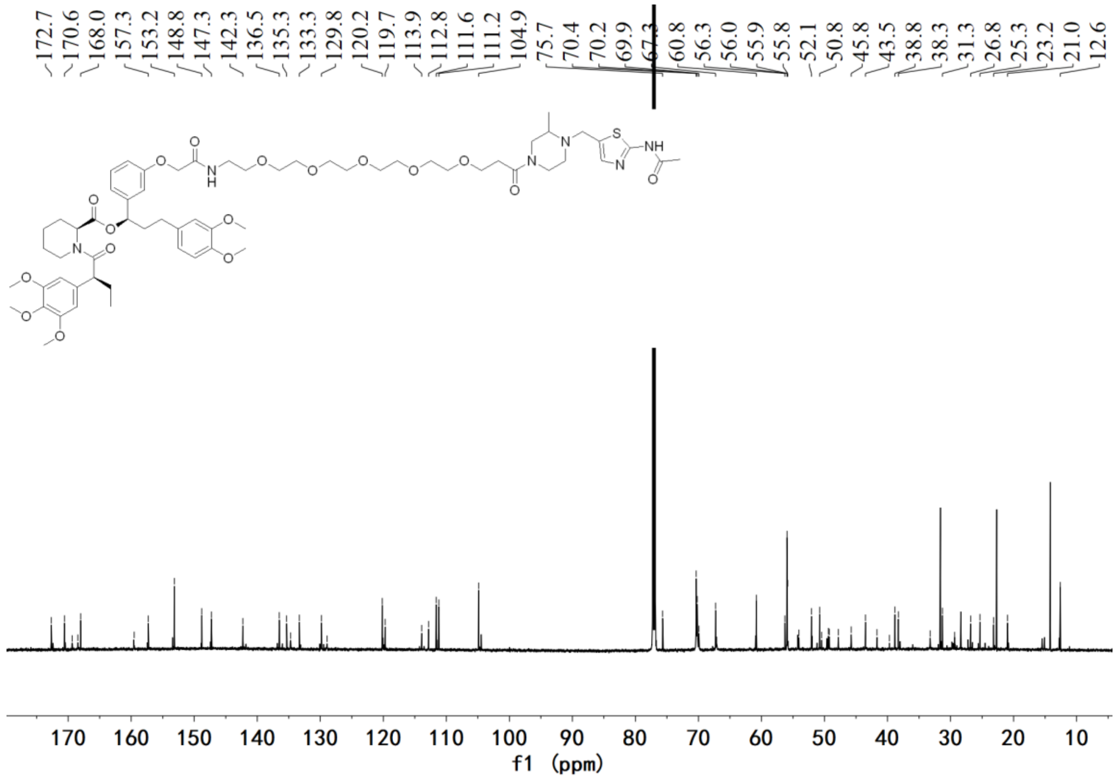

$^{13}C$ NMR spectrum of **D4**

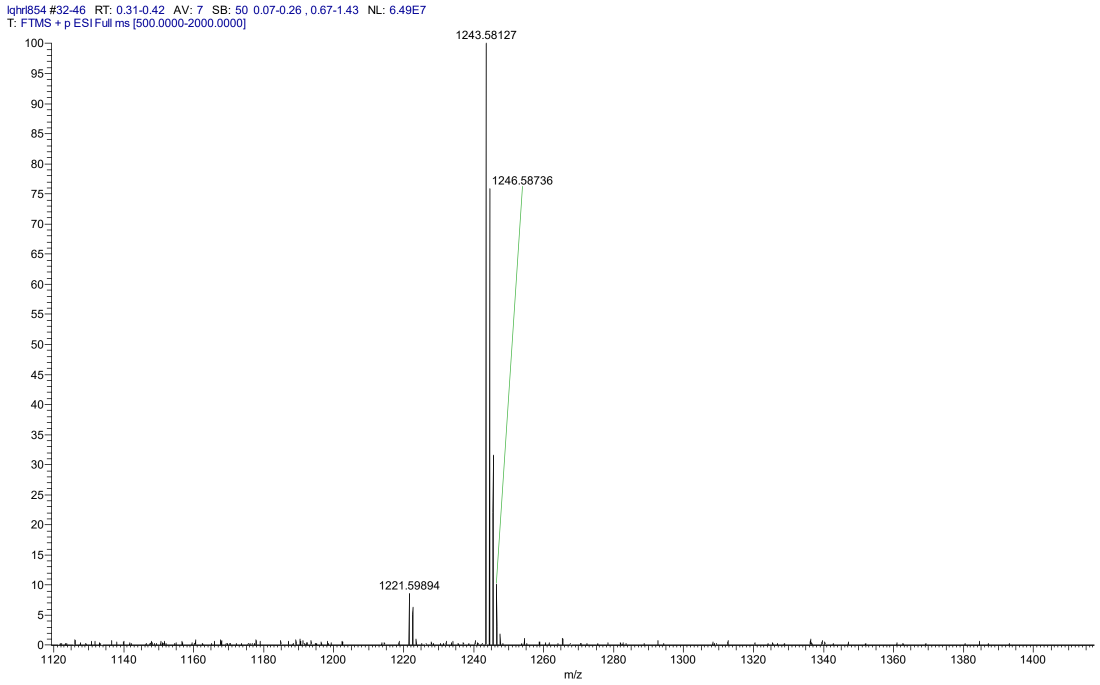


HRMS spectrum of **D4**

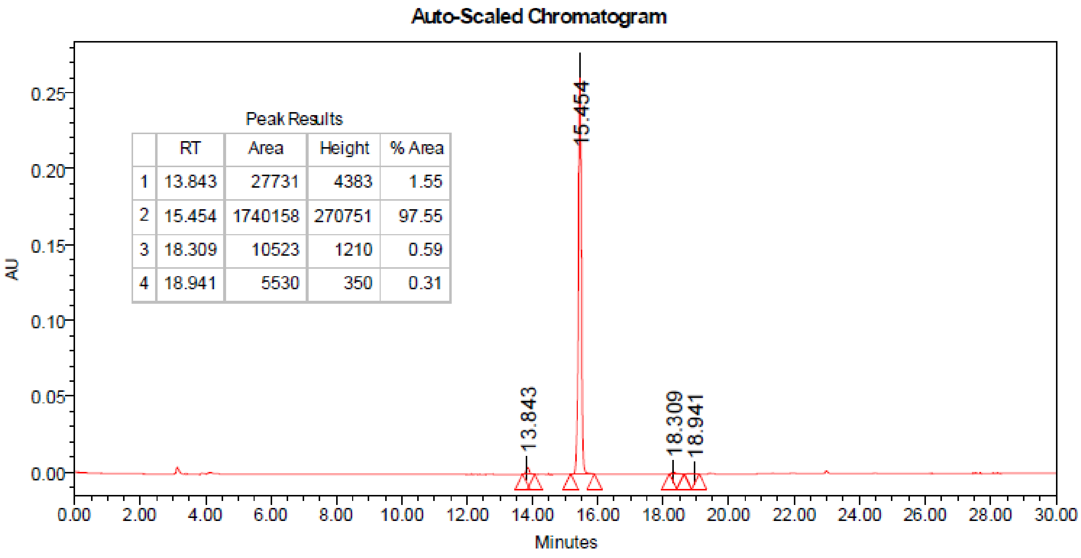


| | RT | Area | Height | % Area |
|---|---|---|---|---|
| 1 | 13.843 | 27731 | 4383 | 1.55 |
| 2 | 15.454 | 1740158 | 270751 | 97.55 |
| 3 | 18.309 | 10523 | 1210 | 0.59 |
| 4 | 18.941 | 5530 | 350 | 0.31 |

HPLC spectrum of **D4**

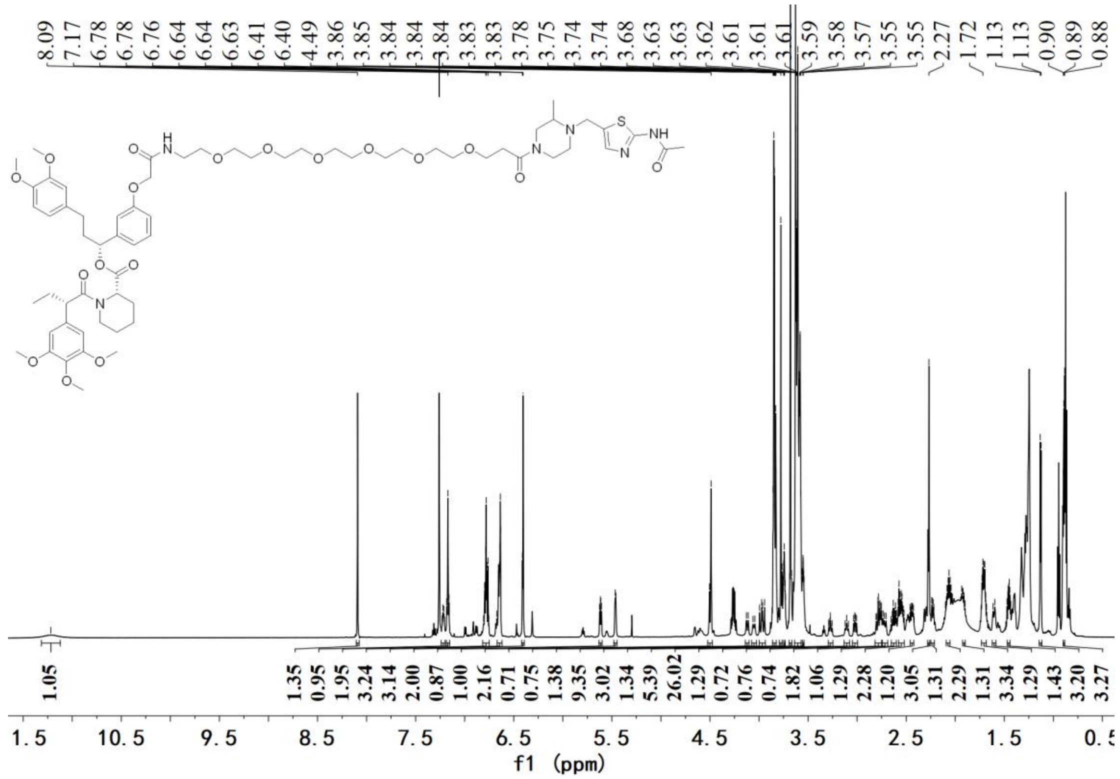

$^{1}H$ NMR spectrum of **D5**

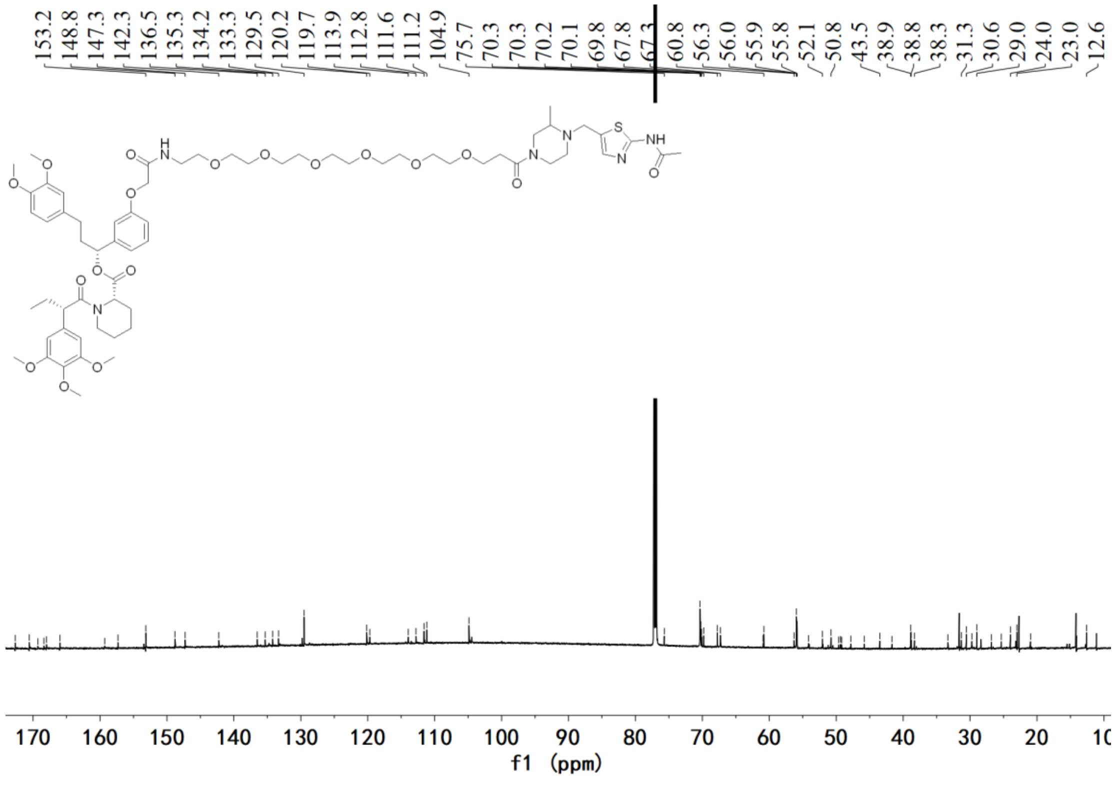

$^{13}C$ NMR spectrum of **D5**

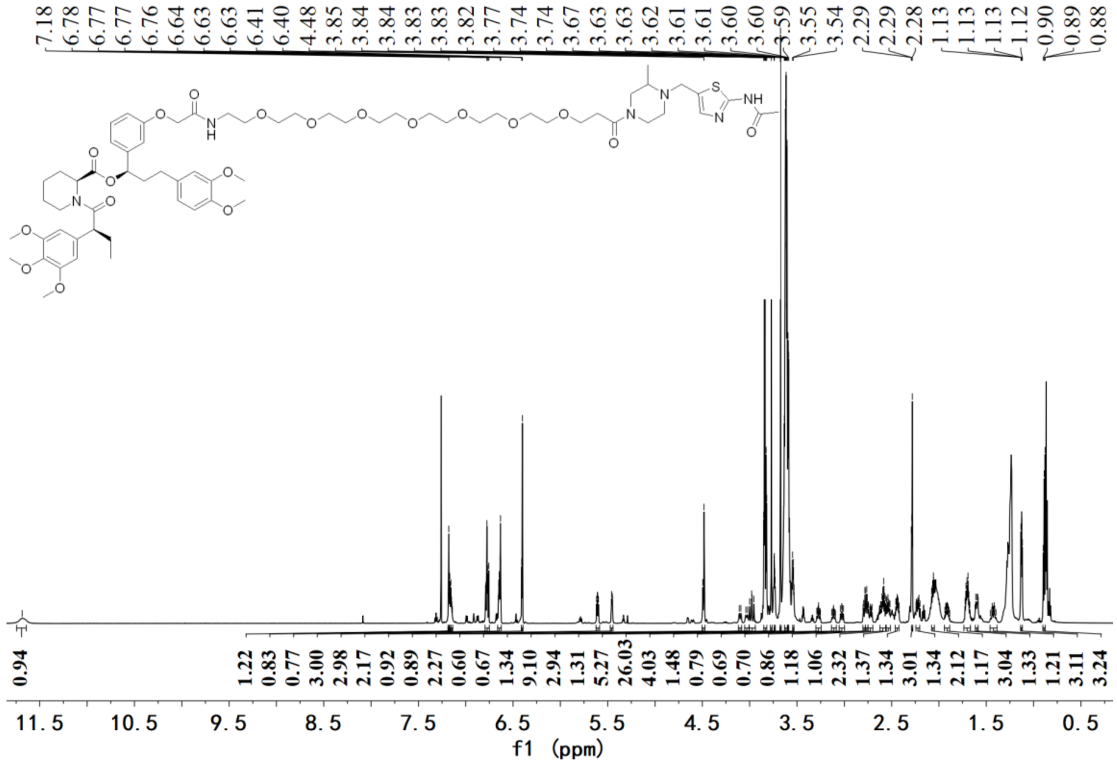


$^{1}H$ NMR spectrum of **D6**

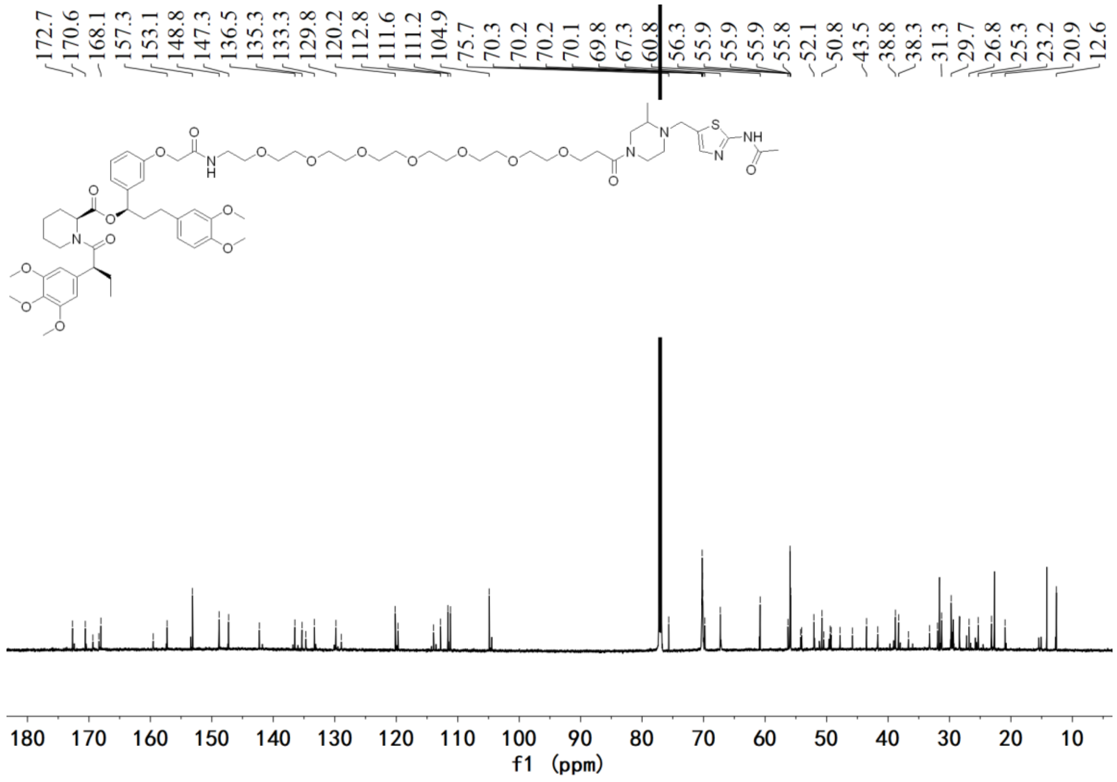


$^{13}C$ NMR spectrum of **D6**

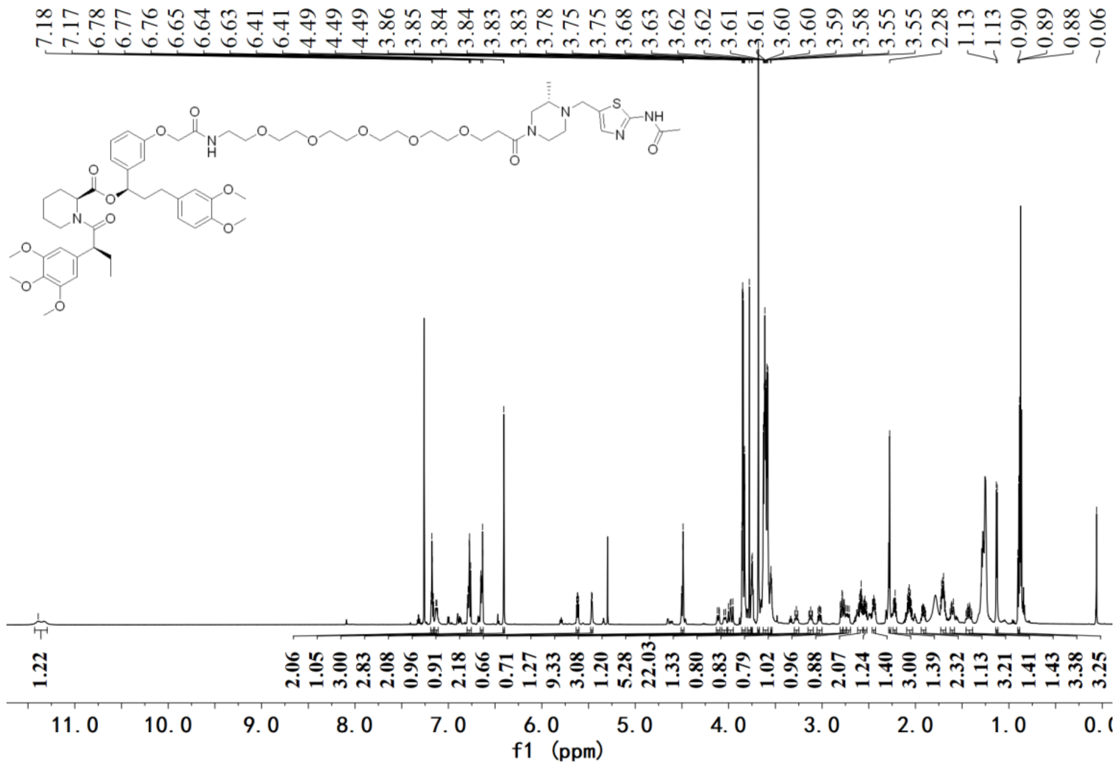

$^1$H NMR spectrum of **E1**

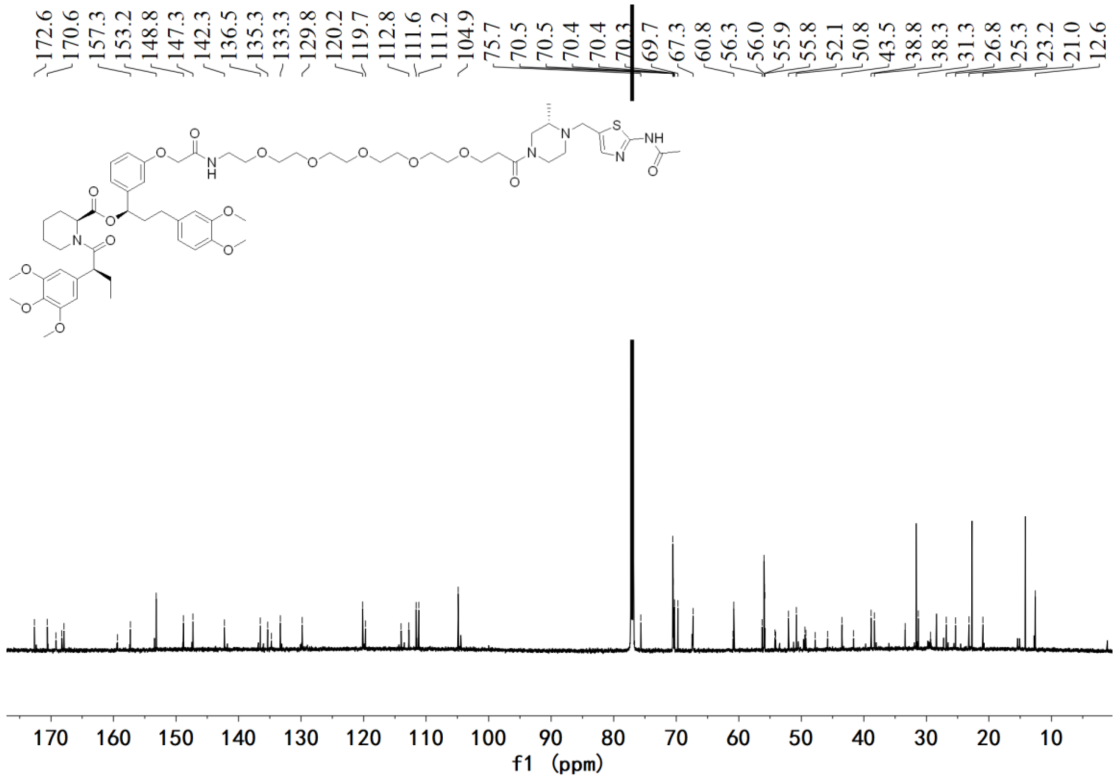

$^{13}$C NMR spectrum of **E1**

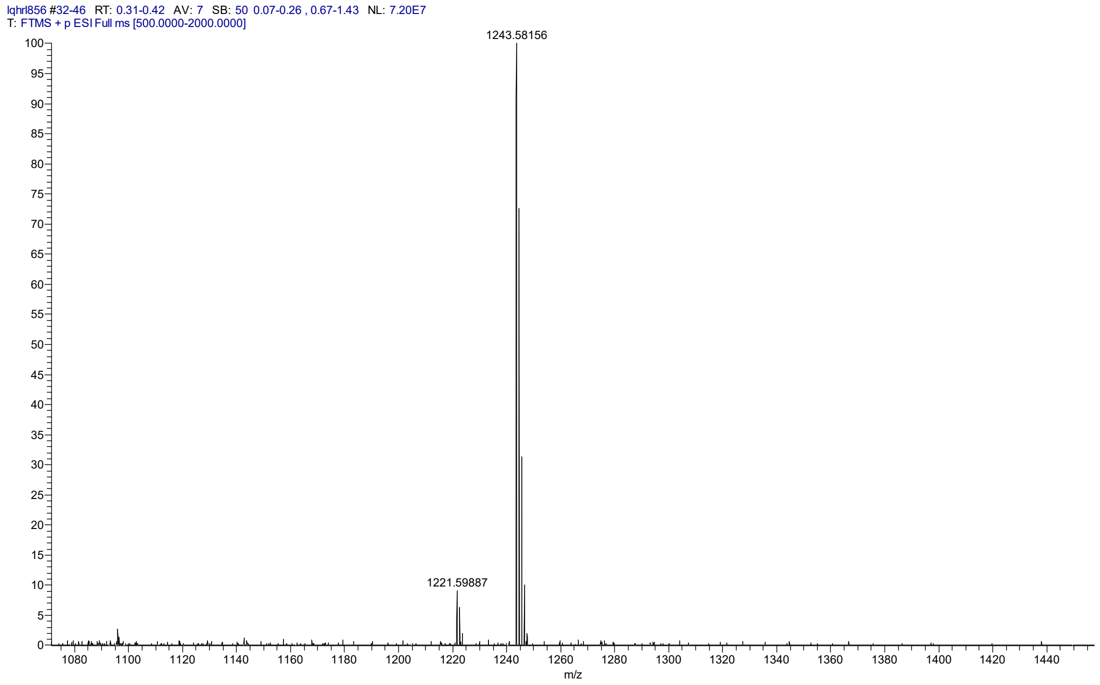


HRMS spectrum of **E1**

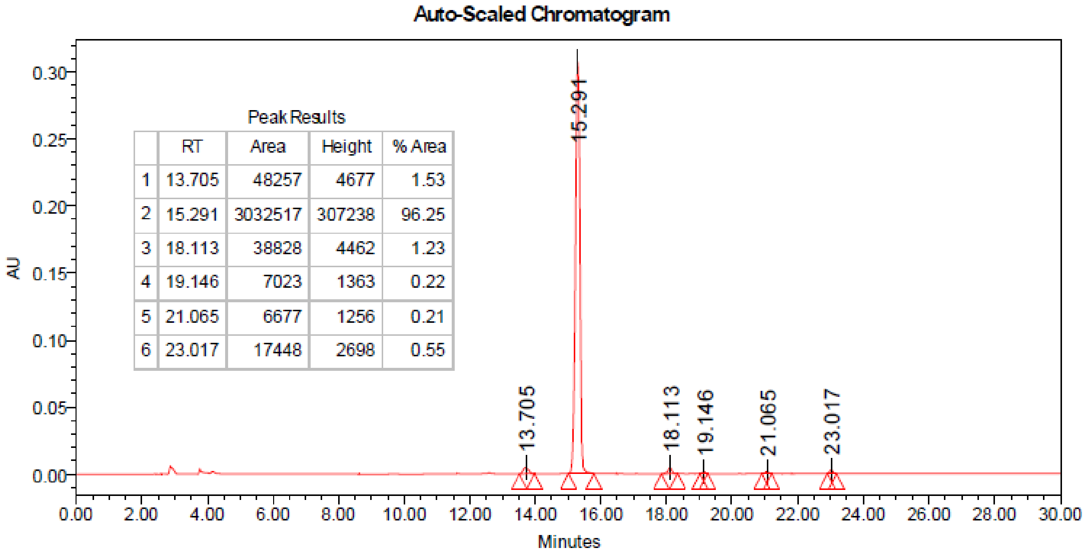


| | RT | Area | Height | % Area |
|---|---|---|---|---|
| 1 | 13.705 | 48257 | 4677 | 1.53 |
| 2 | 15.291 | 3032517 | 307238 | 96.25 |
| 3 | 18.113 | 38828 | 4462 | 1.23 |
| 4 | 19.146 | 7023 | 1363 | 0.22 |
| 5 | 21.065 | 6677 | 1256 | 0.21 |
| 6 | 23.017 | 17448 | 2698 | 0.55 |

HPLC spectrum of **E1**

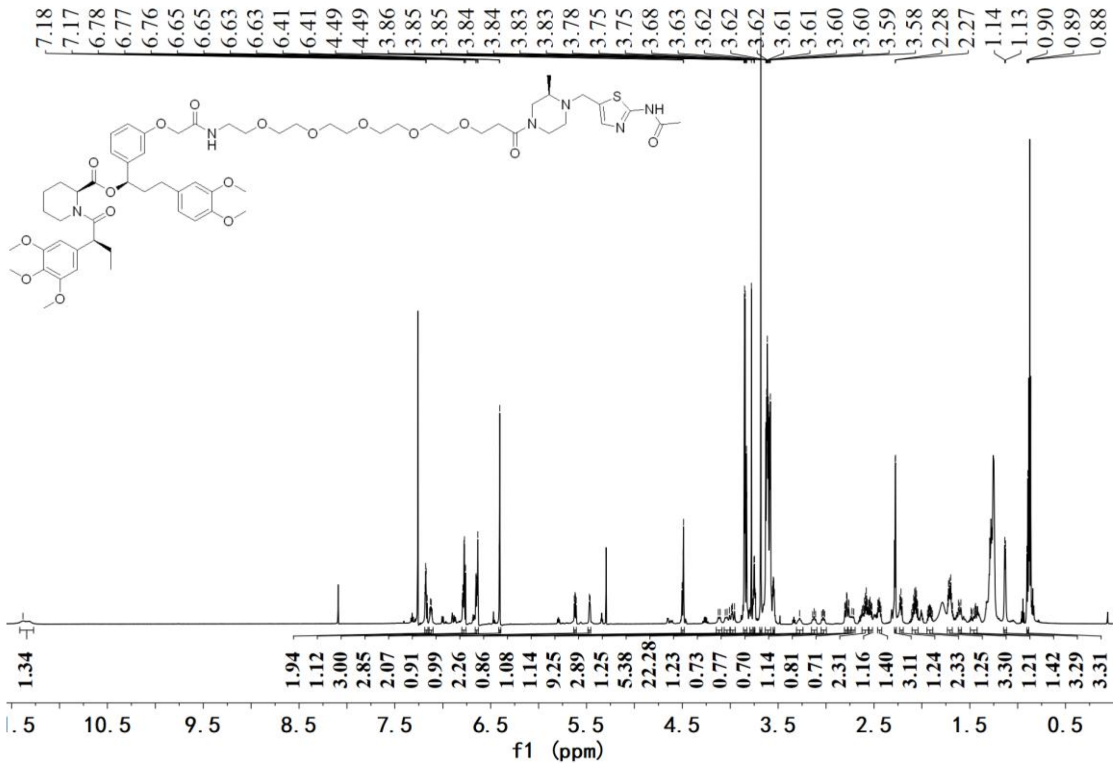

[1]H NMR spectrum of **E2**

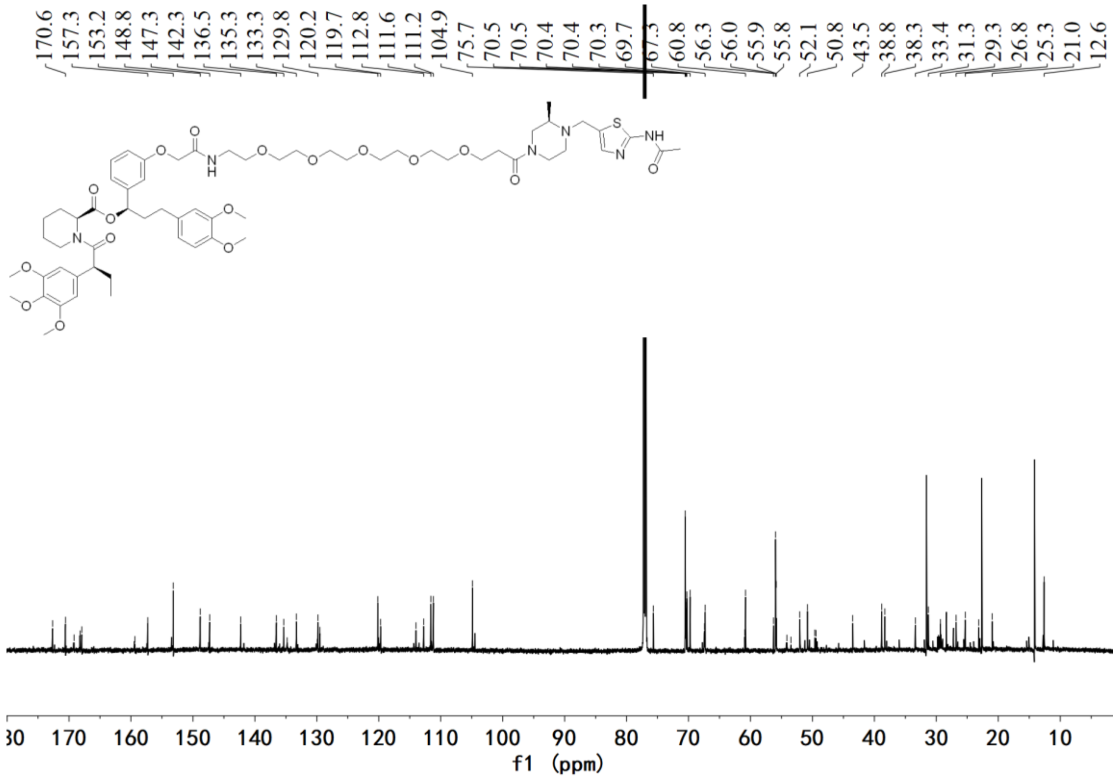

[13]C NMR spectrum of **E2**

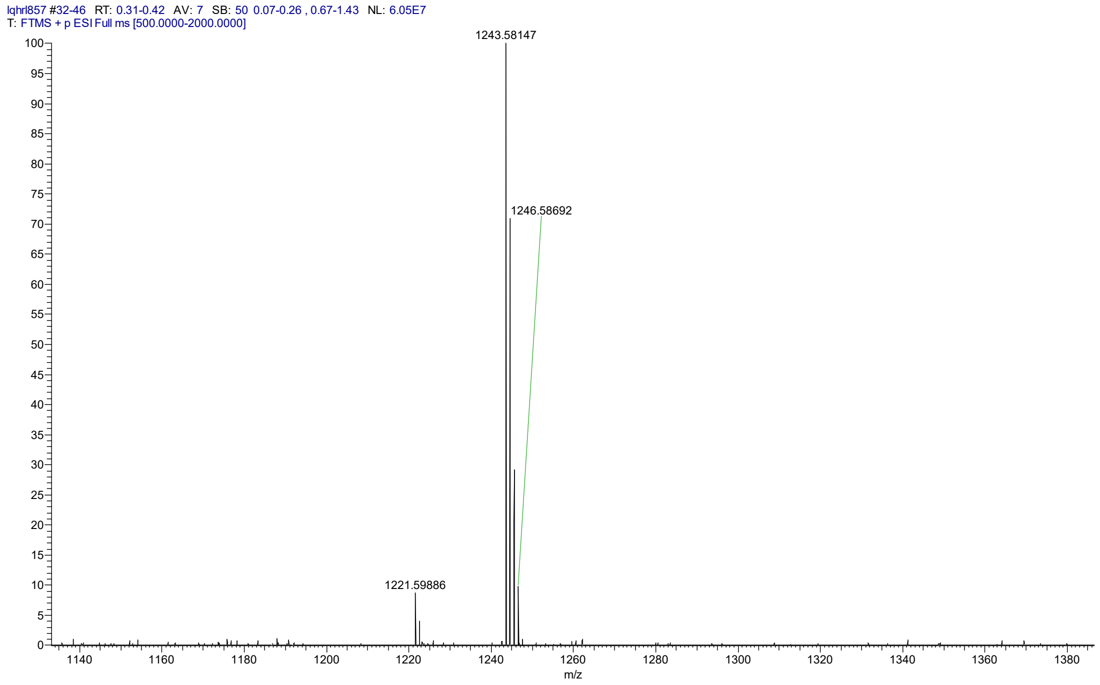


HRMS spectrum of **E2**

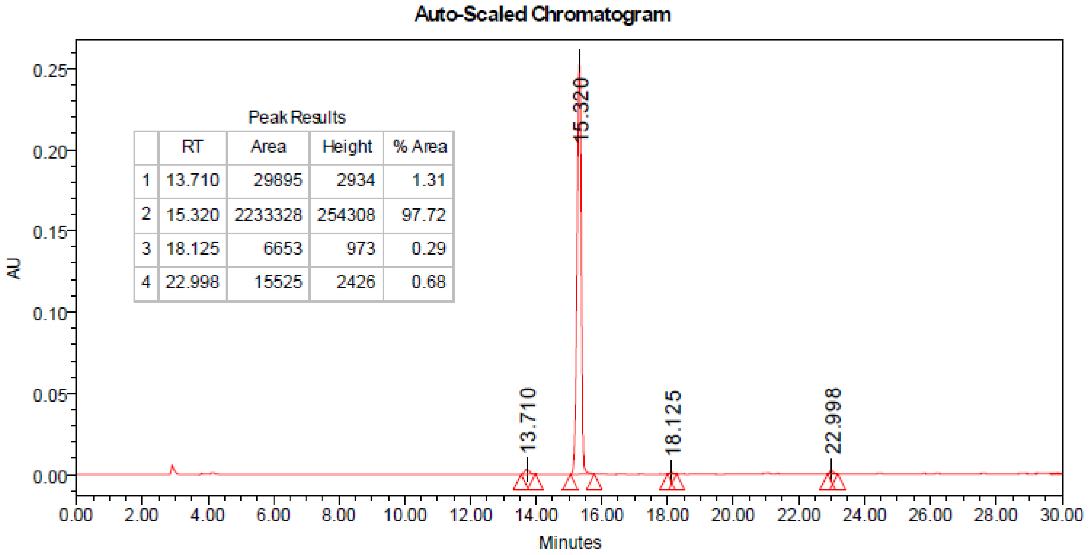


Peak Results

| | RT | Area | Height | % Area |
|---|---|---|---|---|
| 1 | 13.710 | 29895 | 2934 | 1.31 |
| 2 | 15.320 | 2233328 | 254308 | 97.72 |
| 3 | 18.125 | 6653 | 973 | 0.29 |
| 4 | 22.998 | 15525 | 2426 | 0.68 |

HPLC spectrum of **E2**
